# Phylogenomic challenges in polyploid-rich lineages: Insights from paralog processing and reticulation methods using the complex genus *Packera* (Asteraceae: Senecioneae)

**DOI:** 10.1101/2025.09.14.676143

**Authors:** Erika R. Moore-Pollard, Paige A. Ellestad, Jennifer R. Mandel

## Abstract

Phylogenomic discordance is pervasive and cannot always be resolved by increasing the amount of sequencing data alone. Biological processes such as polyploidy, hybridization, and incomplete lineage sorting are major contributors to discordance and must be accounted for to avoid misleading evolutionary interpretations. To better understand how these processes influence phylogenetic reconstruction, we conducted a comprehensive phylogenomic study in the complex genus *Packera*. With over 90 species and varieties, 40% of which exhibit polyploidy, aneuploidy, or other cytological complexities, *Packera* presents significant challenges for phylogenetic reconstruction. Given these complexities, we assessed different published paralog processing methods on the resulting evolutionary relationships and phylogenetic support of this group. We then applied three of these methods to evaluate their impact on tree topology and our understanding of *Packera*’s evolutionary history by constructing a time-calibrated phylogeny, reconstructing historical biogeography, and testing for ancient reticulation. Phylogenetic outcomes varied based on the paralog processing method used, with no method performing the best over others. Our findings highlight the large impact of orthology inference and paralog processing on phylogenomic analyses, particularly in polyploid-rich groups such as *Packera*, and we offer guidance on methodological impacts along with practical recommendations. We note that gaining a robust understanding of *Packera*’s evolutionary history requires more than computational approaches alone. While technological advancements have greatly expanded our ability to analyze genomic data, effective phylogenomic research still relies on strong taxon sampling and detailed species knowledge. Without careful attention to the biological context, such as reproductive boundaries, cytological variation, ecological interactions, and historical biogeographic processes, phylogenomic studies risk misinterpreting evolutionary history and processes. By accounting for these factors, we can begin to improve the accuracy of evolutionary reconstructions and gain deeper insights into the complex history of plant diversification.

## Introduction

The field of phylogenetics is drastically changing and now routinely incorporates multi-locus and genomic data. While this expansion has enabled deeper insights into evolutionary relationships across the tree of life, it has also revealed persistent and complex challenges in phylogenetic inference. One significant challenge is gene tree discordance, where phylogenetic trees estimated from individual loci disagree with each other and/or with the inferred species tree (Degnan and Rosenberg 2009; Mendes et al. 2016). These incongruences are not simply a consequence of limited data; rather, they often reflect differences driven by underlying biological processes such as incomplete lineage sorting (ILS), gene duplication and loss, and hybridization or introgression (Pease et al. 2018). Such processes are especially prevalent in plants, where whole-genome duplications (WGDs), ancient hybridization, and reticulate evolution are widespread and recurrent (Funk 1985; Vision et al. 2000; Wendel and Cronn 2003; Peirson et al. 2012; Panchy et al. 2016; Rose et al. 2021).

Polyploidy, or WGD, is a dominant evolutionary force in angiosperms, with estimates suggesting that up to 70% of extant flowering plants have polyploid ancestry (Grant 1971; Masterson 1994; Soltis and Soltis 1999; One Thousand Plant Transcriptomes Initiative 2019). These events gave rise to extensive paralogy in plant genomes, which poses significant challenges for orthology inference and phylogenetic analysis. Although these issues are particularly pronounced in plants (e.g., Asparagaceae, Bentz and Leebens-Mack 2024; Rosaceae, Morales-Briones et al. 2018, 2021a; and Amaranthaceae, Morales-Briones et al. 2021b), they are by no means exclusive to them. Similar complications have been documented in non-plants, such as African clawed frogs (Evans et al. 2004), desert rodents (Gallardo et al. 2004), alpine butterflies (Nice et al. 2013), and fish (Braasch and Postlethwait 2012), indicating that complex histories of gene duplication and horizontal gene transfer create phylogenetic uncertainty across the tree of life. As the scale and scope of phylogenomic datasets expand, so too does the need to carefully model and interpret the biological processes underlying gene tree discordance. Fortunately, the growing availability of phylogenomic data now makes it possible to start untangling these complex histories in ways that were not previously feasible; however, doing so requires tools and models that explicitly account for these processes.

In response, several bioinformatic tools and pipelines have been developed to infer orthology under increasingly realistic evolutionary models. For instance, softwares such as HybPiper (Johnson et al. 2016) and Phyluce (Faircloth 2016) implement distinct strategies for identifying and handling putative paralogs, with some pipelines retaining only a single gene copy (e.g., HybPiper), while others exclude all loci which exhibit signs of paralogy (e.g., Phyluce). Although Phyluce has shown greater success in some groups (e.g., *Antennaria* Gaertn.; Thapa et al. 2020), HybPiper typically outperforms Phyluce across most plant lineages because of its ability to better resolve issues associated with polyploidy.

Additional pipelines have been developed that implement distinct strategies for identifying and handling putative paralogs using graph- or tree-based methods (e.g., paragone-nf; Yang and Smith 2014; Jackson et al. 2023), whereas others consider paralogy during species tree estimation itself (e.g., ASTRAL-Pro; Zhang et al. 2020; Zhang and Mirarab 2022a). These different approaches can yield markedly different phylogenetic outcomes, particularly in polyploid-rich systems where paralogs may be the rule rather than the exception. Yet, few studies have systematically compared these methods to understand their impact on phylogenetic and downstream evolutionary inferences.

This is especially relevant for groups such as *Packera* Á. Löve & D. Löve (Asteraceae: Senecioneae), a North American genus of about 91 species known for polyploidy, complex patterns of hybridization, and poorly resolved taxonomy. An estimated 40% of *Packera* taxa present polyploidy, aneuploidy, and/or other cytological disturbances (Barkley 1988; Trock 2006), complicating phylogenetic reconstruction of this group (Bain and Golden 2000). Given this, *Packera* provides an ideal system for testing how alternative strategies for handling paralogs influence phylogenomic inference, owing to its extensive polyploidy, ecological and geographic diversity, and history of reticulation; all factors that increase the potential for gene tree discordance.

Here we present the most comprehensive phylogenomic investigation of *Packera* to date, sampling nearly 93% of the genus and leveraging a high-resolution Hyb-Seq dataset. We applied a robust framework for evaluating the impact of gene duplication by comparing multiple paralog processing methods, including pipelines that remove all paralogs, those that model duplication and loss, and those that retain a single representative gene copy. We assessed how these approaches influence gene tree concordance, species tree topology, and node support. We then tested how three of these methods (a standard paralog processing method, a method that removes all paralogs, and a selective paralog-aware method; see methods below) alter the interpretations of downstream macroevolutionary analyses including divergence dating, historical biogeography, and detection of ancient introgression. Our results demonstrate that the choice of paralog processing approach strongly influences inferred species relationships, gene tree concordance, node support, and evolutionary inferences, highlighting the profound effect that paralog-aware decisions can have in phylogenomics. Additionally, this study provides a broadly applicable framework for assessing paralogs and their downstream impacts on phylogenetic inference in systems shaped by gene duplication, reticulation, and rapid diversification.

## Materials & Methods

### Specimen Collection

A nearly complete sampling of currently recognized *Packera* species and varieties were used in this study (62 *Packera* species, 22 varieties, and one nothospecies; totaling 85 *Packera* taxa). *Packera* species were selected based on information published in the International Plant Names Index (IPNI; available at http://www.ipni.org) and then curated from discussions with botanists that have previously specialized in *Packera* (D. Trock & J. Bain, pers. comm.). Outgroup taxa from tribes Senecioneae (n = 19) and Anthemideae (n = 4) were selected by consulting recent studies of the group (Bain and Golden 2000; Pelser et al. 2010; Mandel et al. 2019). Additionally, three taxa from tribe Calenduleae were included for downstream analyses that require more distantly related outgroups that do not have a known WGD event (Barker et al. 2016; Huang et al. 2016; Zhang et al. 2021). A complete list of the 111 sampled species, herbarium vouchers, and their publication status can be found in Supplementary Table 1.

### DNA Extraction and Sequencing

Leaf tissues were obtained from herbarium specimens or silica-dried field collections. Dried leaf tissue (∼25 mg) was ground using a Bead Mill 24 Homogenizer (ThermoFisher Scientific, Atlanta, Georgia, USA), and total DNA was extracted using the Omega Bio-Tek Kit (Atlanta, Georgia, USA) following manufacturer’s protocols with the addition of polyvinylpyrrolidone (PVP) and ascorbic acid to the first extraction buffer (10 mL SQ1 buffer, 100 mg PVP, 90 mg ascorbic acid per reaction). Samples of *Packera* and outgroup species that were obtained from other researchers followed different DNA extraction protocols which are described in Supplementary Table 1. All DNA samples used in this study were quality checked using a Nanodrop 2000 (Thermo Fisher Scientific, Carlsbad, California, USA) and quantified using a Qubit High Sensitivity assay (ThermoFisher Scientific, Oregon, USA). If the DNA had low concentrations or possible contaminants, the samples were cleaned following the E.Z.N.A. Cycle Pure Kit (Omega Biotek, Georgia, USA) and re-quantified with the Qubit High Sensitivity (HS) assay. Up to 1 µg DNA per sample was sonicated with a QSonica machine (ThermoCube, New York, USA) to generate fragment sizes within the 400-500 base pair (bp) range. The amount of DNA used was dependent on the initial total DNA concentration of the targeted sample. Time allotted for sonication of each sample varied depending on approximate length and degree of fragmentation of the DNA. Prior to sonication, total DNA was visualized on a 1% agarose gel in 1X TBE and GelRed 3x (Biotium) and the approximate length was estimated by comparing to a 100 bp GeneRuler DNA ladder (ThermoFisher Scientific, Vilnius, Lithuania). If the DNA sample appeared smaller than 400 bp, it was not sonicated.

Libraries for each sample were prepared using the NEBNext Ultra II DNA Library Prep and NEBNext Multiplex Oligos for Illumina Kits (New England Biolabs, Ipswich, Massachusetts, USA) following the NEBNext Ultra II Version 5 protocol with size selection on DNA fragments at 300-400 bp range. The protocol was adjusted by halving all amounts of reagents and DNA used. The barcoded libraries (NEBNext multiplex oligos) were quality checked with an Agilent Bioanalyzer (Agilent Technologies, Santa Clara, California, USA) and quantified by Qubit HS assay. Libraries were then used for targeted sequence capture with the custom MyBaits probe set, Compositae-1061 (Mandel et al. 2014), from Arbor Biosciences (Ann Arbor, Michigan, USA) following the manufacturer’s protocol (v. 2.3.1). Approximately 500 ng of total DNA library in 6 µl of library, or equal amount of DNA per sample when pooling (maximum of 8 samples), was used in each MyBaits reaction for hybridization with the baits for 36 hours. Hybridized targets were recovered using Dynabeads M-280 Streptavidin (ThermoFisher Scientific, Vilnius, Lithuania) following the manufacturer’s protocol. Captured targets were amplified and quantified using KAPA library quantification kits (Kapa Biosystems, Wilmington, Massachusetts, USA). A 1:4 ratio of library was added to each of the captured target pools to obtain off-target plastid sequence data. Libraries were sequenced on Illumina MiSeq (300 cycles) and NovaSeq 6000 (300 cycles) instruments at Psomagen (Rockville, Maryland, USA) across several runs. Sequence stats were obtained using the pxlssq function in Phyx v. 1.3.1 (Supplementary Table 2) (Brown et al. 2017).

### Sequence Processing

Raw FASTQ data were trimmed of low quality bases and adapter contamination using Trimmomatic v. 0.36 (Bolger et al. 2014). The Sliding Window quality filter was used in this analysis (illuminaclip 2:30:10, leading 20, trailing 20, sliding window 5:20). Cleaned reads with a minimum length of 36 bp and corresponding forward and reverse pairs were retained. FastQC v. 0.11.9 (Andrews 2010) was used to ensure good sequence quality.

### Plastid Assembly

Chloroplast sequences were extracted from trimmed reads using two approaches. First, the GetOrganelle v. 1.7.7.1 toolkit (Jin et al. 2020) was used to assemble circularized plastomes for all sequences with sufficient off-target sequencing data. The recommended parameters for embryophyte plant plastomes were used, except that the number of rounds and the disentangle time limit were increased (-R 75 and --disentangle-time-limit 7200). All complete circular plastomes were annotated using GeSeq (Tillich et al. 2017). From these, a complete *Packera* representative plastome was chosen and used as a reference. Alternatively, all reads were mapped to the reference plastome using Bowtie2 (Langmead and Salzberg 2012), variants were called using SAMtools v. 1.18 and BCFtools v. 1.18 (Danecek et al. 2021), and consensus sequences were created using seqtk v. 1.4 (https://github.com/lh3/seqtk). For the aligned consensus sequences, model selection and maximum likelihood phylogenetic analysis were then conducted using IQ-TREE v. 2.2.2 (Minh et al. 2020) and visualized using ‘ggtree’ (Yu et al. 2017) in R v. 4.3.1 (R Core Team 2016; RStudio 2020).

### Nuclear Ortholog Assembly

Cleaned FASTQ data were assembled using the HybPiper v. 2.1.6 (Johnson et al. 2016) pipeline to match the reads to the target loci contained in the Compositae-1061 probe set (Mandel et al. 2014). In HybPiper, the command ‘assemble’ was used to map the paired-end reads to the target loci using BWA v. 0.7.17 (Li and Durbin 2009. These were then assembled into contigs with specified kmer lengths (21, 33, 55, 77, 99) using SPAdes v. 3.5 (Bankevich et al. 2012). Paralog detection was carried out using the ‘paralog_retriever’ command (https://github.com/mossmatters/HybPiper/wiki/Paralogs), which flags loci that indicate potential paralogs, then extracts the assemblies that were flagged. A supercontig multifasta for each locus was generated and aligned in MAFFT v. 7.407 (Katoh and Standley 2013) using default parameters.

### Phylogenetic Analyses

Given the complex history of hybridization and polyploidization events in *Packera*, along with the large number of paralogs flagged from HybPiper in the current study (Supplementary Table 3), we sought to compare how different published orthology inference methods and species tree estimation tools impacted the inferred evolutionary relationships and support of this group. Our dataset is uniquely adapted to tackle this issue given dense taxon sampling and high locus recovery across both orthologs and paralogs, combined with decades of natural history and systematics work in the group, enabling more thorough investigations into these challenges.

We generated multiple nuclear phylogenies utilizing different paralog processing tools to infer orthology and species tree estimation: Concatenation (individual gene histories not accounted for), HybPiper with ASTRAL-III (hereafter ‘ASTRAL’; putative single-copy orthologous loci are chosen based on coverage depth and sequence similarity concerning the gene references from Compositae-1061, then inferred using coalescent methods), HybPiper with wASTRAL (hereafter ‘wASTRAL’; similar to ASTRAL, but low support gene tree branches are contracted prior to species tree estimation), HybPiper with ASTRAL-Pro (hereafter ‘ASTRAL-Pro’; all single-copy and multi-copy genes identified by HybPiper are considered, and the species tree that optimizes a quartet similarity measure is chosen), and the ‘one-to-one orthologs’ (1-to-1), ‘maximum inclusion’ (MI), ‘monophyletic outgroup’ (MO), and ‘rooted ingroup’ (RT) paralog processing methods by Yang and Smith (2014) with species trees inferred using ASTRAL-III (further details below; refer to Supplementary Table 3, Supplementary Fig. 1). By performing comparisons among methods, we aimed to gain a better understanding of the impact of paralogy on phylogenomic studies, and ultimately increase phylogenomic support and concordance across species of complex lineages.

#### Concatenation

Aligned supercontigs from MAFFT were concatenated with FASconCAT-G v. 1.02 (Kück and Longo 2014), then implemented in PartitionFinder v. 2.1.1 (Lanfear et al. 2012) to determine the most appropriate nucleotide substitution model (GTR, GTR+Γ, GTR+I+Γ). The overall best fitting model (Supplementary Table 4) was used to build a concatenated nuclear maximum likelihood tree in RAxML v. 8.1.3 (Stamatakis 2014) with 1,000 bootstrap (BS) replicates.

#### ASTRAL and wASTRAL

For the ASTRAL and wASTRAL analyses, nuclear gene alignments were separated based on the most suitable nucleotide substitution model for each gene as determined by PartitionFinder (Supplementary Table 4), then individual gene trees were generated using RAxML. For the ASTRAL analysis, the gene trees were used in ASTRAL-III v. 5.7.3 (Zhang et al. 2018) to generate the species trees. For the wASTRAL analysis, or weighted ASTRAL, a species tree was generated using ‘astral-hybrid’ in the ASTER program (https://github.com/chaoszhang/ASTER).

This method has shown to be more accurate than typical ASTRAL methods since wASTRAL incorporates gene tree uncertainty by weighting quartets based on the confidence levels of the input gene trees. In doing so, wASTRAL reduces the influence of erroneous gene trees, leading to a more reliable species tree estimation (Zhang and Mirarab 2022b). Local posterior probability (LPP) values were estimated for branch support and indicate the probability that the branch is the true branch given the set of gene trees provided (Sayyari and Mirarab 2016).

#### ASTRAL-Pro

ASTRAL-Pro2 (ASTRAL for PaRalogs and Orthologs) v. 1.13.1.3 (Zhang et al. 2020; Zhang and Mirarab 2022a), hereafter referred to as ASTRAL-Pro, provides a quartet-based species-tree inference method that allows multi-copy gene trees to be used as input. Thus, all paralogous and orthologous sequences as assigned by HybPiper were used to generate the species tree instead of selecting a single ortholog to represent an individual gene’s history (e.g., ASTRAL method). By accommodating multi-copy gene trees, ASTRAL-Pro accounts for gene duplication and loss events, reducing biases that arise from treating paralogs as true orthologs. This approach is intended to improve species tree accuracy, particularly in datasets with extensive gene family expansions, by leveraging more information from complex gene histories. The ASTRAL-Pro pipeline is similar to ASTRAL, as described above, but differs by having the user generate separate alignments and conduct maximum likelihood analyses on the genes that are flagged as paralogs and single-copy contigs. Single-copy and paralogous gene sequences were independently aligned using MAFFT, and gene trees were generated under the best-fitting nucleotide substitution model from PartitionFinder (Supplementary Table 4) in RAxML with 1,000 BS replicates. The RAxML output files were then used as input for ASTRAL-Pro2 to generate the species tree with LPP values at each node.

#### 1-to-1, MI, MO, and RT methods

Paralog processing was then carried out using the tree-based orthology inference strategies originally proposed by Yang and Smith (2014) via the paragone-nf v. 1.1.0 pipeline (Jackson et al. 2023; https://github.com/chrisjackson-pellicle/paragone-nf). This pipeline investigates the loci flagged as paralogous from HybPiper and determines which loci can be retained or pruned given specified conditions (Yang and Smith 2014; Morales-Briones et al. 2021a). All orthology inference methods proposed by Yang and Smith (2014)–the ‘one-to-one orthologs’ (1-to-1), ‘maximum inclusion’ (MI), ‘monophyletic outgroup’ (MO), and ‘rooted ingroup’ (RT) methods–were tested. Briefly, the 1-to-1 approach is the simplest strategy in that it only retains homologs that have no duplications. The MI approach extracts the subtree with the highest number of taxa without taxon duplications. The MO approach filters for homolog trees where outgroup taxa (here tribe Calenduleae: *Calendula arvensis*, *Tripteris pinnatilobatum* (Norl.) B.Nord., and *Chyrsanthemoides incana* (Burm.f.) Norl.) are monophyletic and non-repeating. The RT approach searches subtrees of ingroup taxa and roots them by the designated outgroups, then removes the ingroup taxa as rooted trees. In turn, inferred gene duplications, or paralogs, are then pruned from root to tip in the rooted tree (Yang and Smith 2014; Morales-Briones et al. 2021a).

To aid in consistency across analyses, the HybPiper run described above was used as input instead of using the hybpiper-nf pipeline (Jackson et al. 2023). The three Calenduleae taxa were considered the internal outgroups since a recent whole genome duplication event has not been reported for this tribe, as is required for the MI and MO analyses (Yang and Smith 2014). The resulting trimmed alignments produced via paragone-nf were used as input for RAxML to generate gene trees, and a species tree with LPP support values was generated using ASTRAL-III following the ‘ASTRAL’ protocol described above.

### Measuring phylogenetic tree support

The normalized quartet score was calculated for all orthology inference methods that generated a species tree with ASTRAL-III using the ‘-q’ option. Branch lengths were recalculated from coalescent units to nucleotide substitutions per codon site for each ASTRAL-III species tree using IQ-TREE. Parsimony informative sites (PIS) and pairwise identity (PI) were calculated for each gene matrix using the ‘parsimony_informative_sites’ and ‘pairwise_identity’ functions, respectively, in phykit v. 2.0.1 (Steenwyk et al. 2021). Nuclear and plastome trees were visualized using the package ‘phytools’ (Revell 2012) in R v. 4.0.5.

### Measuring Discordance

Species trees from concatenation, plastid, ASTRAL, wASTRAL, and paragone-nf (1-to-1, MI, MO, and RT) were analyzed for underlying discordance using Quartet Sampling (Pease et al. 2018). ASTRAL-Pro was not analyzed as Quartet Sampling does not allow multi-copy gene trees as input. Quartet Sampling assesses the confidence, consistency, and informativeness of internal tree relationships, and the reliability of each terminal branch by evaluating if a phylogeny has a lack of support due to lack of information, discordance from ILS or introgression, or misplaced or erroneous taxa. Three scores were calculated for each internal branch of the focal tree which determines the consistency of information (Quartet Concordance [QC]), the presence of secondary evolutionary histories (Quartet Differential [QD]), and the amount of information (Quartet Informativeness [QI]). Quartet Fidelity (QF) scores were calculated at each tip to report how frequently a taxon is included in concordant topologies. Using the rooted species trees and concatenated gene alignments from FASconCAT-G (described above), Quartet Sampling was run using default settings on the designated analyses. The concatenation, plastid, ASTRAL, wASTRAL, 1-to-1, MI, and MO species trees were rooted to the outgroup Calenduleae taxon, *Tripteris pinnatilobatum*, while the RT tree was rooted to *Doronicum pardalianches* L. using the *pxrr* function in Phyx (Supplementary Table 3).

Additionally, PhyParts v. 0.0.1 (Smith et al. 2015) was used with default settings to quantify and visualize discordance in the ASTRAL, wASTRAL, and paragone-nf (1-to-1, MI, MO, and RT) trees. Similar to Quartet Sampling, ASTRAL-Pro could not be analyzed because of multi-copy gene trees. Additionally, the concatenation and plastid trees could not be analyzed since those analyses do not use individual gene histories as input. Resulting trees containing pie charts indicating whether gene trees were supported, conflicted, or uninformative at each node were generated using a custom python script (https://github.com/mossmatters/phyloscripts/tree/master/phypartspiecharts). PhyParts requires rooted gene trees and a rooted species tree as input. Similar to Quartet Sampling, ASTRAL, 1-to-1, MI, and MO gene trees were rooted to *Tripteris pinnatilobatum* and the RT tree was rooted to *Doronicum pardalianches* using the *pxrr* function in Phyx. Rooting gene trees for PhyParts decreased the number of genes used as input since not all gene trees contained the targeted outgroup taxon (see Supplementary Table 3). For phylogenies generated with ASTRAL technologies– ASTRAL, wASTRAL, ASTRAL-Pro, and paragone-nf trees (Supplementary Table 3)– introgression and ILS were visualized across lineages using phytop v. 0.3.1 (Shang et al. 2025) with default settings. The species trees were re-ran with the flag ‘-t 2’ added to be used as input for phytop.

### Tree Topology Comparison

To evaluate similarity among the nuclear gene trees, pairwise adjusted Robinson-Foulds (RF_adj_) distances were calculated as outlined by (Mitchell et al. 2017) between all trees using the *RF.dist()* function in package ‘phangorn’ (Schliep 2011) in R, with unrooted trees as input and setting the “normalize” argument to TRUE (Steel and Penny 1993). The Calenduleae outgroup taxa were trimmed from the trees in R using the *drop.tip()* function in package ‘ape’ (Paradis and Schliep 2019) to make concatenation, ASTRAL, ASTRAL-Pro, and paragone-nf trees directly comparable. The RF_adj_ values range from zero to one, indicating whether the tree topologies are identical or completely dissimilar, respectively. A principal component analysis (PCA) was generated from all RF_adj_ values in R to aid in visualization.

All nuclear and plastid phylogenies and their associated analyses described above were compared, but for simplicity only the ASTRAL, 1-to-1, and MO trees were analyzed in further analyses. ASTRAL was chosen as it selects the putative single-copy orthologs less rigorously than the remaining methods, creating a baseline method for comparison. We then compared ASTRAL to a method that disregards all genes containing paralogs (1-to-1) and a more rigorous paralog-processing method (MO). We chose MO over MI and RT due to its more stringent ortholog identification, which minimizes the inclusion of paralogs and reduces phylogenetic noise. Additionally, the conservative filtering of MO enhances alignment quality and computational efficiency, providing a more reliable foundation for downstream phylogenetic analyses (Yang and Smith 2014; Morales-Briones et al. 2021a; Jackson et al. 2023).

### Molecular Clock Dating

Estimates of divergence times were calculated on the ASTRAL, 1-to-1, and MO trees with RelTime (Tamura et al. 2012, 2018) using the command-line version of the MEGA software, MEGA-CC v. 11.0.11 (Kumar et al. 2012, 2016), with necessary input files generated with MEGA-X (Kumar et al. 2018; Mello 2018). The three Calenduleae taxa were used as outgroup taxa for this analysis.

There are no fossils available within *Packera* or tribe Senecioneae, so instead, multiple dating scenarios were tested utilizing both primary and secondary age constraints (Supplementary Table 5). Briefly, a maximum (59.95 MY) and minimum (31 MY) age was tested for the Anthemideae/Senecioneae split given those dates are the highest and lowest values within the 95% CI from a previous diversification rate analysis of Asteraceae (Mandel et al. 2019). Next, a maximum age of 34.06 MY was set at the *Packera* crown node from the upper age limit provided by (Mandel et al. 2019). Additionally, a minimum age constraint of 12 MY was tested within the outgroup (the Anthemideae clade) given the presence of a fossil pollen (Wang 2004; Hobbs and Baldwin 2013; Palazzesi et al. 2022). Finally, a maximum age constraint of 14.3 MY was tested for *Pericallis* D.Don (Senecioneae), a genus endemic to the Macaronesia islands (Jones et al. 2014), since that is the estimated age of Porto Santo, the oldest volcanic archipelago within the Madeira Islands (McDougall and Schmincke 1976; Geldmacher et al. 2000). Results of all tested scenarios were compared for biological relevance, though only one scenario was chosen for the discussion. This approach allowed us to focus on the topology that best aligned with prior systematic understanding and showed the strongest overall support across analytical frameworks.

### Biogeography

Ancestral ranges were estimated on the dated phylogenies using the package BioGeoBEARS v. 1.1 (Matzke 2014) in R. BioGeoBEARS estimates historical, biogeographic ranges by implementing popular models in a likelihood framework. Some of the most common models used are dispersal-extinction-cladogenesis (DEC; Ree and Smith 2008), the likelihood version of DIVA (DIVALIKE; Ronquist 1997), and the BayArea likelihood version of the range evolution model (BAYAREALIKE; Landis et al. 2013). In BioGeoBEARS, founder-event speciation can be added to any of these models (*+J*) with flexibility in determining relative probability. Likelihood Ratio Test (LRT) and Akaike Information Criterion (AIC) were used to determine which model best fits the data.

All *Packera* taxa, along with their four most closely related outgroup genera (*Pericallis*, *Elekmania* B.Nord., *Werneria* Kunth, and *Xenophyllum* V.A.Funk), were coded as present or absent in ten geographic ranges outlined in Takhtajan (1986) (Supplementary Table 6): Arctic Province, Canadian Province, Appalachian Province, Atlantic and Gulf Coast Plain Province, North American Prairies Province, Rocky Mountain and Vancouverian Province, Madrean Region, Asia, Europe, and Central America. Distributions were determined from previously published literature (Freeman and Barkley 1995; Trock 1999, 2006; Gramling 2006; Mahoney and Kowal 2008; Elven et al. 2011; Yeatts et al. 2011). Only four of the 26 outgroup taxa were included to limit the number of geographic ranges used in the analysis. The remaining outgroup taxa were pruned from the dated tree generated with RelTime using the *pxrmt* function in Phyx. Each taxon was limited to a maximum distribution of four ranges.

### Reticulation

The Species Networks applying Quartets (SNaQ) function in the Julia v. 1.8.1 (Bezanson et al. 2018) package PhyloNetworks v. 0.15.2 (Solís-Lemus and Ané 2016; Solís-Lemus et al. 2017) was used to investigate reticulate evolution among *Packera* taxa. This package uses unrooted gene trees to infer phylogenetic networks in a pseudolikelihood framework with multi-locus data, while accounting for ILS and reticulation. We tested the fit of multiple models and gradually increased the number of reticulation events (h) from 0 to 10 to find which pseudolikelihood score reached a plateau, given recommendations by (Solís-Lemus et al. 2017). This process is computationally intense, so only clades containing 14 taxa or less that had counter-support nodes in the ASTRAL, 1-to-1, and/or MO trees, as determined by a negative QC score resulting from Quartet Sampling, were used as input for PhyloNetworks, with the remaining taxa pruned from the tree using the *pxrmt* function in Phyx. Given this, five Packera clades were investigated for ancient reticulation events (Figs. 2-4). Additionally, a single species from each major clade was then used to represent backbone relationships throughout the entire phylogeny, totaling six separate PhyloNetwork runs that will be referenced as follows: Backbone, representing 11 taxa that span the entire phylogeny; Eastern, a clade of 12, 13, or 15 species (MO, 1-to-1, and ASTRAL, respectively) either distributed along eastern North America or having known affiliations with those species; California, eight taxa with distributions mainly within California; Mexico, seven (MO and ASTRAL) to eight (1-to-1) taxa with distributions mainly within Mexico; Rocky, a clade of five taxa that are distributed in the Central US, most within the Rocky Mountains; and Arctic, nine taxa that are present in colder climates along the Pacific Northwest and into Canada (Supplementary Table 7). Rocky could not be run for 1-to-1 since fewer than five taxa were present in that target clade.

## Results

### Sequencing and Matrix Results

Illumina sequencing resulted in approximately 733.3 million reads across 111 taxa. Read numbers ranged from 535,934 in *Werneria aretioides* Wedd. to 20,685,576 in *Senecio vulgaris* L. (full list in Supplementary Table 2). HybPiper retained 1,058 of the targeted 1,061 loci across all 111 taxa with 846 of those loci having paralog warnings (∼80%). The number of loci recovered for each taxon ranged from 566 in *Packera paupercula var. paupercula* (Michx.) Á.Löve & D.Löve to 1,054 in *Calendula arvensis* L. (Supplementary Table 8). The number of gene trees varied across analyses, from 285 in the 1-to-1 analysis to 7,388 in the MI analysis (Supplementary Table 3). Parsimony informative sites (PIS) also ranged from 2.11% in the plastid (mapping) tree to 99.64% in the MO tree (Supplementary Table 3). For species trees generated with ASTRAL-III, normalized quartet scores were used to measure gene tree discordance, which ranged from 0.516 for the RT tree to 0.534 for the MO tree (Supplementary Table 3), indicating that ∼52-53% of the gene trees matched the species tree in those analyses.

Off-target plastid sequencing resulted in partial (i.e., mapping; n = 111) to whole (i.e., complete; n = 18) plastomes. Complete plastomes from 17 *Packera* and one *Senecio* species ranged from 150,772 bp to 151,728 bp in size (Supplementary Fig. 2). For each consensus sequence mapped to the reference plastid assembly, *Packera cymbalaria*, the percentage of missing data varied drastically, with *Tephroseris newcombei* exhibiting the most missing data (93.1%; Supplementary Table 9, Supplementary Fig. 3). The mapped plastid supermatrix with 111 taxa had 2.11% PIS (Supplementary Table 3).

### Topological Comparisons and Discordance

Phylogenetic topologies varied depending on sequence type (e.g., nuclear or plastid) as well as paralog processing and phylogenetic tree inference method (i.e., ASTRAL, ASTRAL-Pro, 1-to-1, etc.) (Fig. 1, Supplementary Fig. 4). When building phylogenies with nuclear data, *Packera* was generally placed as the sister lineage to a clade comprising *Pericallis* and *Elekmania*, with the exception of ASTRAL-Pro which placed *Packera* as sister to *Tephroseris* and in a larger clade with *Pericallis* and *Elekmania* (Fig. 1). Interestingly, the plastid data showed a similar relationship to ASTRAL-Pro; however, *Pericallis* was placed unexpectedly in a separate clade with other Senecionineae taxa. All nuclear phylogenies recovered monophyletic tribes and subtribes (e.g., Senecionineae and Tussilaginineae s.str.), though the relationships within the subtribes varied. The mapped plastid tree generally separated the subtribes, apart from *Roldana* (Senecionineae) in Tussilaginineae s. str. (Fig. 1, Supplementary Fig. 4). The placement of *Tephroseris* (tribe Tephroseridineae) varied across phylogenies, with its most consistent placement as sister to Senecioneae (Fig. 1). *Doronicum* was consistently placed sister to Senecioneae, providing support for its elevation to tribal status (Doroniceae; Panero 2005; Fu et al. 2016).

**Figure 1.**
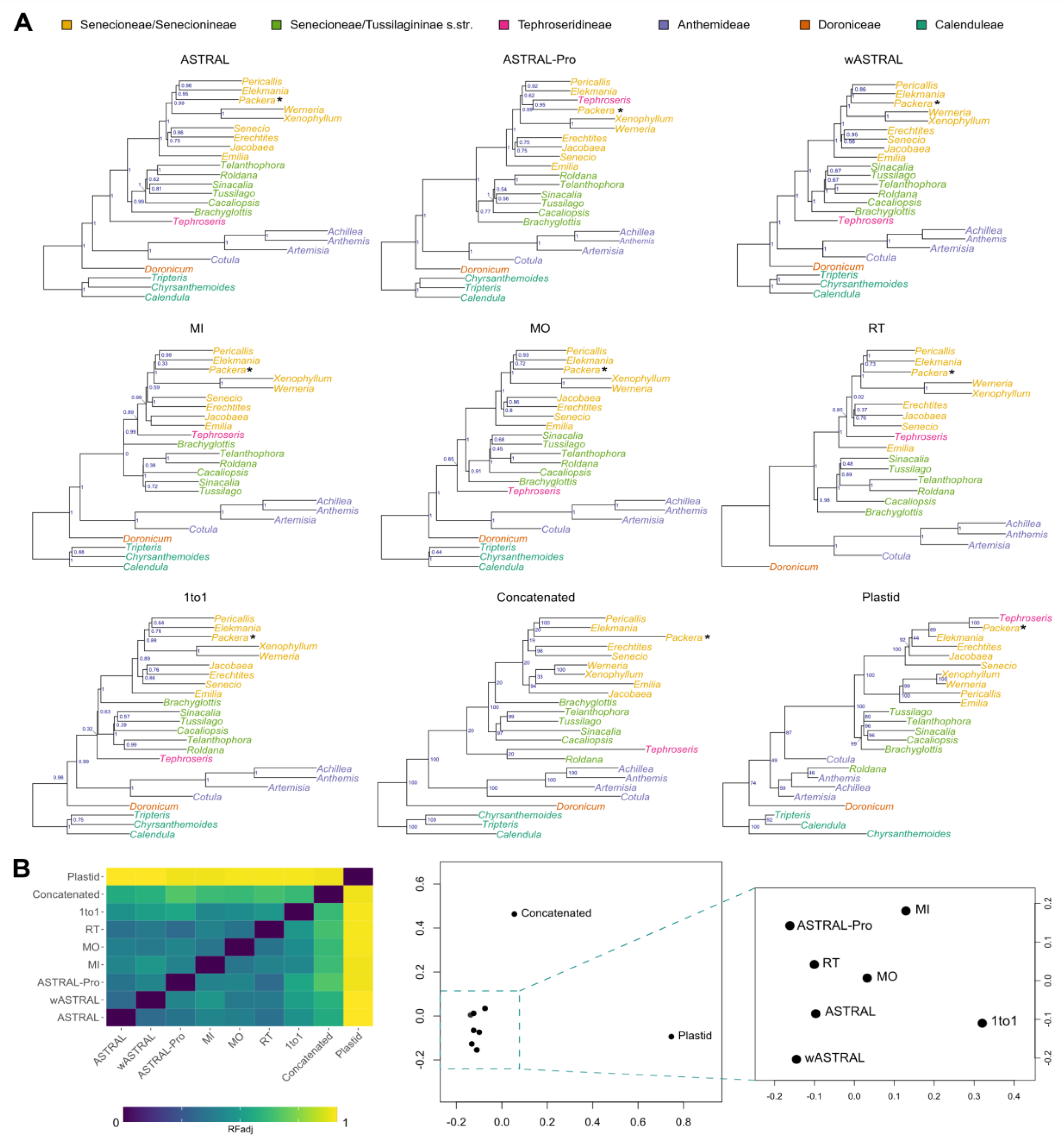
Genus-level phylogenies of all trees generated in this study. **A**) Genera are colored according to their tribal/subtribal classifications following the key on the top of the figure. *Packera* is marked with a black asterisk next to its placement in the trees. Navy blue text at node represents support values (LPP or BS). **B**) Bottom left is a matrix of RF_adj_ values between the nine different trees, with most similar topologies as dark blue squares (0) and least similar as yellow squares (1). The bottom right shows the RF_adj_ values plotted as a principal components analysis (PCA). Points closer together indicate higher similarity, while dots further away indicate higher dissimilarity.

Phylogenetic topologies remained highly incongruent independent of methods used (Figs. 2-4, Supplementary Table 10). The ASTRAL and wASTRAL tree topologies were most similar (RF_adj_ = 0. 324) (Fig. 1, Supplementary Table 10), while the plastid tree was consistently the most dissimilar to the remaining trees (RF_adj_ ranging from 0.895 to 0.914), with the concatenated tree being the next most dissimilar (RF_adj_ ranging from 0.562 to 0. 648; Supplementary Table 10). Additionally, the normalized quartet scores from ASTRAL indicate that only about half of the gene trees match the species trees in the ASTRAL and paragone-nf analyses (Supplementary Table 3). Most of the topological conflict was among *Packera* taxa, with minor differences among the remaining Senecioneae taxa and outgroups (Supplementary Fig. 4). Support values varied depending on which sequence type and method was used, though generally remained low. The wASTRAL and MI trees showed the highest node support with 58 out of 110 nodes (52.7%) having ≥ 0.95 LPP (Supplementary Table 3, Supplementary Fig. 4). In contrast, the plastid and concatenated trees showed much lower resolution with the fewest number of nodes having high support (36 of 110 nodes, 32.7%, have ≥ 95 BS; Supplementary Table 3, Supplementary Fig. 4).

**Figure 2.**
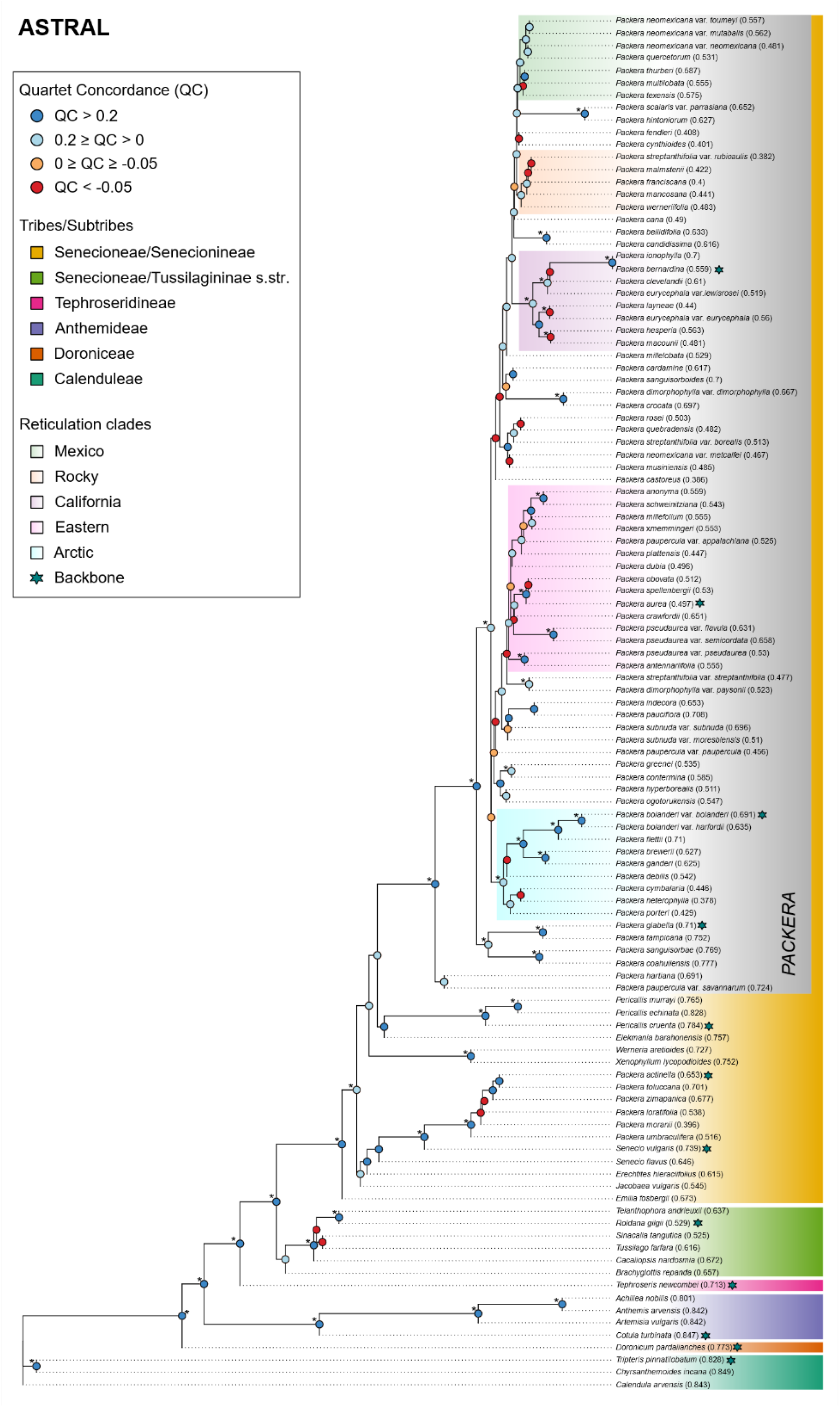
Full phylogeny of the ASTRAL analysis. Tribal/subtribal status is indicated on the right of the phylogeny and follows the color code as indicated in the key. Numbers next to tip labels correspond to the Quartet Fidelity (QF) score from Quartet Sampling, which ranges from 0 to 1, with values closer to 0 indicating erroneous taxa. Circles at nodes show the Quartet Concordance (QC) factors from Quartet Sampling, where red and orange indicate counter-support nodes. Asterisks next to QC circles indicate high support (≥ 95 LPP) at that node. Highlighted clades and teal stars next to some tip labels, indicate clades where reticulation events were tested, following the color key on the left.

Quartet Sampling indicated that underlying gene tree discordance remained high, independent of the method used. ASTRAL and concatenated trees had the most fully supported nodes (QS = 1/–/1) when compared to the remaining phylogenies (Supplementary Fig. 5), though the number was still notably low (∼10%). Even so, the concatenated tree also had the highest proportion of counter-support nodes (QC < 0; 26.4%; see Supplementary Fig. 5 for all Quartet Sampling results). PhyParts indicated that the ASTRAL and MO trees had the most concordant gene trees. The wASTRAL analysis had the highest percentages of gene trees supporting the top alternative bipartition. The MI tree harbored the highest discordance and uninformative data (Supplementary Fig. 6).

This underlying discordance could be caused by ILS, which was common in all tested ASTRAL trees (Supplementary Fig. 7). Introgressive hybridization (IH) was not detected without the presence of ILS in any of the nodes, though IH was most common in the nodes that were shown to be discordant (Supplemental Figs. 5, 7). Interestingly, crown *Packera* consistently had high IH values, potentially indicating introgression at that node. Backbone nodes were generally considered well resolved with high confidence (ILS index < 50% and IH index < 10%; Shang et al. 2025), especially in the outgroups, whereas *Packera* taxa appeared less resolved.

### Molecular Clock Dating

The resulting divergence rate estimates differed among ASTRAL, 1-to-1, and MO, with estimated ages of the MO and 1-to-1 trees older and sometimes almost double the ASTRAL ages. For ASTRAL, the mean crown age for *Packera* ranged from 11.4 MY in Scenario 5 and 6 to 13.5 MY in Scenario 1-4. Alternatively, the mean crown age of *Packera* were almost doubled in the 1-to-1 and MO analyses, with ranges from ∼21 MY in Scenario 5 to 35.5–35.7 MY in Scenario 4 (Supplementary Table 11, Supplementary Fig. 8). Scenario 2 was chosen as the best-fitting dating model because it incorporated all available primary calibration points and yielded divergence time estimates that align well with known patterns of evolutionary history (Pelser et al. 2010; Mandel et al. 2019), while minimizing reliance on secondary calibrations (Supplementary Table 5) (Müller and Reisz 2005; Benton and Donoghue 2007; Parham et al. 2012). The dated trees from all six scenarios can be found in Supplementary Figure 8.

### Biogeography

Evaluation of the six different model choices demonstrated that BAYAREALIKE was the best fit given our data based on its lowest log likelihood (Ln L) and highest AIC score, as well as the highest weighted AICc score for all three of our analyses (Supplementary Table 12). Generally, models including jump speciation (+J) provided a better fit than those without in the DEC and DIVALIKE models, and were the same or very similar in the BAYAREALIKE models (Supplementary Table 12), indicating a potential for jump dispersal or founder-event speciation in our data. Trees of all model runs are in Supplementary Figure 9.

The historical biogeography analysis revealed distinct geographic patterns across the different orthology methods (Fig. 5, Supplementary Fig. 9). ASTRAL identified the ancestral range as concentrated in the North American Prairies Province (N), Rocky Mountain and Vancouverian Province (R), and Madrean (M) regions. In contrast, MO showed the ancestral regions being only the Rocky Mountain and Vancouverian Province (R), and Madrean (M) regions. 1-to-1 was intermediate with crown *Packera* being ancestral to the North American Prairies Province (N), Rocky Mountain and Vancouverian Province (R), and Madrean (M) regions, similar to ASTRAL; however, the remaining backbone was Rocky Mountain and Vancouverian Province (R), and Madrean (M) regions, similar to MO. Notably, all analyses showed one major expansion into the eastern United States–Appalachian Province (P) and Atlantic and Gulf Coast Plain Province (T)–represented by the Eastern clade (Fig. 5, Supplementary Fig. 9), with a few instances of individual introductions to the eastern United States. Many of these taxa originated initially in the North American Prairies Province (N) and/or Rocky Mountain and Vancouverian Province (R) ranges. ASTRAL and MO showed two early expansions into the Arctic Province (I) and Canadian Province (C) at the base of the phylogeny, while 1-to-1 showed at least three expansions north. 1-to-1 exhibited the most geographic inconsistency, followed by MO. Even so, geography proved to be a reliable indicator for larger clades across all analyses, although widespread species were scattered across the phylogeny and did not always align consistently within geographic clades (Figs. 2-4, Supplementary Fig. 9).

### Reticulation events

Five *Packera* clades were tested for ancient reticulation events given a large number of counter-support (QC < 0) nodes from Quartet Sampling on the ASTRAL, 1- to-1, and MO trees (Figs. 2-4). Additionally, a single species from each major clade was then used to represent backbone relationships throughout the entire phylogeny, totaling six separate PhyloNetwork runs named: Arctic, Backbone, California, Eastern, Mexico, and Rocky (Supplementary Table 13). Log-likelihood values indicated that the most likely number of reticulation events (h) varied depending on the run (Supplementary Fig. 10). Additionally, Arctic, Mexico, and Backbone had varying numbers of optimal reticulation events between the paralog-processing methods (Supplementary Table 7, Supplementary Fig. 10). For ASTRAL, Arctic and Mexico had the highest number of reticulation events (h = 3), followed by California and Backbone (h = 2), then Eastern and Rocky (h = 1). For 1-to-1, Backhone had the most (h = 6), then Mexico (h = 3), Arctic and California (h = 2), and then Eastern (h = 1). For MO, Arctic, California, and Backbone had the highest number of reticulations (h = 2) with Eastern, Mexico, and Rocky having the lowest (h = 1) (Fig. 6, Supplementary Fig. 10, Supplementary Table 7). The taxa also differed between analyses given different phylogenetic relationships between clades (Supplementary Table 13), impacting some of the networks identified.

Some networks showed different events depending on the phylogenetic method and subset of data (Fig. 6); however, some consistent patterns emerged. For example, the Eastern clade continuously showed the same reticulation event regardless of the paralog processing method, and parentage percentages were very similar. In contrast, Rocky did not show the same reticulation events, but the networks generally involved *Packera mancosana* and *P. streptanthifolia*. The California clade showed differing results across methods: both ASTRAL and 1-to-1 analyses identified *P. bernardina* as the product of reticulation with the same parentage, whereas MO did not yield the same results, and no consistent trends emerged. The Mexico clade was generally inconsistent, but featured reticulation events involving *P. neomexicana* and its varieties. Additionally, *P. texensis* and *P. quercetorum* were involved in reticulation events in two of the three paralog processing networks. Among all the clades, the Arctic clade exhibited the most variability in the number of reticulation events, with the *P. bolanderi* varieties consistently involved in some form of reticulation. However, the specifics of these events varied depending on sampling and paralog processing methods. In the Backbone subtree analysis, *Tephroseris* was frequently identified as a product of reticulation, with 1-to-1 and MO showing the same parentage. In contrast, ASTRAL did not show the same reticulation events but did identify a reticulation event involving *Tephroseris* with a different parentage. Notably, the parentage of *Tephroseris* always included a *Packera* taxon, though the specific taxon varied.

## Discussion

Gene duplication was pervasive in *Packera*, with more than 80% of nuclear loci showing evidence of paralogy (Supplementary Table 3). Such widespread paralogy poses major challenges for species tree inference, especially in Hyb-Seq datasets, where probes may unintentionally enrich for multiple homeologs (Salmon et al. 2012; Akama et al. 2014; Crameri et al. 2022), complicating the recovery of true single-copy orthologs for phylogenetic reconstructions. Although some paralogy is expected in target enrichment studies (Siniscalchi et al. 2021), the extent observed in *Packera* is unusually high and reflects lineage-specific genome dynamics, rather than a general feature of all species. This has important methodological implications, as HybPiper’s orthology inference relies largely on single-copy filtering (Johnson et al. 2016), and using its outputs alone can introduce noise and bias into gene tree reconstruction, ultimately distorting species tree inference. Consistent with our study, using only putative single-copy orthologs designated by HybPiper was insufficient for accurate phylogenetic inference in *Packera*, a group characterized by extensive polyploidy and hybridization (Fig. 2) (Trock 2006). These challenges are not unique to *Packera*, as many plant and non-plant lineages also experience difficulties reconstructing phylogenies due to hybridization and gene/genome duplications. Together, these examples highlight the complexity of orthology inference in phylogenomics and the need for rigorous paralog processing across both plant and non-plant systems.

The most recent phylogeny of *Packera* prior to this study was based on ITS data and revealed extremely low sequence variation, with only 0–5% divergence and several *Packera* taxa sharing identical sequences (Bain and Golden 2000). This led to a largely unresolved tree, likely due to few phylogenetically informative sites, sparse species sampling, and potential reticulation events (Funk 1985; Linder and Rieseberg 2004). By using Hyb-Seq data and broader taxon sampling, we substantially improved phylogenetic resolution across *Packera*, including within major clades. However, our analyses also revealed that gene tree concordance and species tree resolution varied substantially across methods (Supplementary Fig. 4), exemplifying the influence of paralog processing strategies on downstream phylogenetic outcomes. Methods that explicitly model gene duplication and loss, such as those implemented in paragone-nf (Yang and Smith 2014; Jackson et al. 2023), generally yielded more concordant gene trees and better-resolved species trees (Figs. 3, 4). However, substantial conflict persisted across the trees, indicating that even sophisticated modeling approaches cannot fully resolve deep relationships in groups with complex histories like *Packera*. These discrepancies may in part reflect differences in the number and identity of genes retained after filtering (Supplementary Table 3). For instance, the 1-to-1 method, which excludes all loci with paralogs and resembles approaches used in tools like Phyluce, eliminates a significant portion of the dataset and may oversimplify the data, especially in duplication-rich systems such as *Packera*. While Phyluce and 1-to-1 filtering both remove paralogs, the underlying methods differ (Faircloth 2016; Jackson et al. 2023), causing Phyluce to outperform HybPiper in certain cases (Mandel et al. 2019; Thapa et al. 2020). Similarly, the RT method naturally loses information since it removes outgroups to determine orthology (Yang and Smith 2014; Morales-Briones et al. 2021b, 2021a) and potentially limits phylogenetic signals. These findings highlight that while some approaches have potential to improve concordance, none fully resolved the underlying discordance of *Packera*, emphasizing that the choice of method, underlying data, and biological context must guide ortholog selection in phylogenomic analyses.

**Figure 3.**
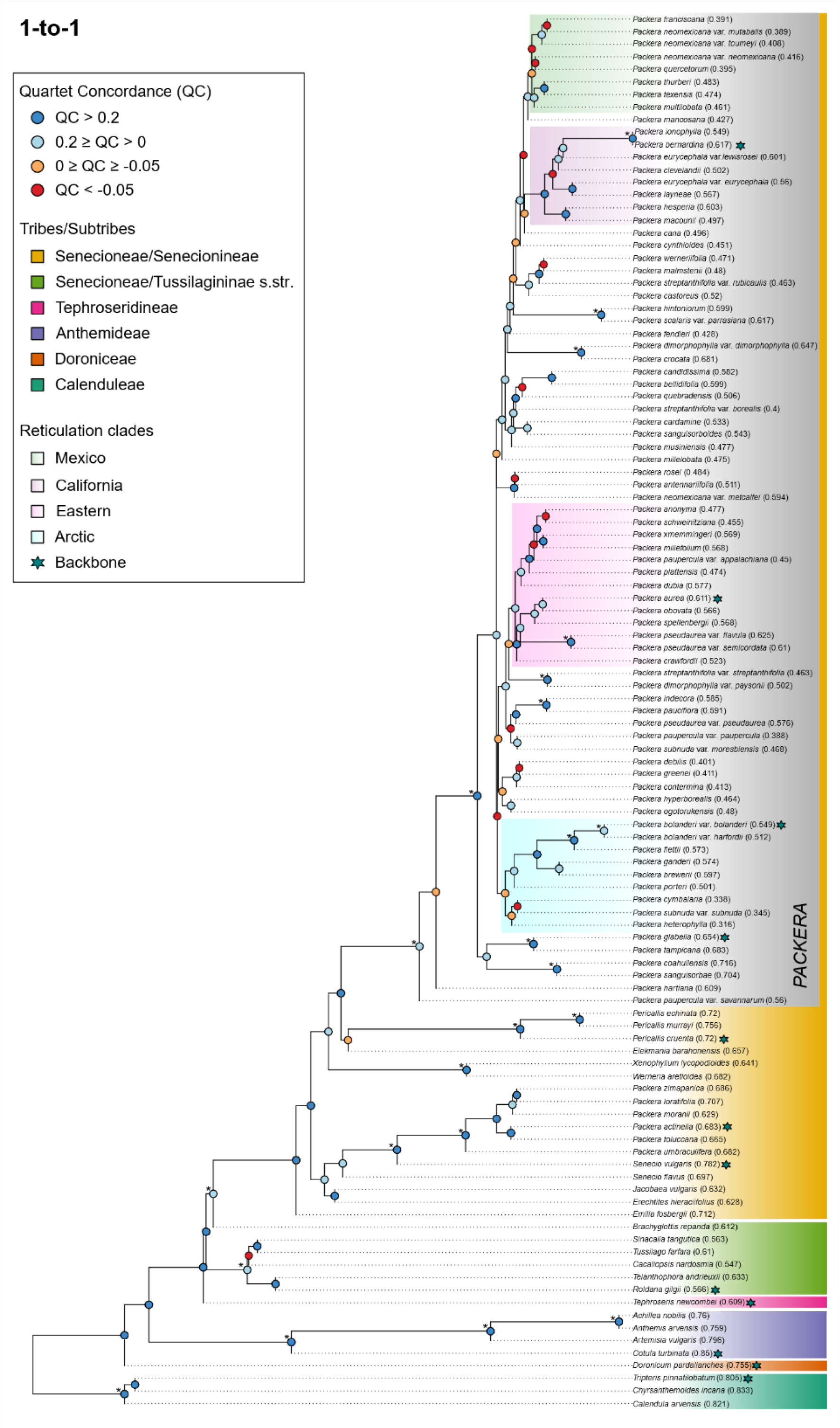
Full phylogeny of the ‘one-to-one ortholog’ (1-to-1) analysis. Tribal/subtribal status is indicated on the right of the phylogeny and follows the color code as indicated in the key. Numbers next to tip labels correspond to the Quartet Fidelity (QF) score from Quartet Sampling, which ranges from 0 to 1, with values closer to 0 indicating erroneous taxa. Circles at nodes show the Quartet Concordance (QC) factors from Quartet Sampling, where red and orange indicate counter-support nodes. Asterisks next to QC circles indicate high support (≥ 95 LPP) at that node. Highlighted clades and teal stars next to some tip labels, indicate clades where reticulation events were tested, following the color key on the left.

Results from a previous phylogenetic study of *Packera* showed that its evolutionary relationships tend to loosely follow a geographic pattern (Bain and Golden 2000), which is consistent with our findings (Figs. 2-4, Supplementary Fig. 9). Our nuclear phylogenies showed that major clades tended to contain taxa within similar geographic ranges, independent of which paralog processing method was utilized. Additionally, most taxa within these major clades were conserved across methods, though their placement along the backbone, and some taxa, did vary (Figs. 2-4). For example, the taxa in the Eastern clade was fairly consistent between the orthology inference methods, although some taxa did move in and out of the clade (Supplementary Fig. 4). Interestingly, several of these taxa are widespread and engage in interspecific hybridization with other *Packera* species where their ranges overlap. For example, *P. pseudaurea* and its varieties are typically found in central North America, though their phylogenetic placement is generally in the Eastern clade. These results are unsurprising as they have an extensive history of hybridization with other *Packera* taxa in the Eastern clade, which is believed to have created much taxonomic uncertainty (Barkley 1962; Trock 1999), potentially explaining their inconsistent phylogenetic placements (Supplementary Fig. 4). However, conflicting signals may also reflect additional evolutionary processes beyond hybridization, such as cryptic speciation within widespread taxa or the influence of unsampled and/or extinct lineages.

**Figure 4.**
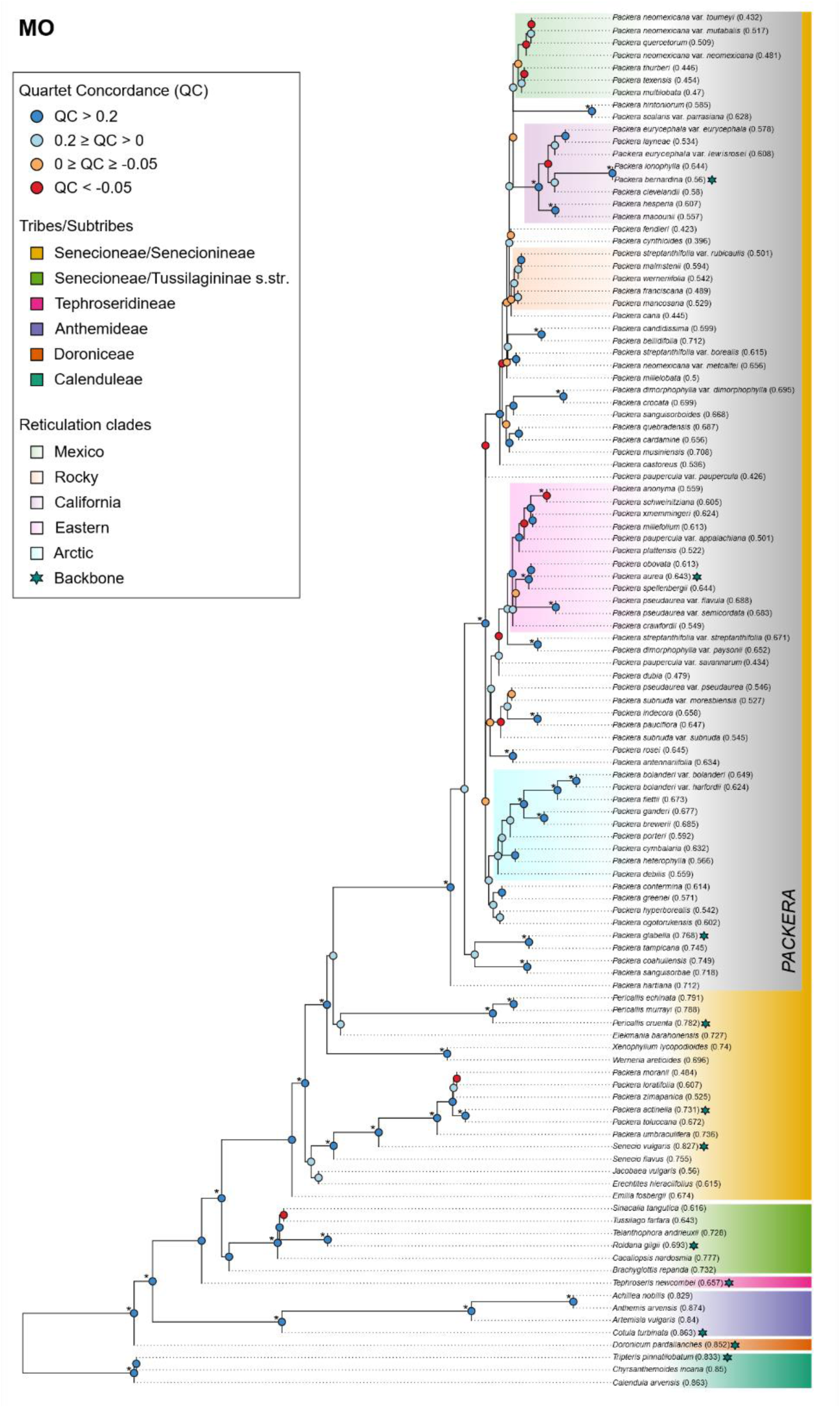
Full phylogeny of the ‘monophyletic outgroup’ (MO) analysis. Tribal/subtribal status is indicated on the right of the phylogeny and follows the color code as indicated in the key. Numbers next to tip labels correspond to the Quartet Fidelity (QF) score from Quartet Sampling, which ranges from 0 to 1, with values closer to 0 indicating erroneous taxa. Circles at nodes show the Quartet Concordance (QC) factors from Quartet Sampling, where red and orange indicate counter-support nodes. Asterisks next to QC circles indicate high support (≥ 95 LPP) at that node. Highlighted clades and teal stars next to some tip labels, indicate clades where reticulation events were tested, following the color key on the left.

Our results also reveal that evolutionary interpretations such as divergence dating and biogeographic reconstruction are highly sensitive to the paralog processing approach used to infer the underlying phylogeny. Divergence time estimates varied across methods, with some species trees suggesting that crown *Packera*’s diversification occurred more recently (ASTRAL: 13.5 MY), while others placed key divergences deeper in time (1-to-1: 24.2 MY and MO: 25 MY; Supplementary Fig. 8, Supplementary Table 11). These differences in divergence times are not trivial, as they affect our understanding of how *Packera* taxa respond to climatic shifts, mountain uplift, and other historical events, and have broader implications for comparative studies across Asteraceae and other radiating plant lineages. For example, tribal- and family-wide studies predicted the crown age of *Packera* to be fairly young, between 7.8 MY and 20.2 MY (Pelser et al. 2010; Mandel et al. 2019; Zhang et al. 2021), which is more in-line with the ASTRAL analysis. While more *Packera*-specific predictions are older, with (Barkley 1988) predicting the origin of *Packera* in the later stages of the mid-Tertiary (∼22-37 MY), closely aligning with the paragone-nf analyses (Supplementary Fig. 8). Similarly, biogeographic patterns shifted depending on the inferred topology, with each analysis showing different migration patterns over time (Fig. 5, Supplementary Fig. 9). Even so, the broad pattern is relatively unchanged. For example, all topologies generally agreed that *Packera* originated in western North America with one major expansion east and then later expansions north from the western clades. From there, individual taxa expanded or became specialized to certain regions, with most of the diversity present in western North America. Because the topologies are consistent across methods, we can be more confident that they likely reflect the true evolutionary history of *Packera* instead of methodological artifacts.

**Figure 5.**
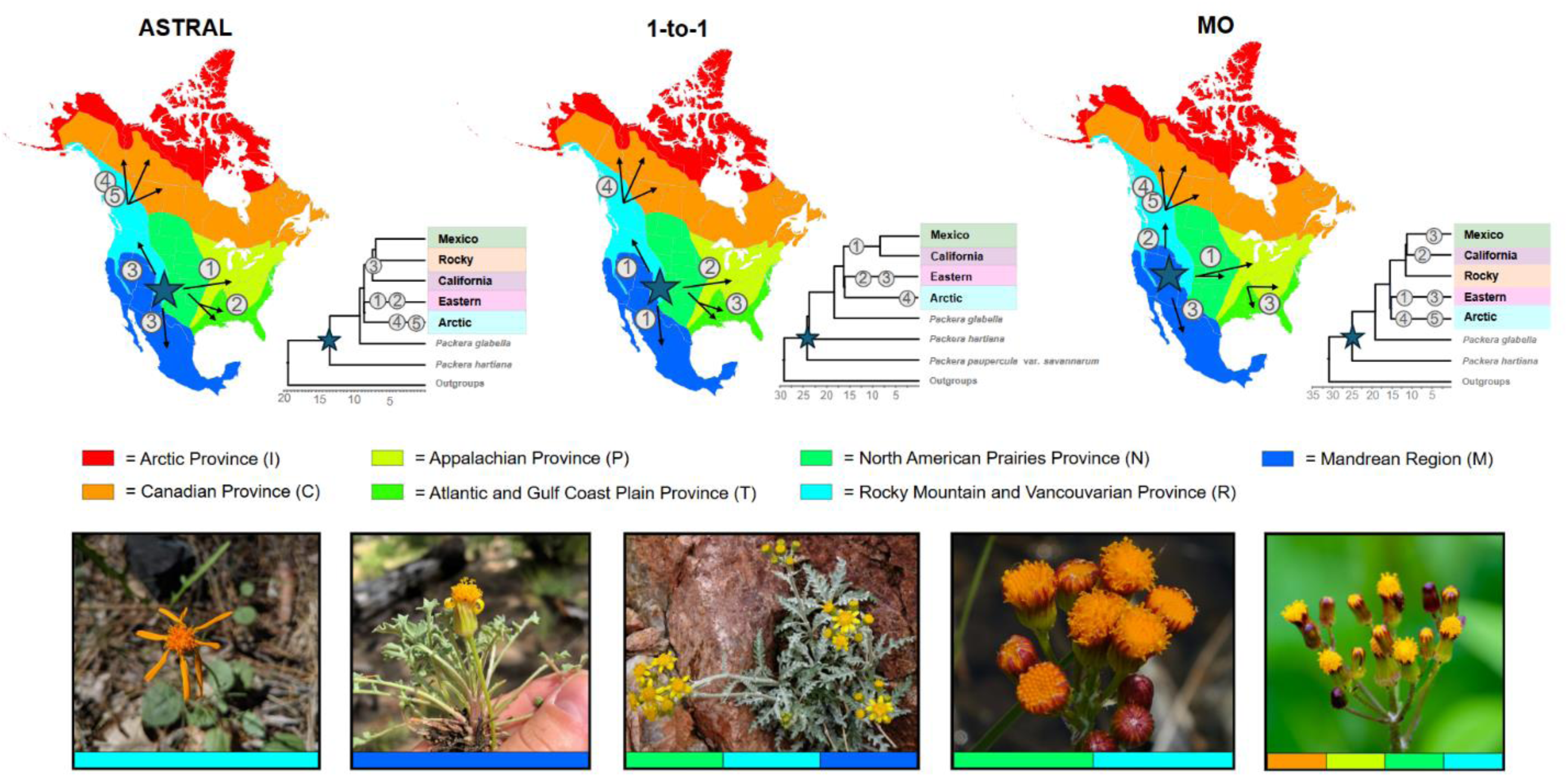
Predicted biogeographic region summaries generated from BioGeoBEARS of the ASTRAL, 1-to-1, and MO analyses. The maps of North America are colored according to the ten geographic regions outlined by Takhtajan (1986): Arctic Province (I), Canadian Province (C), Appalachian Province (P), Atlantic and Gulf Coast Plain Province (T), North American Prairies Province (N), Rocky Mountain and Vancouvarian Province (R), Madrean Region (M), Asia (A), Europe (E), and Central America (S). Stars indicate the predicted ancestral range of crown *Packera*. Arrows and numbers correspond to their predicted range expansions in chronological order, with 5 being the most recent major expansion. Numbers also correspond to the dated tree to the right of the phylogeny, which indicates the nodes the expansions occurred in relation to the major geographic clades designated in Figures 2-4. All BioGeoBears results can be found in Supplementary Figure 9. Images of *Packera* species highlighting some of the morphological diversity seen in the group. Colored bars on the bottom of the image correspond to their current distributions, following the color scheme of the map and key. From left to right: Flame ragwort, *P. greenei* (A.Gray) W.A.Weber & Á.Löve; Tehachapi ragwort, *P. ionophylla* (Greene) W.A.Weber & Á.Löve; Fendler’s ragwort, *Packera fendleri* (A.Gray) W.A.Weber & Á.Löve; Weak groundsel, *P. debilis* (Nutt.) W.A.Weber & Á.Löve; and Rayless mountain ragwort, *P. indecora* (Greene) Á.Löve & D.Löve. Image credits: Erika R. Moore-Pollard and Robert Lagier.

**Figure 6.**
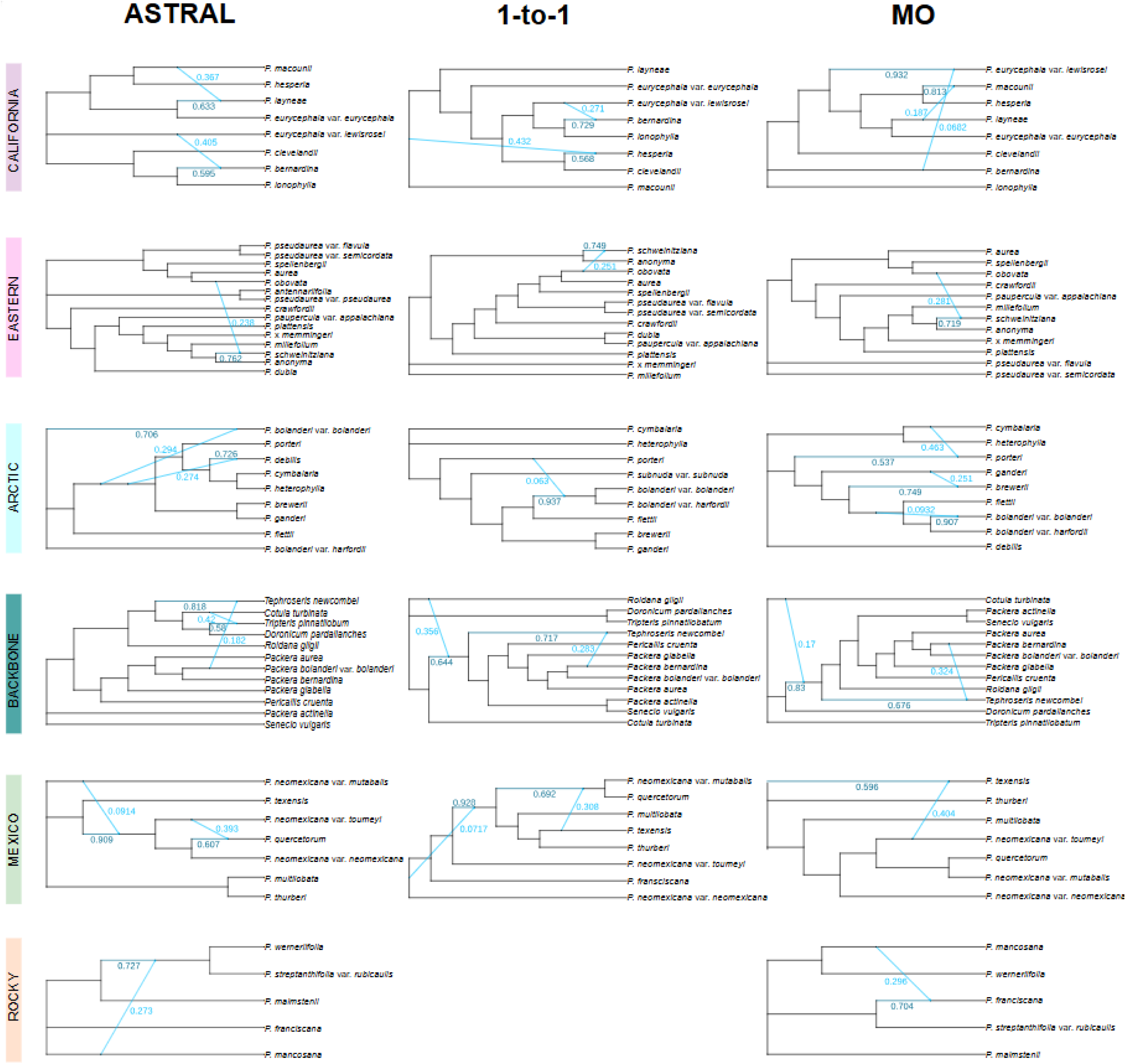
PhyloNetworks figures of the three tested analyses: ASTRAL, 1-to-1, and MO. The various areas of the phylogeny that were tested are shown on the left and can be read as a row. Dark green edges indicate the major hybrid edge, while the light blue shows the minor hybrid edge. The associated numbers next to each branch correspond to the percentage of parentage given by that branch. Number of hybridizations correspond with the best log likelihood value. The 1-to-1 analysis does not have a Rocky analysis since the clade in that tree only had four taxa.

Despite improvements from paralogy-aware filtering, the species trees for *Packera* were not fully resolved, indicating that underlying biological processes such as ILS, hybridization, and introgression also play significant roles in shaping *Packera*’s evolutionary history. This is supported by strong cytonuclear discordance, where plastid phylogenetic relationships vary significantly from the nuclear trees (Fig. 1, Supplementary Fig. 4), consistent with chloroplast capture or ancient hybridization events (Doyle 1992; Maddison 1997; Galtier and Daubin 2008; Soltis and Soltis 2009; Smith et al. 2015; Stull et al. 2020). Directly testing for gene flow revealed multiple introgression events, particularly along short internal branches susceptible to ILS (Supplementary Fig. 7). However, the specific reticulation patterns recovered varied depending on the paralog processing method used. These discrepancies highlight a critical point: gene selection and paralog processing directly shapes reticulation inference. Each gene has its own evolutionary trajectory, shaped by recombination and gene duplication and loss (Betrán 2015), and filtering strategies that exclude paralogs or collapse duplications can erase signals of past hybridization. As a consequence of different orthology and paralog processing methods, some reticulation events, such as those involving Eastern *Packera* lineages, were consistently recovered across datasets, while others appeared only in specific analyses.

Network analyses, such as PhyloNetworks, are inherently sensitive to sampling bias (Solís-Lemus and Ané 2016). Even though we sampled 85 out of a targeted 91 *Packera* taxa (∼93% coverage of the genus), missing taxa may have impacted our results. Including the remaining and unsampled *Packera* taxa in future studies could help increase our understanding of these relationships. Additionally, limiting the analyses to specified clades can ignore other potential reticulation events occurring across groups. We recognize this limitation; however, expanding sampling to more taxa or loci is currently restricted by the intensive and often unfeasible computational requirements of reticulation network analyses (see Hejase and Liu 2016). With recent bioinformatic advances and increased interest in non-bifurcating trees, we anticipate that less computationally intense and more efficient phylogenetic and network tools will soon be developed, allowing researchers to test for reticulations across large phylogenies.

Ghost introgressions, which are ancient introgression events that leave genetic signatures of extinct species in present-day species (Ottenburghs 2020), also pose challenges to inferences made by hybridization networks. Accounting for ghost introgressions in macroevolutionary studies may help improve the accuracy of trait evolution and biogeographic reconstructions, refine divergence time estimates, and reveal hidden contributions of hybridization to diversification (Ottenburghs 2020; Tricou et al. 2022). Network methods such as PhyloNetworks can detect ghost introgressions, but these methods do not specify that the hybrid edge is a ghost introgression, so it is up to the researcher to recognize its presence, which can be difficult especially without background knowledge of the group (Solís-Lemus and Ané 2016; Solís-Lemus et al. 2017).

Our findings emphasize that accurately reconstructing reticulate evolutionary histories requires embracing the independent histories of genes and carefully considering how orthology is defined. In *Packera*, gene tree heterogeneity is expected and evolutionarily informative rather than mere noise. Methods that oversimplify gene relationships, such as pipelines restricted to single-copy orthologs, risk masking the very patterns needed to understand complex evolutionary processes. By comparing alternative strategies for handling paralogs and integrating gene flow analyses, we show that approaches which explicitly model gene duplication and loss provide modest improvements in resolution and concordance but are insufficient on their own to fully resolve relationships. Direct tests for reticulation and ILS revealed the extent to which *Packera*’s evolutionary history has been shaped by hybridization and gene flow, underscoring the broader challenge of capturing complexity in phylogenomic datasets. Within polyploid systems, the tradeoff between excluding paralogs to reduce conflict and retaining them to preserve evolutionary signals becomes especially pronounced.

These results also highlight how paralog handling influences not only phylogenetic reconstruction but also the biological inferences drawn from the inferred relationships. Macroevolutionary analyses such as divergence dating, trait evolution, and biogeography are highly sensitive to orthology inference strategies and are only as reliable as the phylogenetic framework on which they are built (e.g., Magalhaes et al. 2021). Improving that framework will require both methodological innovation and deeper biological insight. For example, targeted probe sets designed to enrich truly single-copy loci (e.g., Compositae-ParaLoss-1272; Moore-Pollard et al. 2024) offer one path forward. Yet entirely excluding paralogs may discard informative loci, while retaining them without appropriate modeling risks misleading inference. Integrating morphological, ecological, and geographic evidence is therefore essential for assessing whether patterns inferred from large-scale datasets reflect genuine evolutionary history or methodological artifacts.

Rather than treating paralogy as a technical obstacle to be circumvented in phylogenetic studies, our findings underscore its potential to illuminate fundamental evolutionary processes, including gene duplication, polyploidy, and reticulation, that have shaped the diversification of lineages across the tree of life. While these challenges are common to many plant and non-plant groups, our densely sampled *Packera* dataset provides a uniquely powerful system for testing and improving phylogenomic methods. As the field advances toward increasingly nuanced and biologically realistic models of genome evolution, systems like *Packera* will be essential for developing tools that can fully capture the intricacy of complex evolutionary histories.

## Supporting information

Individual tree files

Alignment files

## Acknowledgments

The authors thank Drs. Carol Siniscalchi and Matthew Pollard for their bioinformatic help, Drs. Randall Bayer, L. Ellie Buckland, Duane McKenna, Anna Pardo, and Linda Watson for reading and editing prior to submission, and Dr. John Bain, Dr. Ed Schilling, and Brandon Fuller for donating genetic material for this study. We would also like to thank the following herbaria for allowing us to visit and/or destructively sample leaf tissue: ALA, ARIZ, ASU, CAS, MEM, NY, OSC, TEX, UNC, UCR, WTU. Permits to collect in the San Bernardino National Forest and Shasta-Trinity National Forest in California were obtained prior to collection of specimens through the USDA Forest Service. We also thank the University of Memphis High-Performance Cluster (HPC) administrators, Eric Spangler and Kristian Skjervold, for their assistance with the HPC and willingness to help. This work was supported by the Center for Biodiversity Research at the University of Memphis.

## Data Availability Statement

Raw sequence data are available in the National Center for Biotechnology Information (NCBI) Sequence Read Archive (SRA) under BioProjects PRJNA907383, PRJNA978568, PRJNA540287, and PRJNA516161.

---

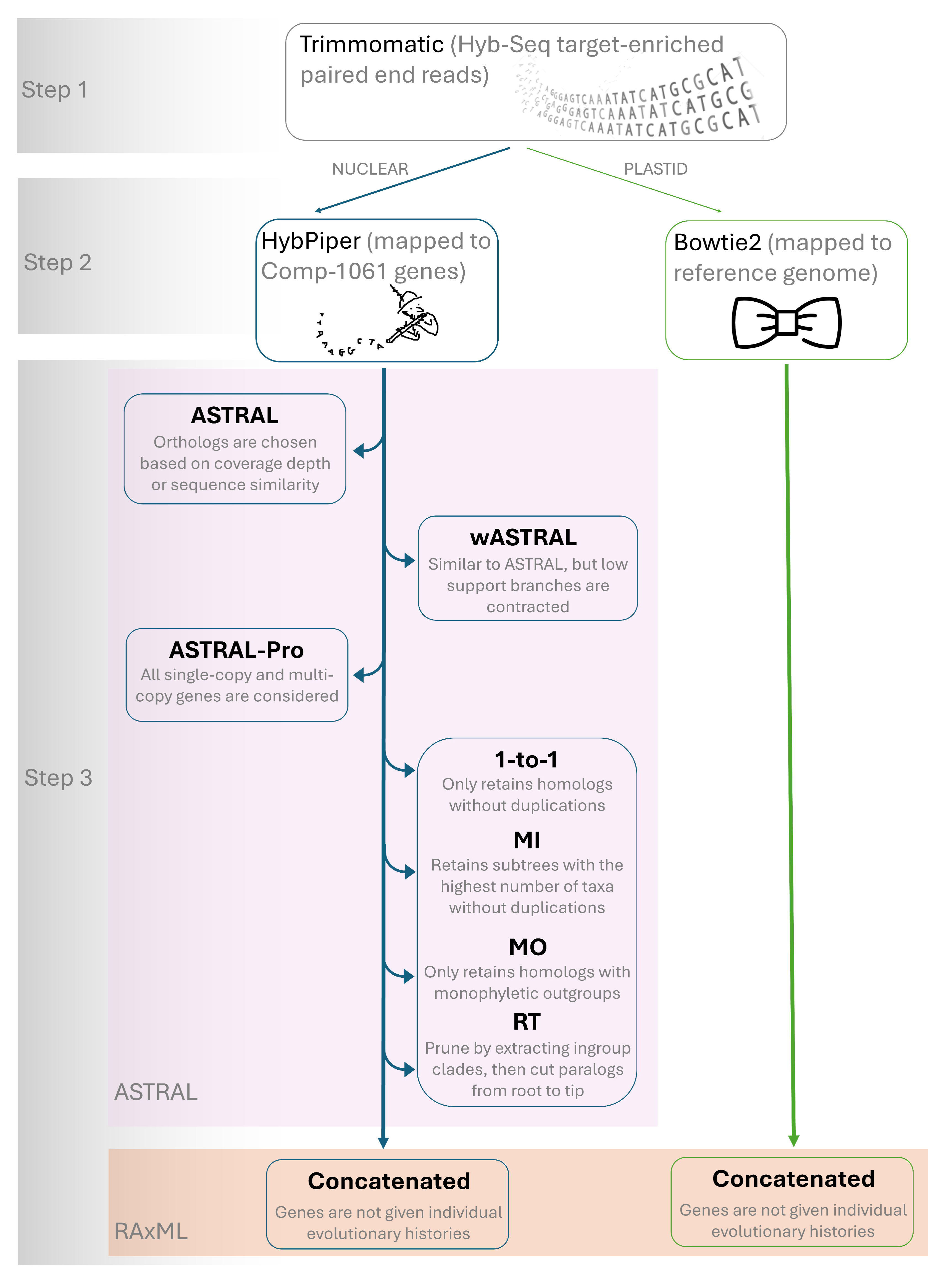

