## Supplementary Figure 2 for "Phylogenomic challenges in polyploid-rich lineages: Insights from paralog processing and reticulation methods using the complex genus *Packera* (Asteraceae: Senecioneae)"

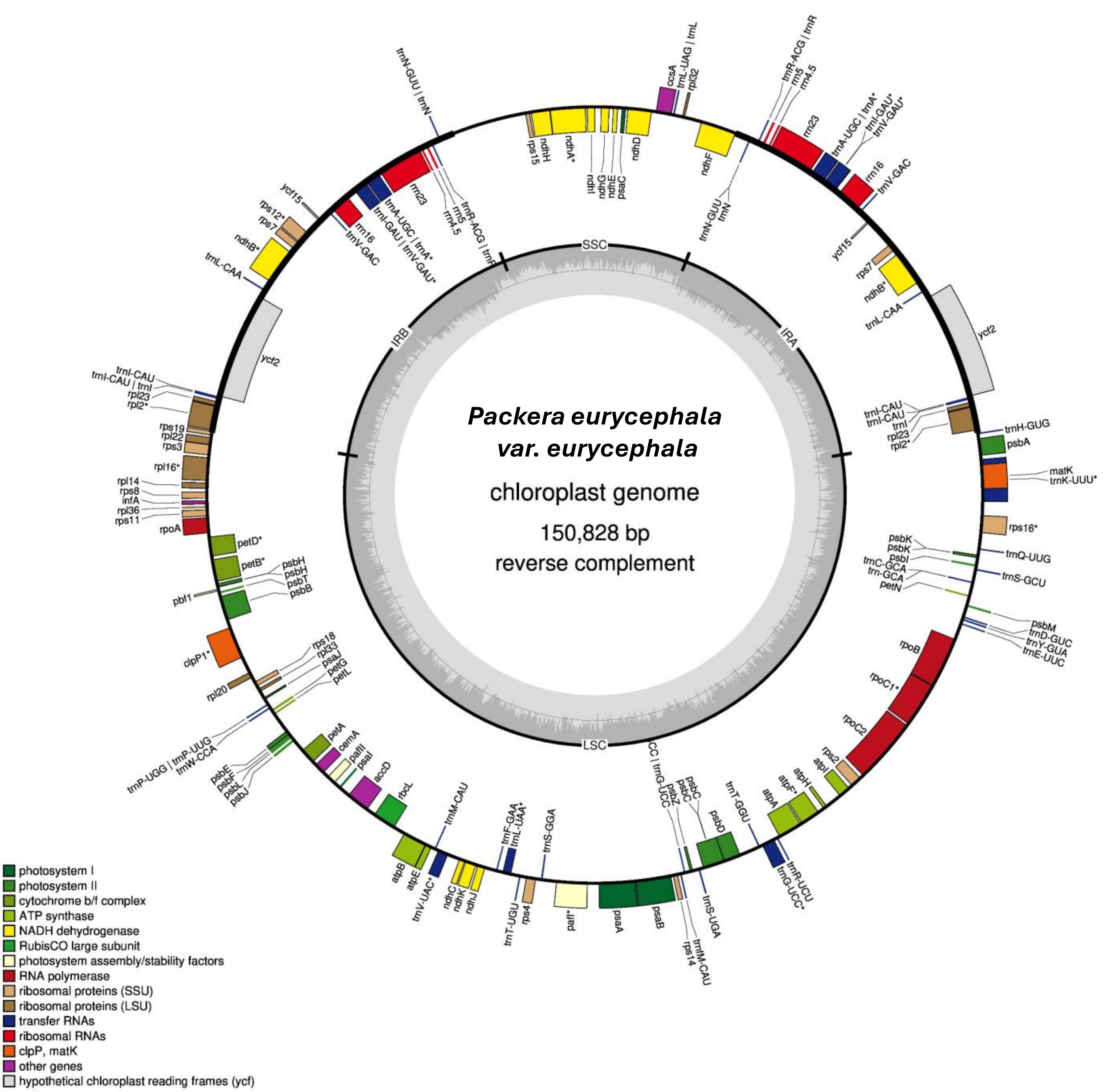

Packera eurycephala
var. eurycephala
chloroplast genome
150,828 bp
reverse complement
SSC
IRA
IRB
LSC
ccsA
trnL-UAG | trnL
rpl32
ndhF
trnN-GUU
trnN
trnR-ACG | trnR
rrn5
rrn4.5
rrn23
trnA-UGC | trnA*
trnI-GAU*
trnV-GAU*
rrn16
trnV-GAC
ycf15
rps7
ndhB*
trnL-CAA
ycf2
trnI-CAU
trnI-CAU
trnI
rpl23
rpl2*
trnH-GUG
psbA
matK
trnK-UUU*
rps16*
psbK
psbK
psbI
trnC-GCA
trn-GCA
petN
trnQ-UUG
trnS-GCU
psbM
trnD-GUC
trnY-GUA
trnE-UUC
rpoB
rpoC1*
rpoC2
rps2
atpI
atpH
atpF*
atpA
trnT-GGU
trnR-UCU
trnG-UCC*
psbD
psbC
psbC
psbZ
trnG-UCC
trnS-UGA
trnfM-CAU
rps14
psaB
psaA
paf*
rps4
trnT-UGU
trnS-GGA
trnL-UAA*
trnF-GAA
ndhJ
ndhK
ndhC
trnV-UAC*
trnM-CAU
atpE
atpB
rbcL
accD
psaI
pafII
cemA
petA
psbJ
psbL
psbF
psbE
trnW-CCA
trnP-UGG | trnP-UUG
rpl20
clpP1*
rps18
rpl33
psaJ
petG
petL
pbf1
psbB
psbT
psbH
psbH
petB*
petD*
rpoA
rps11
rpl36
infA
rps8
rpl14
rpl16*
rps3
rpl22
rps19
rpl2*
rpl23
trnI-CAU | trnI
trnI-CAU
ycf2
trnL-CAA
ndhB*
rps7
rps12*
ycf15
rrn16
trnV-GAC
trnI-GAU | trnV-GAU*
trnA-UGC | trnA*
rrn23
rrn4.5
rrn5
trnR-ACG | trnR
trnN-GUU | trnN
rps15
ndhH
ndhA*
ndhI
ndhG
ndhE
psaC
ndhD
photosystem I
photosystem II
cytochrome b/f complex
ATP synthase
NADH dehydrogenase
RubisCO large subunit
photosystem assembly/stability factors
RNA polymerase
ribosomal proteins (SSU)
ribosomal proteins (LSU)
transfer RNAs
ribosomal RNAs
clpP, matK
other genes
hypothetical chloroplast reading frames (ycf)

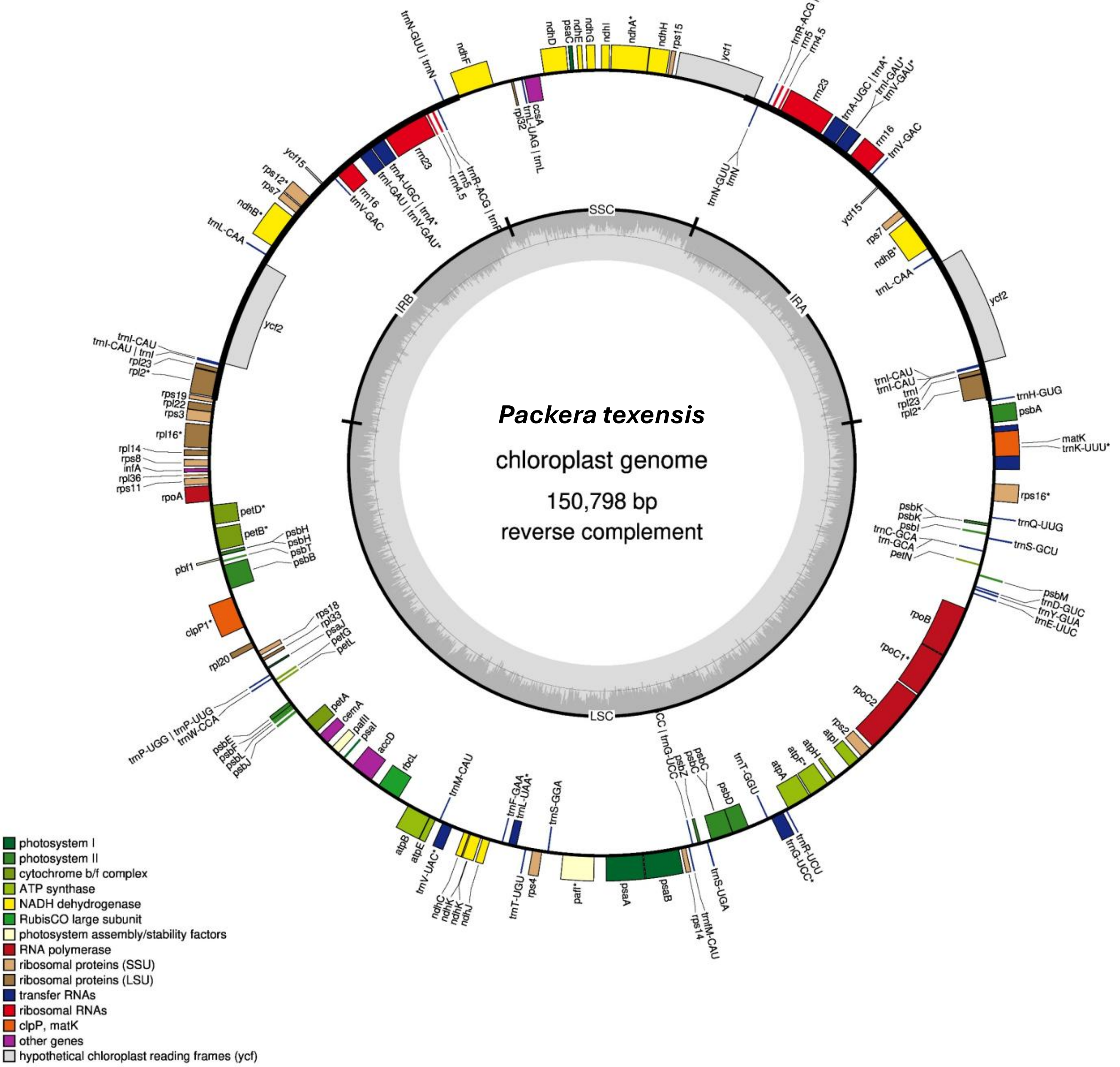

*Packera texensis*
chloroplast genome
150,798 bp
reverse complement
SSC
IRA
IRB
LSC
trnN-GUU | trnN
ndhF
rpl32
trnL-UAG | trnL
ccsA
ndhD
psaC
ndhE
ndhG
ndhI
ndhA*
ndhH
rps15
ycf1
trnR-ACG | trnR
rrn5
rrn4.5
rrn23
trnA-UGC | trnA*
trnI-GAU*
trnV-GAU*
rrn16
trnV-GAC
trnN-GUU
trnN
ycf15
rps7
ndhB*
trnL-CAA
ycf2
trnI-CAU
trnI
rpl23
rpl2*
trnH-GUG
psbA
matK
trnK-UUU*
rps16*
psbK
psbI
trnQ-UUG
trnC-GCA
trn-GCA
trnS-GCU
petN
psbM
trnD-GUC
trnY-GUA
trnE-UUC
rpoB
rpoC1*
rpoC2
rps2
atpI
atpH
atpF*
atpA
trnR-UCU
trnG-UCC*
trnT-GGU
psbD
psbC
psbZ
trnS-UGA
trnfM-CAU
rps14
psaB
psaA
pafI*
rps4
trnT-UGU
trnS-GGA
trnF-GAA
trnL-UAA*
ndhC
ndhK
ndhJ
trnV-UAC*
trnM-CAU
atpE
atpB
rbcL
accD
psaI
pafII
cemA
petA
psbJ
psbL
psbF
psbE
trnP-UGG | trnP-UGG
trnW-CCA
rpl20
rps18
rpl33
psaJ
petG
petL
clpP1*
pbf1
psbB
psbT
psbH
petB*
petD*
rpoA
rps11
rpl36
infA
rps8
rpl14
rpl16*
rps3
rpl22
rps19
trnI-CAU | trnI
rrn23
rrn5
rrn4.5
trnR-ACG | trnR
trnA-UGC | trnA*
trnI-GAU | trnV-GAU*
rrn16
trnV-GAC
rps12*
photosystem I
photosystem II
cytochrome b/f complex
ATP synthase
NADH dehydrogenase
RubisCO large subunit
photosystem assembly/stability factors
RNA polymerase
ribosomal proteins (SSU)
ribosomal proteins (LSU)
transfer RNAs
ribosomal RNAs
clpP, matK
other genes
hypothetical chloroplast reading frames (ycf)

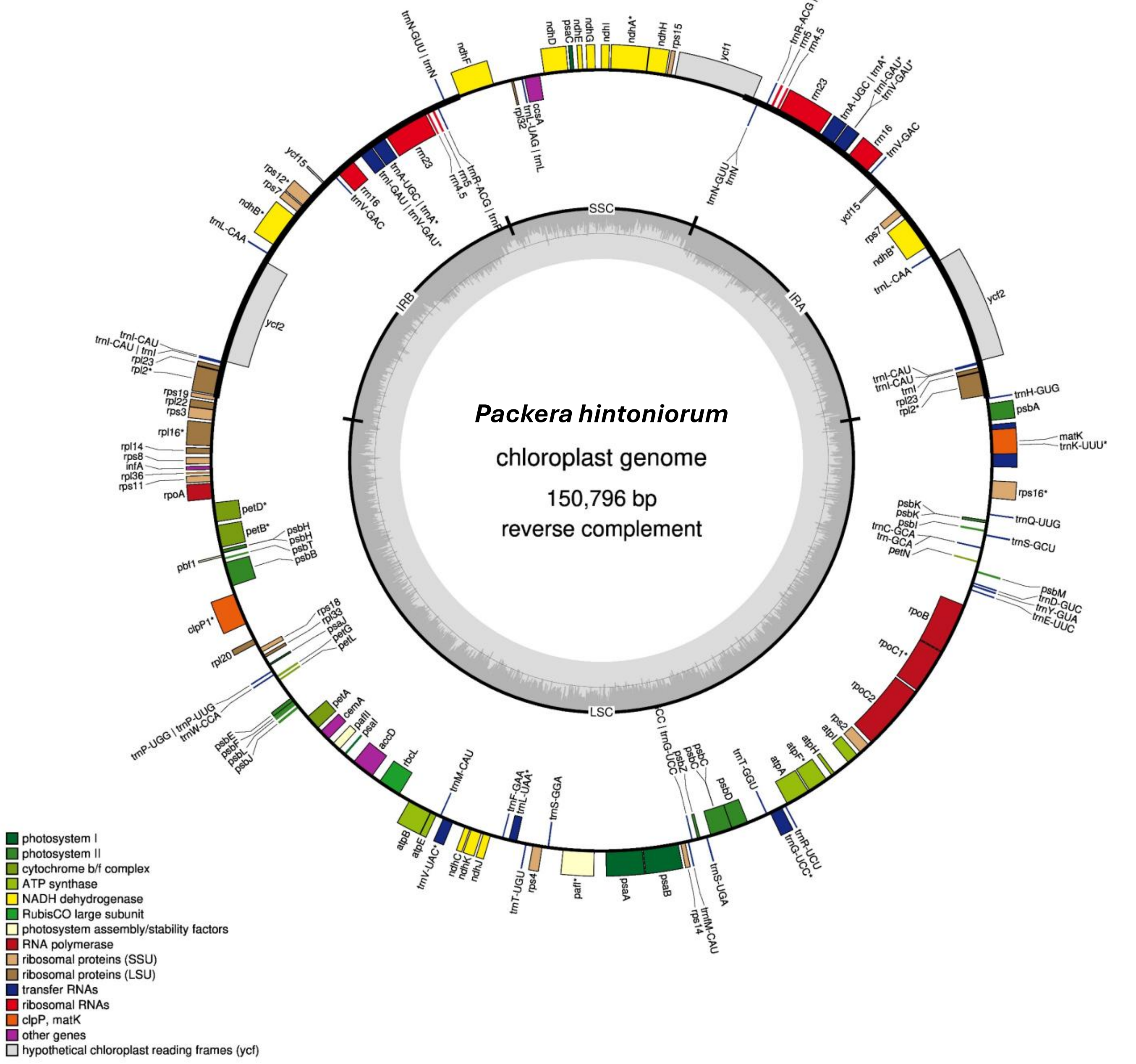

*Packera hintoniorum*
chloroplast genome
150,796 bp
reverse complement
SSC
IRA
LSC
IRB
trnN-GUU | trnN
ndhF
rpl32
trnL-UAG | trnL
ccsA
ndhD
psaC
ndhE
ndhG
ndhI
ndhA*
ndhH
rps15
ycf1
trnR-ACG | trnR
rrn5
rrn4.5
rrn23
trnA-UGC | trnA*
trnI-GAU*
trnV-GAU*
rrn16
trnV-GAC
trnN-GUU
trnN
ycf15
rps7
ndhB*
trnL-CAA
ycf2
trnI-CAU
trnI
rpl23
rpl2*
trnH-GUG
psbA
matK
trnK-UUU*
rps16*
psbK
psbI
trnQ-UUG
trnC-GCA
trn-GCA
trnS-GCU
petN
psbM
trnD-GUC
trnY-GUA
trnE-UUC
rpoB
rpoC1*
rpoC2
rps2
atpI
atpH
atpF*
atpA
trnR-UCU
trnG-UCC*
trnT-GGU
psbD
psbC
psbZ
trnG-UCC
trnS-UGA
trnfM-CAU
rps14
psaB
psaA
pafI*
rps4
trnT-UGU
trnS-GGA
trnF-GAA
trnL-UAA*
ndhJ
ndhK
ndhC
trnV-UAC*
trnM-CAU
atpE
atpB
rbcL
accD
psaI
pafII
cemA
petA
psbJ
psbL
psbF
psbE
trnP-UGG | trnP-UGG
trnW-CCA
rpl20
rps18
rpl33
psaJ
petG
petL
clpP1*
pbf1
psbH
psbT
psbB
petB*
petD*
rpoA
rps11
rpl36
infA
rps8
rpl14
rpl16*
rps3
rpl22
rps19
trnI-CAU | trnI
rrn16
trnV-GAC
trnI-GAU | trnV-GAU*
rps12*
photosystem I
photosystem II
cytochrome b/f complex
ATP synthase
NADH dehydrogenase
RubisCO large subunit
photosystem assembly/stability factors
RNA polymerase
ribosomal proteins (SSU)
ribosomal proteins (LSU)
transfer RNAs
ribosomal RNAs
clpP, matK
other genes
hypothetical chloroplast reading frames (ycf)

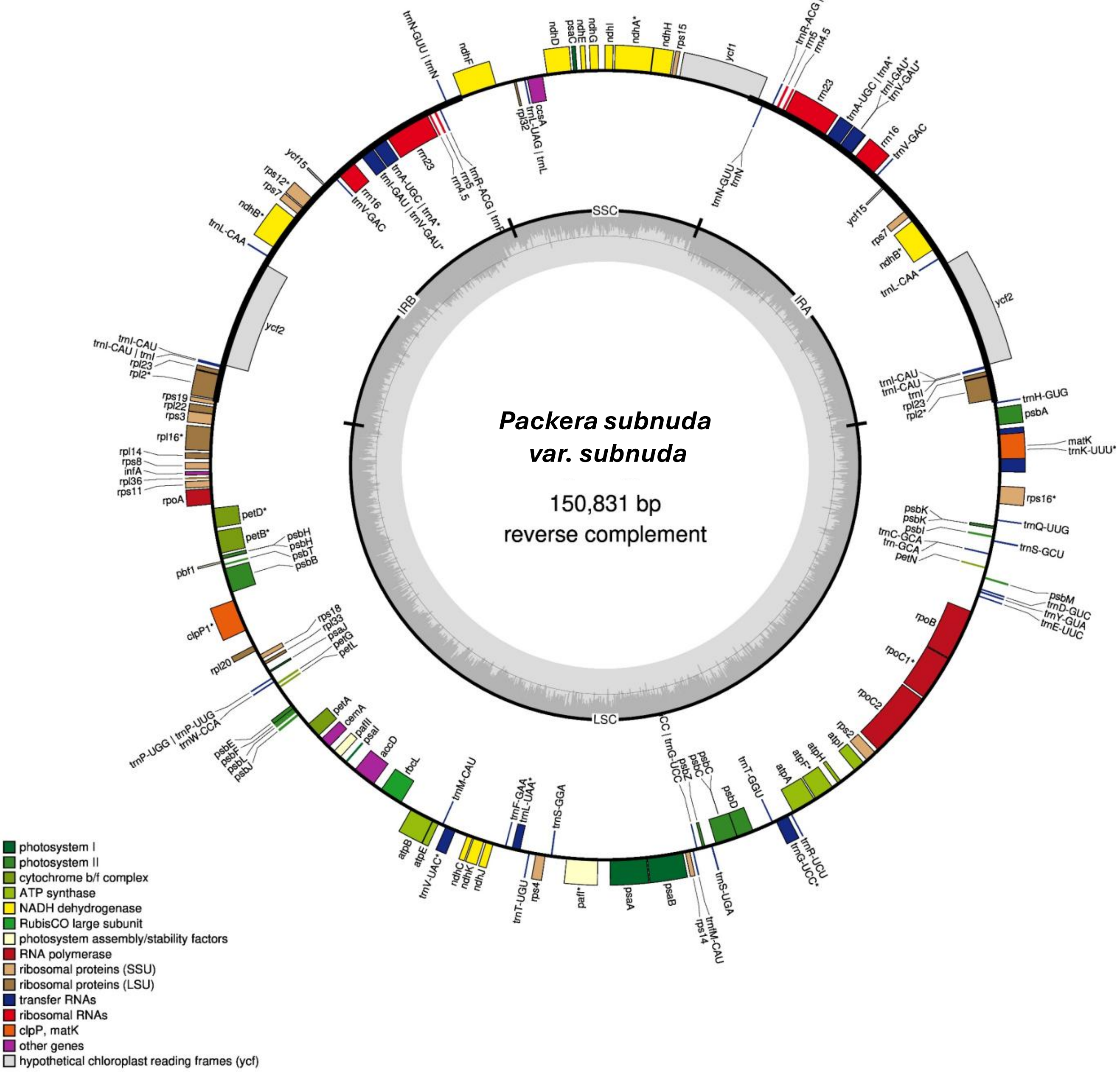
Packera subnuda
var. subnuda
150,831 bp
reverse complement
SSC
IRA
IRB
LSC
trnN-GUU | trnN
ndhF
ndhD
psaC
ndhE
ndhG
ndhI
ndhA*
ndhH
rps15
ycf1
rpl32
trnL-UAG | trnL
ccsA
trnR-ACG | trnR
rrn5
rrn4.5
rrn23
trnA-UGC | trnA*
trnI-GAU*
trnV-GAU*
rrn16
trnV-GAC
trnN-GUU
trnN
ycf15
rps7
ndhB*
trnL-CAA
ycf2
trnI-CAU
trnI-CAU
trnI
rpl23
rpl2*
trnH-GUG
psbA
matK
trnK-UUU*
rps16*
psbK
psbK
psbI
trnC-GCA
trn-GCA
petN
trnQ-UUG
trnS-GCU
psbM
trnD-GUC
trnY-GUA
trnE-UUC
rpoB
rpoC1*
rpoC2
rps2
atpI
atpH
atpF*
atpA
trnR-UCU
trnG-UCC*
trnT-GGU
psbD
psbC
psbC
psbZ
trnG-UCC
trnS-UGA
trnfM-CAU
rps14
psaB
psaA
pafI*
rps4
trnT-UGU
trnS-GGA
trnL-UAA*
trnF-GAA
ndhJ
ndhK
ndhC
trnV-UAC*
trnM-CAU
atpE
atpB
rbcL
accD
psaI
pafII
cemA
petA
psbJ
psbL
psbF
psbE
trnP-UGG | trnP-UUG
trnW-CCA
rpl20
rps18
rpl33
psaJ
petG
petL
clpP1*
pbf1
psbH
psbH
psbT
psbB
petB*
petD*
rpoA
rps11
rpl36
infA
rps8
rpl14
rpl16*
rps3
rpl22
rps19
rpl2*
rpl23
trnI-CAU | trnI
trnI-CAU
ycf2
trnL-CAA
ndhB*
rps7
rps12*
ycf15
rrn16
trnV-GAC
trnI-GAU | trnV-GAU*
trnA-UGC | trnA*
rrn23
rrn5
rrn4.5
trnR-ACG | trnR
photosystem I
photosystem II
cytochrome b/f complex
ATP synthase
NADH dehydrogenase
RubisCO large subunit
photosystem assembly/stability factors
RNA polymerase
ribosomal proteins (SSU)
ribosomal proteins (LSU)
transfer RNAs
ribosomal RNAs
clpP, matK
other genes
hypothetical chloroplast reading frames (ycf)

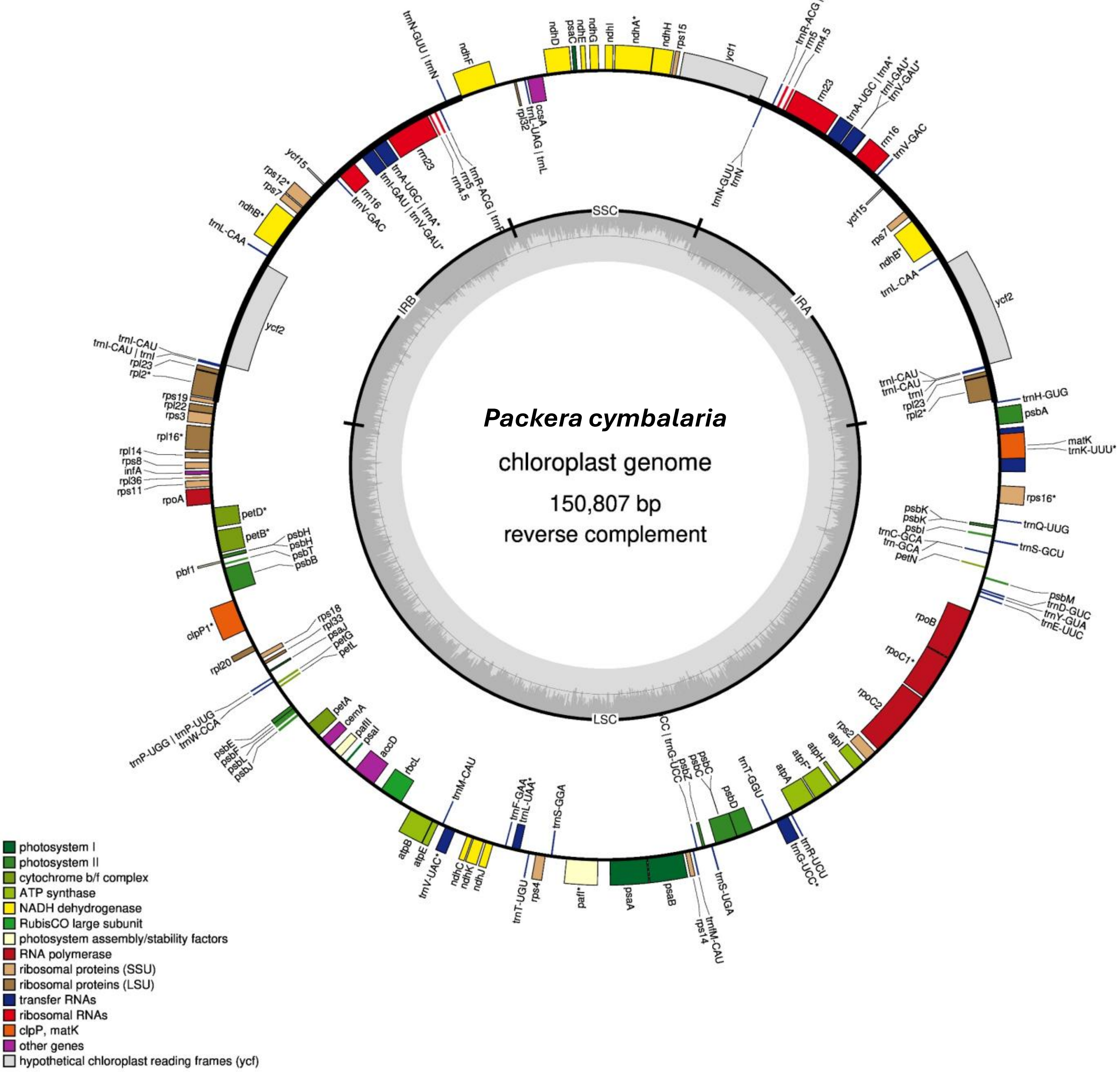

Packera cymbalaria
chloroplast genome
150,807 bp
reverse complement
SSC
IRA
IRB
LSC
photosystem I
photosystem II
cytochrome b/f complex
ATP synthase
NADH dehydrogenase
RubisCO large subunit
photosystem assembly/stability factors
RNA polymerase
ribosomal proteins (SSU)
ribosomal proteins (LSU)
transfer RNAs
ribosomal RNAs
clpP, matK
other genes
hypothetical chloroplast reading frames (ycf)
trnN-GUU | trnN
ndhF
ndhD
psaC
ndhE
ndhG
ndhI
ndhA*
ndhH
rps15
ycf1
rpl32
trnL-UAG | trnL
ccsA
trnR-ACG | trnR
rrn5
rrn4.5
rrn23
trnA-UGC | trnA*
trnI-GAU*
trnV-GAU*
rrn16
trnV-GAC
trnN-GUU
trnN
ycf15
rps7
ndhB*
trnL-CAA
ycf2
trnI-CAU
trnI-CAU
trnI
rpl23
rpl2*
trnH-GUG
psbA
matK
trnK-UUU*
rps16*
psbK
psbK
psbI
trnC-GCA
trn-GCA
petN
trnQ-UUG
trnS-GCU
psbM
trnD-GUC
trnY-GUA
trnE-UUC
rpoB
rpoC1*
rpoC2
rps2
atpI
atpH
atpF*
atpA
trnR-UCU
trnG-UCC*
trnT-GGU
psbD
psbC
psbC
psbZ
trnG-UCC
trnS-UGA
trnfM-CAU
rps14
psaB
psaA
pafI*
rps4
trnT-UGU
trnS-GGA
trnF-GAA
trnL-UAA*
ndhJ
ndhK
ndhC
trnV-UAC*
trnM-CAU
atpE
atpB
rbcL
accD
psaI
pafII
cemA
petA
psbJ
psbL
psbF
psbE
trnP-UGG | trnP-UGG
trnW-CCA
rpl20
rps18
rpl33
psaJ
petG
petL
clpP1*
pbf1
psbH
psbH
psbT
psbB
petB*
petD*
rpoA
rps11
rpl36
infA
rps8
rpl14
rpl16*
rps3
rpl22
rps19
rpl2*
rpl23
trnI-CAU | trnI
trnI-CAU
ycf2
trnL-CAA
ndhB*
rps7
rps12*
ycf15
rrn16
trnV-GAC
trnA-UGC | trnA*
trnI-GAU | trnV-GAU*
rrn23
rrn5
rrn4.5
trnR-ACG | trnR

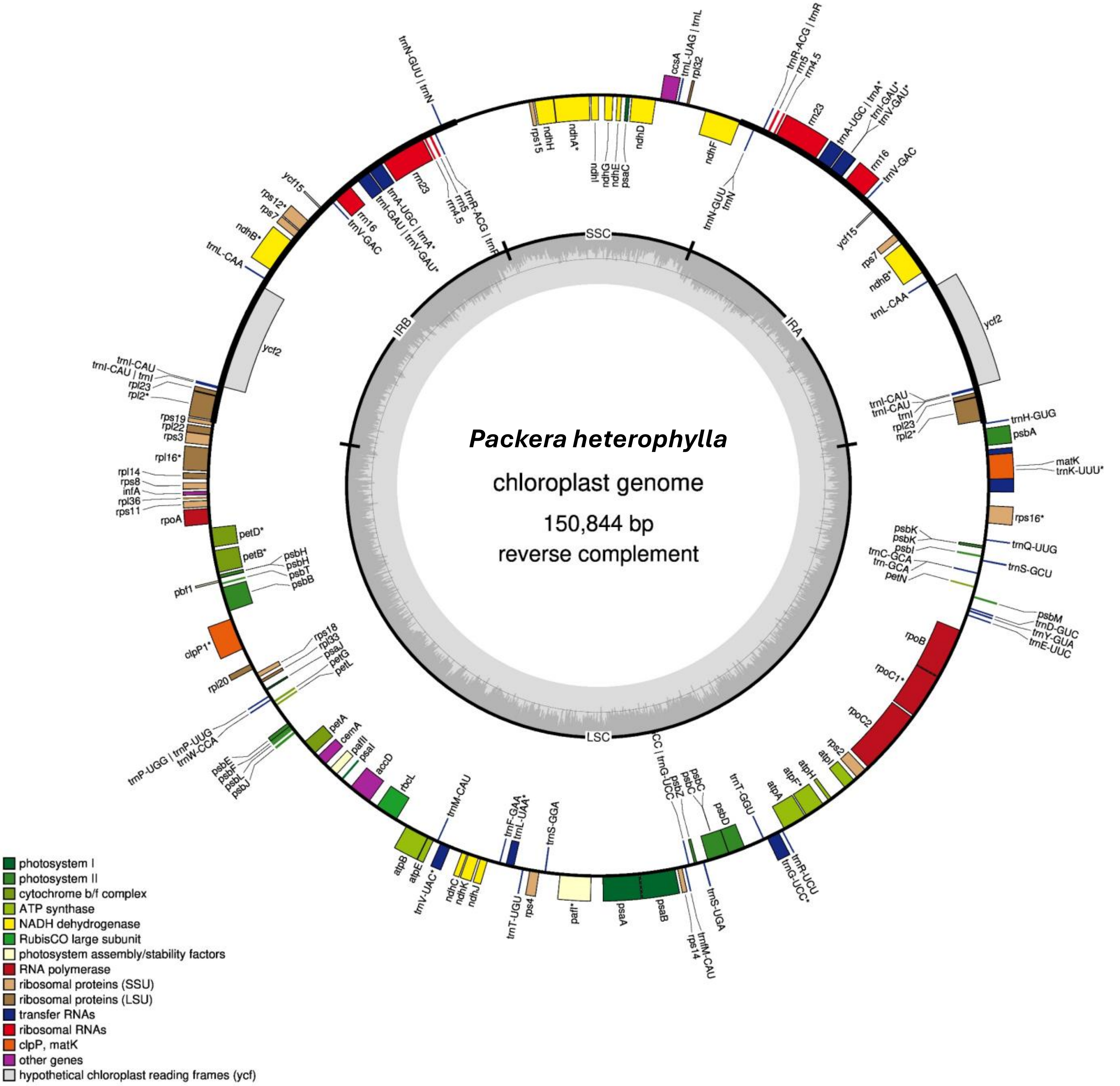
Packera heterophylla
chloroplast genome
150,844 bp
reverse complement
SSC
IRA
IRB
LSC
ndhF
ccsA
trnL-UAG | trnL
rpl32
ndhD
psaC
ndhE
ndhG
ndhI
ndhA*
ndhH
rps15
trnR-ACG | trnR
rrn5
rrn4.5
rrn23
trnA-UGC | trnA*
trnI-GAU*
trnV-GAU*
rrn16
trnV-GAC
trnN-GUU
trnN
ycf15
rps7
ndhB*
trnL-CAA
ycf2
trnI-CAU
trnI
rpl23
rpl2*
trnH-GUG
psbA
matK
trnK-UUU*
rps16*
psbK
psbI
trnQ-UUG
trnC-GCA
trn-GCA
trnS-GCU
petN
psbM
trnD-GUC
trnY-GUA
trnE-UUC
rpoB
rpoC1*
rpoC2
rps2
atpI
atpH
atpF*
atpA
trnR-UCU
trnG-UCC*
trnT-GGU
psbD
psbC
psbZ
trnG-UCC
trnS-UGA
trnfM-CAU
rps14
psaB
psaA
paf*
rps4
trnT-UGU
trnS-GGA
trnL-UAA*
trnF-GAA
ndhJ
ndhK
ndhC
trnV-UAC*
trnM-CAU
atpE
atpB
rbcL
accD
psaI
pafII
cemA
petA
psbJ
psbL
psbF
psbE
trnW-CCA
trnP-UGG | trnP-UGG
petL
petG
psaJ
rpl33
rps18
rpl20
clpP1*
pbf1
psbB
psbT
psbH
petB*
petD*
rpoA
rps11
rpl36
infA
rps8
rpl14
rpl16*
rps3
rpl22
rps19
trnI-CAU | trnI
rps12*
trnN-GUU | trnN
rrn16
trnV-GAC
trnI-GAU | trnV-GAU*
trnR-ACG | trnR
photosystem I
photosystem II
cytochrome b/f complex
ATP synthase
NADH dehydrogenase
RubisCO large subunit
photosystem assembly/stability factors
RNA polymerase
ribosomal proteins (SSU)
ribosomal proteins (LSU)
transfer RNAs
ribosomal RNAs
clpP, matK
other genes
hypothetical chloroplast reading frames (ycf)

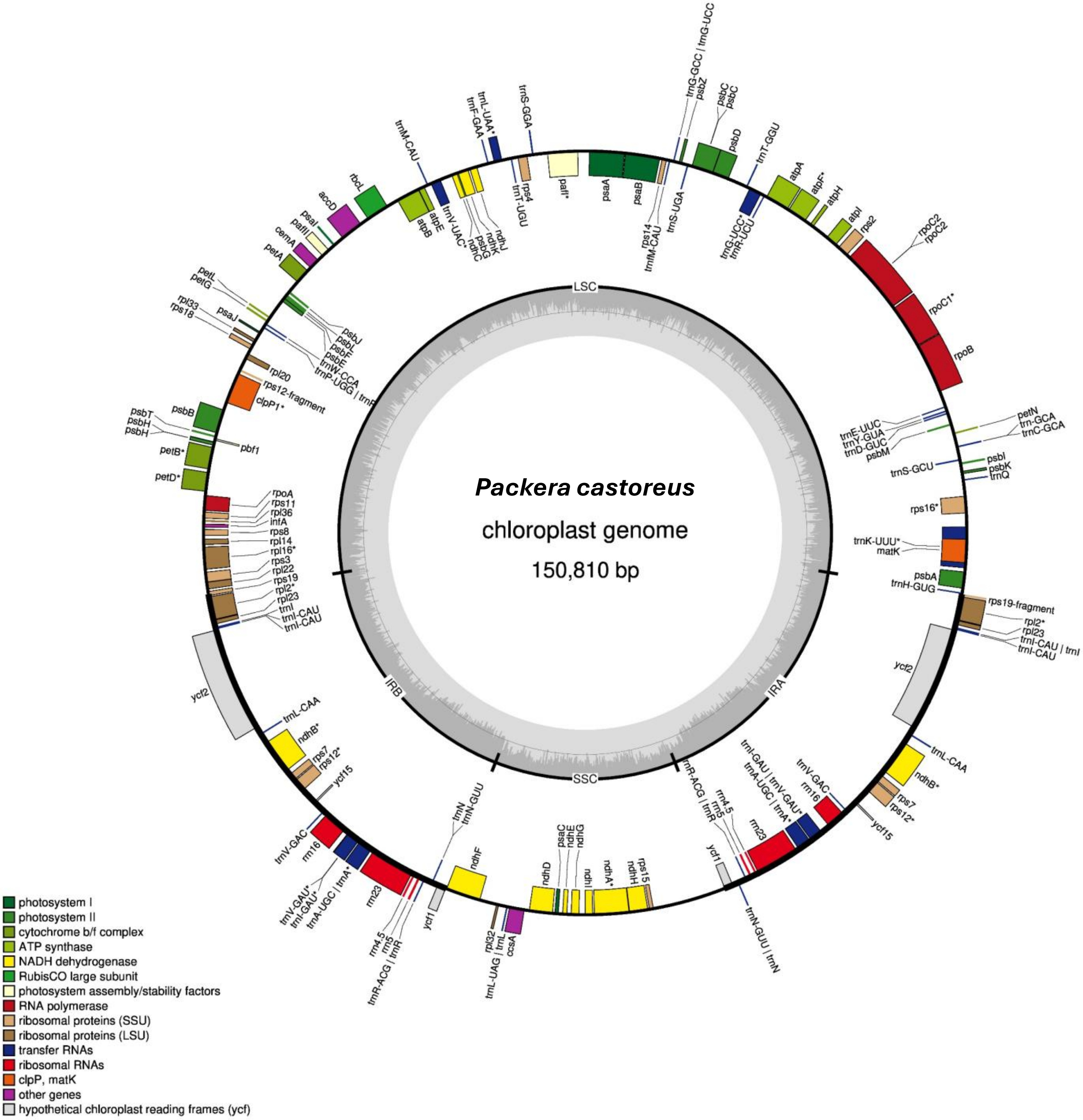

Packera castoreus
chloroplast genome
150,810 bp
LSC
IRA
SSC
IRB
psaA
psaB
pafI*
rps4
trnT-UGU
trnS-GGA
trnL-UAA*
trnF-GAA
trnM-CAU
ndhJ
ndhK
ndhC
psbG
trnV-UAC*
atpE
atpB
rbcL
accD
psaI
pafII
cemA
petA
petL
petG
rpl33
rps18
psaJ
psbJ
psbL
psbF
psbE
trnW-CCA
trnP-UGG | trnP
rpl20
rps12-fragment
clpP1*
psbB
psbT
psbH
pbf1
petB*
petD*
rpoA
rps11
rpl36
infA
rps8
rpl14
rpl16*
rps3
rpl22
rps19
rpl2*
rpl23
trnI
trnI-CAU
ycf2
trnL-CAA
ndhB*
rps7
rps12*
ycf15
trnV-GAC
rrn16
trnV-GAU*
trnI-GAU*
trnA-UGC | trnA*
rrn23
rrn4.5
rrn5
trnR-ACG | trnR
ycf1
trnN
trnN-GUU
ndhF
rpl32
trnL-UAG | trnL
ccsA
ndhD
psaC
ndhE
ndhG
ndhI
ndhA*
ndhH
rps15
trnN-GUU | trnN
trnI-GAU | trnV-GAU*
rps14
trnfM-CAU
trnS-UGA
trnG-GCC | trnG-UCC
psbZ
psbC
psbD
trnT-GGU
trnG-UCC*
trnR-UCU
atpA
atpF*
atpH
atpI
rps2
rpoC2
rpoC1*
rpoB
petN
trn-GCA
trnC-GCA
trnE-UUC
trnY-GUA
trnD-GUC
psbM
psbI
psbK
trnQ
trnS-GCU
rps16*
trnK-UUU*
matK
psbA
trnH-GUG
rps19-fragment
trnI-CAU | trnI
photosystem I
photosystem II
cytochrome b/f complex
ATP synthase
NADH dehydrogenase
RubisCO large subunit
photosystem assembly/stability factors
RNA polymerase
ribosomal proteins (SSU)
ribosomal proteins (LSU)
transfer RNAs
ribosomal RNAs
clpP, matK
other genes
hypothetical chloroplast reading frames (ycf)

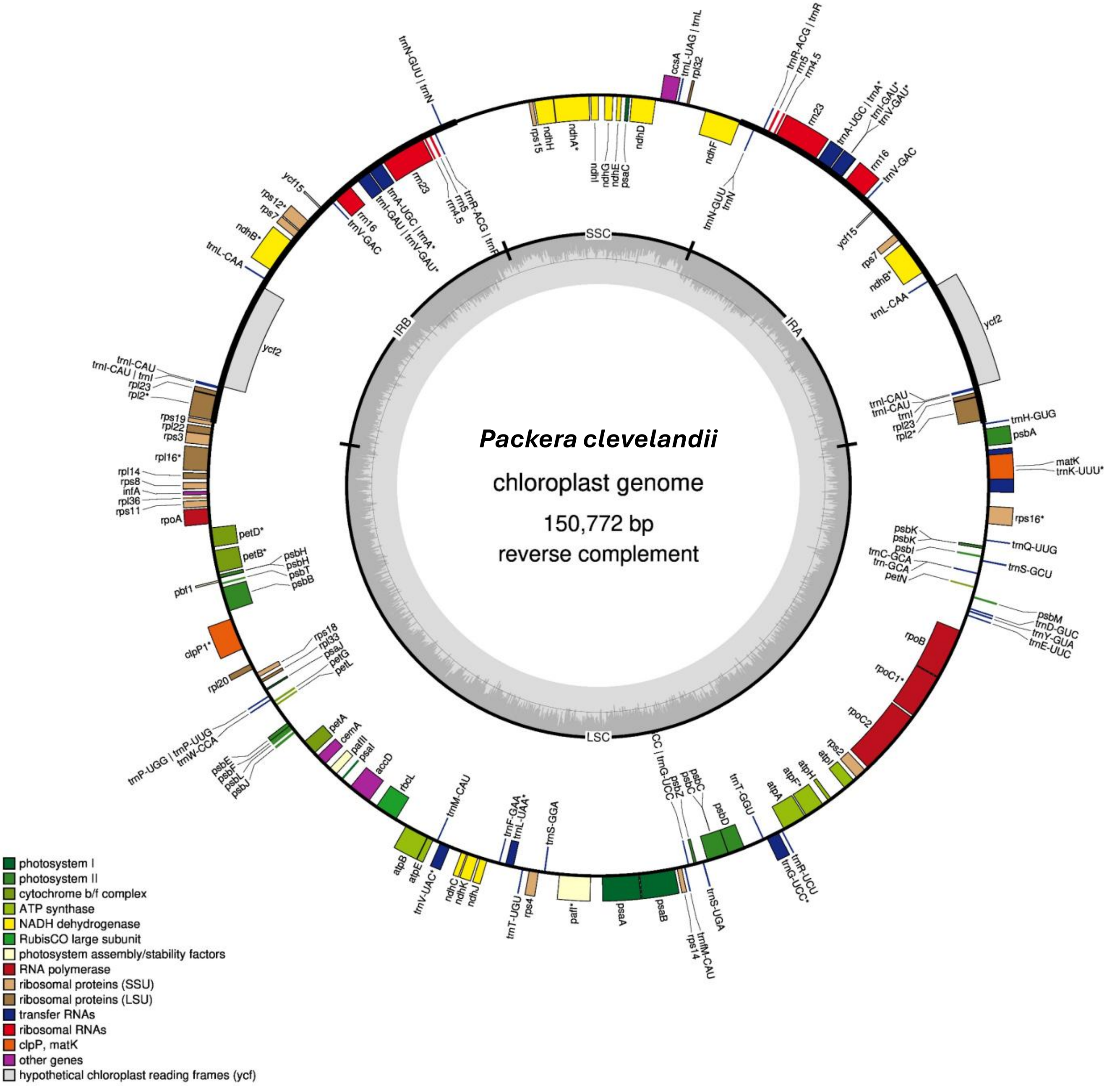

*Packera clevelandii*
chloroplast genome
150,772 bp
reverse complement
SSC
IRA
IRB
LSC
photosystem I
photosystem II
cytochrome b/f complex
ATP synthase
NADH dehydrogenase
RubisCO large subunit
photosystem assembly/stability factors
RNA polymerase
ribosomal proteins (SSU)
ribosomal proteins (LSU)
transfer RNAs
ribosomal RNAs
clpP, matK
other genes
hypothetical chloroplast reading frames (ycf)

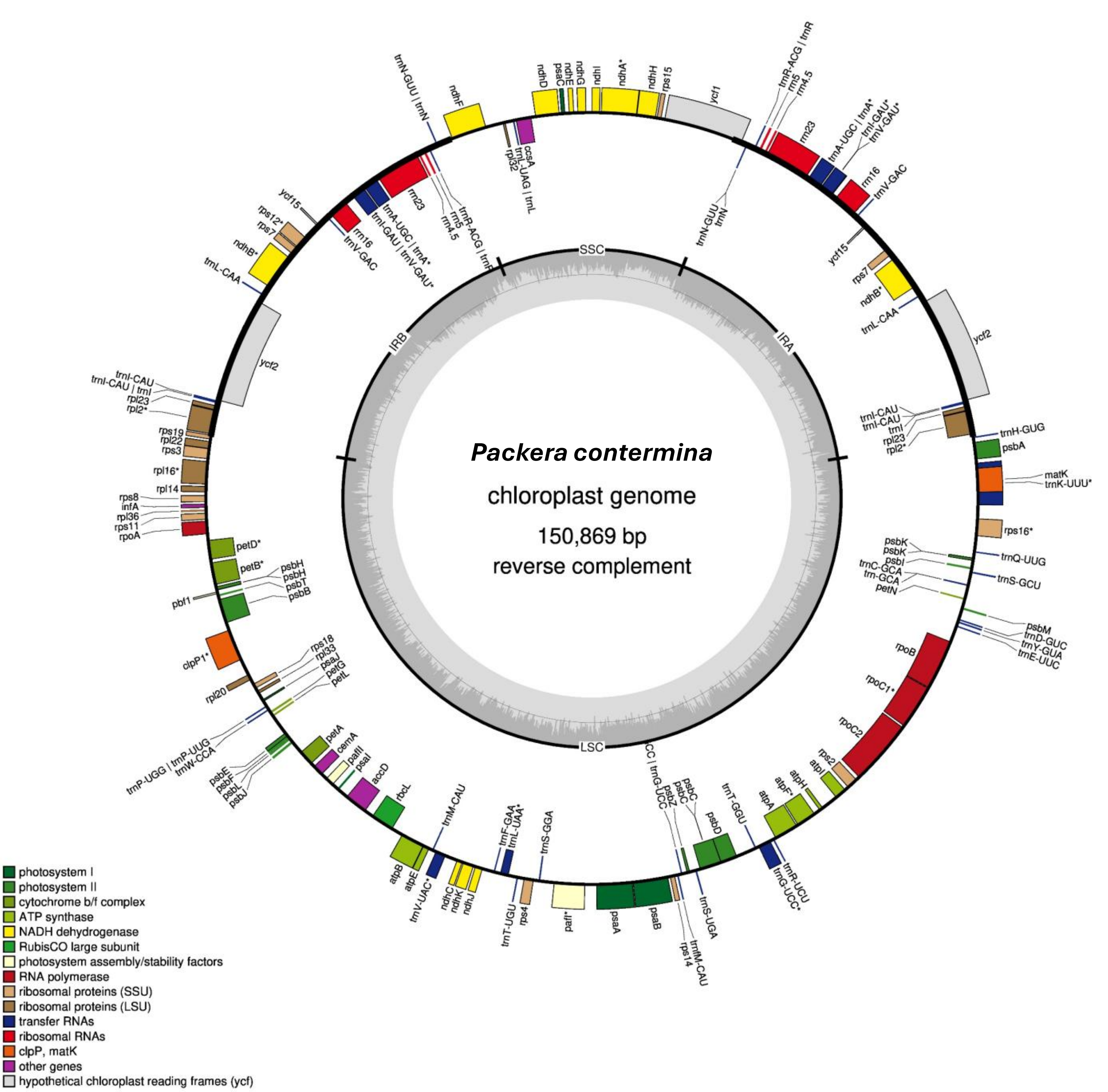
*Packera contermina*
chloroplast genome
150,869 bp
reverse complement
SSC
IRA
IRB
LSC
photosystem I
photosystem II
cytochrome b/f complex
ATP synthase
NADH dehydrogenase
RubisCO large subunit
photosystem assembly/stability factors
RNA polymerase
ribosomal proteins (SSU)
ribosomal proteins (LSU)
transfer RNAs
ribosomal RNAs
clpP, matK
other genes
hypothetical chloroplast reading frames (ycf)

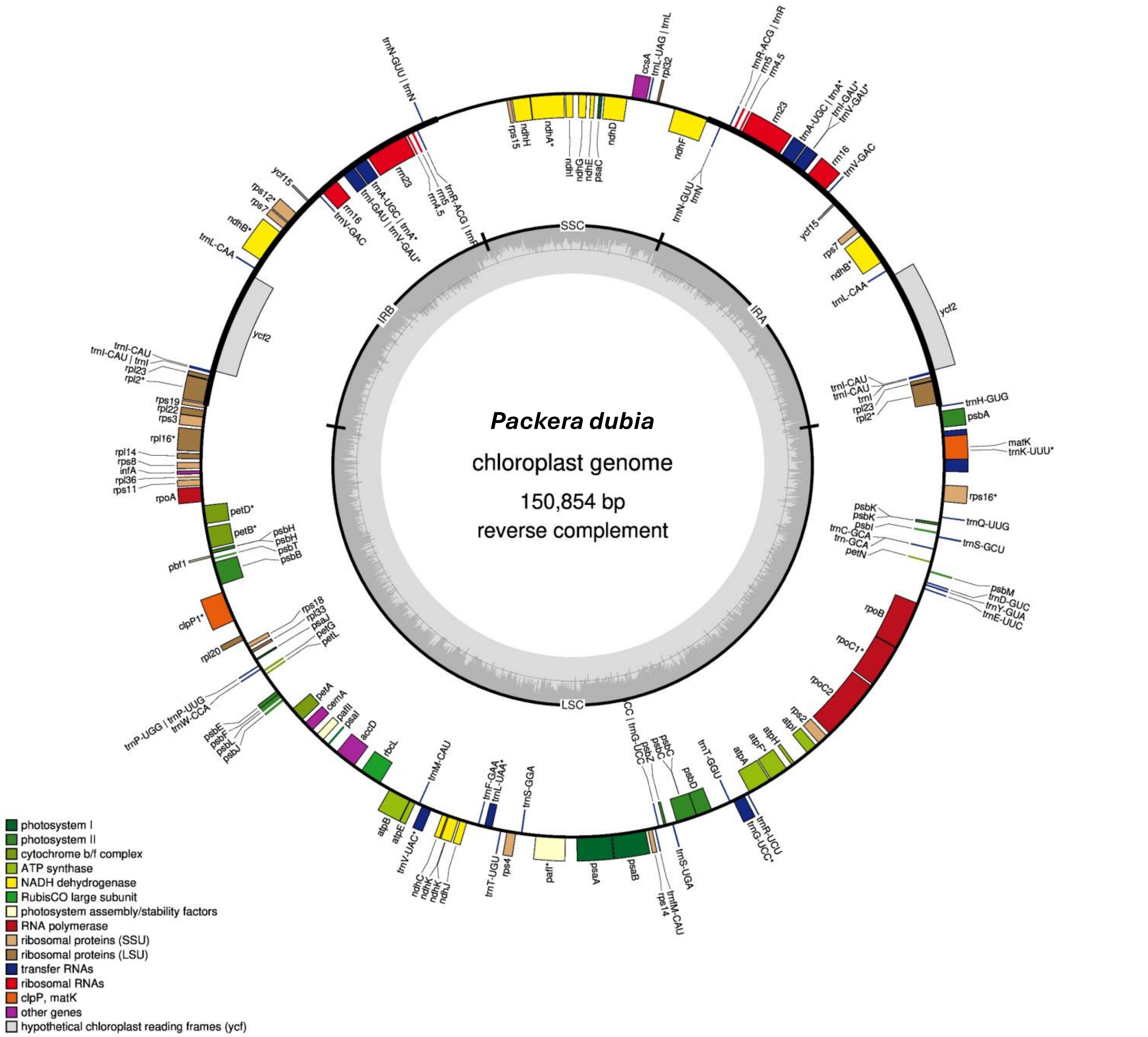
Packera dubia
chloroplast genome
150,854 bp
reverse complement
SSC
IRA
IRB
LSC
trnN-GUU | trnN
ccsA
trnL-UAG | trnL
rpl32
trnR-ACG | trnR
rrn5
rrn4.5
rrn23
trnA-UGC | trnA*
trnI-GAU*
trnV-GAU*
rrn16
trnV-GAC
rps15
ndhH
ndhA*
ndhI
ndhG
ndhE
psaC
ndhD
ndhF
trnN-GUU
trnN
ycf15
rps12*
rps7
ndhB*
trnL-CAA
ycf2
rrn16
trnV-GAC
trnA-UGC | trnA*
trnI-GAU | trnV-GAU*
rrn23
rrn5
rrn4.5
trnR-ACG | trnR
trnI-CAU
trnI-CAU | trnI
rpl23
rpl2*
rps19
rpl22
rps3
rpl16*
rpl14
rps8
infA
rpl36
rps11
rpoA
petD*
petB*
psbH
psbH
psbT
psbB
pbf1
clpP1*
rps18
rpl33
psaJ
petG
petL
rpl20
trnP-UGG | trnP-UGG
trnW-CCA
psbE
psbF
psbL
psbJ
petA
cemA
pafII
psaI
accD
rbcL
trnM-CAU
atpB
atpE
trnV-UAC*
ndhC
ndhK
ndhK
ndhJ
trnF-GAA
trnL-UAA*
trnT-UGU
rps4
trnS-GGA
pafI*
psaA
psaB
rps14
trnfM-CAU
trnS-UGA
trnG-UCC
psbZ
psbC
psbC
psbD
trnT-GGU
trnR-UCU
trnG-UCC*
atpA
atpF*
atpH
atpI
rps2
rpoC2
rpoC1*
rpoB
psbM
trnD-GUC
trnY-GUA
trnE-UUC
psbK
psbK
psbI
trnC-GCA
trn-GCA
petN
trnQ-UUG
trnS-GCU
rps16*
matK
trnK-UUU*
psbA
trnH-GUG
trnI-CAU
trnI-CAU
trnI
rpl23
rpl2*
ycf2
trnL-CAA
ndhB*
rps7
ycf15
photosystem I
photosystem II
cytochrome b/f complex
ATP synthase
NADH dehydrogenase
RubisCO large subunit
photosystem assembly/stability factors
RNA polymerase
ribosomal proteins (SSU)
ribosomal proteins (LSU)
transfer RNAs
ribosomal RNAs
clpP, matK
other genes
hypothetical chloroplast reading frames (ycf)

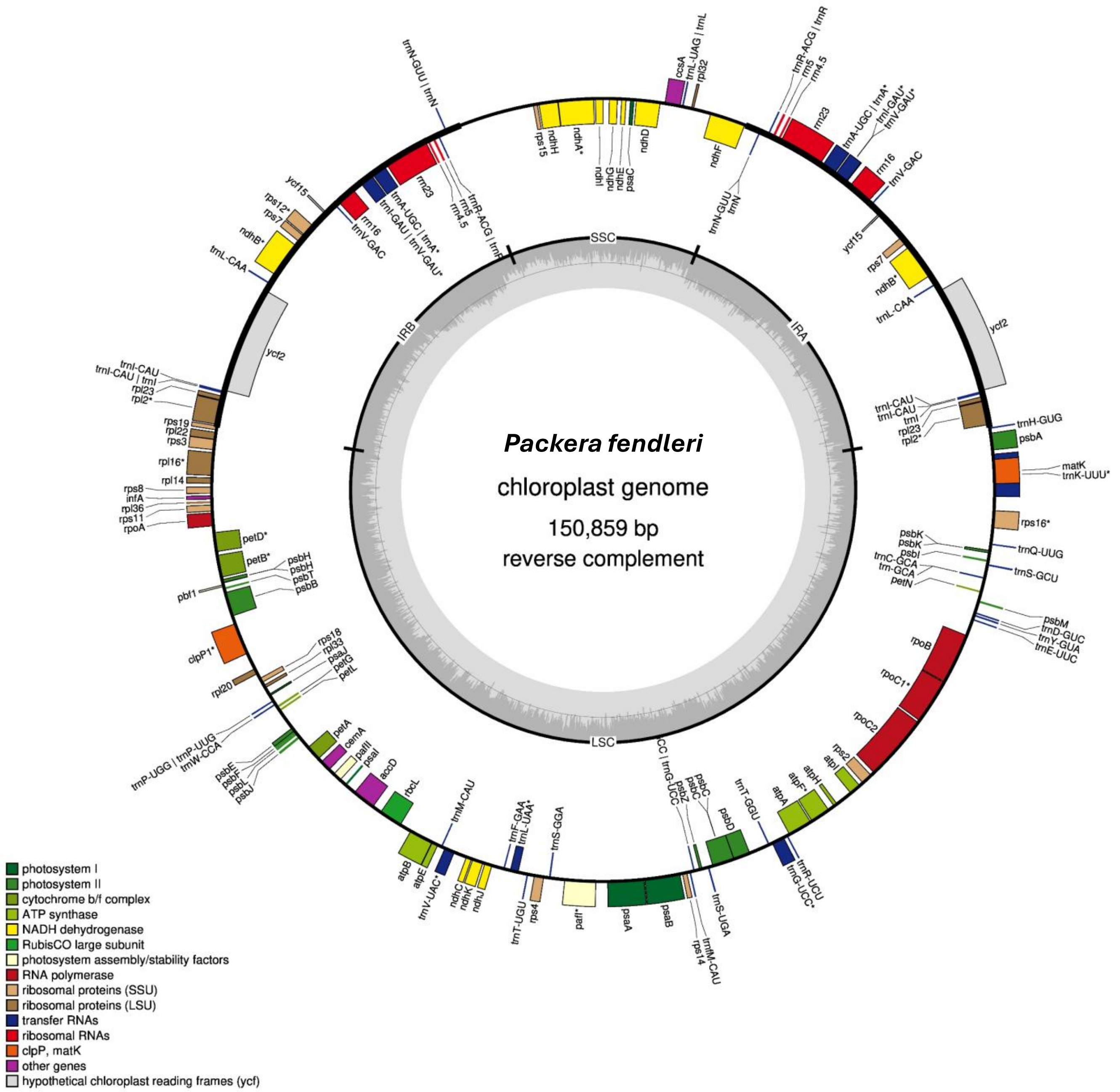

Packera fendleri
chloroplast genome
150,859 bp
reverse complement
SSC
IRA
IRB
LSC
photosystem I
photosystem II
cytochrome b/f complex
ATP synthase
NADH dehydrogenase
RubisCO large subunit
photosystem assembly/stability factors
RNA polymerase
ribosomal proteins (SSU)
ribosomal proteins (LSU)
transfer RNAs
ribosomal RNAs
clpP, matK
other genes
hypothetical chloroplast reading frames (ycf)
trnH-GUG
psbA
matK
trnK-UUU*
rps16*
trnQ-UUG
trnS-GCU
psbM
trnD-GUC
trnY-GUA
trnE-UUC
rpoB
rpoC1*
rpoC2
rps2
atpI
atpH
atpF*
atpA
trnT-GGU
psbD
psbC
psbZ
trnS-UGA
trnfM-CAU
rps14
psaB
psaA
paf*
rps4
trnT-UGU
trnS-GGA
trnF-GAA
trnL-UAA*
ndhJ
ndhK
ndhC
trnV-UAC*
atpE
atpB
trnM-CAU
rbcL
accD
psaI
pafII
cemA
petA
psbJ
psbL
psbF
psbE
trnP-UGG | trnP-UGG
trnW-CCA
rpl20
clpP1*
rps18
rpl33
psaJ
petG
petL
pbf1
psbB
psbT
psbH
petB*
petD*
rpoA
rps11
rpl36
infA
rps8
rpl14
rpl16*
rps3
rpl22
rps19
rpl2*
rpl23
trnI-CAU
ycf2
trnL-CAA
ndhB*
rps7
rps12*
ycf15
rrn16
trnV-GAC
rrn23
rrn5
rrn4.5
trnN-GUU | trnN
rps15
ndhH
ndhA*
ndhI
ndhG
ndhE
psaC
ndhD
ccsA
trnL-UAG | trnL
rpl32
ndhF
trnN-GUU
trnR-ACG | trnR

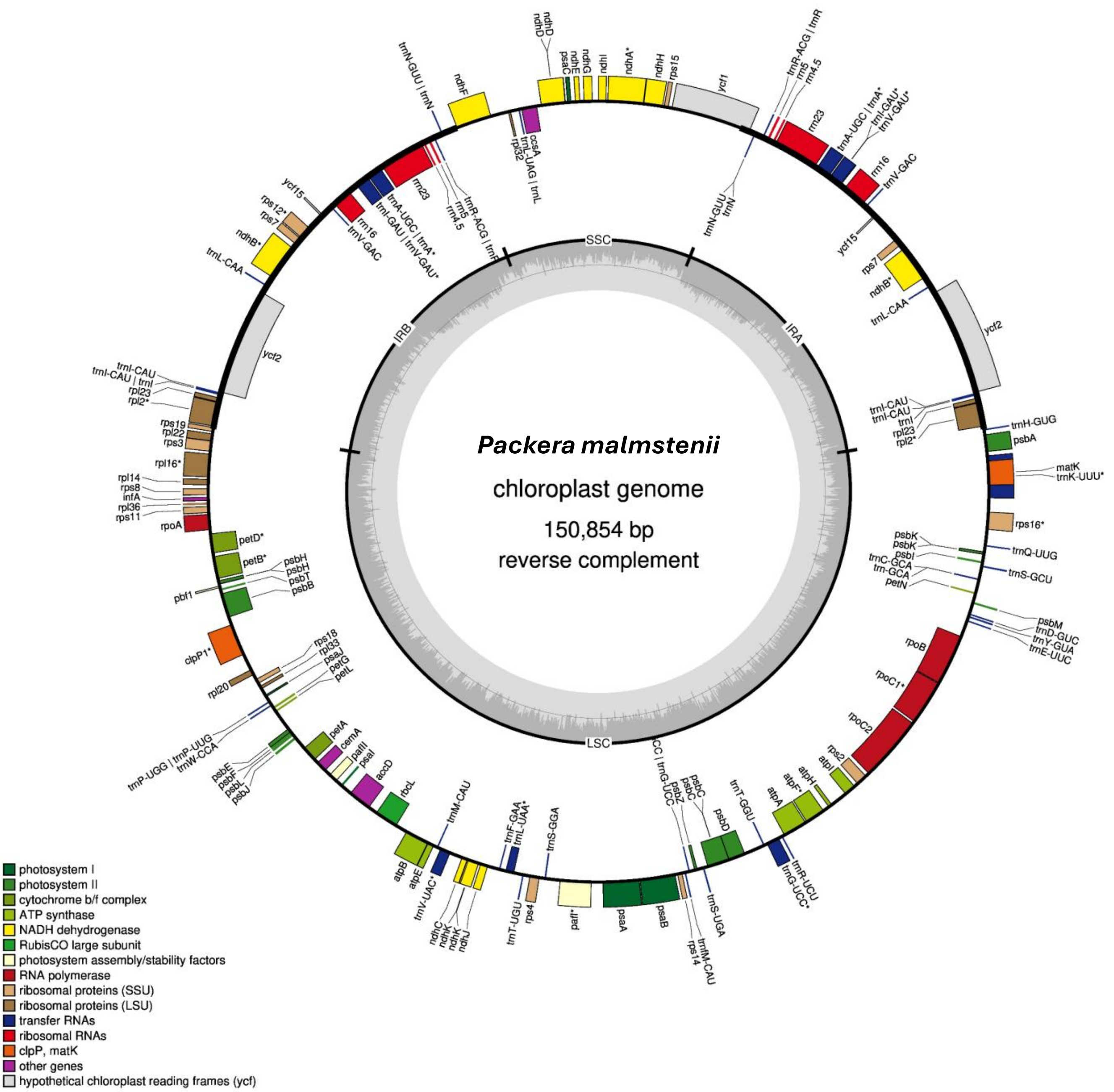
Packera malmstenii
chloroplast genome
150,854 bp
reverse complement
SSC
IRA
LSC
IRB
photosystem I
photosystem II
cytochrome b/f complex
ATP synthase
NADH dehydrogenase
RubisCO large subunit
photosystem assembly/stability factors
RNA polymerase
ribosomal proteins (SSU)
ribosomal proteins (LSU)
transfer RNAs
ribosomal RNAs
clpP, matK
other genes
hypothetical chloroplast reading frames (ycf)
ndhD
ndhD
psaC
ndhE
ndhG
ndhI
ndhA*
ndhH
rps15
ycf1
ndhF
trnN-GUU | trnN
ccsA
trnL-UAG | trnL
rpl32
trnR-ACG | trnR
rrn5
rrn4.5
rrn23
trnA-UGC | trnA*
trnI-GAU*
trnV-GAU*
rrn16
trnV-GAC
trnN-GUU
trnN
ycf15
rps7
ndhB*
trnL-CAA
ycf2
trnI-CAU
trnI-CAU
trnI
rpl23
rpl2*
trnH-GUG
psbA
matK
trnK-UUU*
rps16*
psbK
psbK
psbI
trnC-GCA
trn-GCA
petN
trnQ-UUG
trnS-GCU
psbM
trnD-GUC
trnY-GUA
trnE-UUC
rpoB
rpoC1*
rpoC2
rps2
atpI
atpH
atpF*
atpA
trnT-GGU
trnR-UCU
trnG-UCC*
psbD
psbC
psbC
psbZ
trnS-UGA
trnfM-CAU
rps14
psaB
psaA
pafI*
rps4
trnT-UGU
trnS-GGA
trnF-GAA
trnL-UAA*
trnM-CAU
trnV-UAC*
ndhC
ndhK
ndhK
ndhJ
atpE
atpB
rbcL
accD
psaI
pafII
cemA
petA
psbE
psbF
psbL
psbJ
trnP-UGG | trnP-UUG
trnW-CCA
rpl20
clpP1*
rps18
rpl33
psaJ
petG
petL
pbf1
psbH
psbH
psbT
psbB
petB*
petD*
rpoA
rps11
rpl36
infA
rps8
rpl14
rpl16*
rps3
rpl22
rps19
rpl2*
rpl23
trnI-CAU | trnI
trnI-CAU
ycf2
trnL-CAA
ndhB*
rps7
rps12*
ycf15
rrn16
trnV-GAC
trnA-UGC | trnA*
trnI-GAU | trnV-GAU*
rrn23
rrn5
rrn4.5
trnR-ACG | trnR

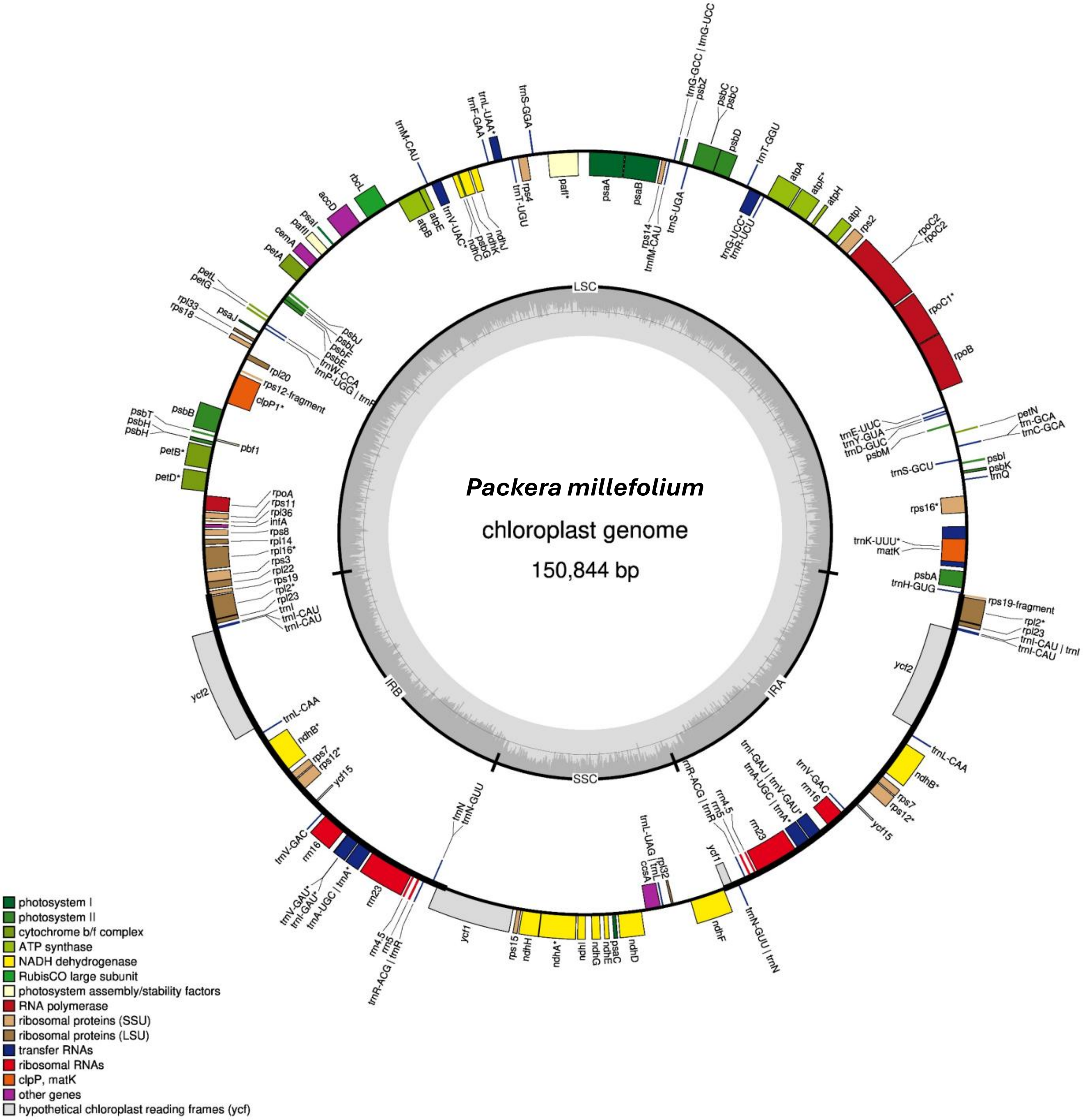
Packera millefolium
chloroplast genome
150,844 bp
LSC
IRB
IRA
SSC
trnG-GCC | trnG-UCC
psbZ
psbC
psbC
psbD
trnT-GGU
atpA
atpF*
atpH
atpI
rps2
rpoC2
rpoC2
rpoC1*
rpoB
trnS-GGA
trnL-UAA*
trnF-GAA
trnM-CAU
rbcL
accD
psaI
pafII
cemA
petA
psaA
psaB
pafI*
rps4
trnT-UGU
ndhJ
ndhK
ndhC
psbG
trnV-UAC*
atpE
atpB
rps14
trnfM-CAU
trnS-UGA
trnG-UCC*
trnR-UCU
petL
petG
rpl33
rps18
psaJ
psbJ
psbL
psbF
psbE
trnW-CCA
trnP-UGG | trnP
rpl20
rps12-fragment
clpP1*
psbB
psbT
psbH
psbH
pbf1
petB*
petD*
rpoA
rps11
rpl36
infA
rps8
rpl14
rpl16*
rps3
rpl22
rps19
rpl2*
rpl23
trnI
trnI-CAU
trnI-CAU
ycf2
trnL-CAA
ndhB*
rps7
rps12*
ycf15
trnV-GAC
rrn16
trnV-GAU*
trnI-GAU*
trnA-UGC | trnA*
rrn23
rrn4.5
rrn5
trnR-ACG | trnR
trnN
trnN-GUU
ycf1
rps15
ndhH
ndhA*
ndhI
ndhG
ndhE
psaC
ndhD
trnL-UAG | trnL
ccsA
rpl32
ndhF
trnN-GUU | trnN
ycf1
trnR-ACG | trnR
rrn4.5
rrn5
rrn23
trnI-GAU | trnV-GAU*
trnA-UGC | trnA*
trnV-GAC
rrn16
ycf15
rps12*
rps7
ndhB*
trnL-CAA
ycf2
trnI-CAU | trnI
trnI-CAU
rpl23
rpl2*
rps19-fragment
trnH-GUG
psbA
matK
trnK-UUU*
rps16*
trnQ
psbK
psbI
trnS-GCU
psbM
trnD-GUC
trnY-GUA
trnE-UUC
petN
trn-GCA
trnC-GCA
photosystem I
photosystem II
cytochrome b/f complex
ATP synthase
NADH dehydrogenase
RubisCO large subunit
photosystem assembly/stability factors
RNA polymerase
ribosomal proteins (SSU)
ribosomal proteins (LSU)
transfer RNAs
ribosomal RNAs
clpP, matK
other genes
hypothetical chloroplast reading frames (ycf)

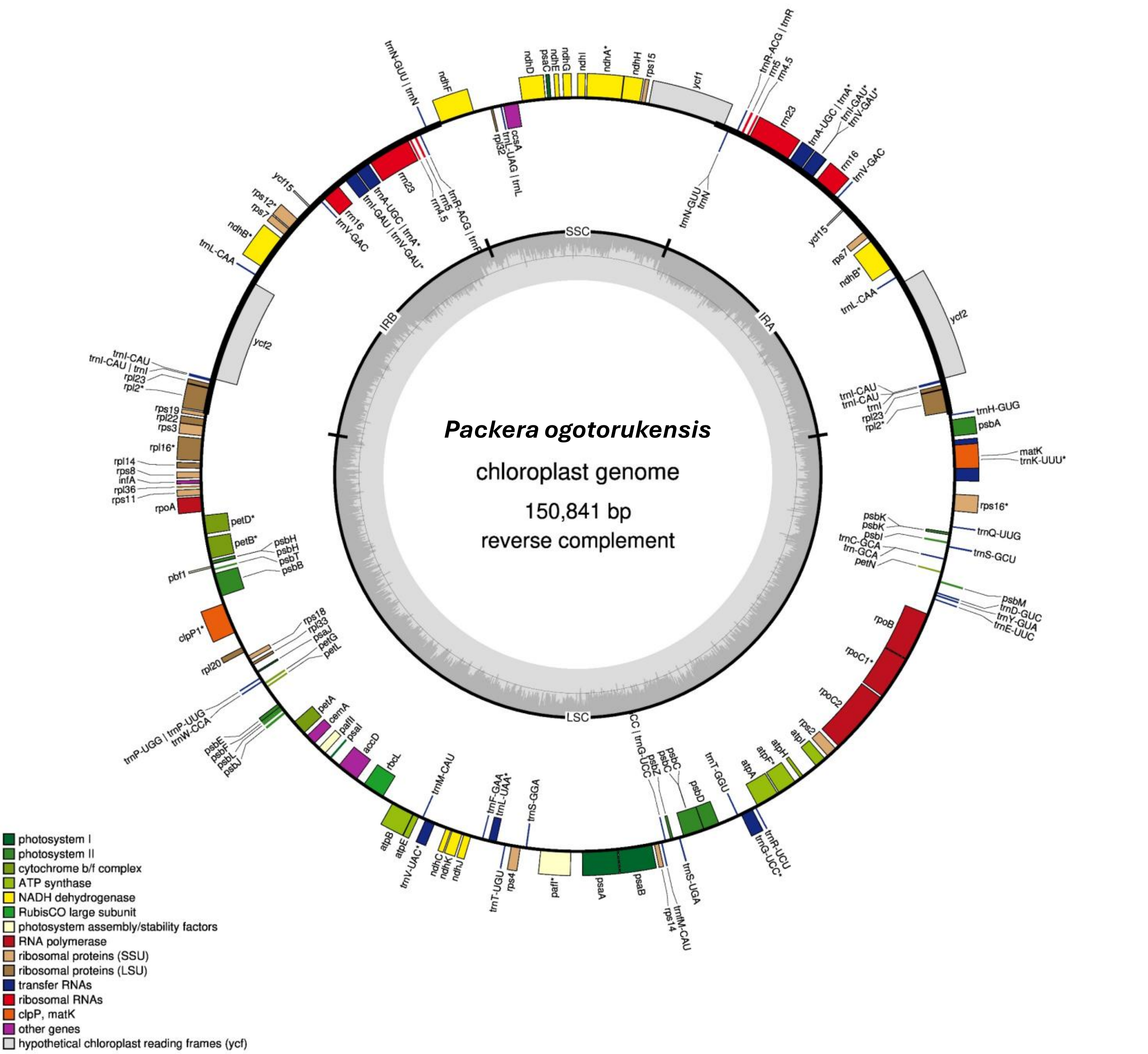
Packera ogotorukensis
chloroplast genome
150,841 bp
reverse complement
SSC
IRA
IRB
LSC
photosystem I
photosystem II
cytochrome b/f complex
ATP synthase
NADH dehydrogenase
RubisCO large subunit
photosystem assembly/stability factors
RNA polymerase
ribosomal proteins (SSU)
ribosomal proteins (LSU)
transfer RNAs
ribosomal RNAs
clpP, matK
other genes
hypothetical chloroplast reading frames (ycf)

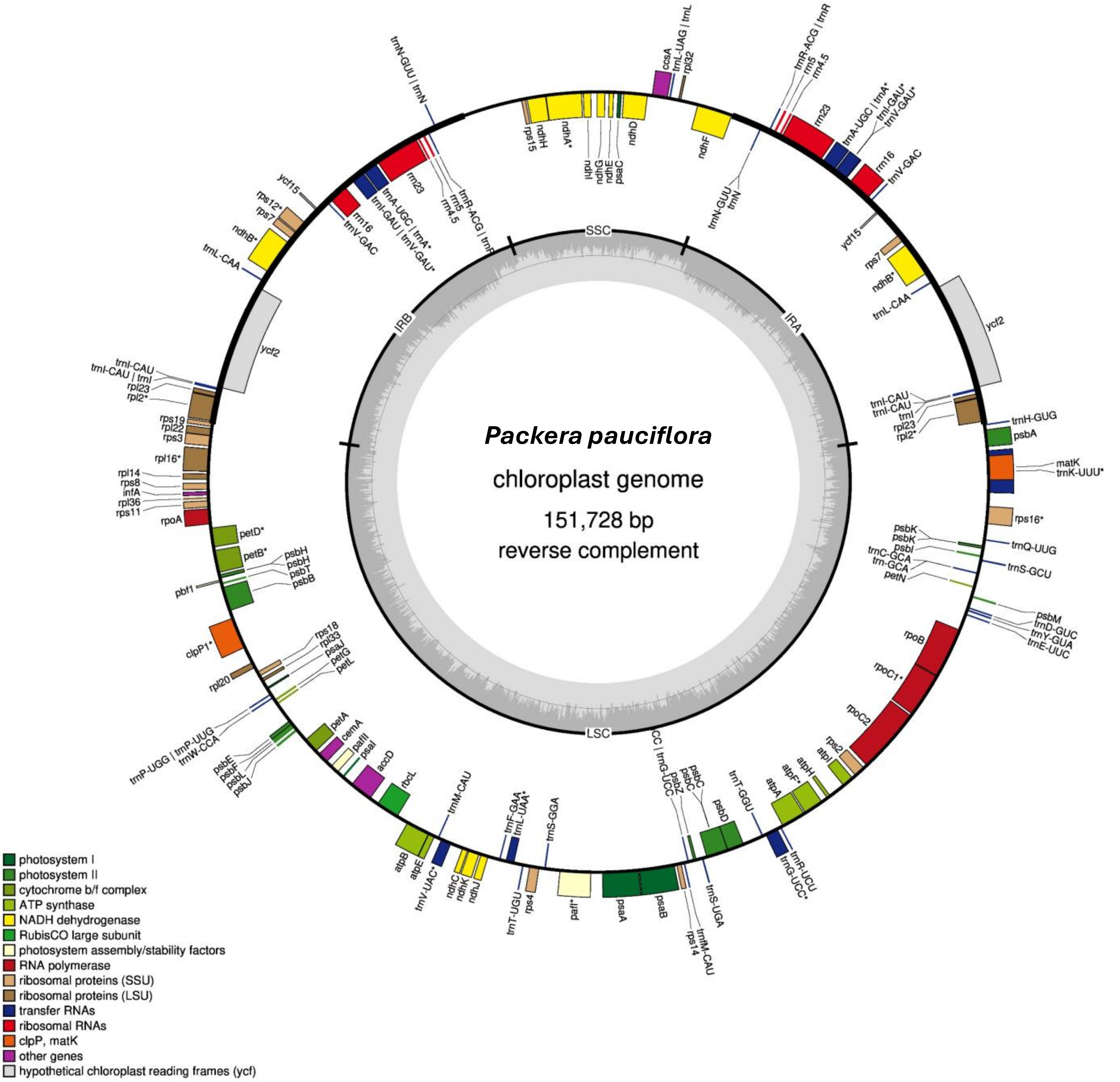

*Packera pauciflora*
chloroplast genome
151,728 bp
reverse complement
SSC
IRA
LSC
IRB
photosystem I
photosystem II
cytochrome b/f complex
ATP synthase
NADH dehydrogenase
RubisCO large subunit
photosystem assembly/stability factors
RNA polymerase
ribosomal proteins (SSU)
ribosomal proteins (LSU)
transfer RNAs
ribosomal RNAs
clpP, matK
other genes
hypothetical chloroplast reading frames (ycf)

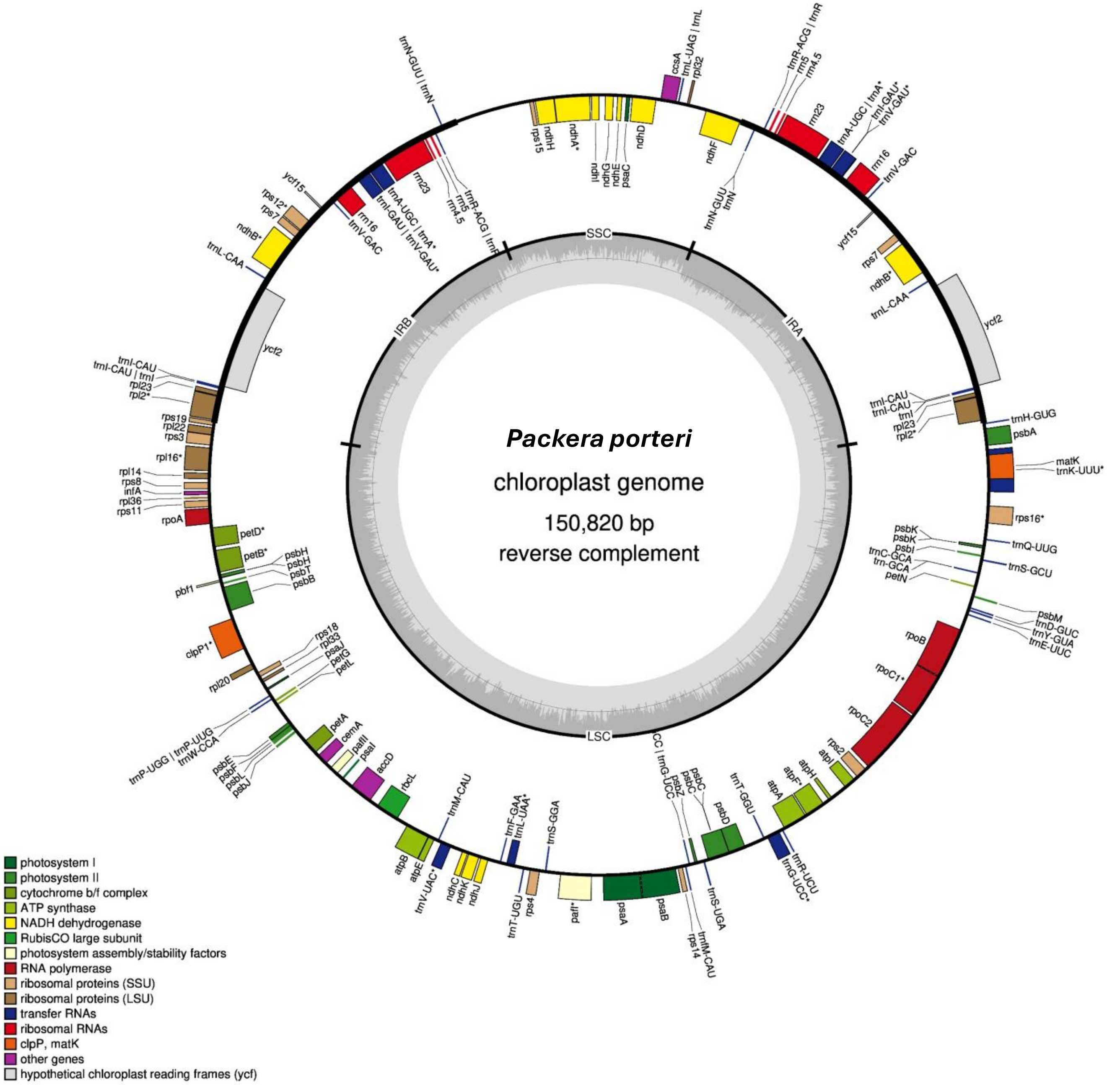

Packera porteri
chloroplast genome
150,820 bp
reverse complement
SSC
IRA
LSC
IRB
photosystem I
photosystem II
cytochrome b/f complex
ATP synthase
NADH dehydrogenase
RubisCO large subunit
photosystem assembly/stability factors
RNA polymerase
ribosomal proteins (SSU)
ribosomal proteins (LSU)
transfer RNAs
ribosomal RNAs
clpP, matK
other genes
hypothetical chloroplast reading frames (ycf)

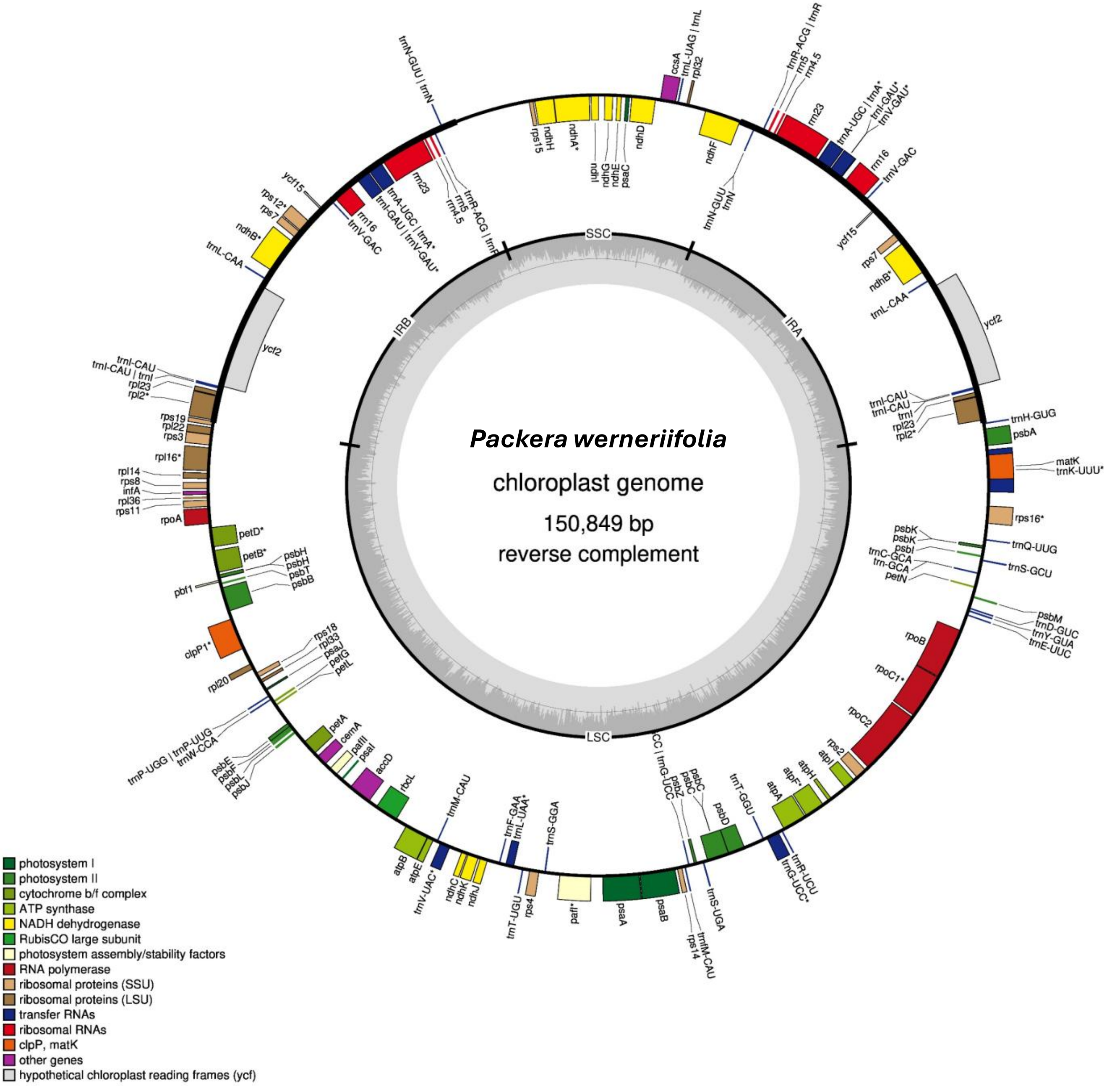

Packera werneriifolia
chloroplast genome
150,849 bp
reverse complement
SSC
IRA
IRB
LSC
photosystem I
photosystem II
cytochrome b/f complex
ATP synthase
NADH dehydrogenase
RubisCO large subunit
photosystem assembly/stability factors
RNA polymerase
ribosomal proteins (SSU)
ribosomal proteins (LSU)
transfer RNAs
ribosomal RNAs
clpP, matK
other genes
hypothetical chloroplast reading frames (ycf)

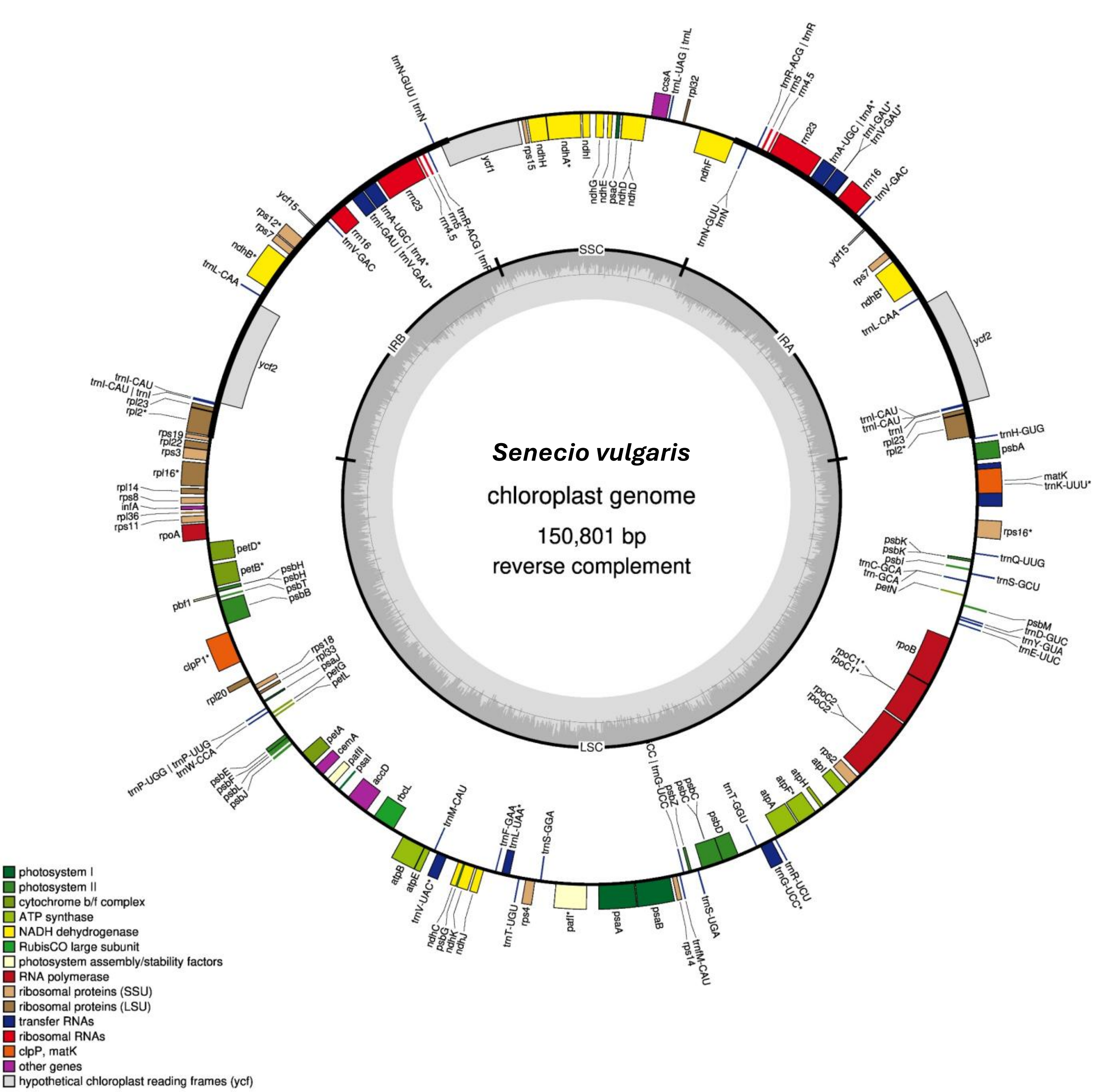
Senecio vulgaris
chloroplast genome
150,801 bp
reverse complement
SSC
IRA
IRB
LSC
ccsA
trnL-UAG | trnL
rpl32
ndhF
trnN-GUU
trnN
trnR-ACG | trnR
rrn5
rrn4.5
rrn23
trnA-UGC | trnA*
trnI-GAU*
trnV-GAU*
rrn16
trnV-GAC
ycf15
rps7
ndhB*
trnL-CAA
ycf2
trnI-CAU
trnI
rpl23
rpl2*
trnH-GUG
psbA
matK
trnK-UUU*
rps16*
psbK
psbI
trnQ-UUG
trnC-GCA
trn-GCA
trnS-GCU
petN
psbM
trnD-GUC
trnY-GUA
trnE-UUC
rpoB
rpoC1*
rpoC2
rps2
atpI
atpH
atpF*
atpA
trnT-GGU
trnR-UCU
trnG-UCC*
psbD
psbC
psbZ
trnS-UGA
trnfM-CAU
rps14
psaB
psaA
pafI*
rps4
trnT-UGU
trnS-GGA
trnF-GAA
trnL-UAA*
trnV-UAC*
ndhC
psbG
ndhK
ndhJ
trnM-CAU
atpE
atpB
rbcL
accD
psaI
pafII
cemA
petA
psbE
psbF
psbL
psbJ
trnP-UGG | trnP-UUG
trnW-CCA
rpl20
rps18
rpl33
psaJ
petG
petL
clpP1*
pbf1
psbB
psbT
psbH
petB*
petD*
rpoA
rps11
rpl36
infA
rps8
rpl14
rpl16*
rps3
rpl22
rps19
trnI-CAU | trnI
rps12*
trnR-ACG | trnR
rrn4.5
trnA-UGC | trnA*
trnI-GAU | trnV-GAU*
trnN-GUU | trnN
ycf1
rps15
ndhH
ndhA*
ndhI
ndhG
ndhE
psaC
ndhD
photosystem I
photosystem II
cytochrome b/f complex
ATP synthase
NADH dehydrogenase
RubisCO large subunit
photosystem assembly/stability factors
RNA polymerase
ribosomal proteins (SSU)
ribosomal proteins (LSU)
transfer RNAs
ribosomal RNAs
clpP, matK
other genes
hypothetical chloroplast reading frames (ycf)
