## Supplementary figures and images for "Phylogenomic challenges in polyploid-rich lineages: Insights from paralog processing and reticulation methods using the complex genus *Packera* (Asteraceae: Senecioneae)"

### Supplementary Figure 3

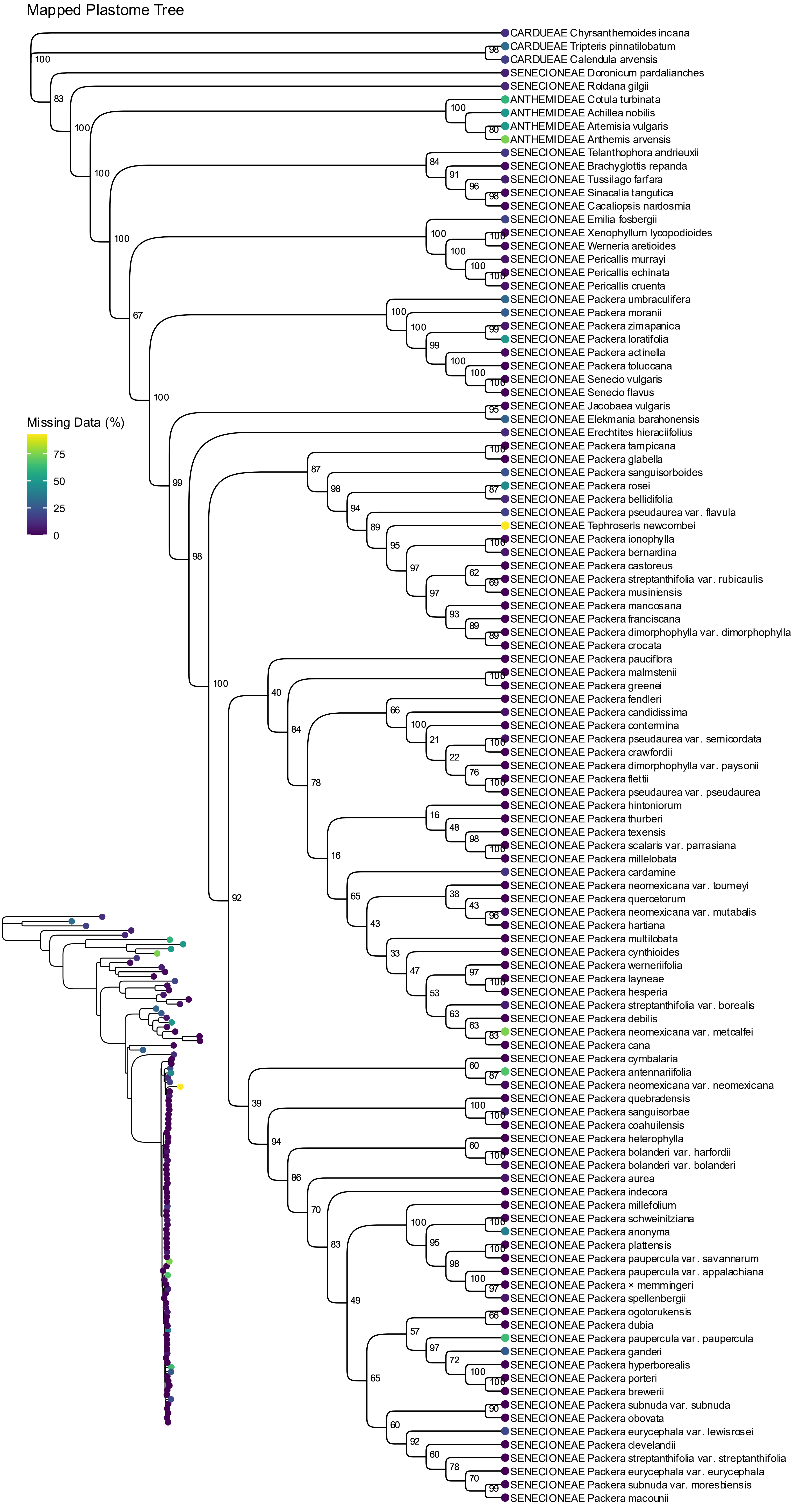

### Supplementary Figure 4

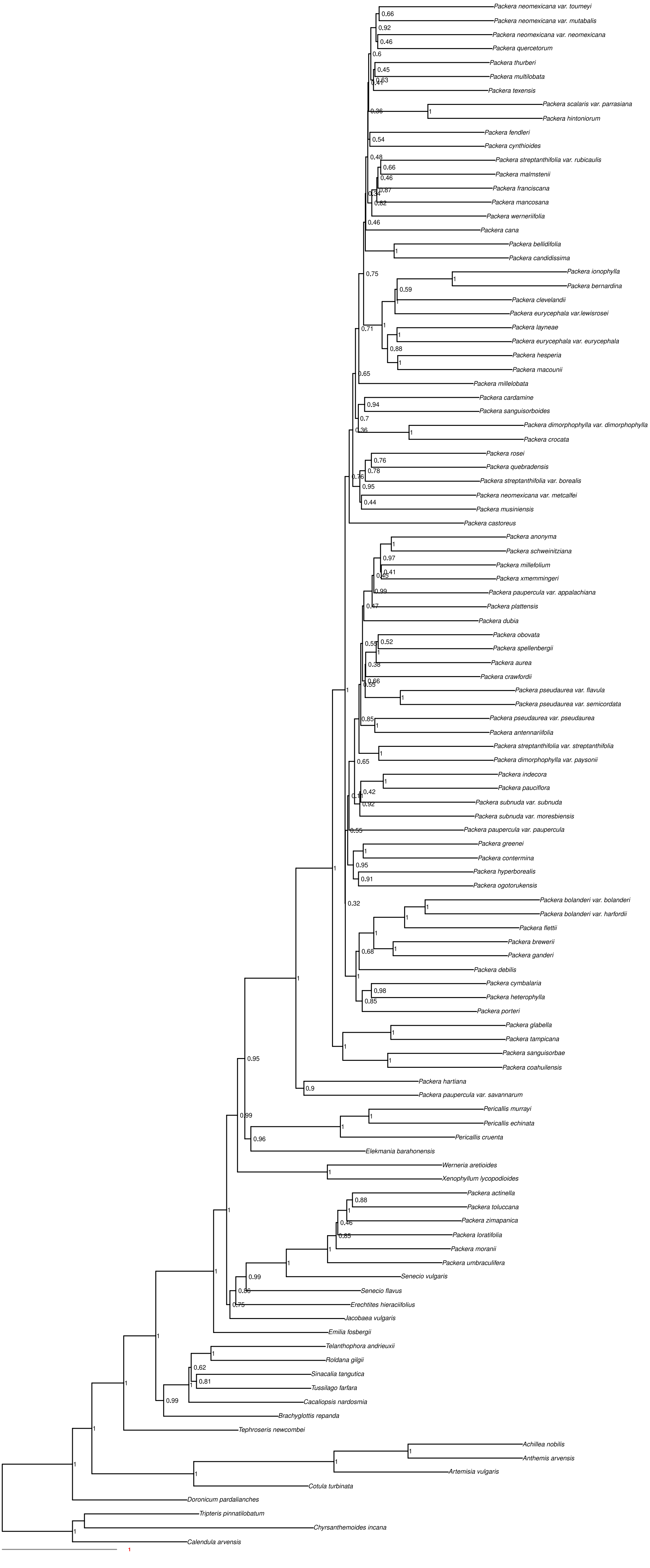

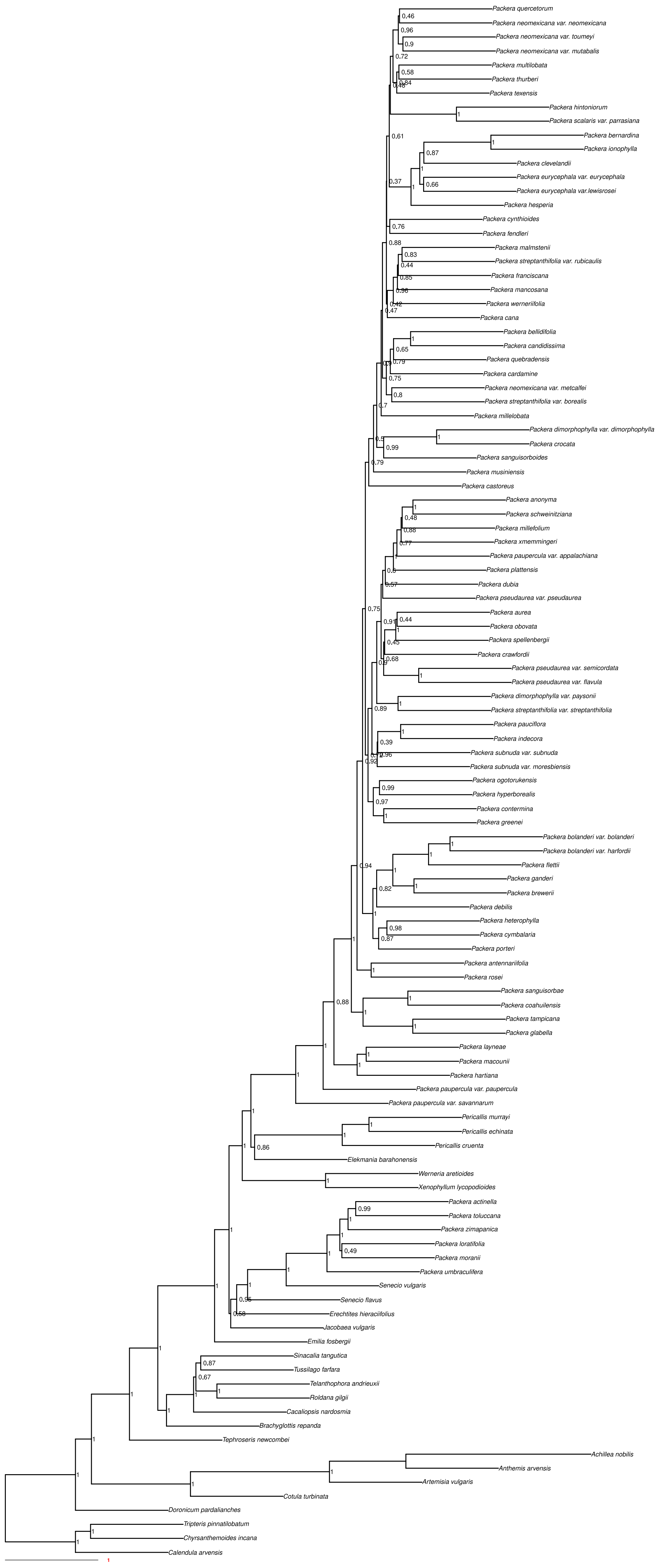

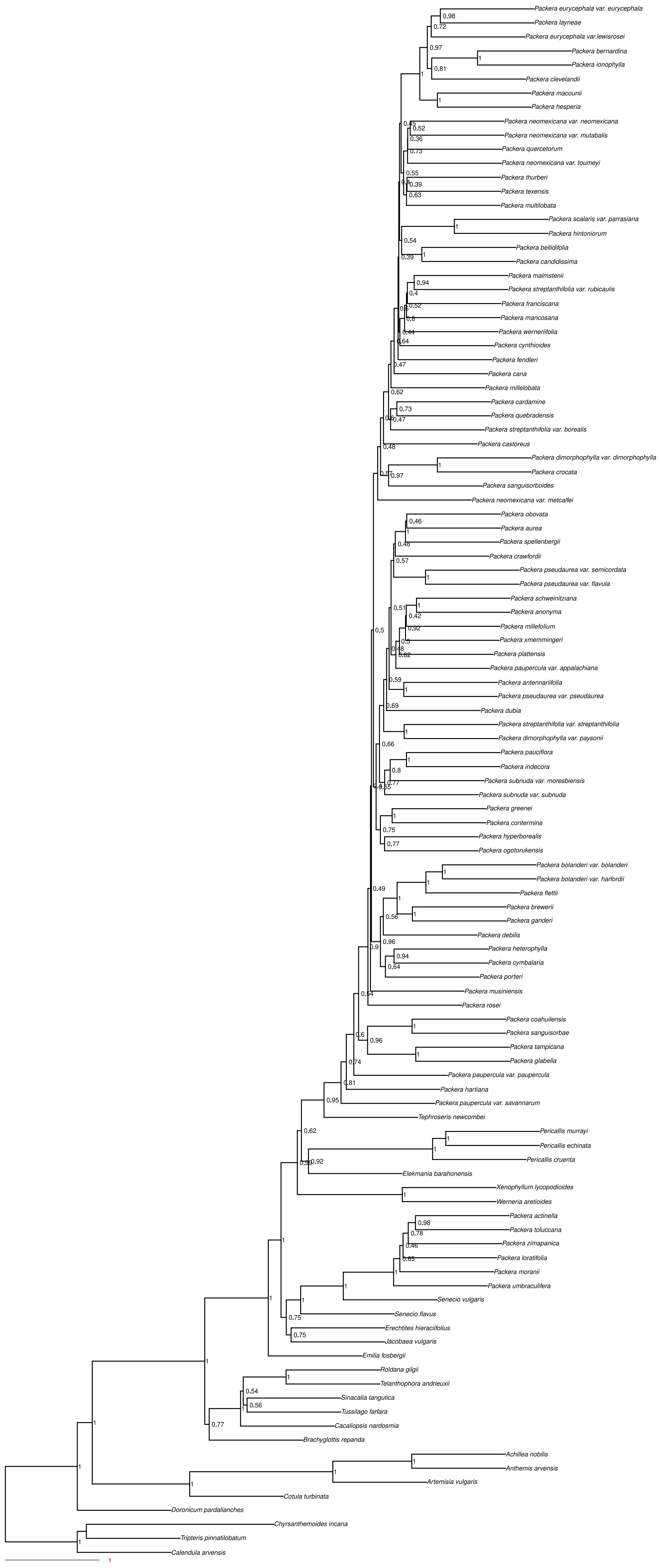

#### paragone−nf: MI

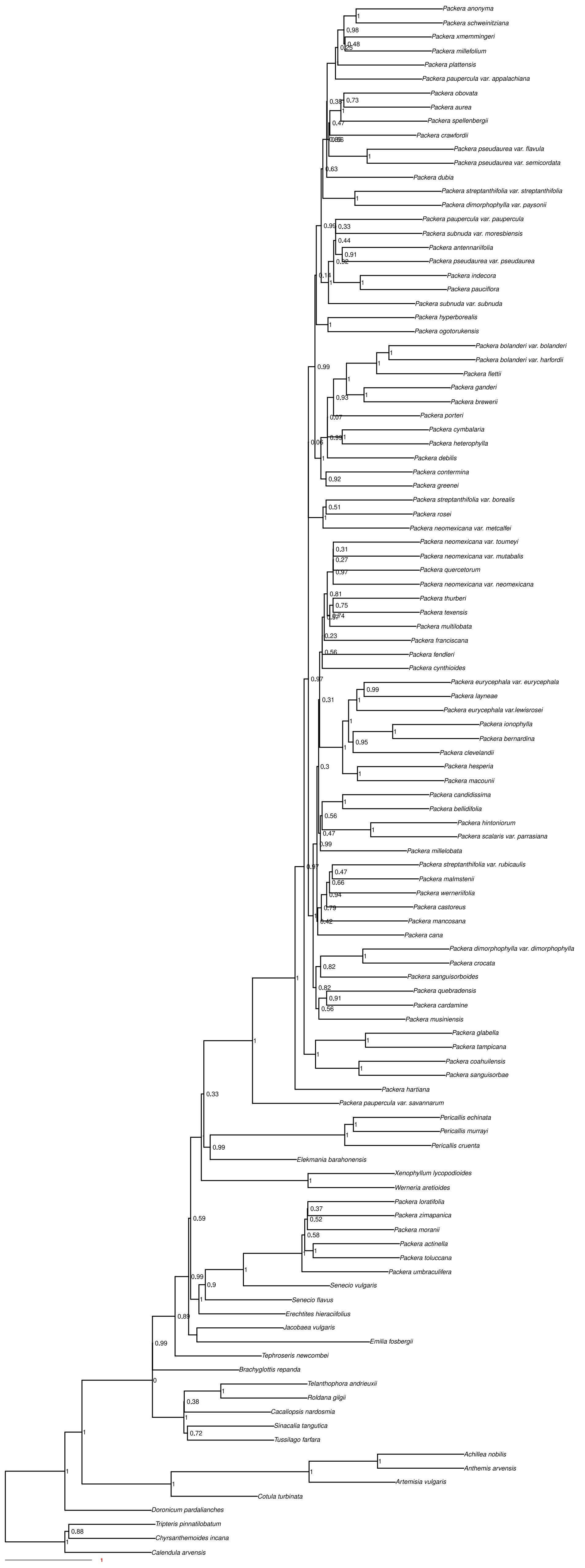

paragone−nf: MO

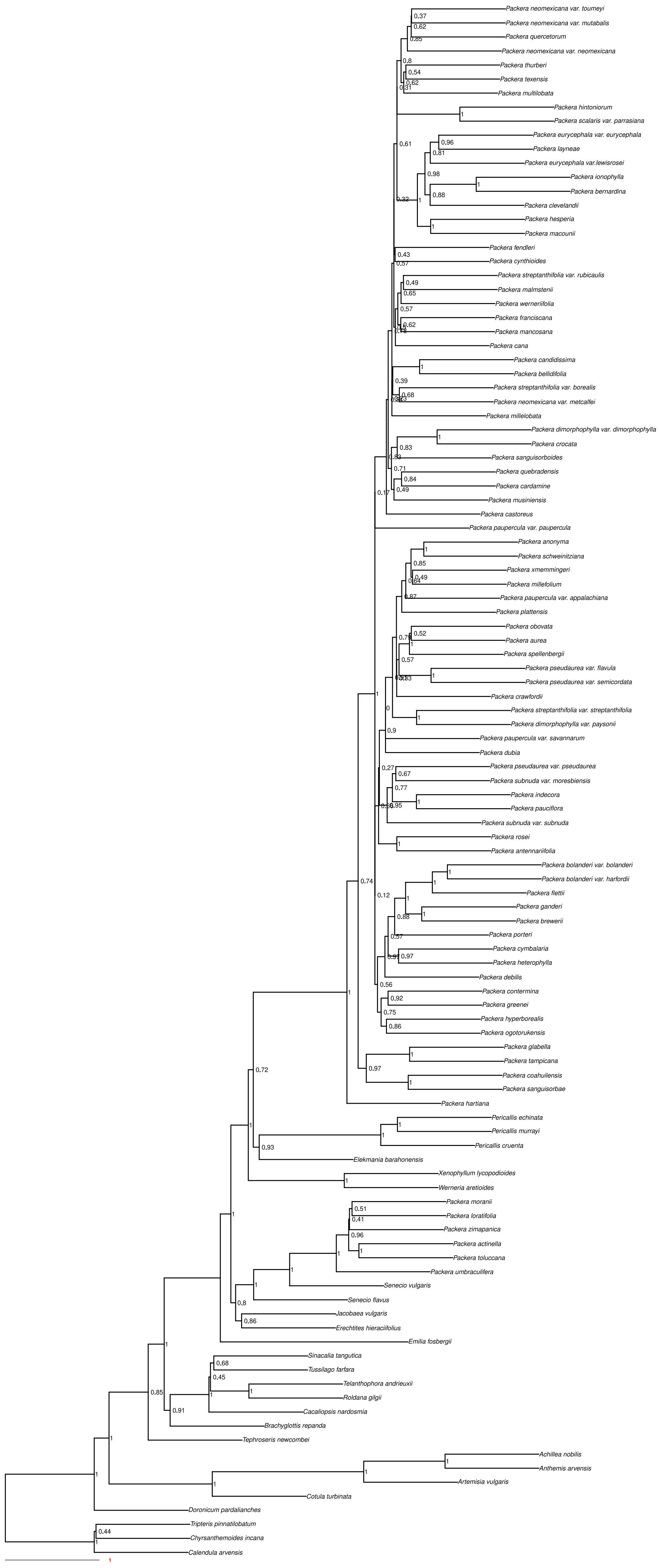

#### paragone−nf: RT

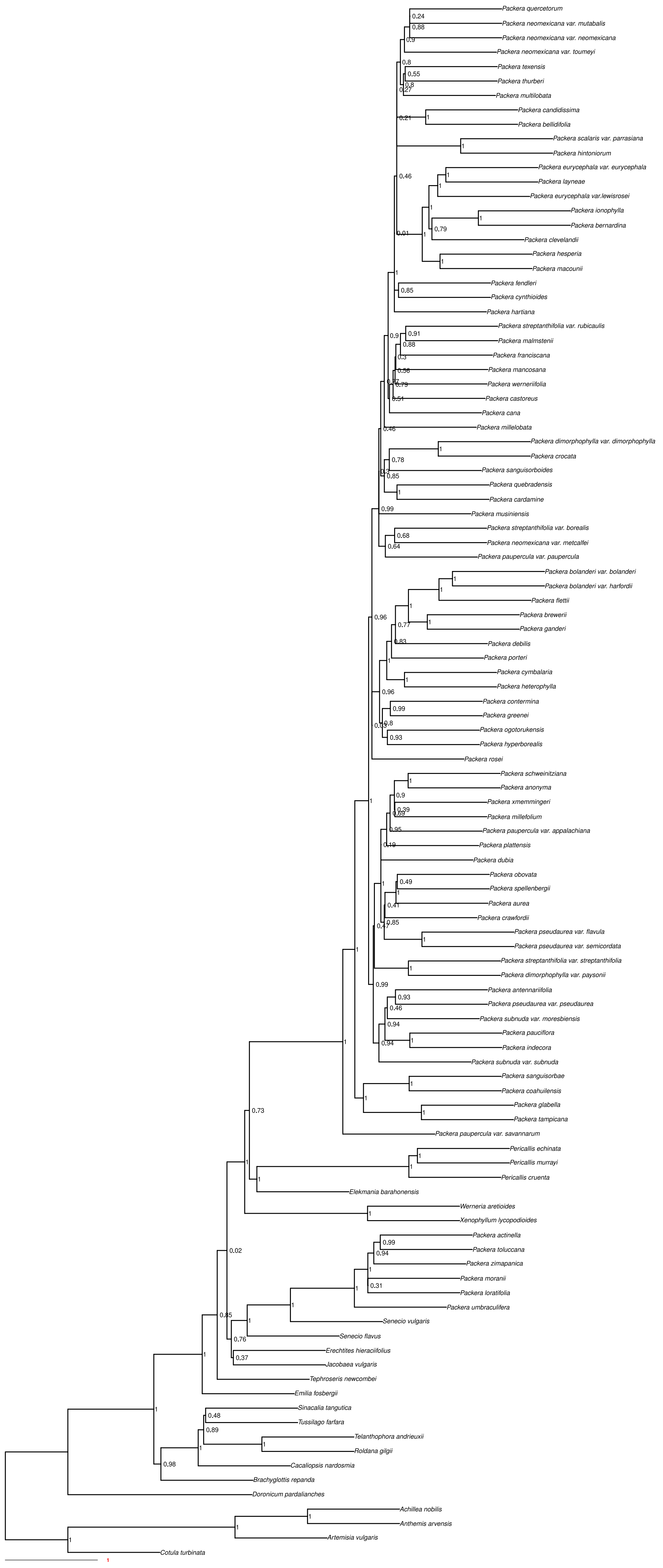

##### paragone−nf: 1to1

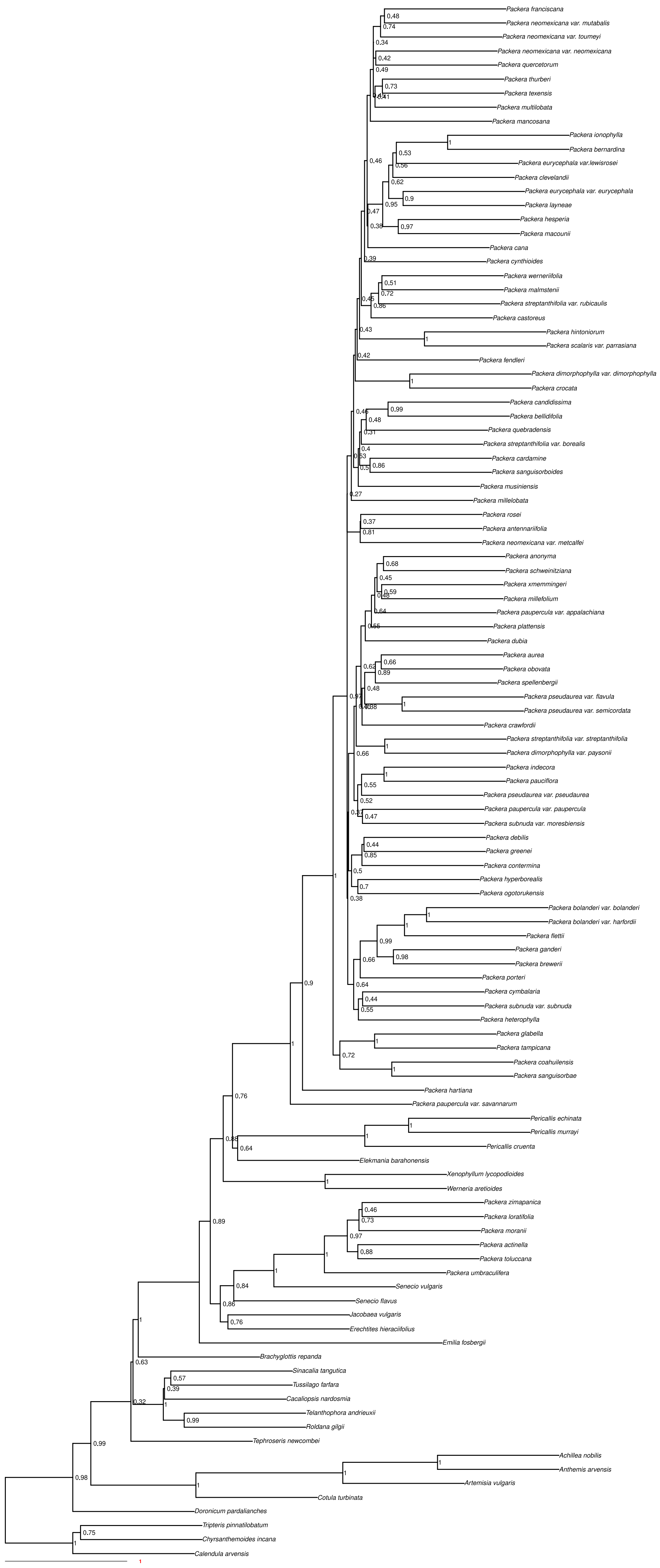

#### Concatenated

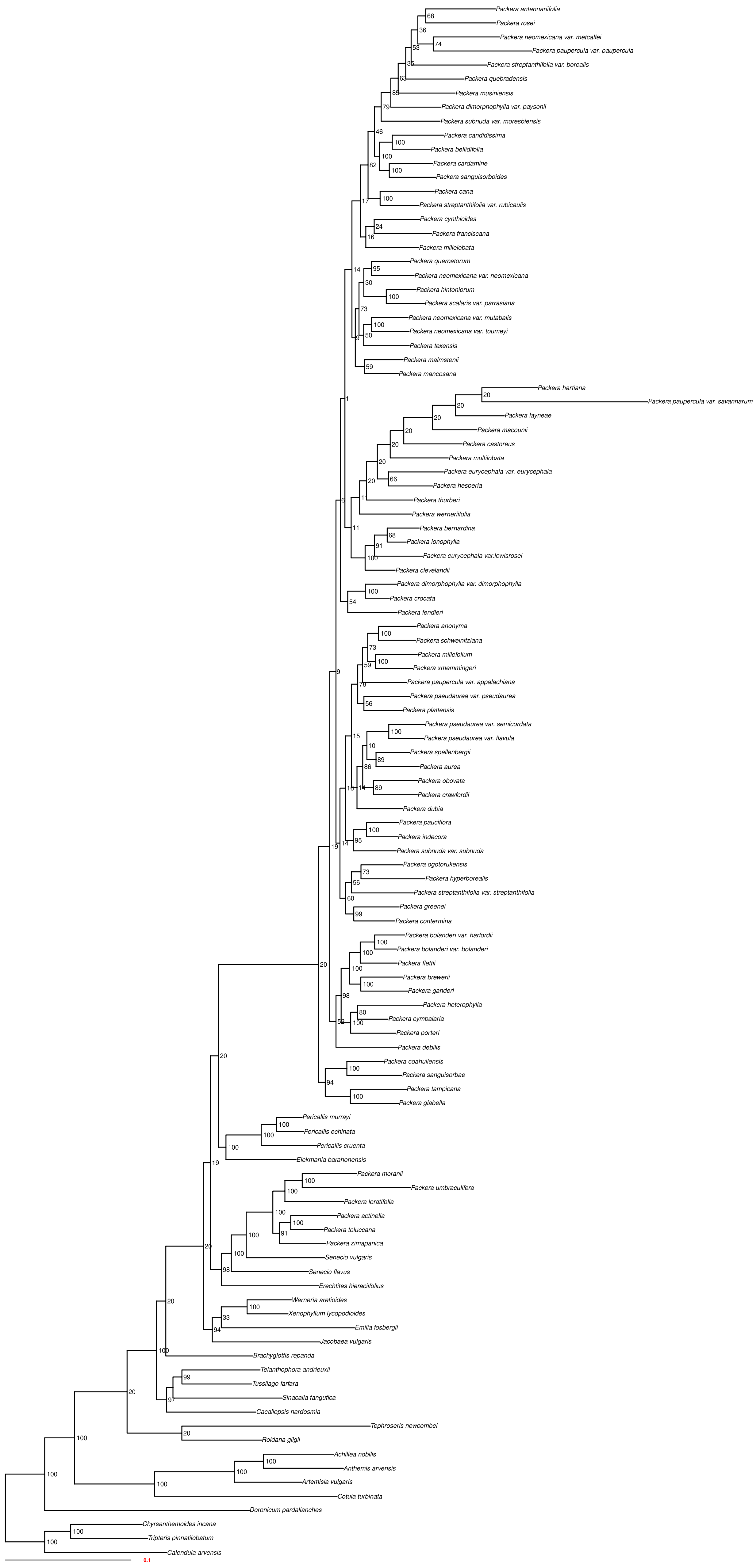

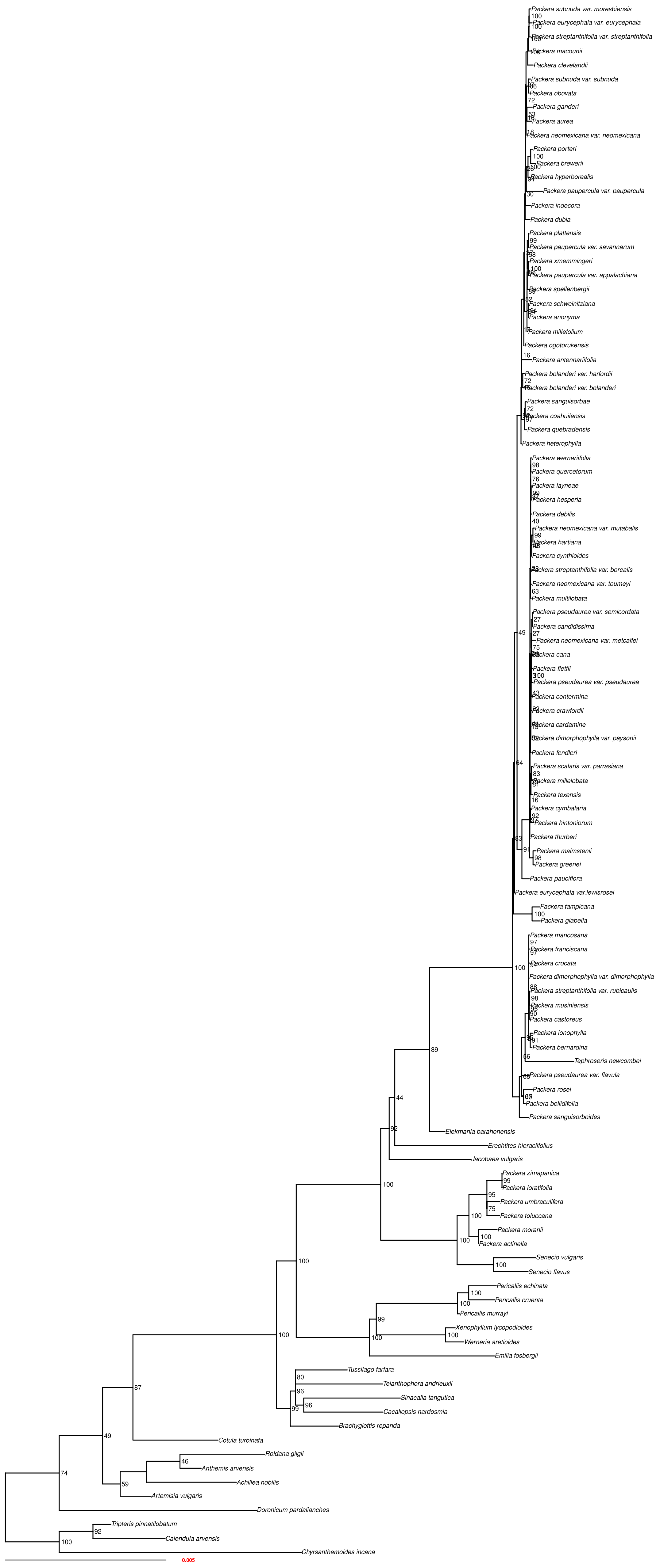

### Supplementary Figure 5

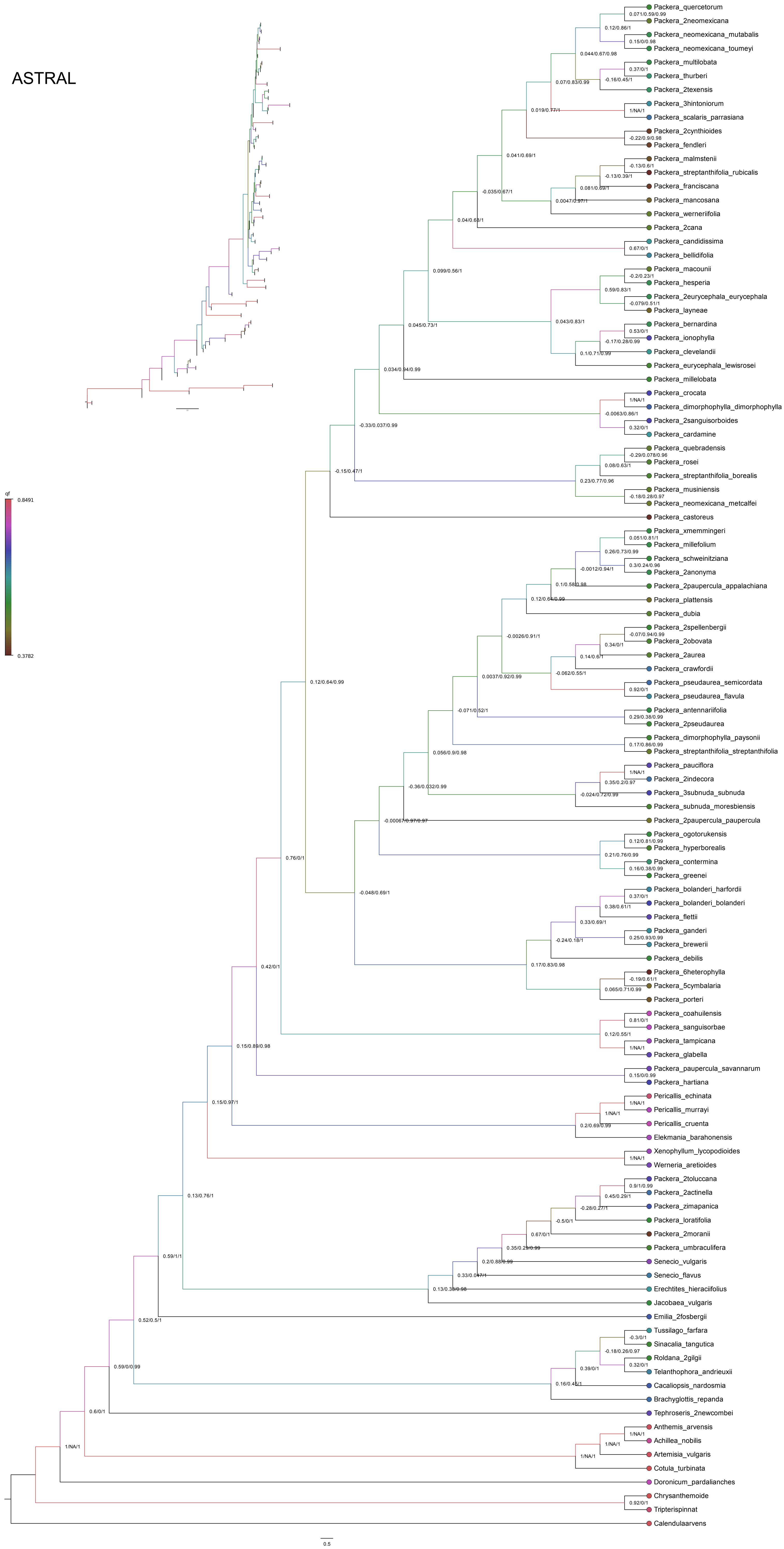

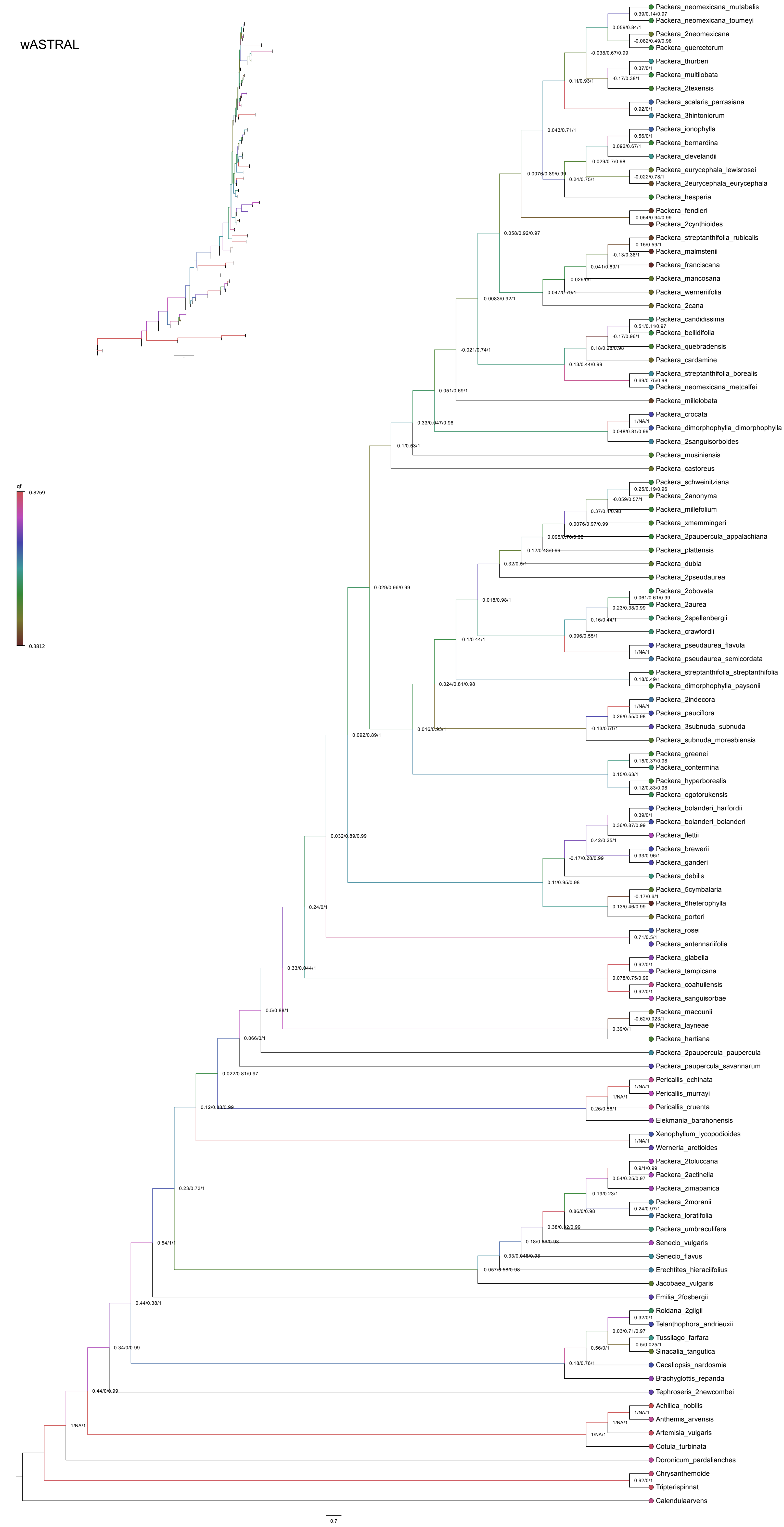

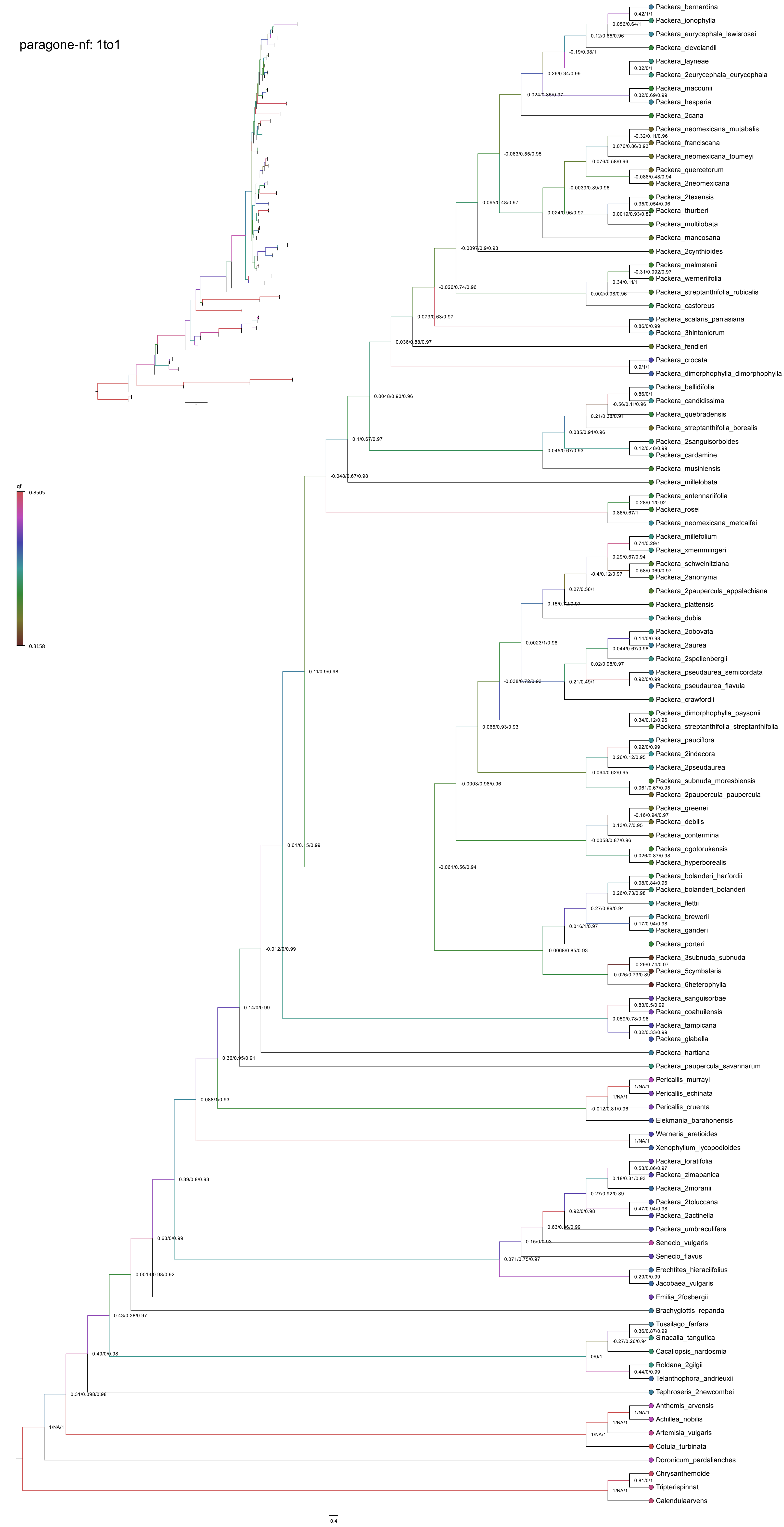

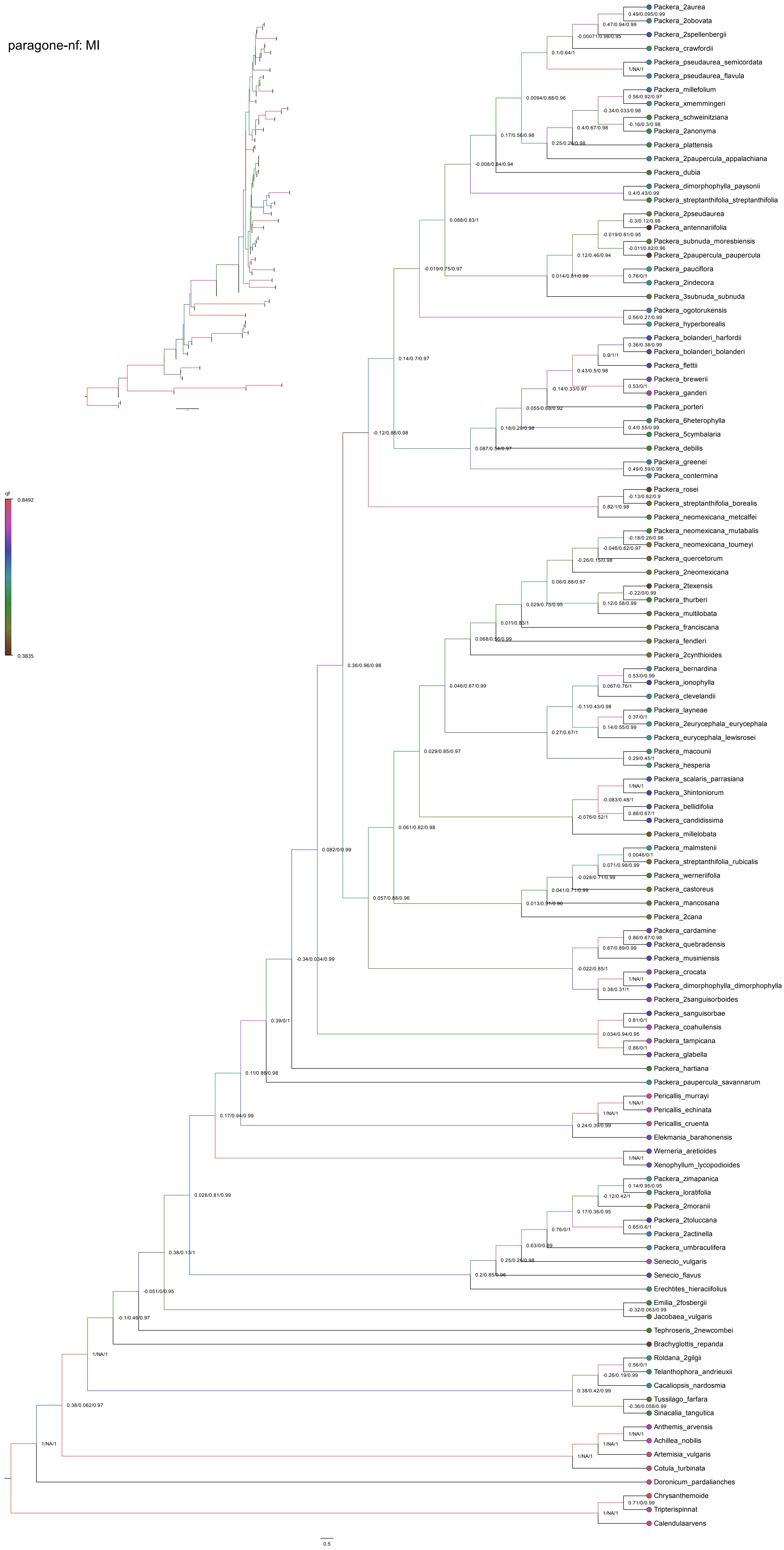

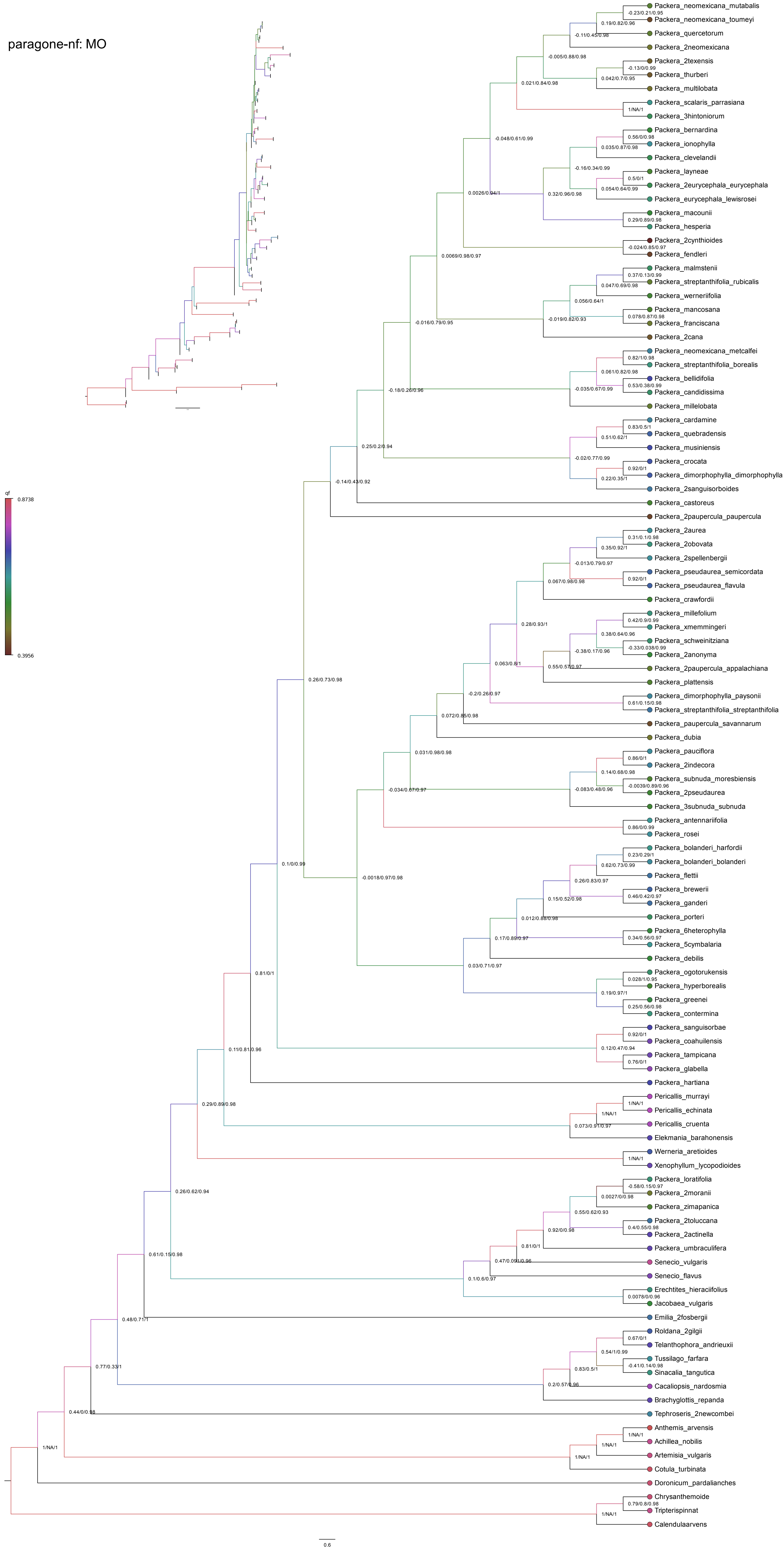

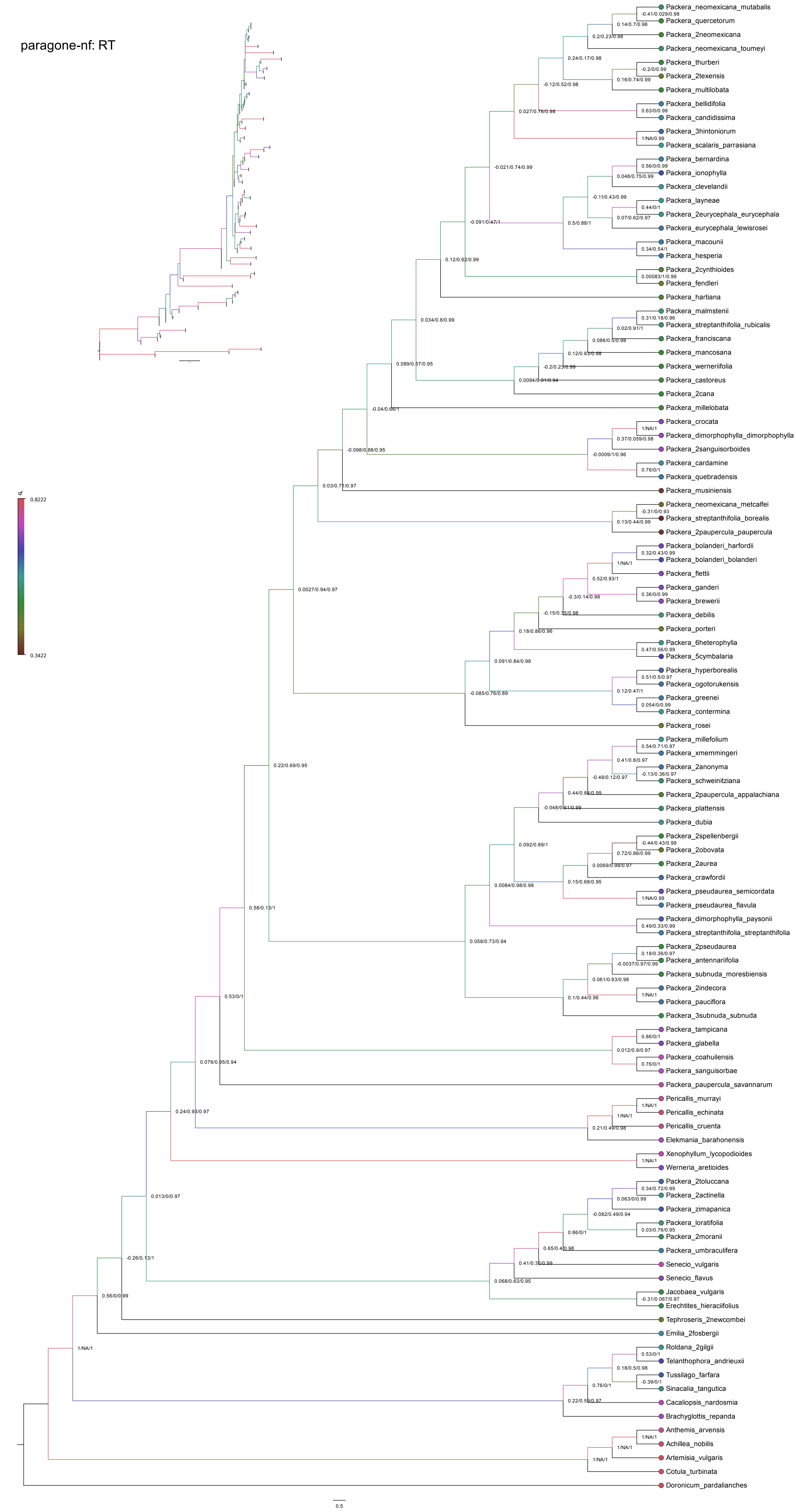

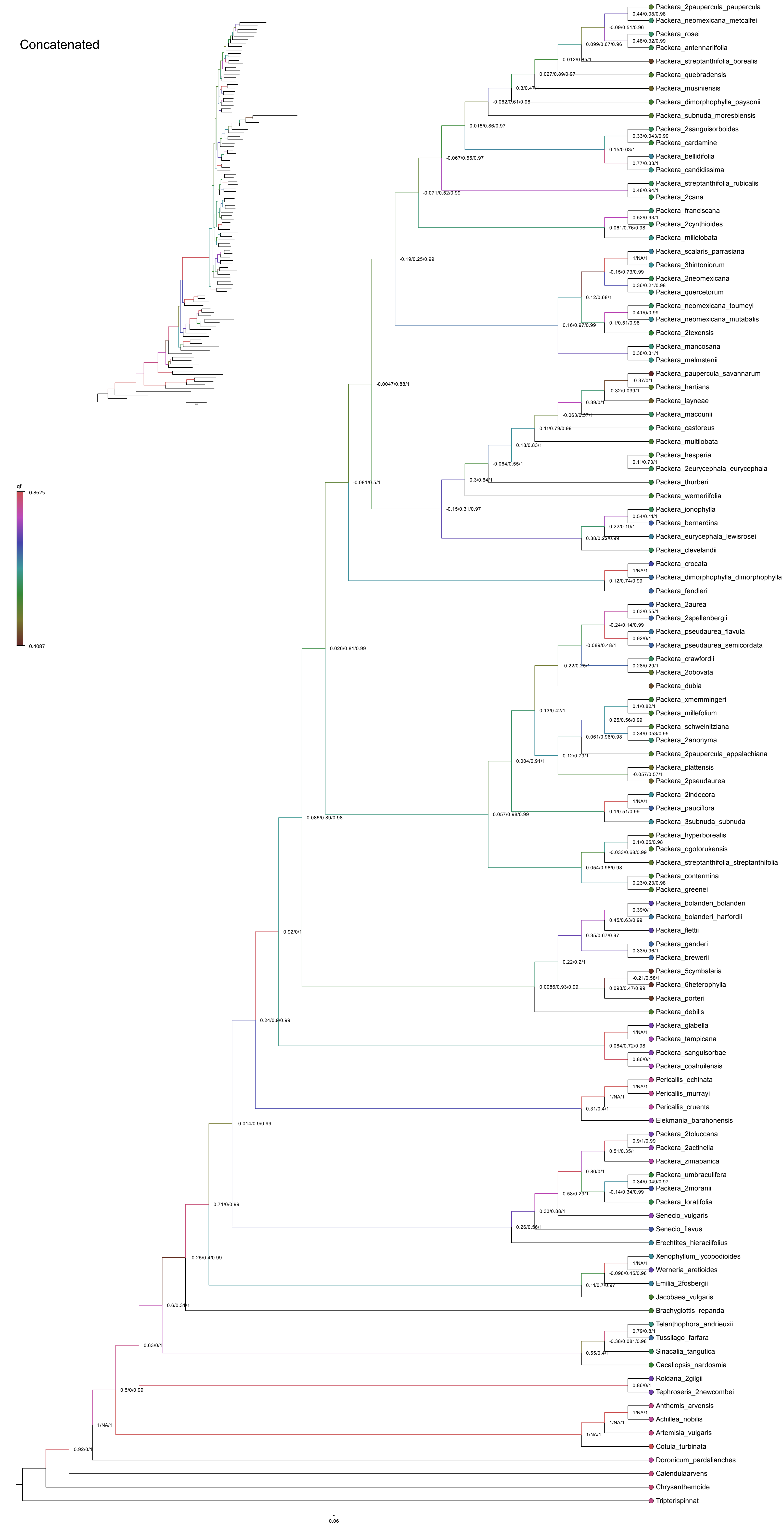

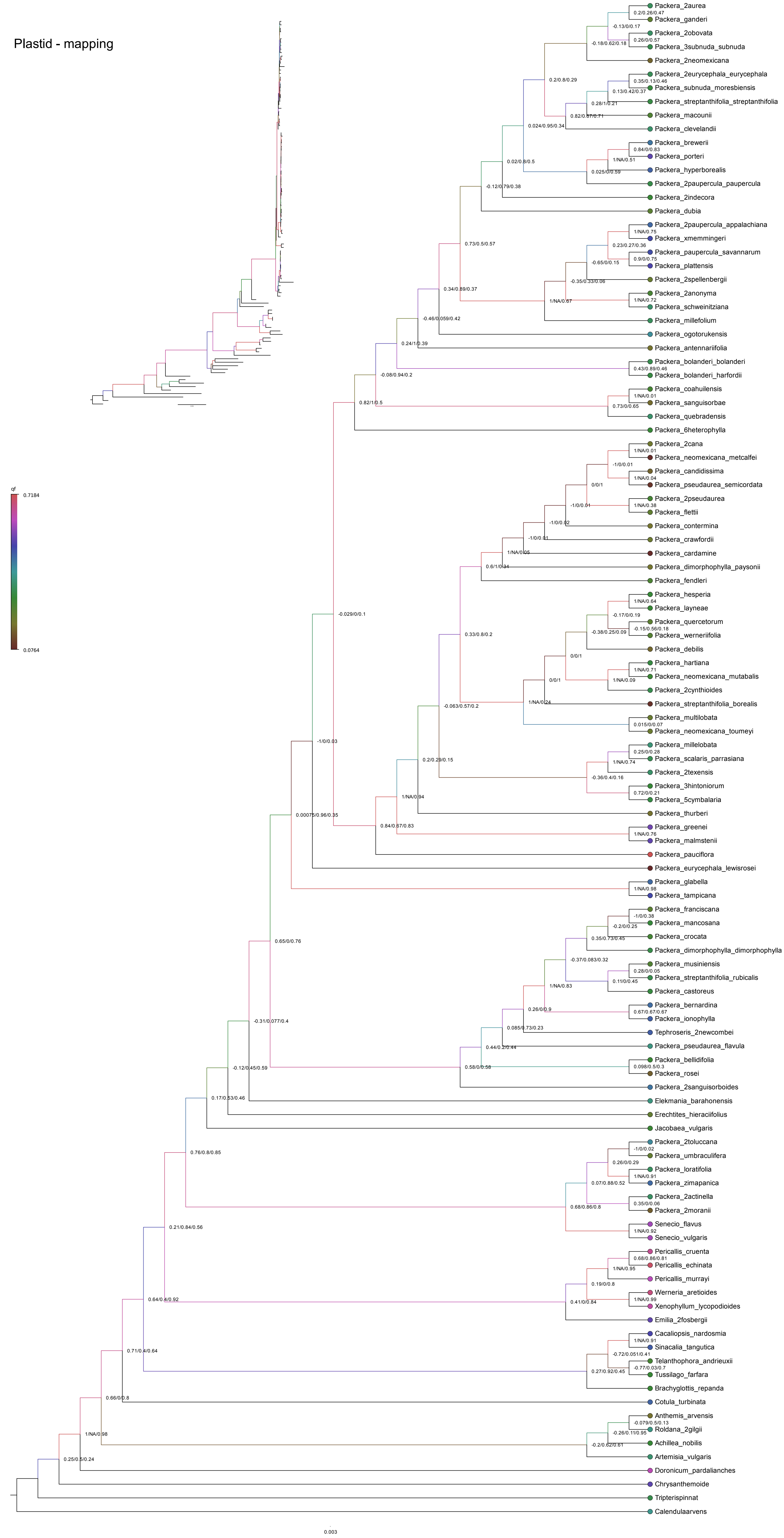

### Supplementary Figure 6

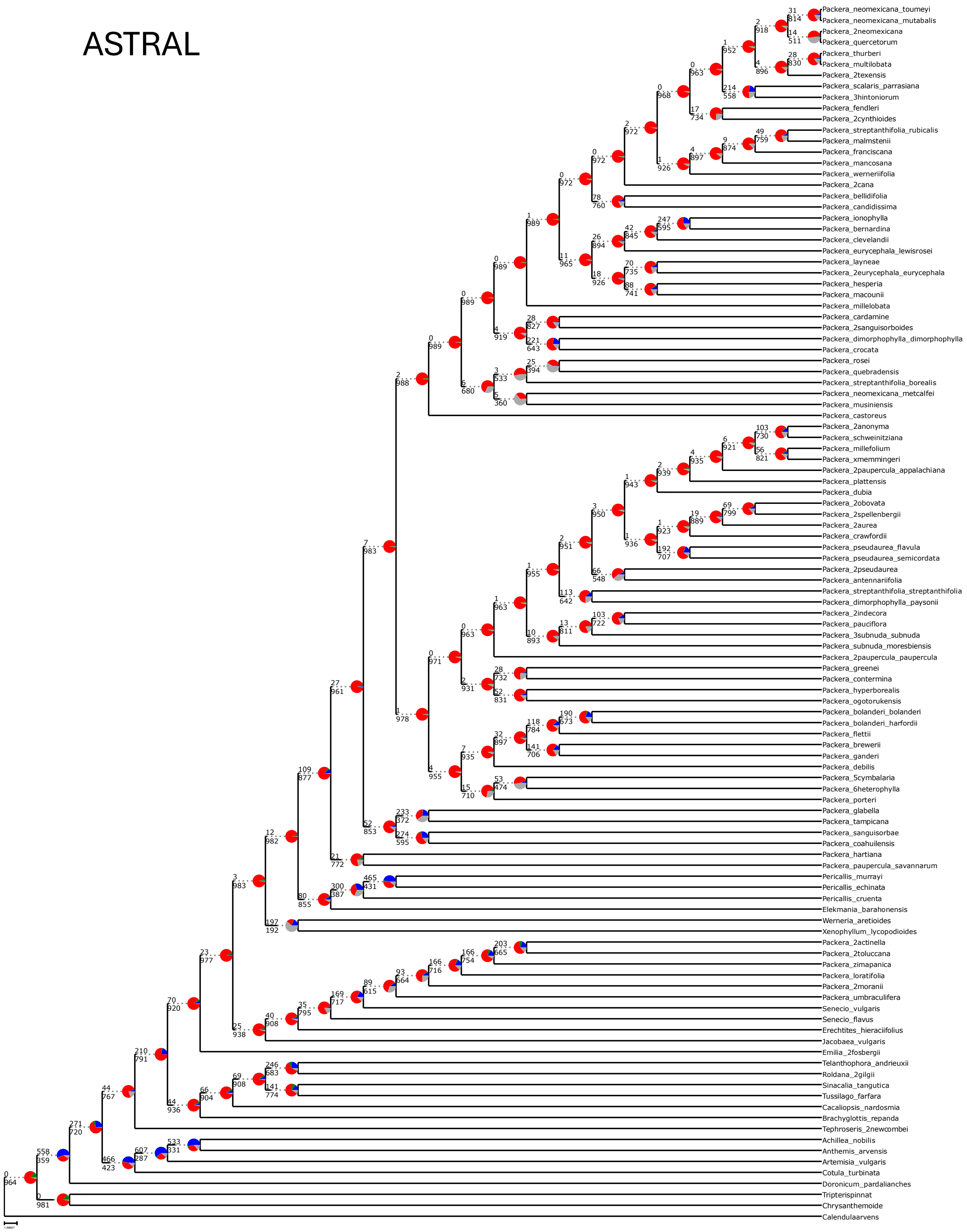

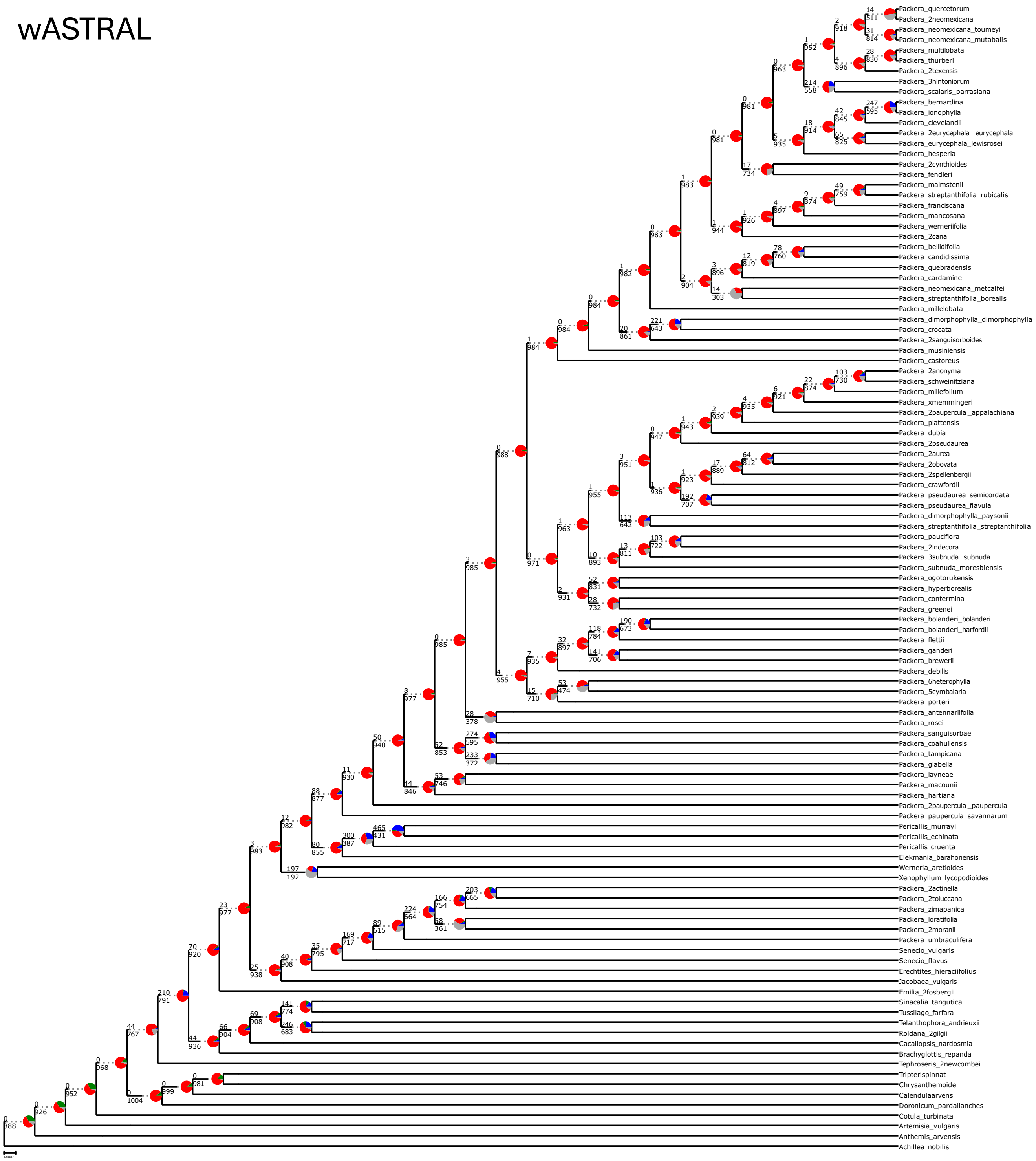

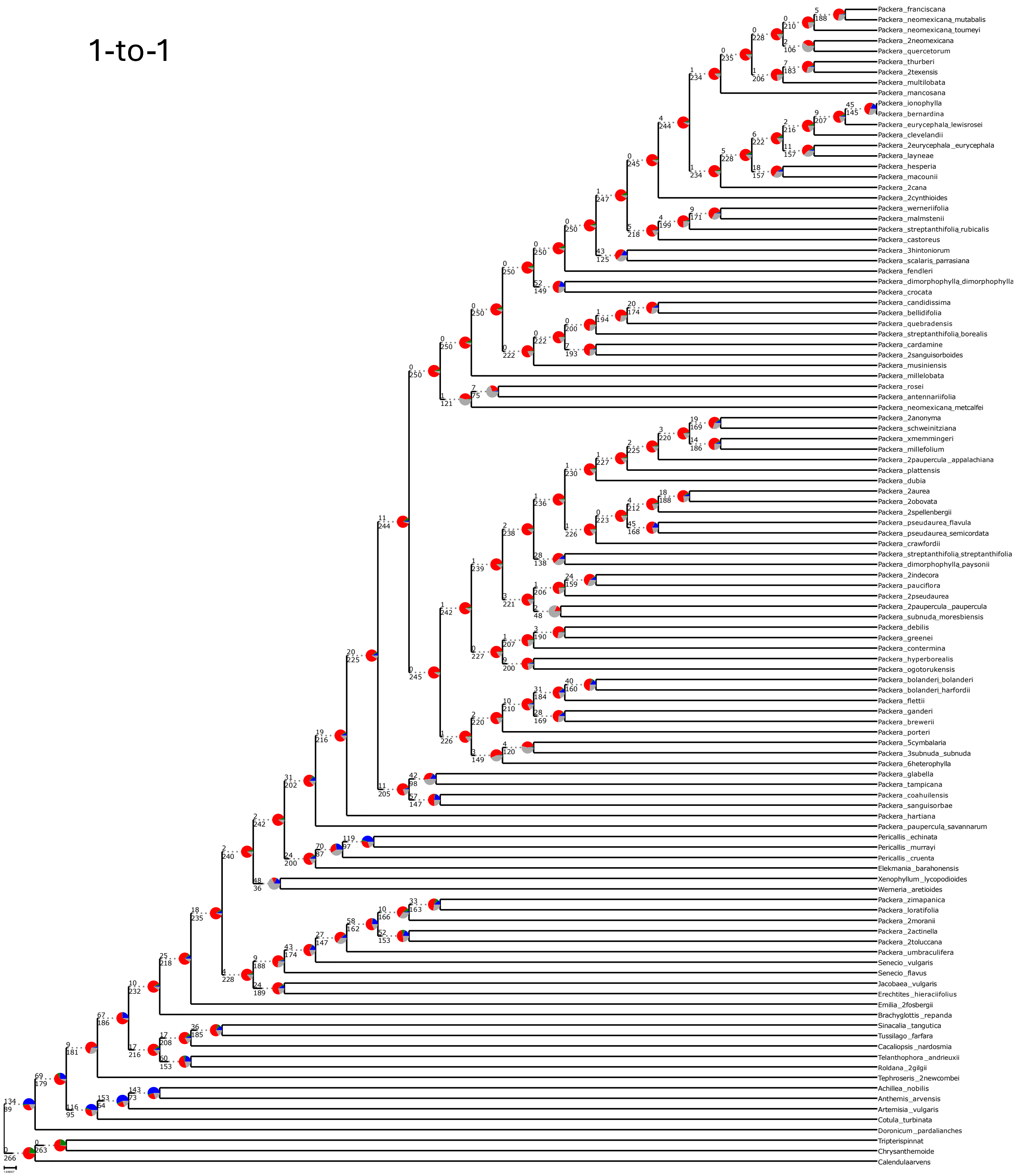

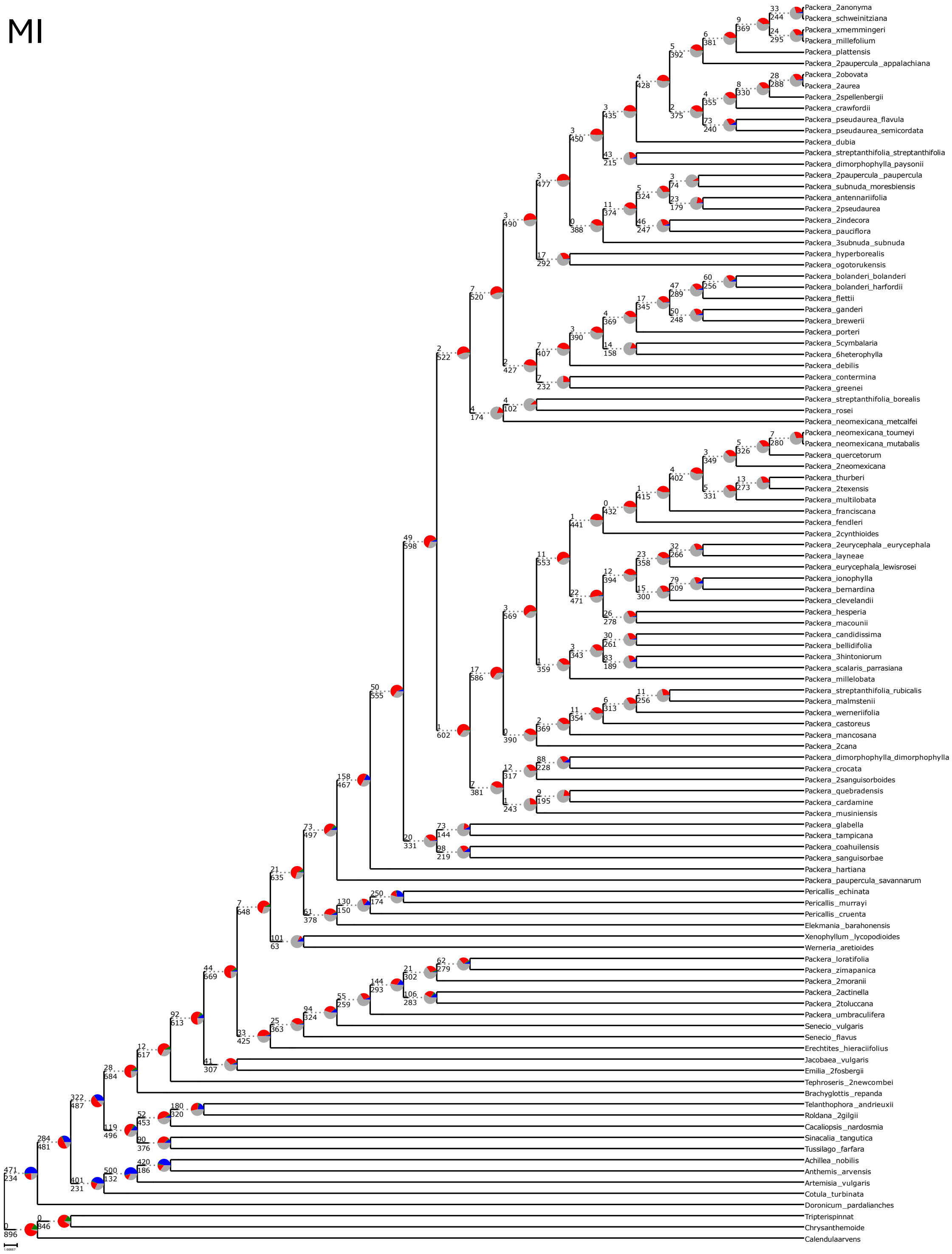

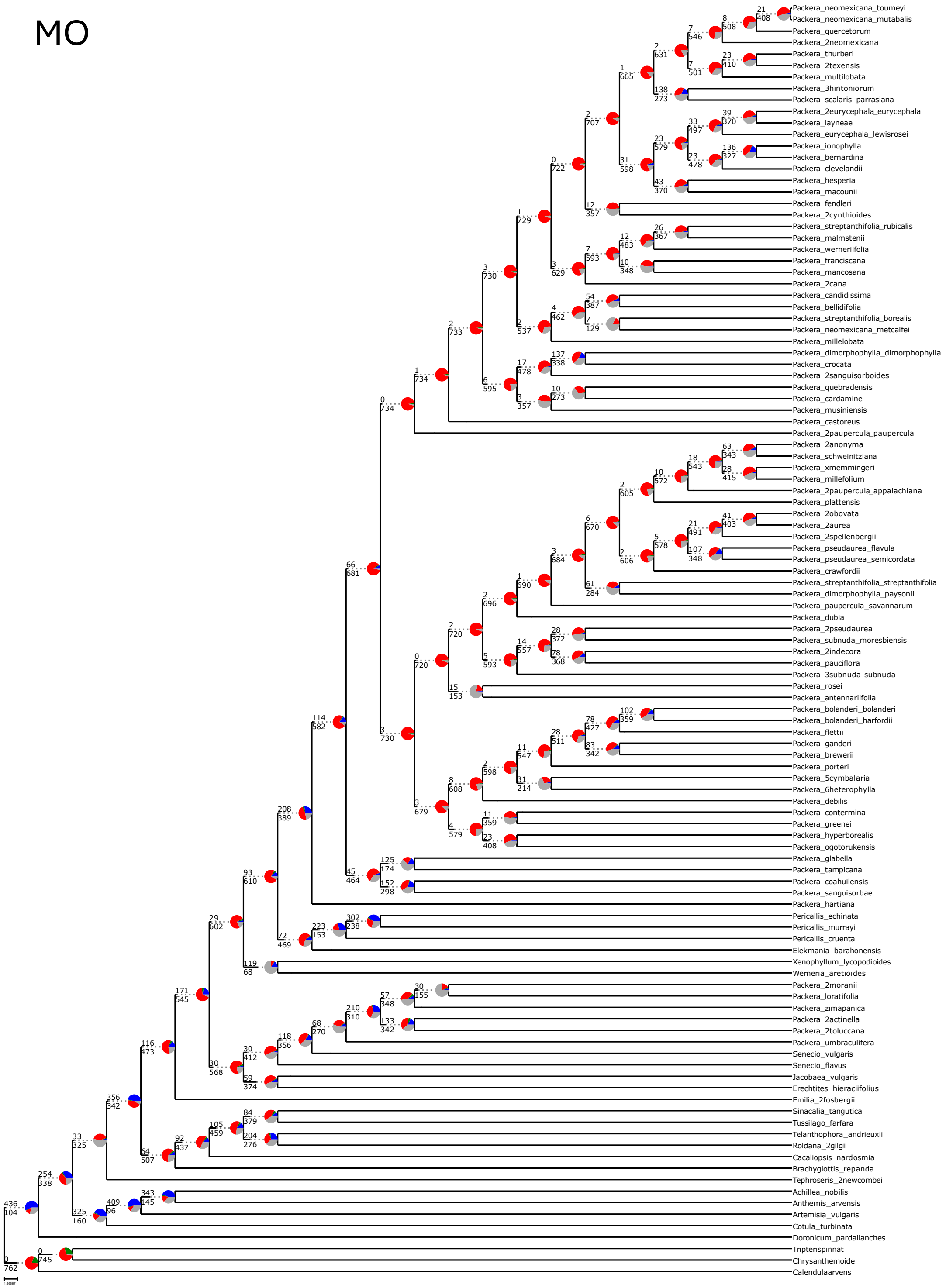

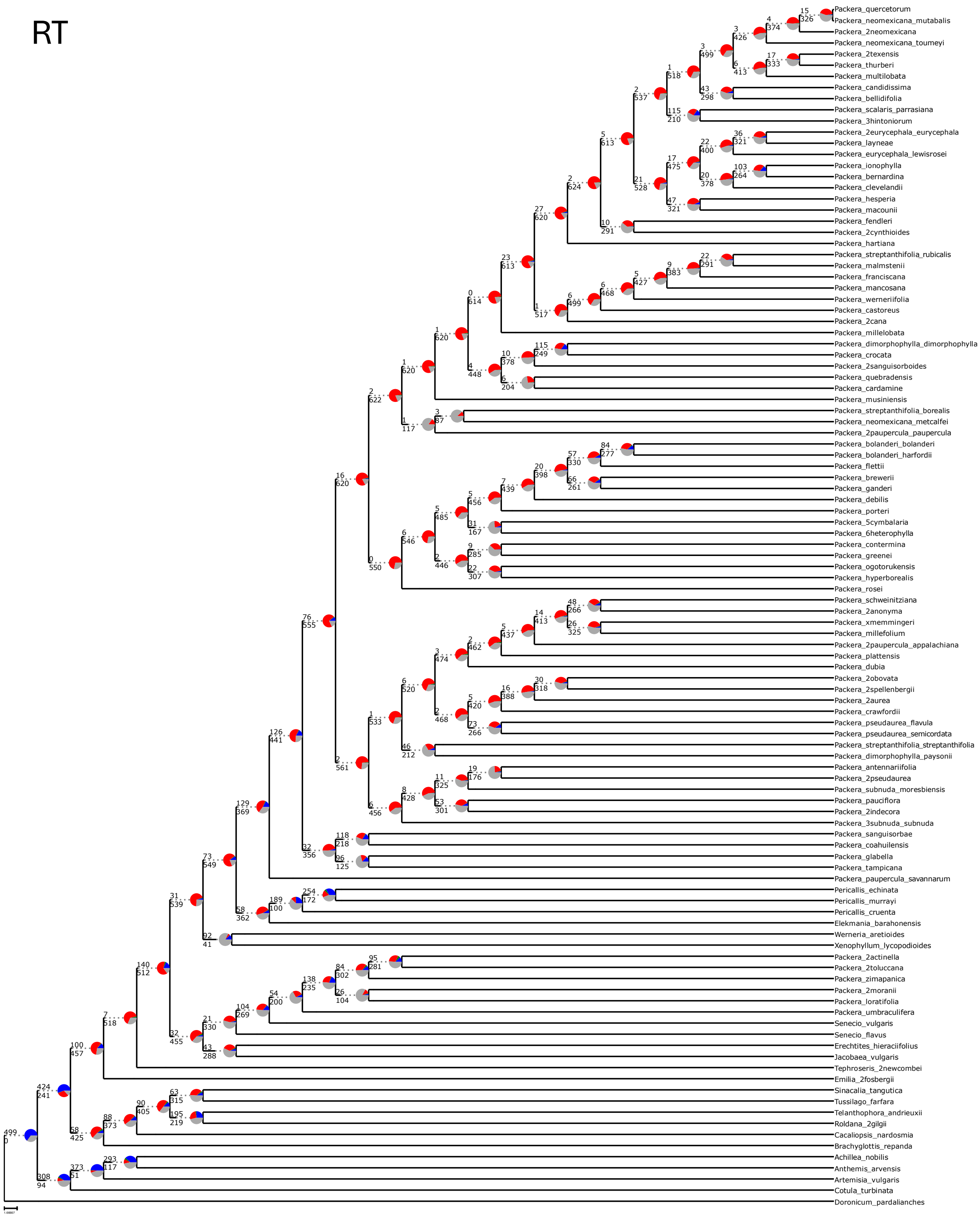

### Supplementary Figure 7

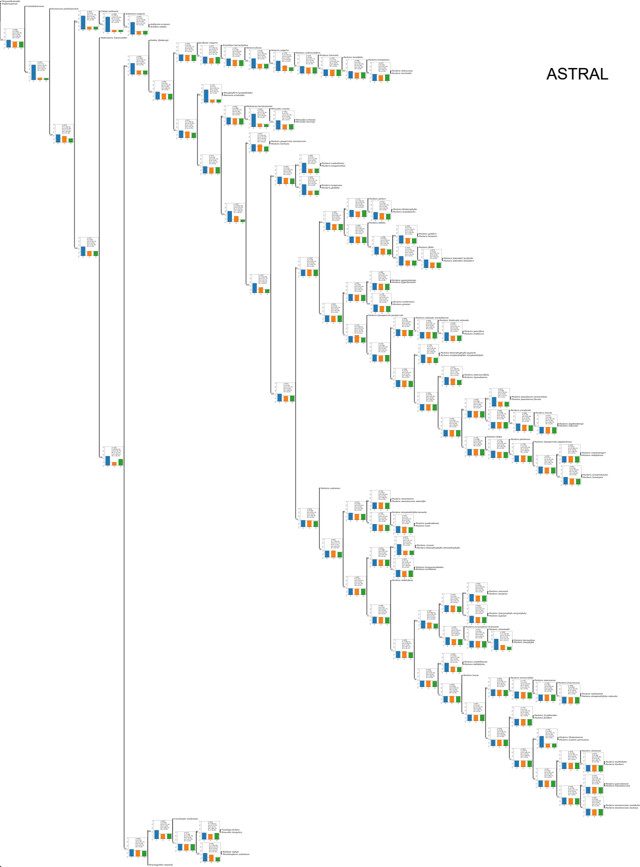

#### wASTRAL

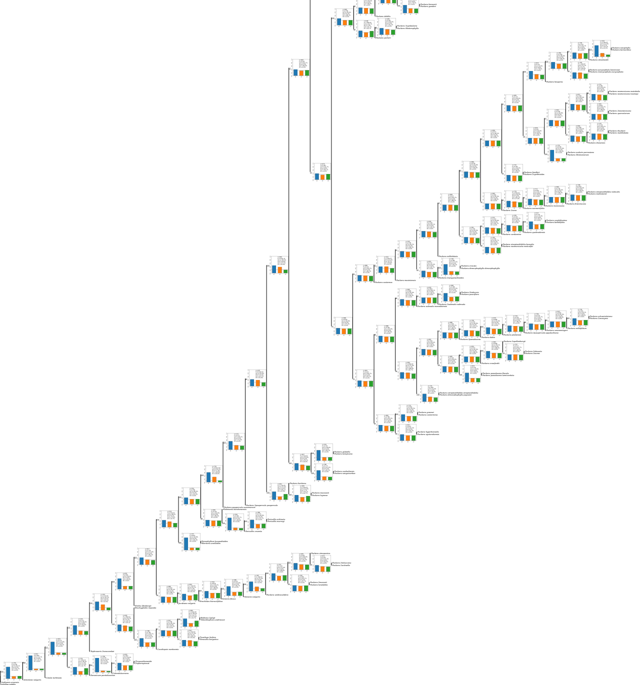

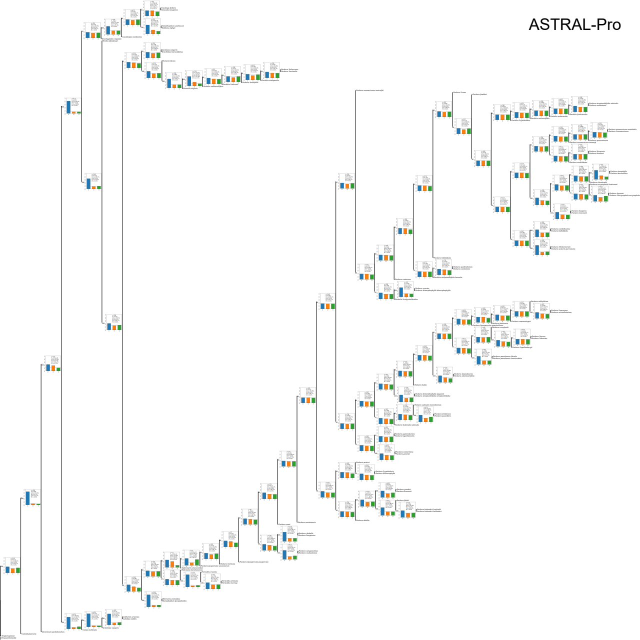

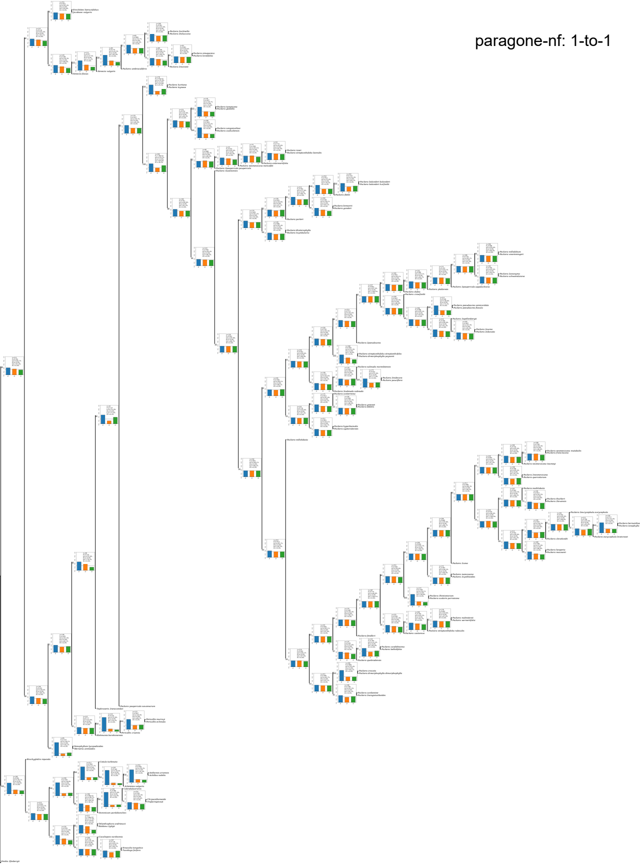

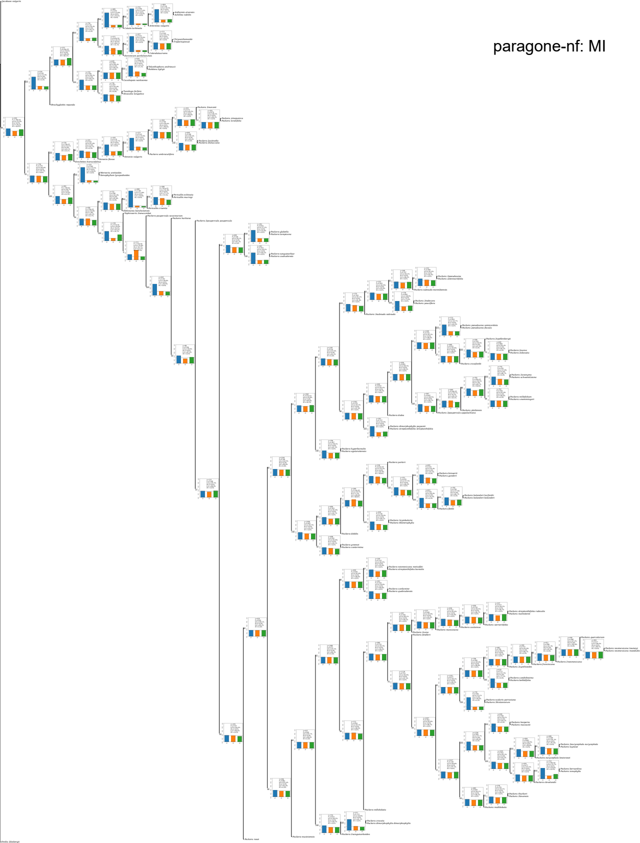

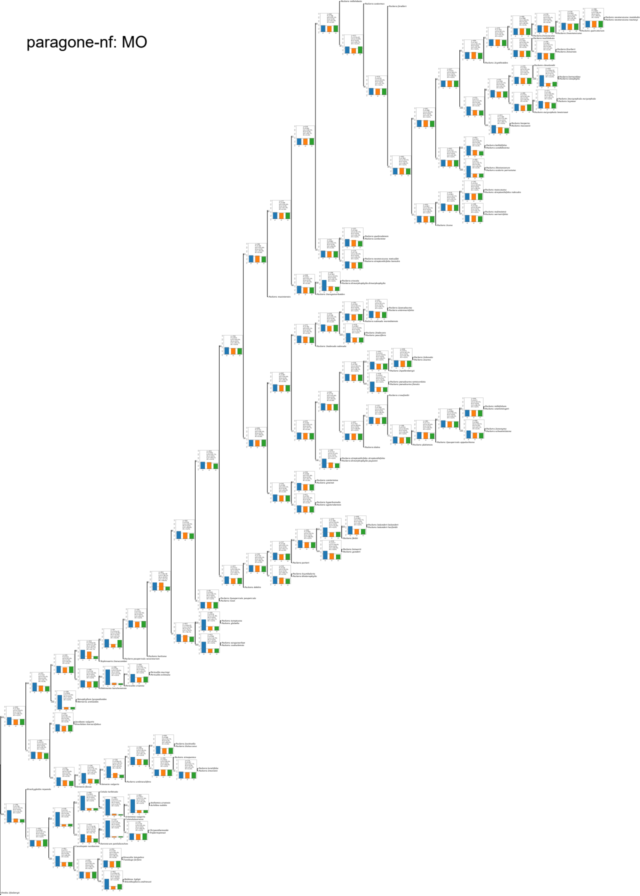

paragone-nf: RT

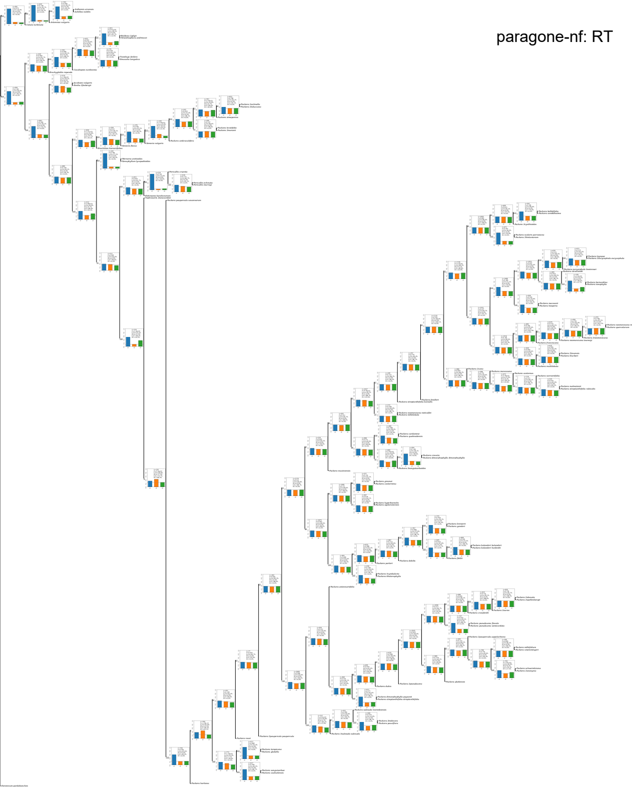

### Supplementary Figure 10

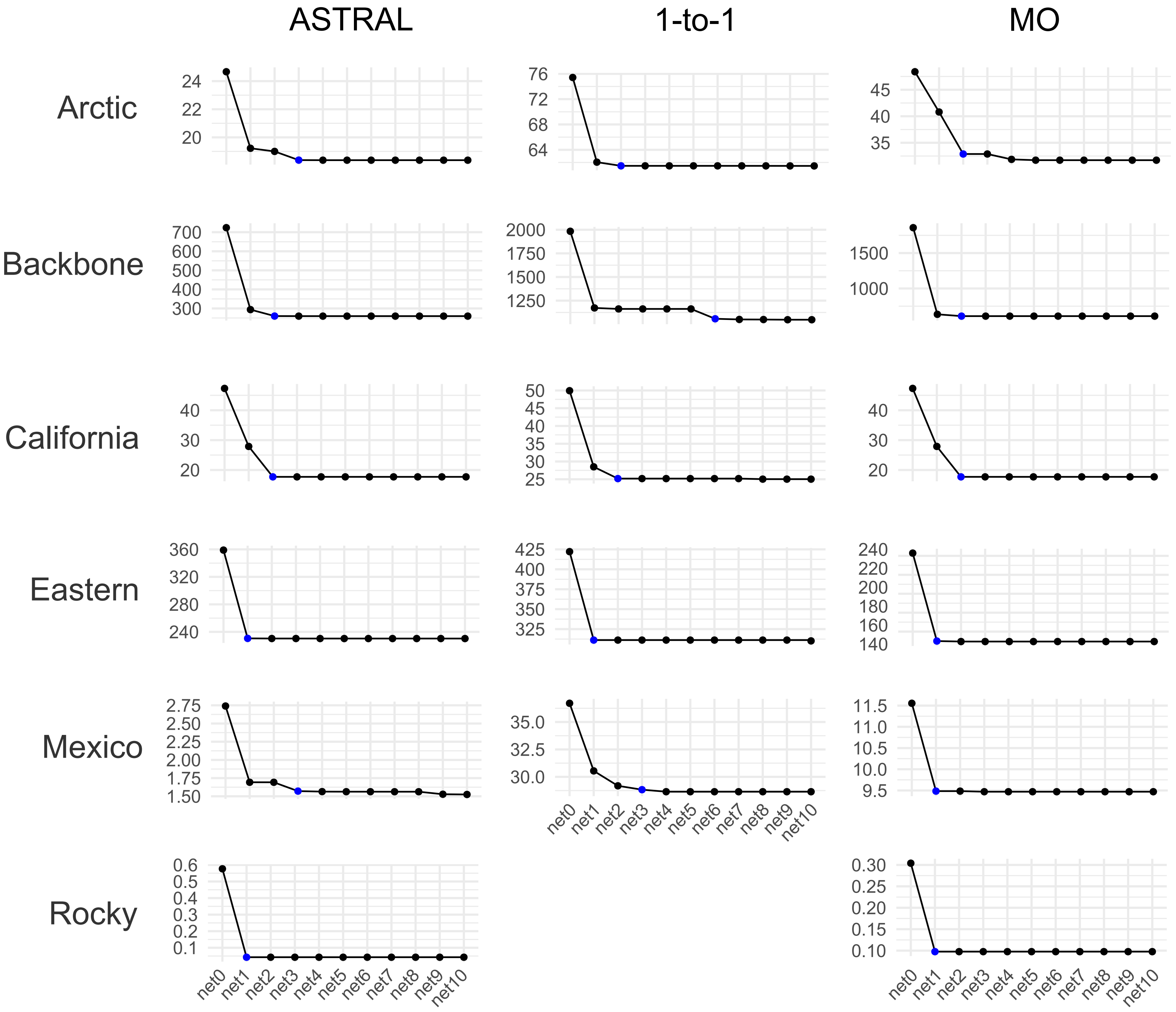
