## Supplementary Figure 8 for "Phylogenomic challenges in polyploid-rich lineages: Insights from paralog processing and reticulation methods using the complex genus *Packera* (Asteraceae: Senecioneae)"

### ASTRAL – Scenario 1

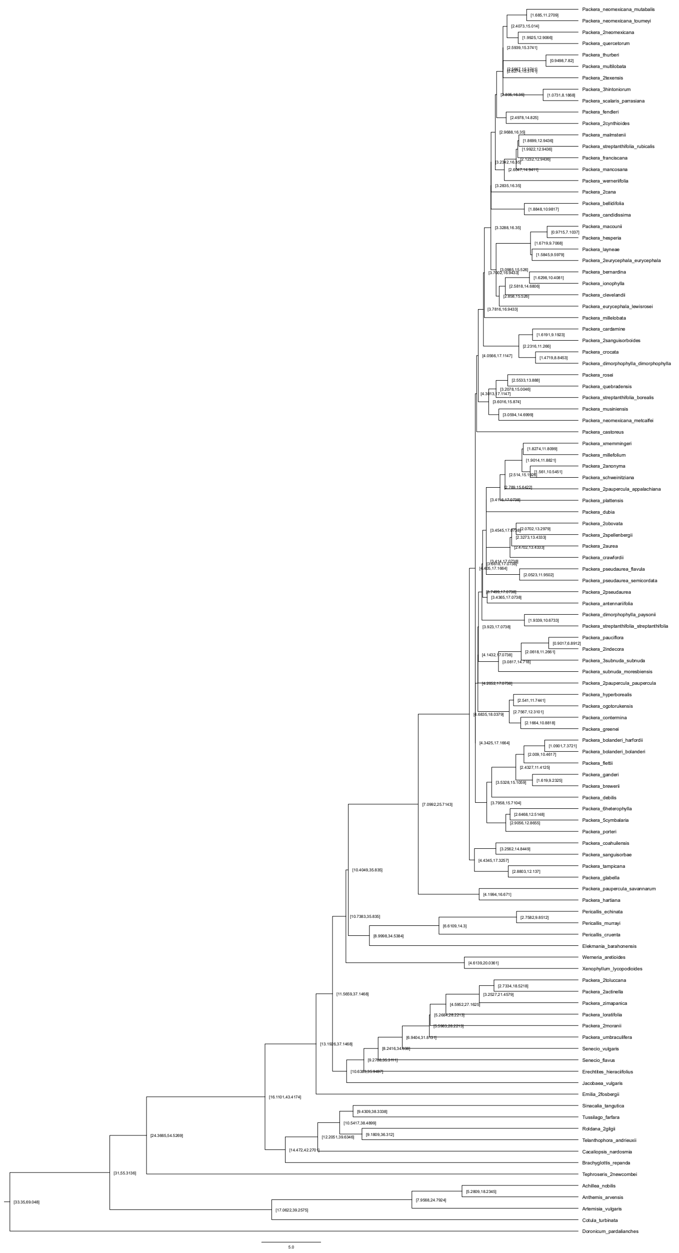


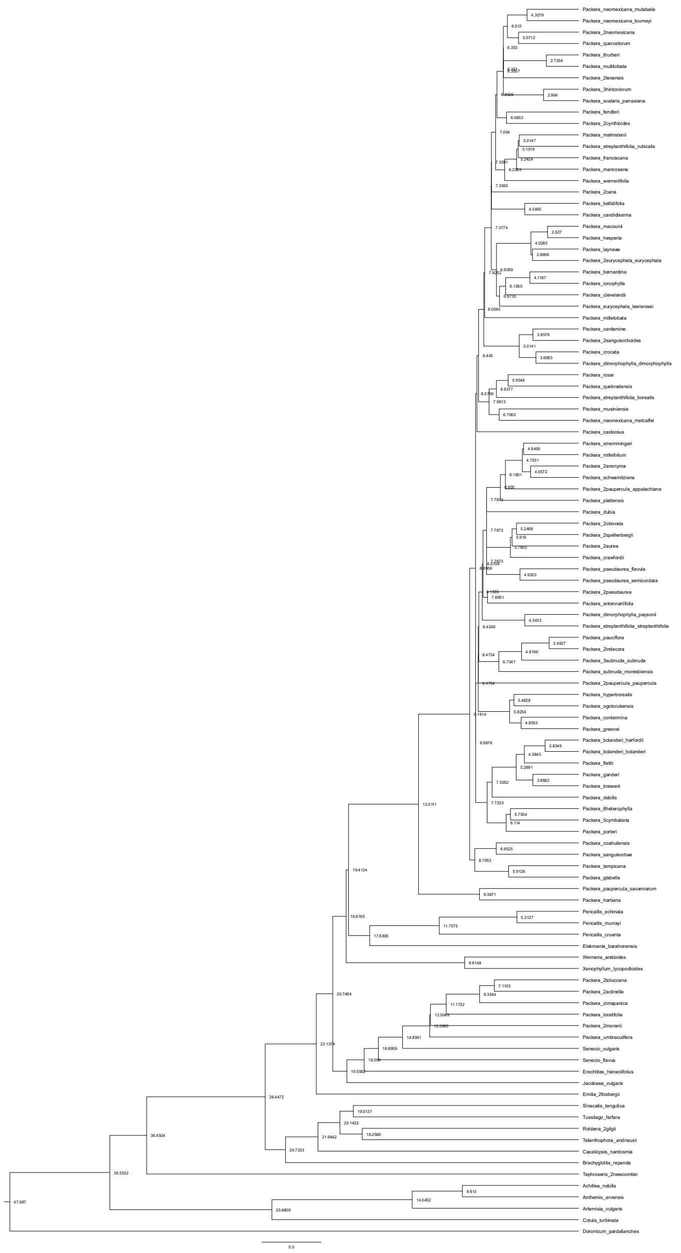

### ASTRAL – Scenario 2

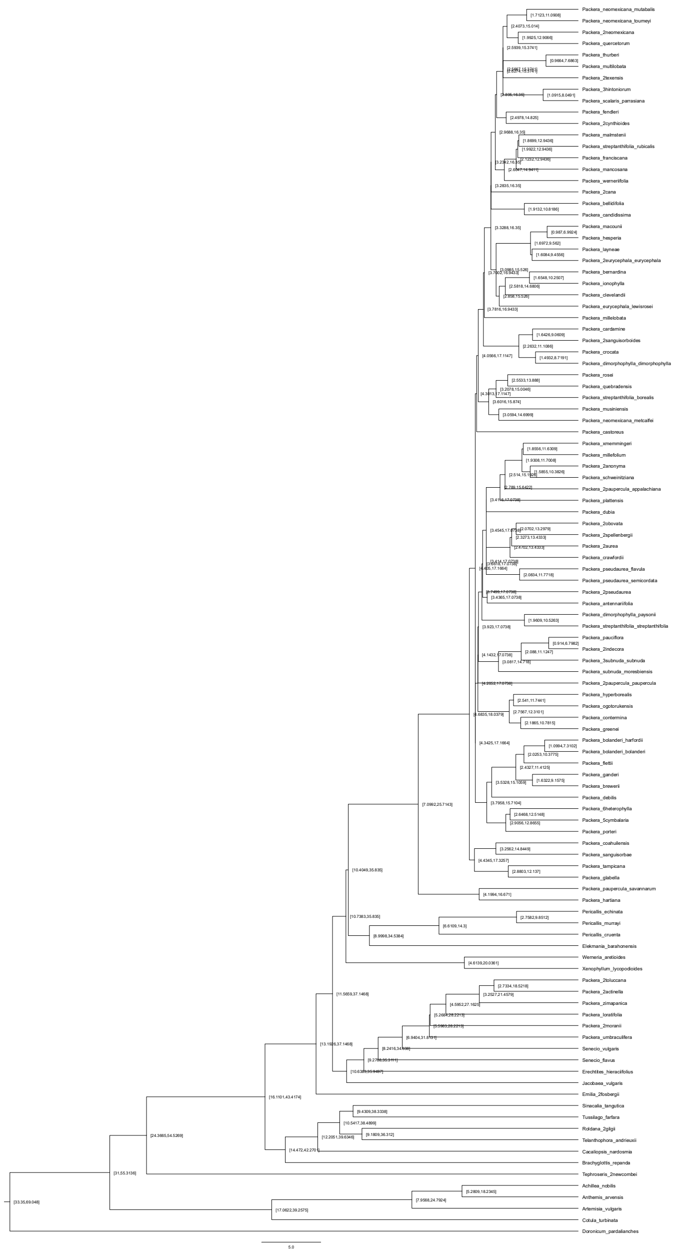

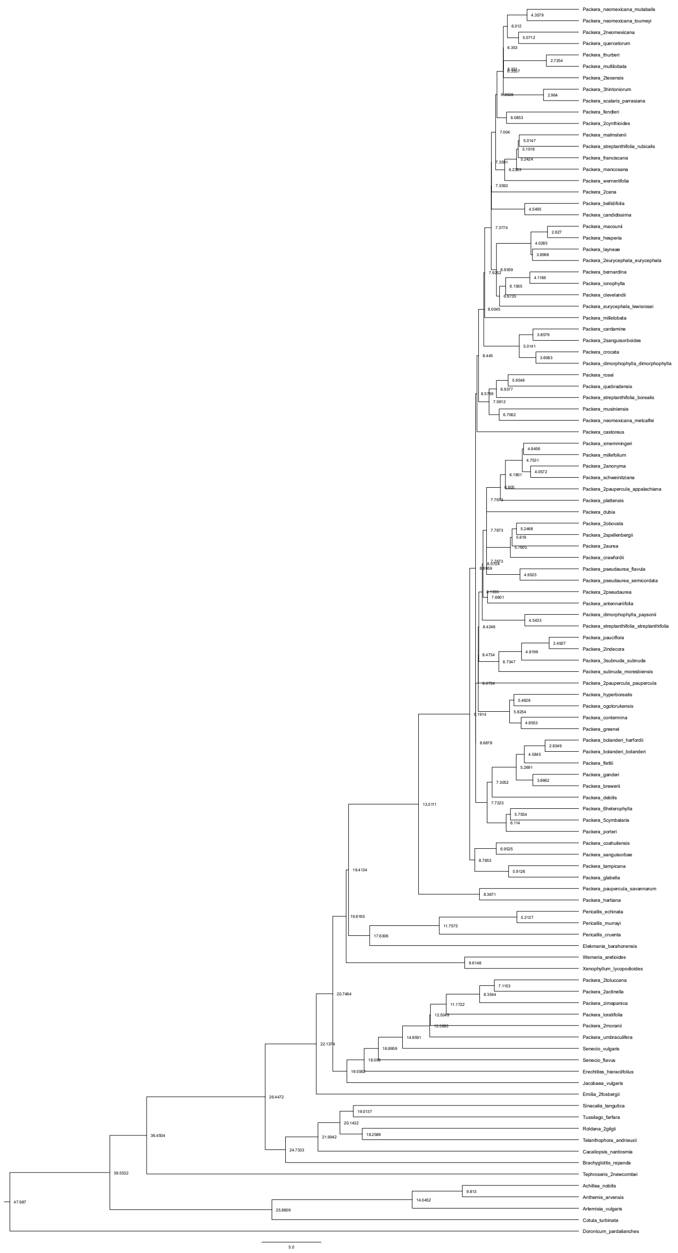

### ASTRAL – Scenario 3

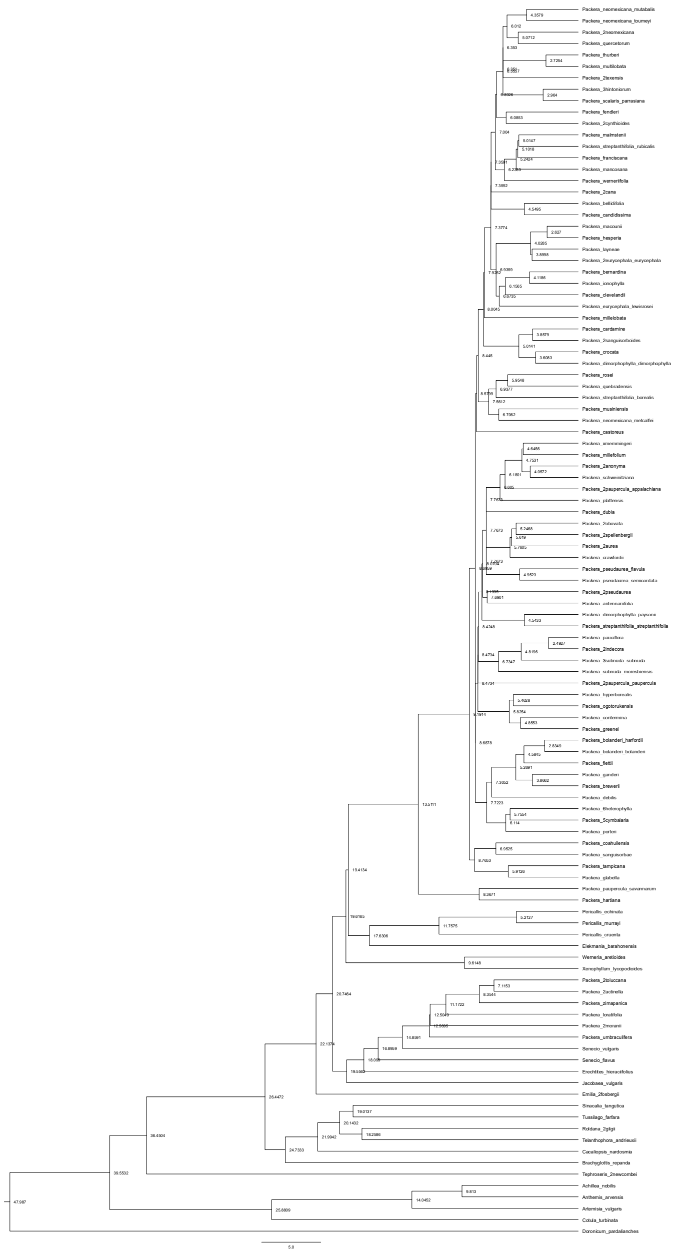

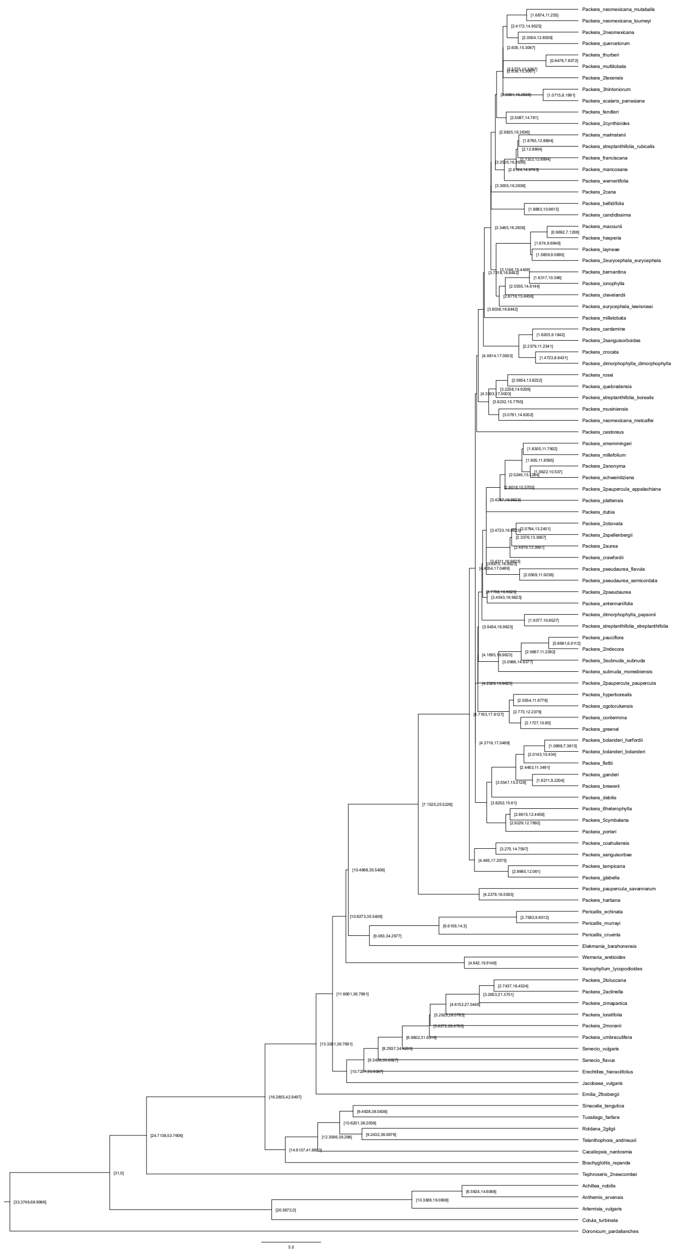

### ASTRAL – Scenario 4

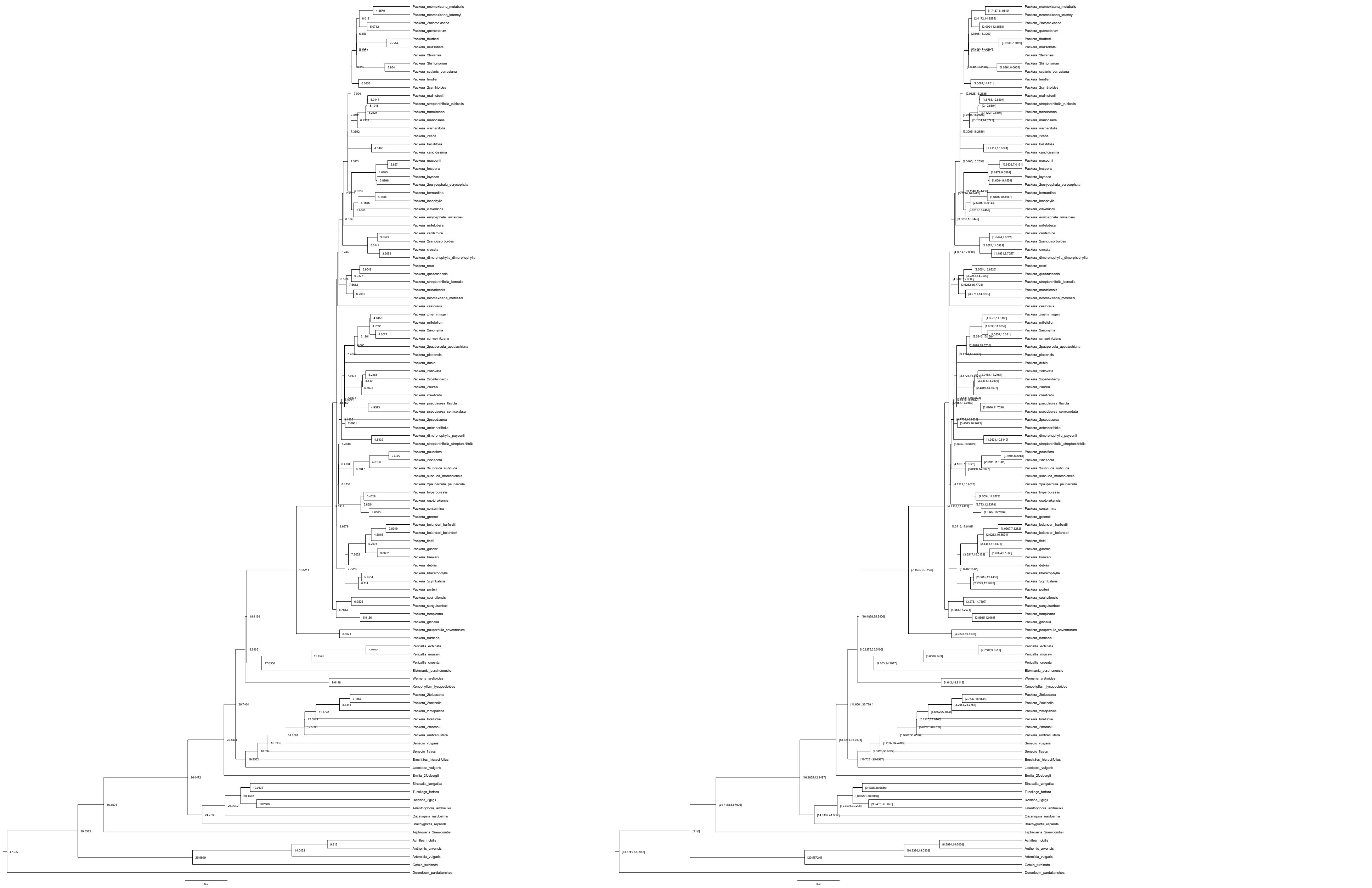

### ASTRAL – Scenario 5

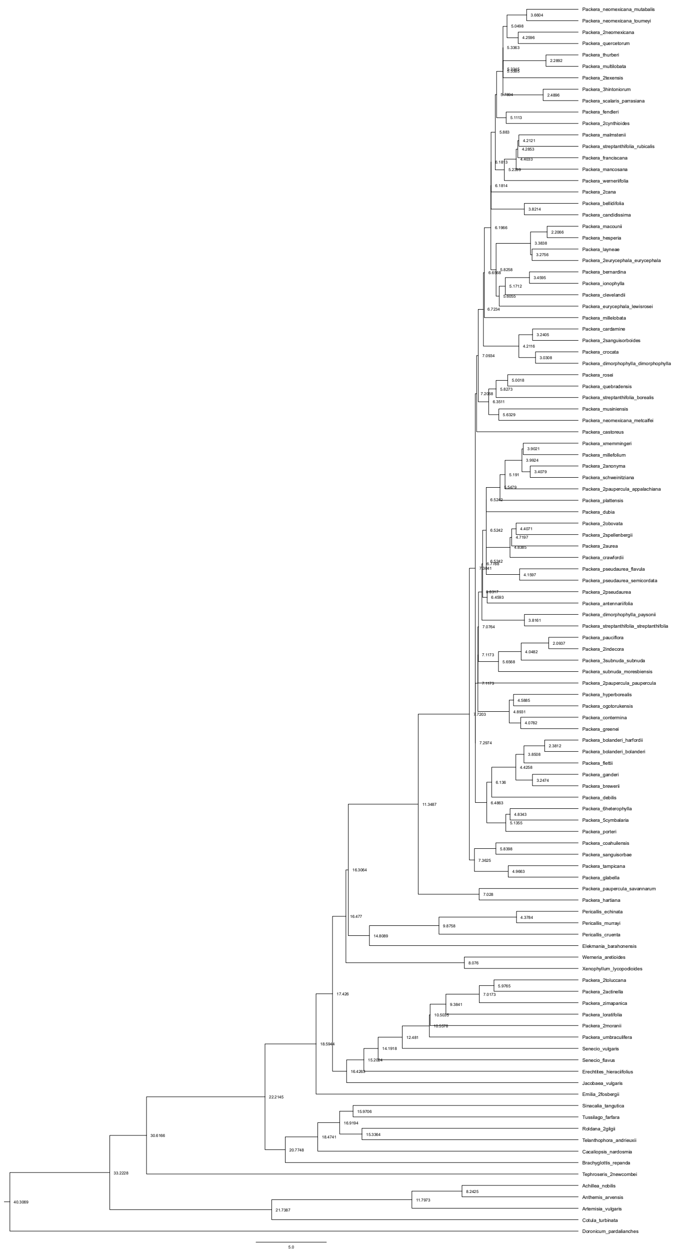

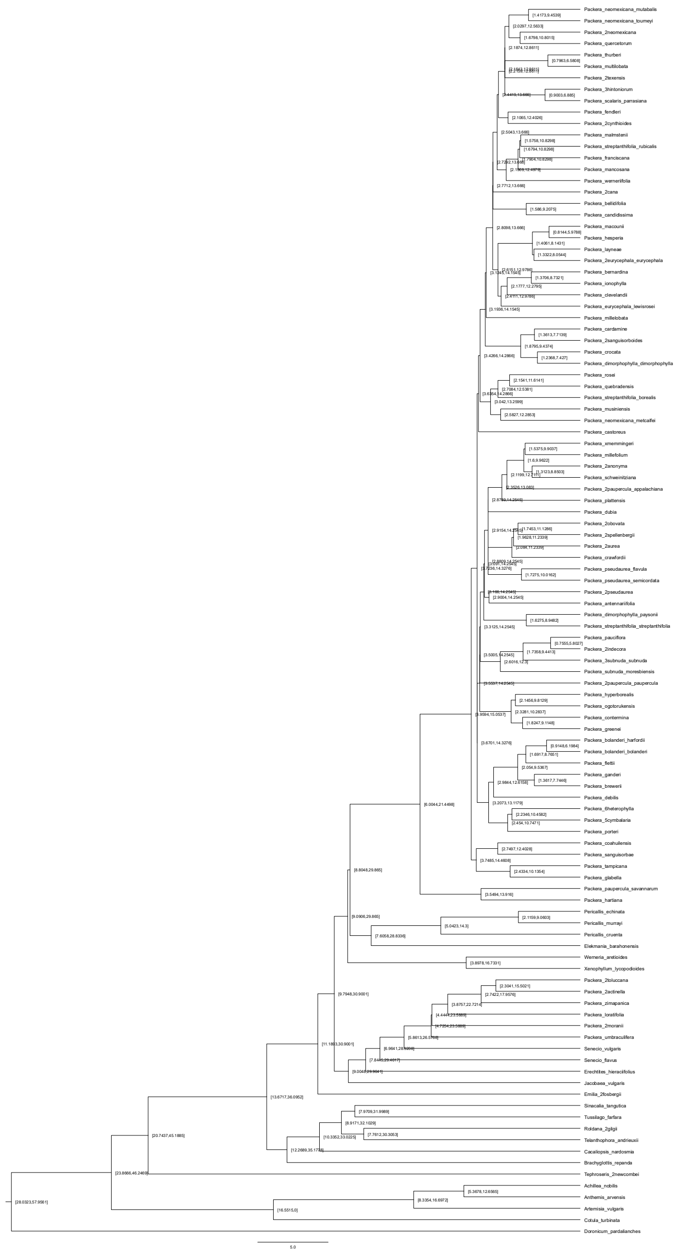

### ASTRAL – Scenario 6

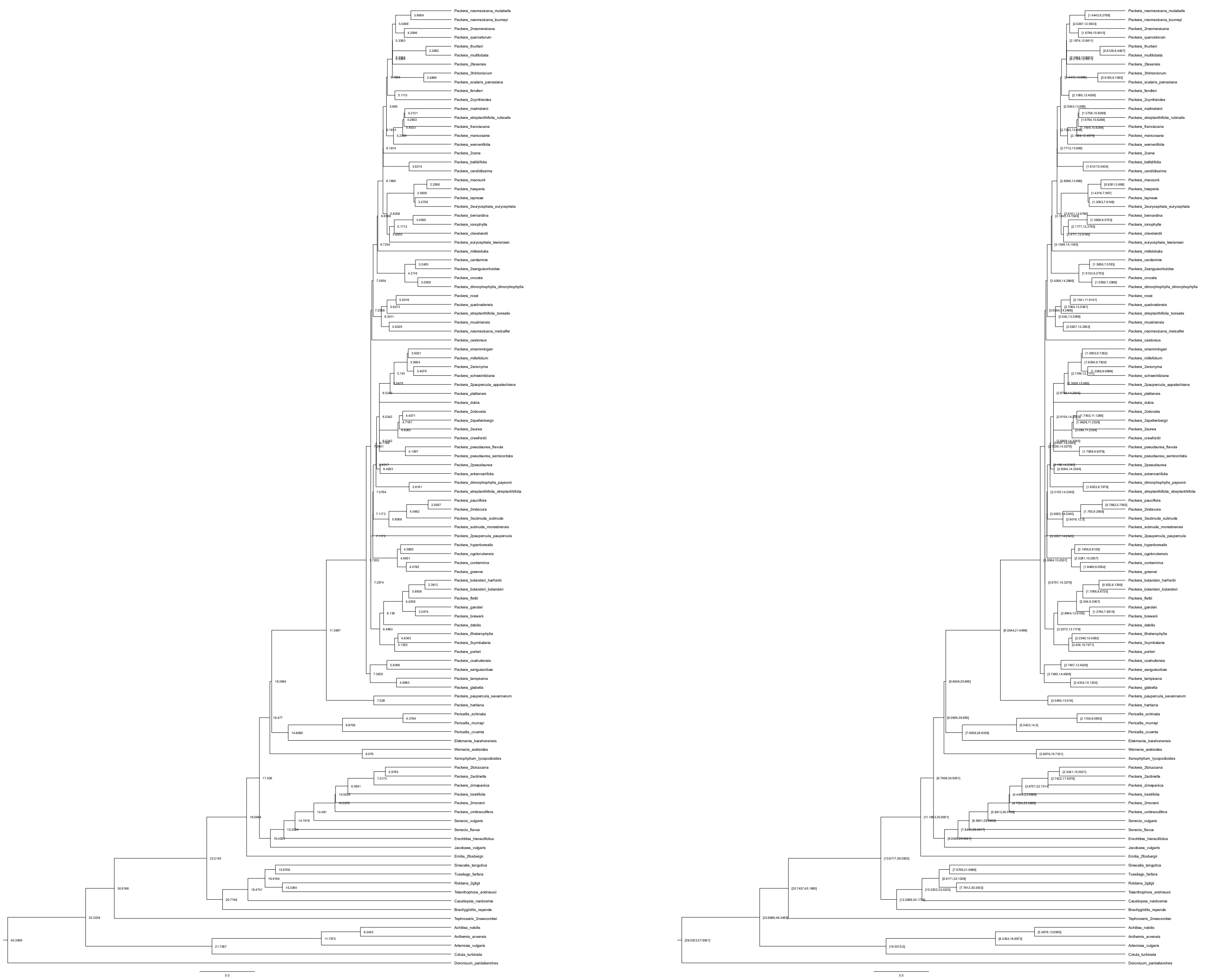

### paragone-nf: 1-to-1 – Scenario 1

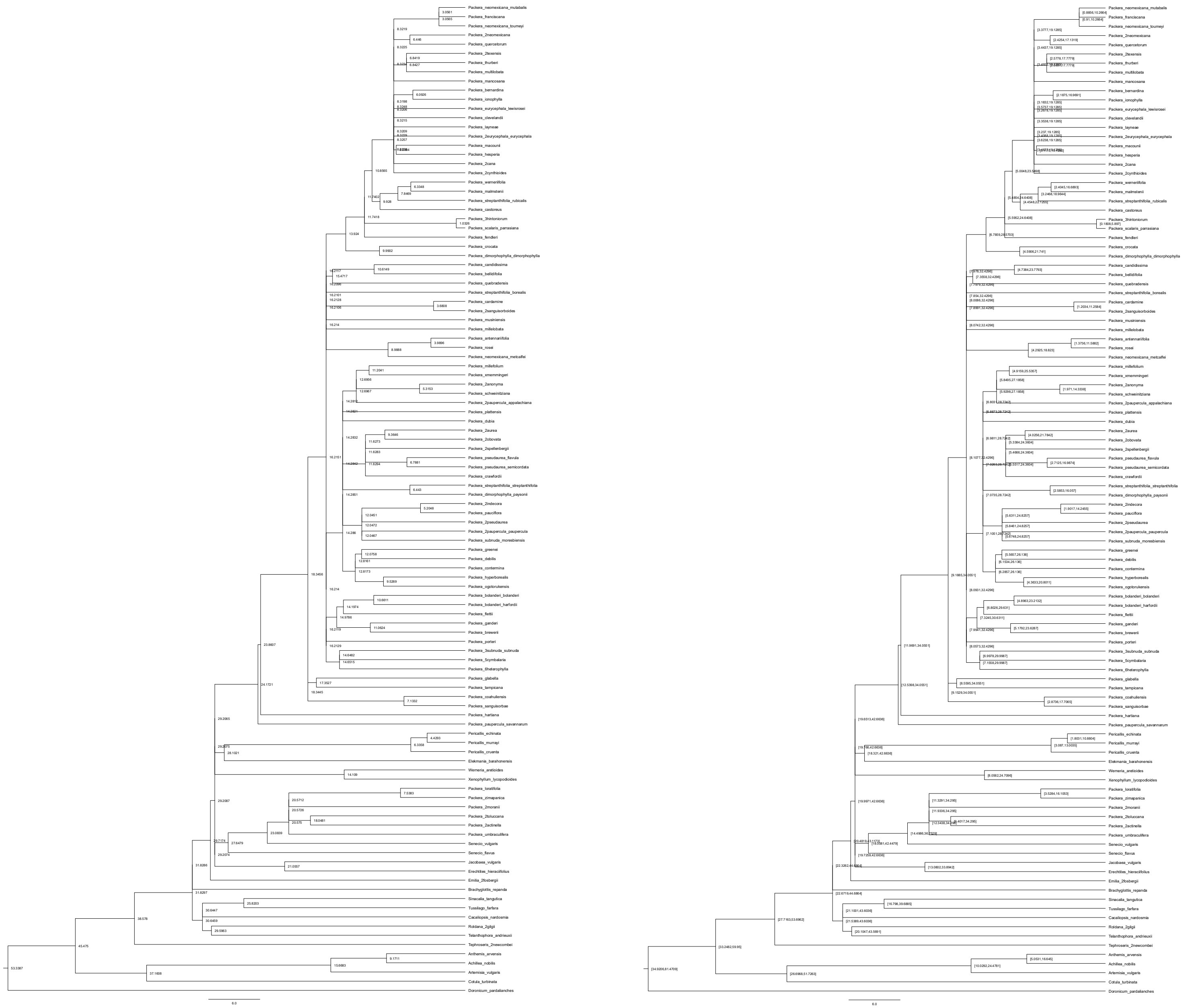

### paragone-nf: 1-to-1 – Scenario 2

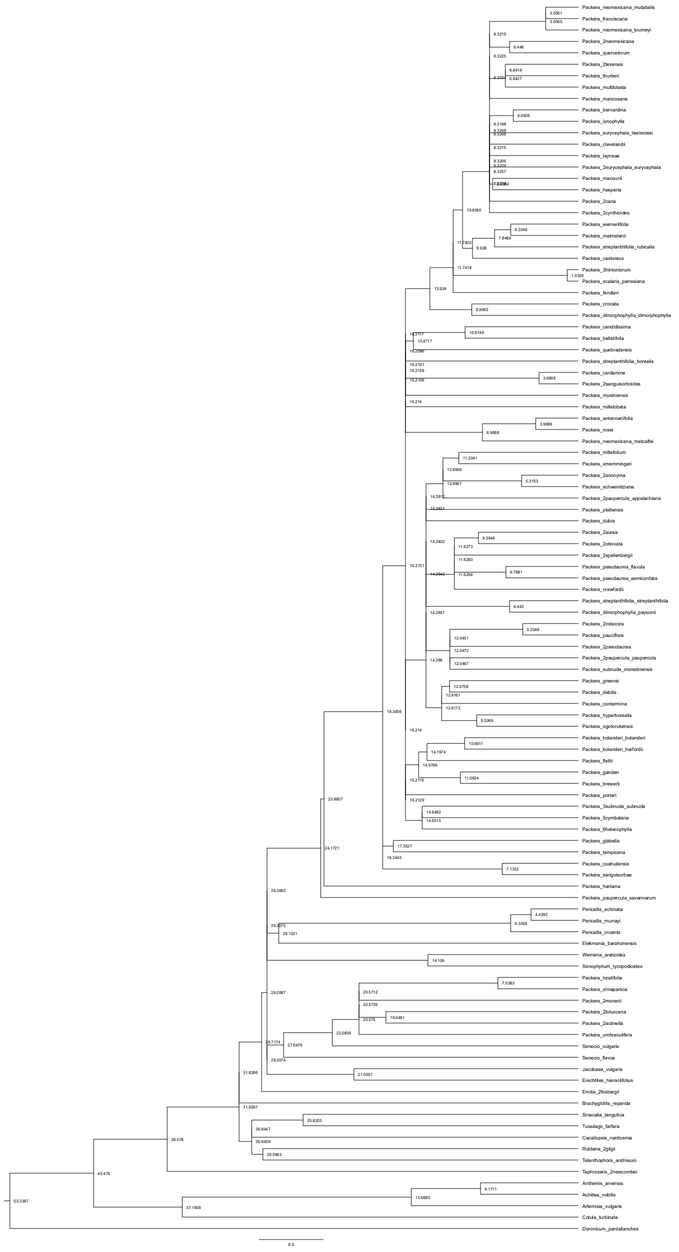

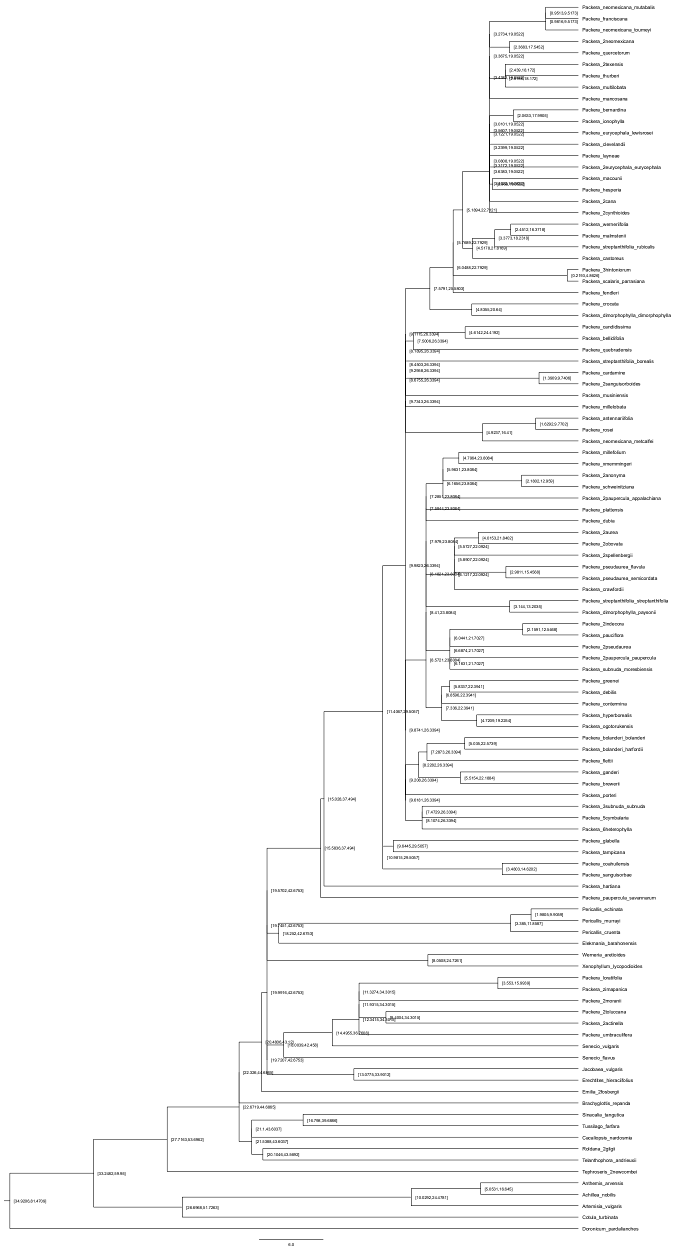

### paragone-nf: 1-to-1 – Scenario 3

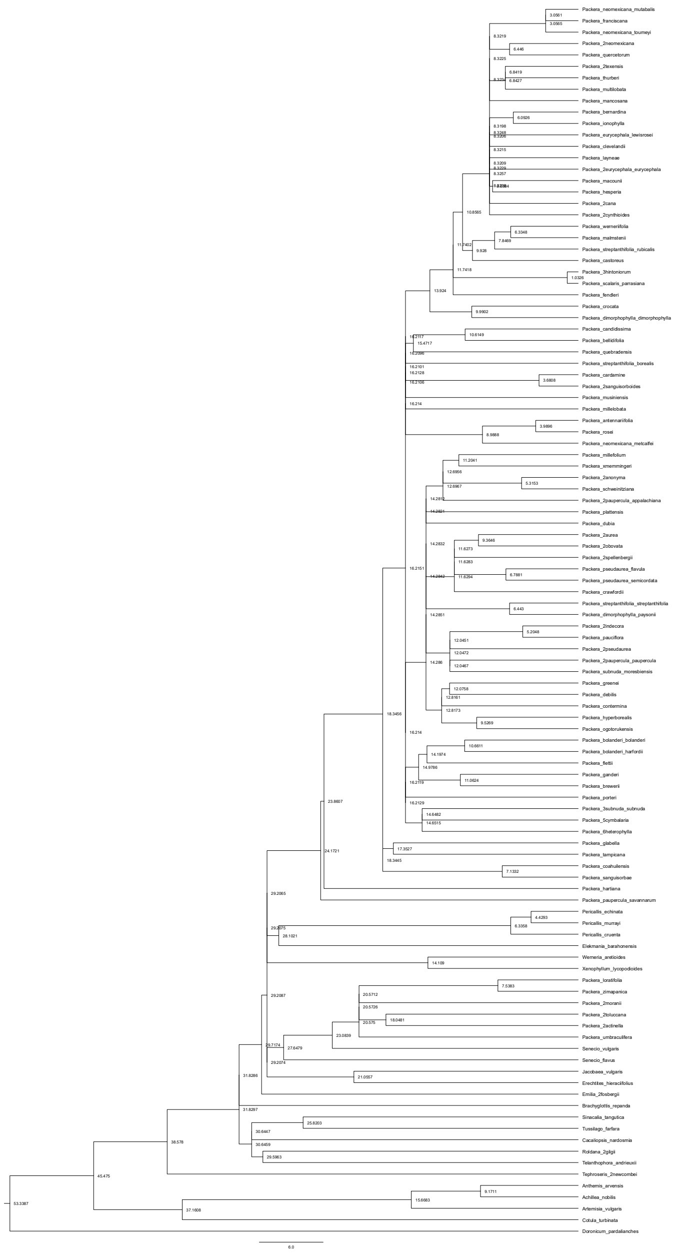

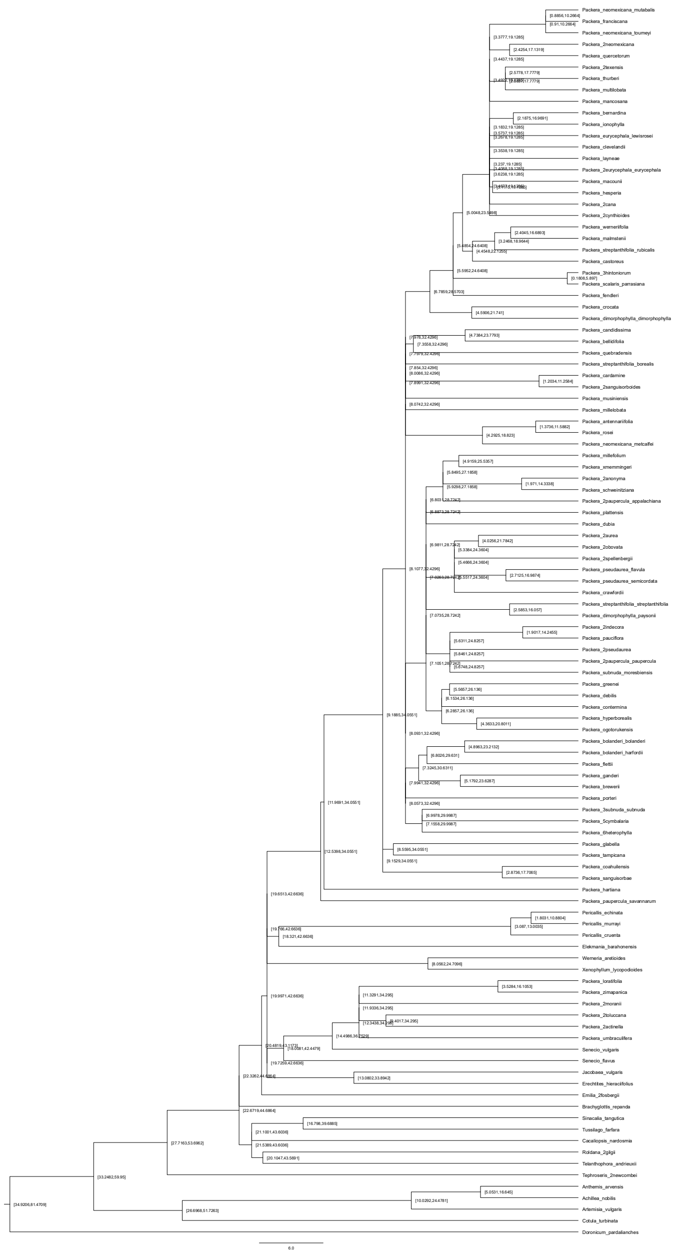

### paragone-nf: 1-to-1 – Scenario 4

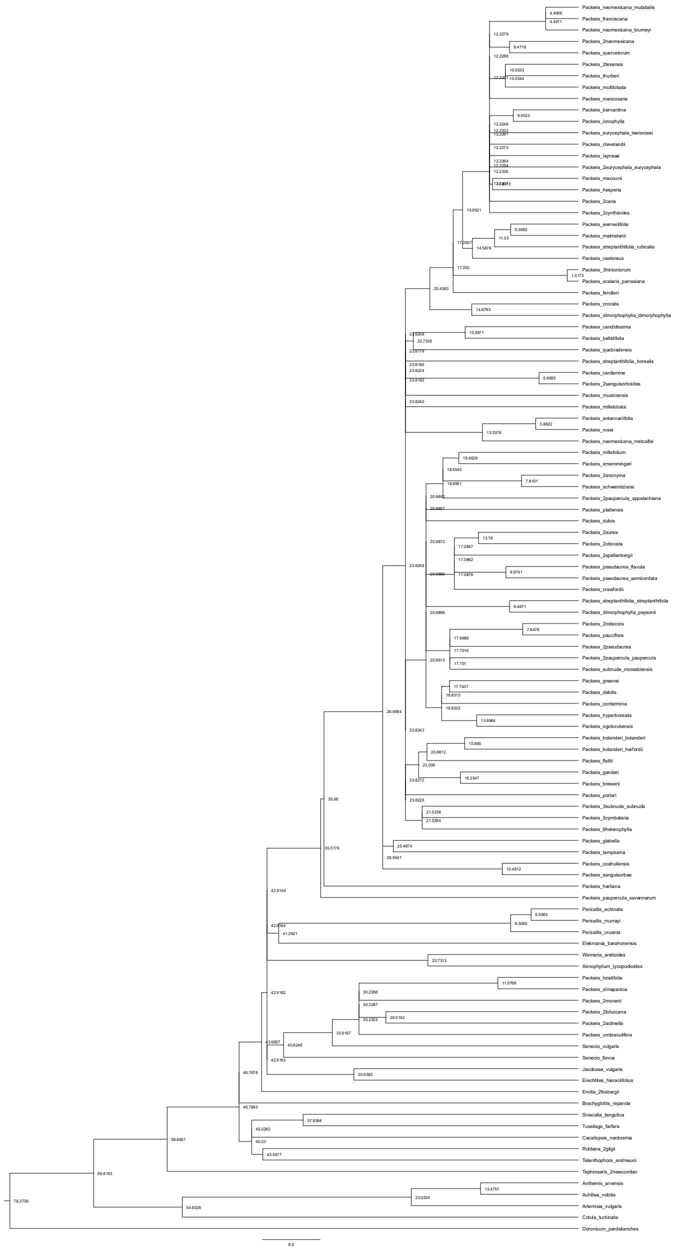


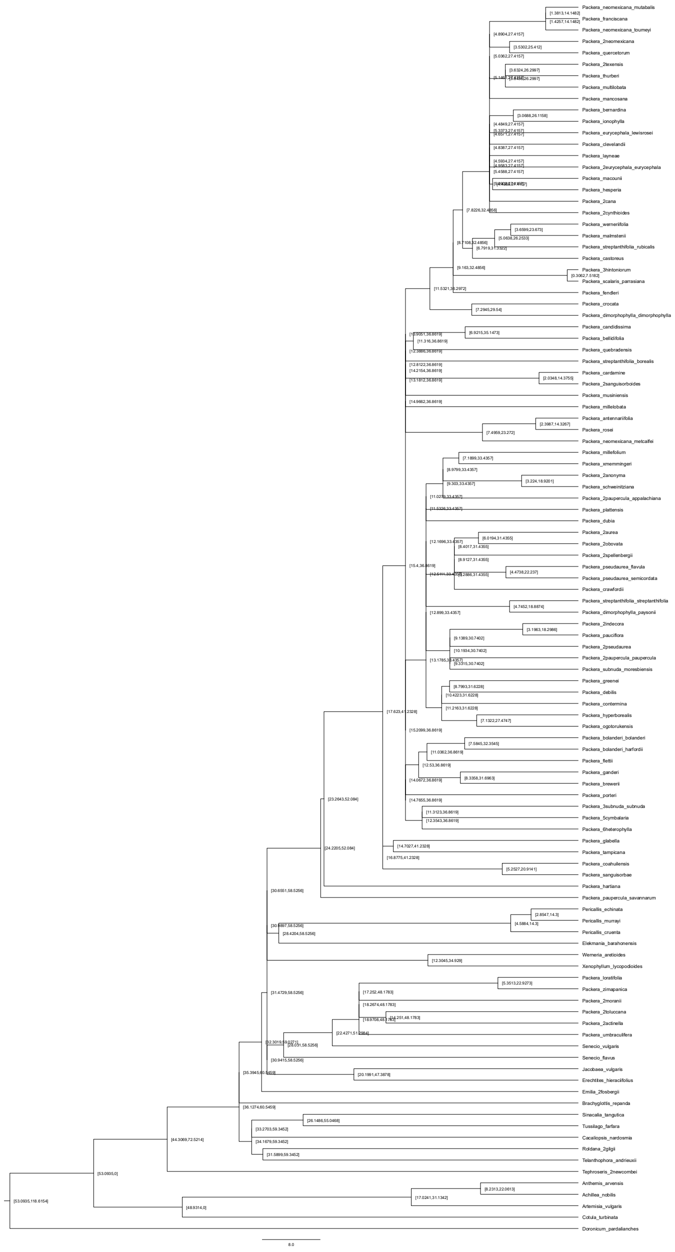

### paragone-nf: 1-to-1 – Scenario 5

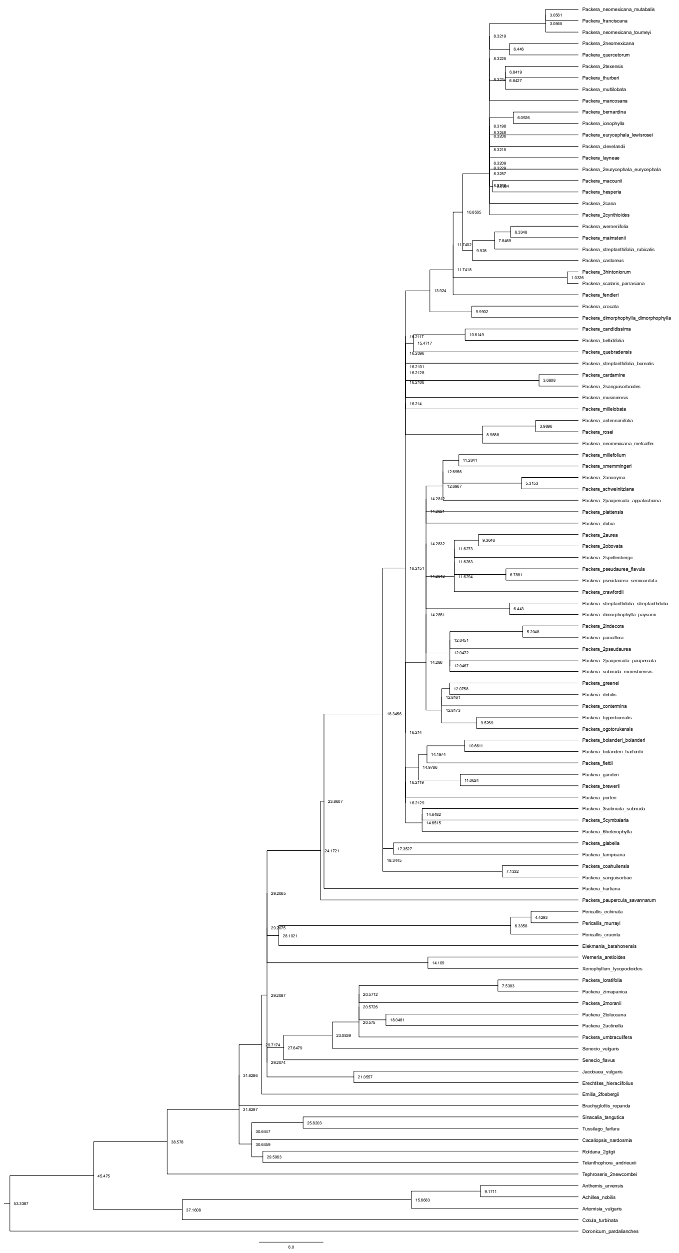


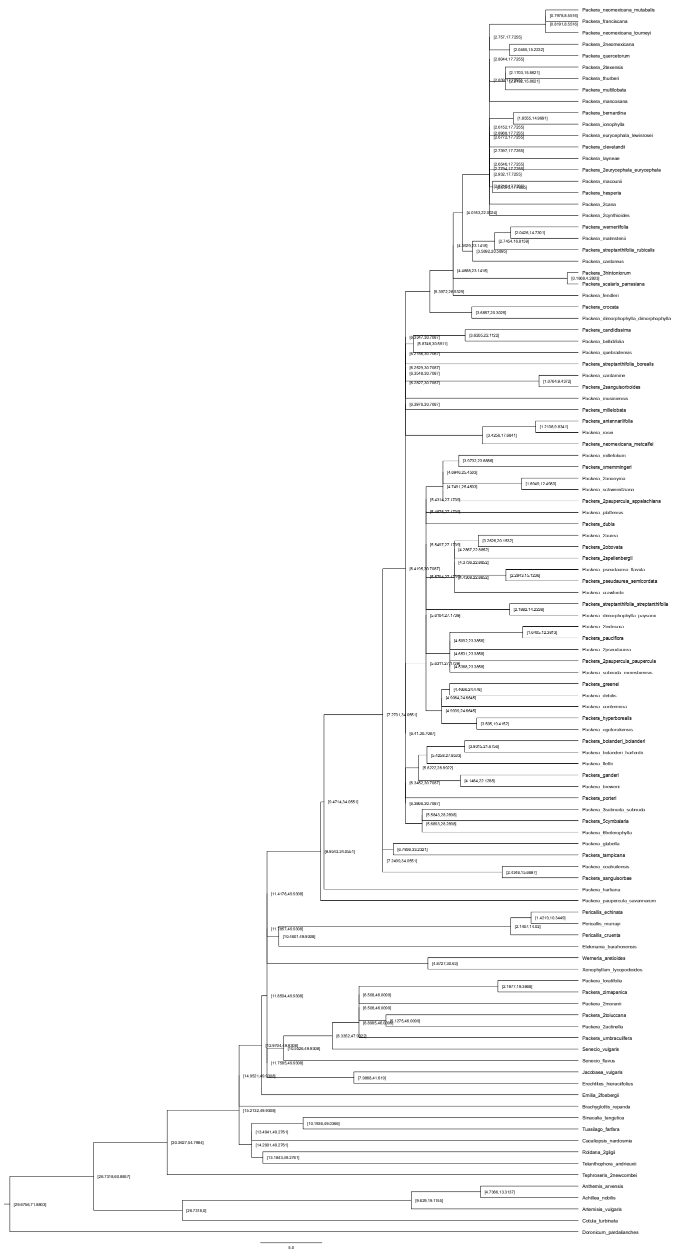

### paragone-nf: 1-to-1 – Scenario 6

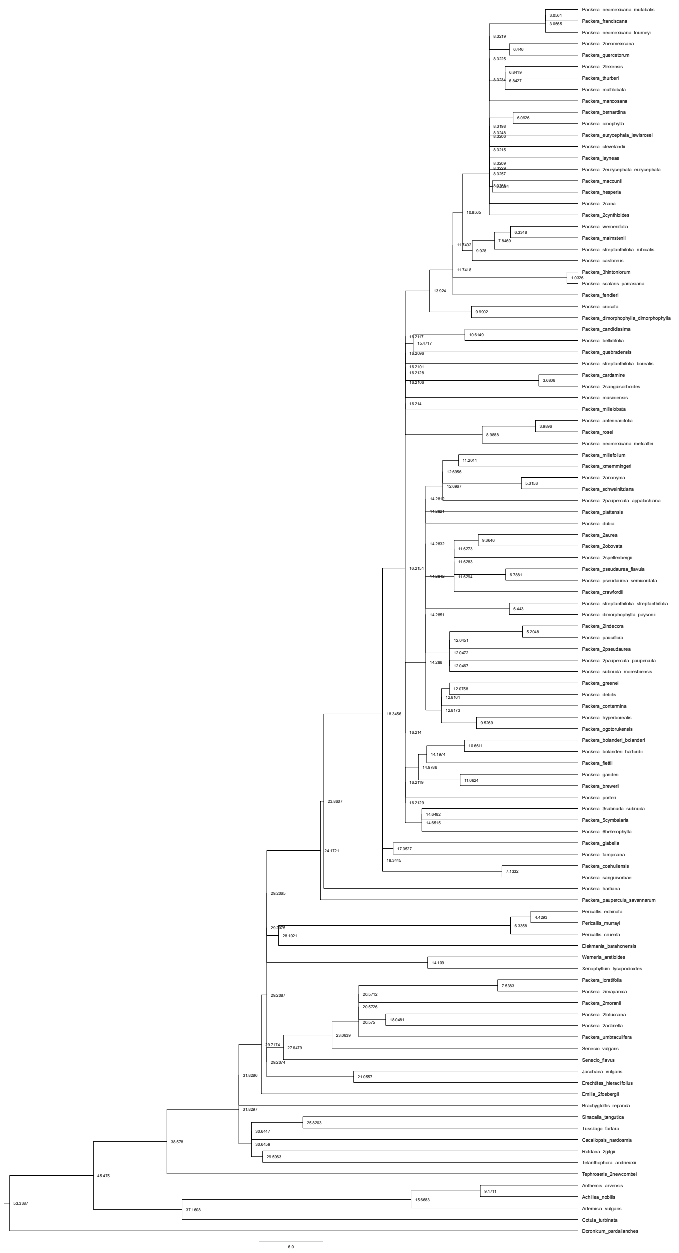


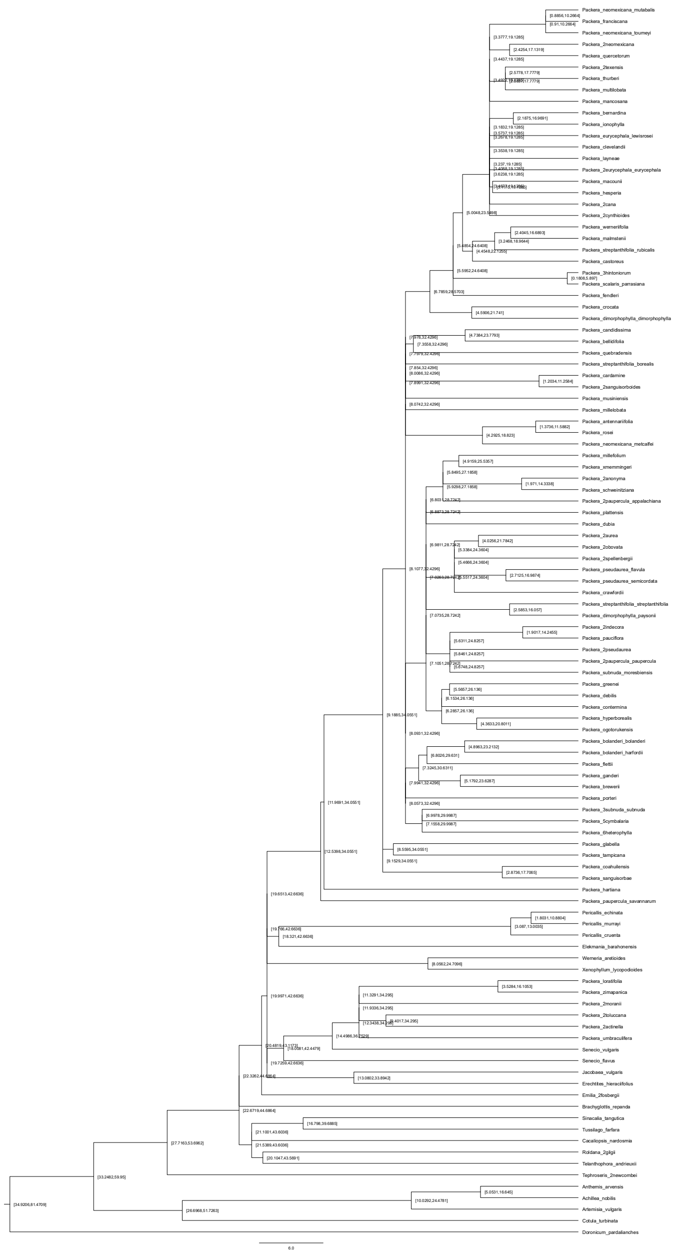

### paragone-nf: MO – Scenario 1

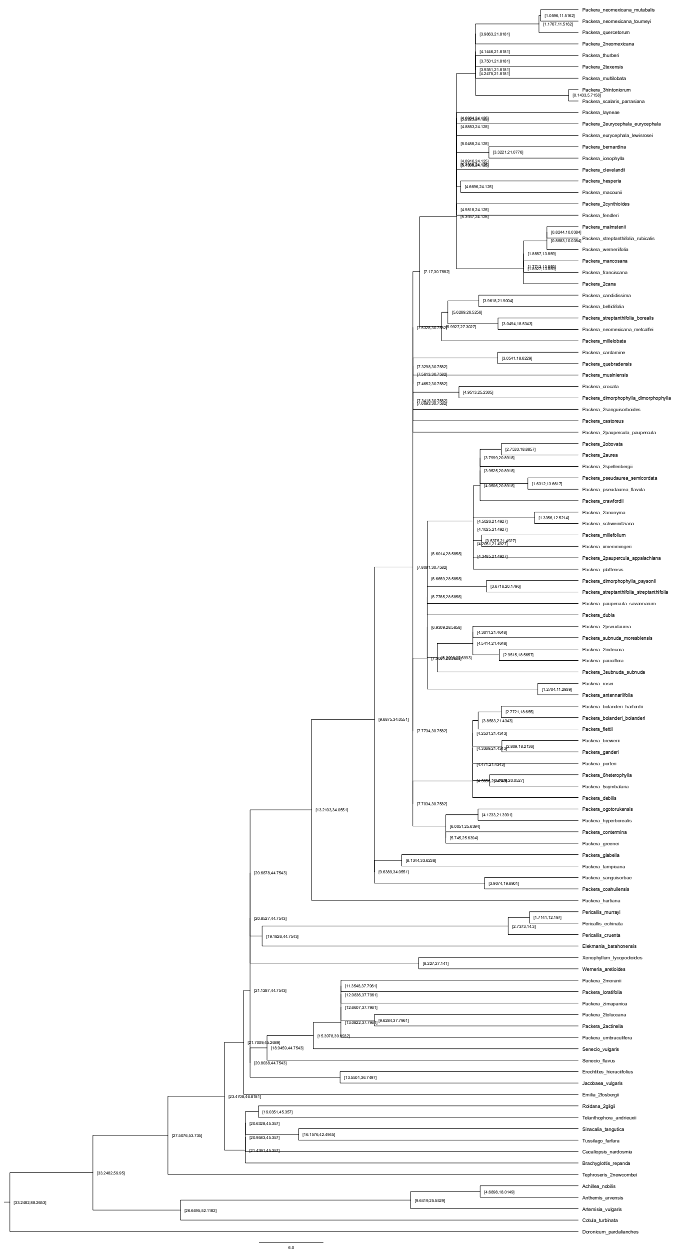

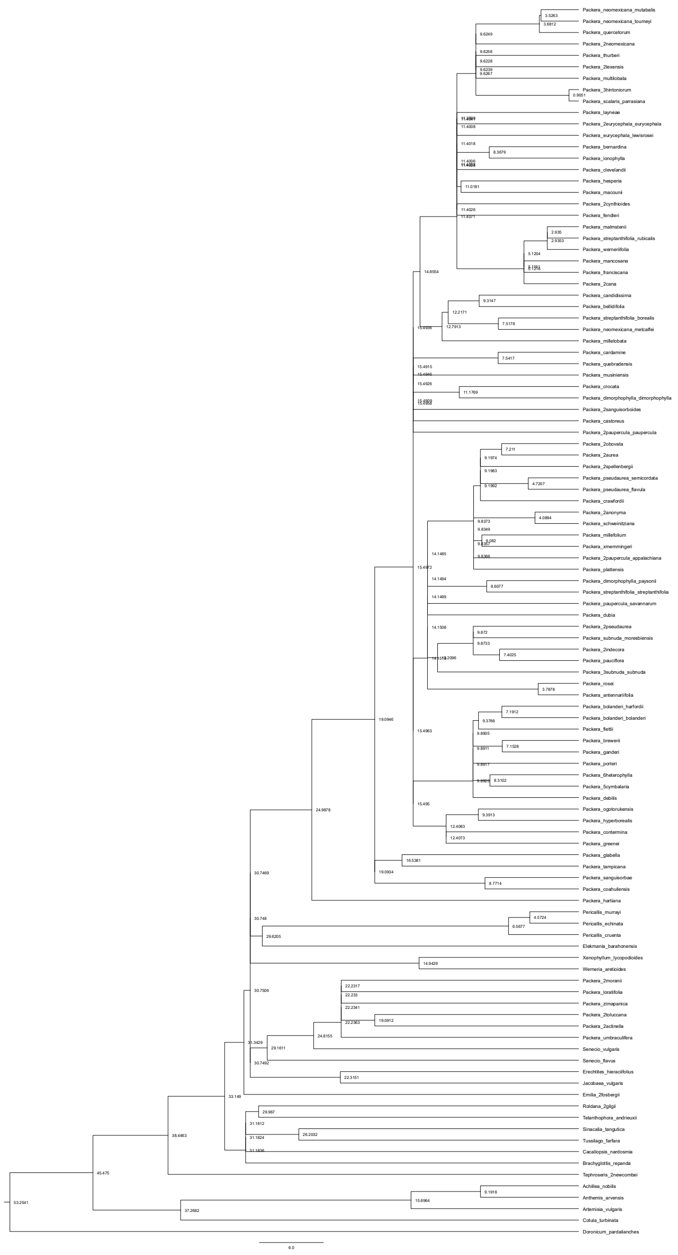

### paragone-nf: MO – Scenario 2

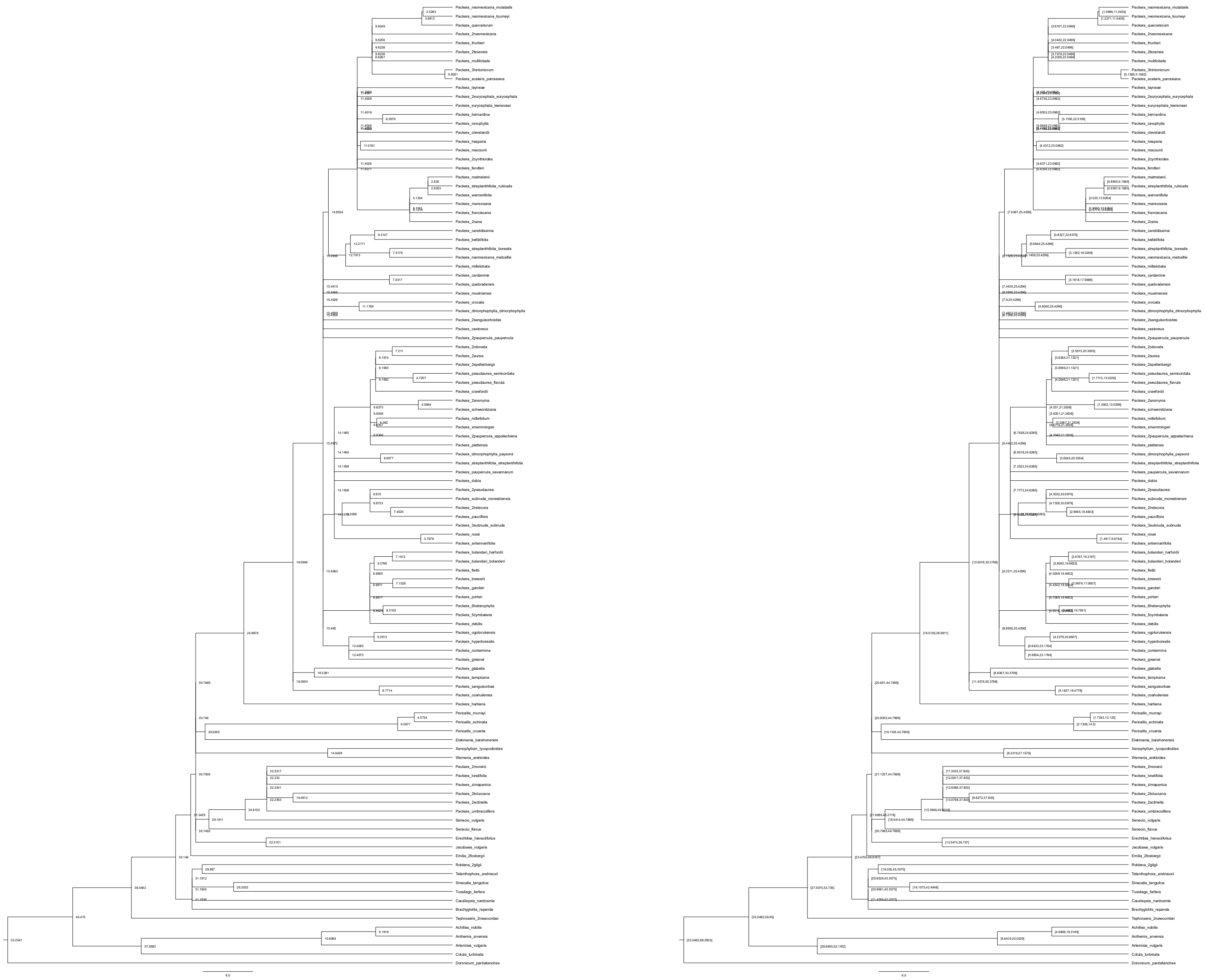

### paragone-nf: MO – Scenario 3

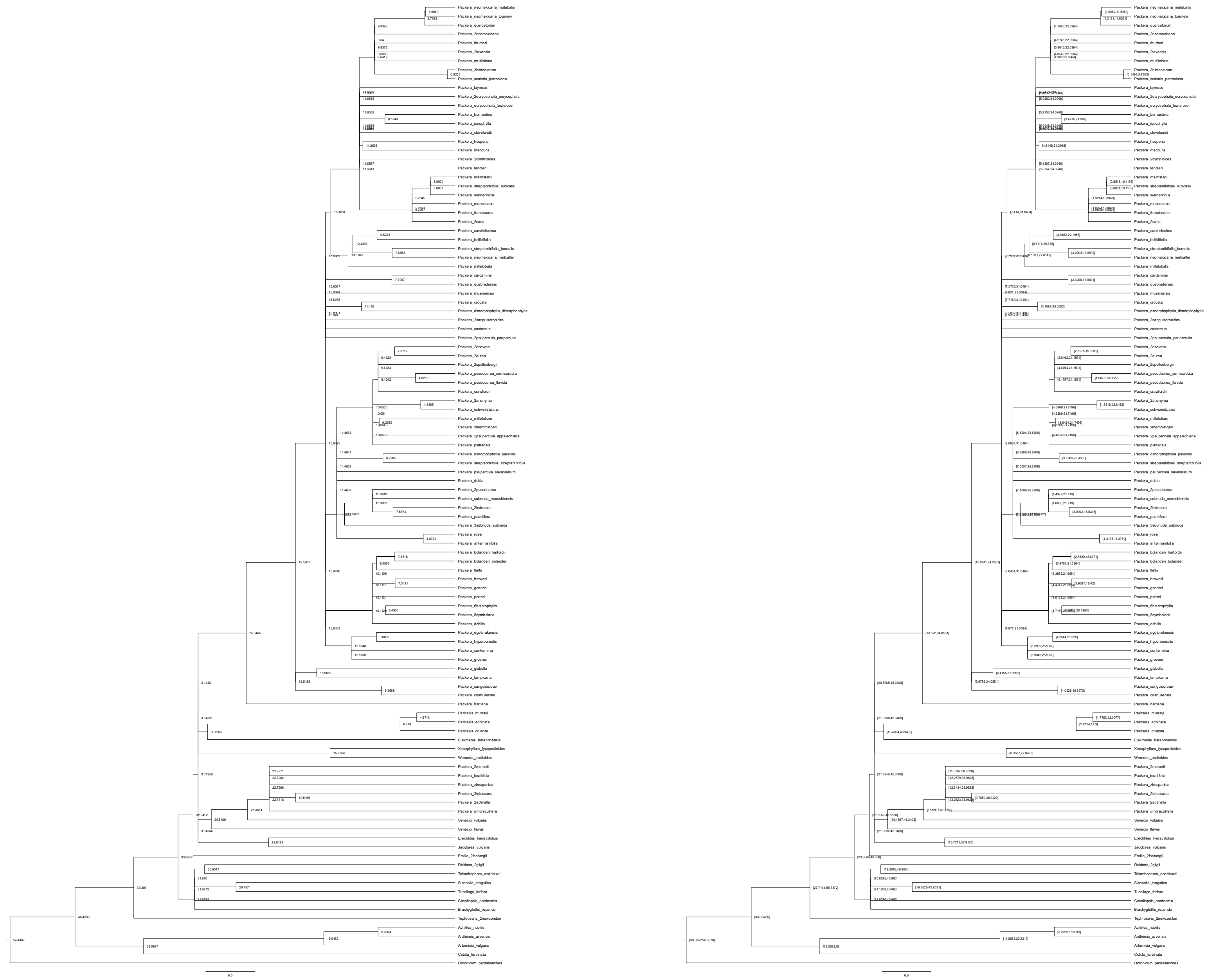

### paragone-nf: MO – Scenario 4

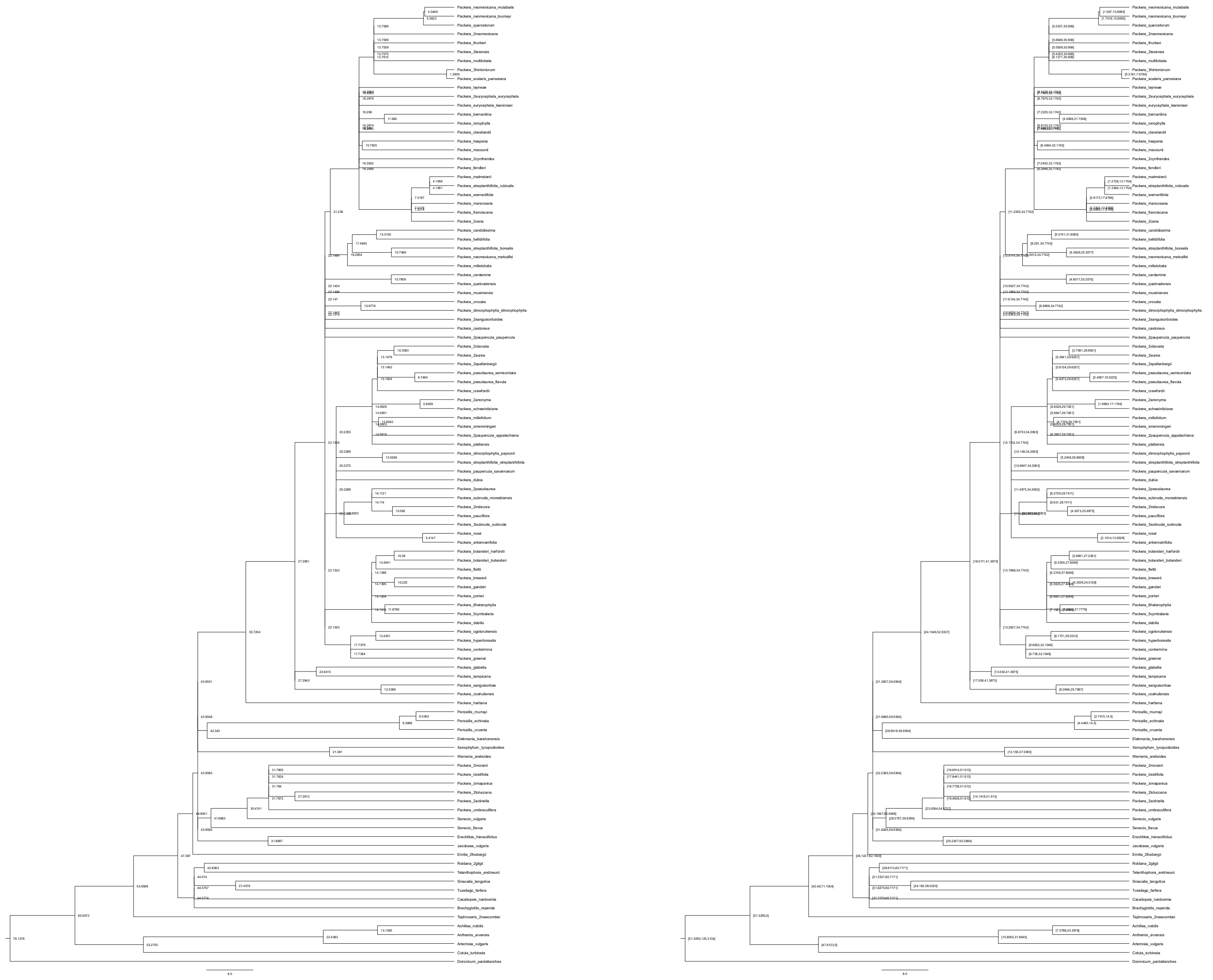

### paragone-nf: MO – Scenario 5

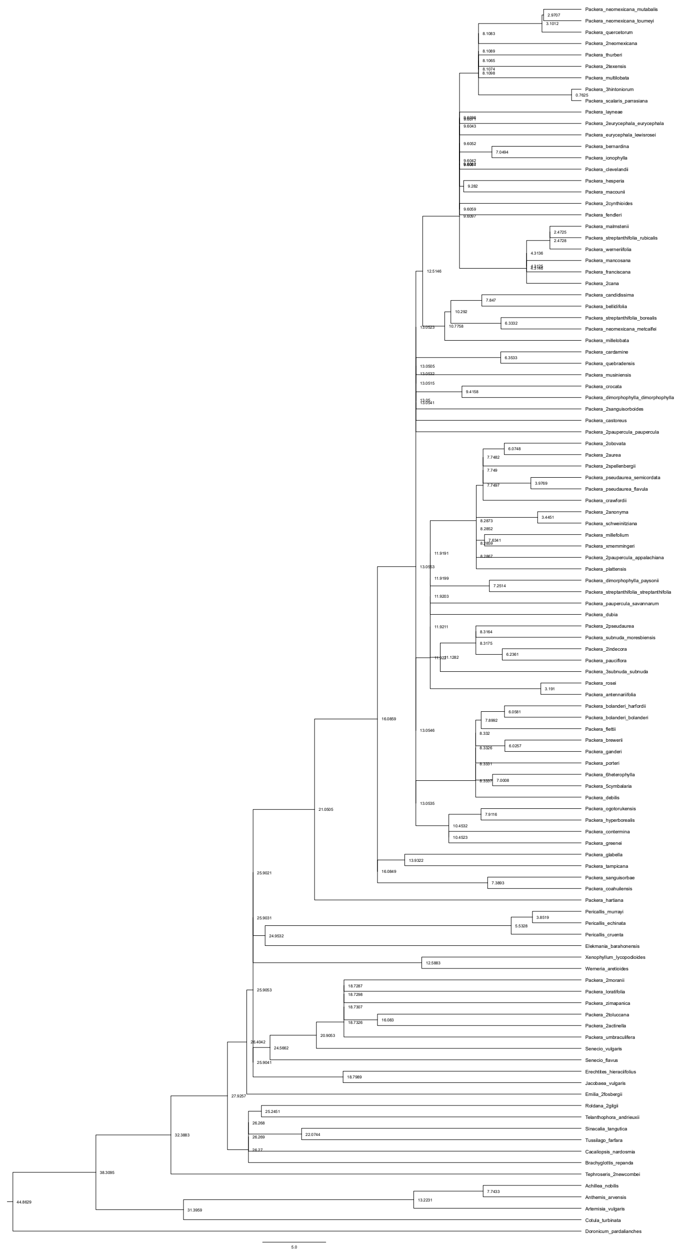

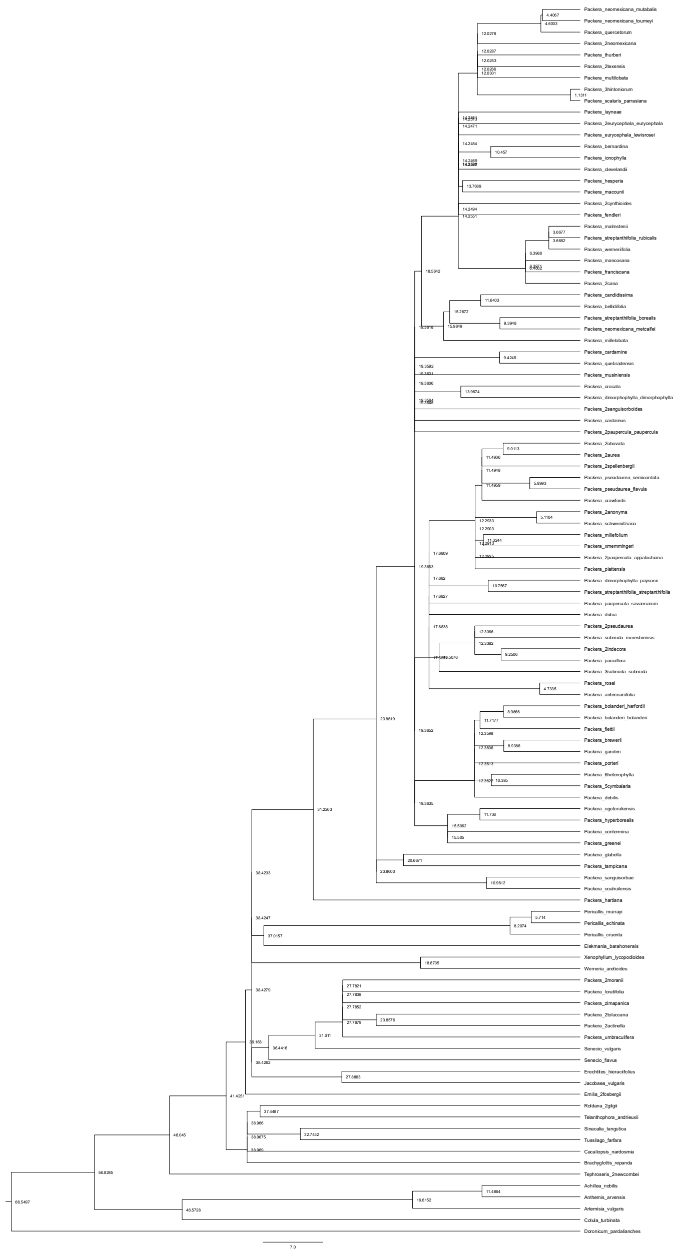

### paragone-nf: MO – Scenario 6

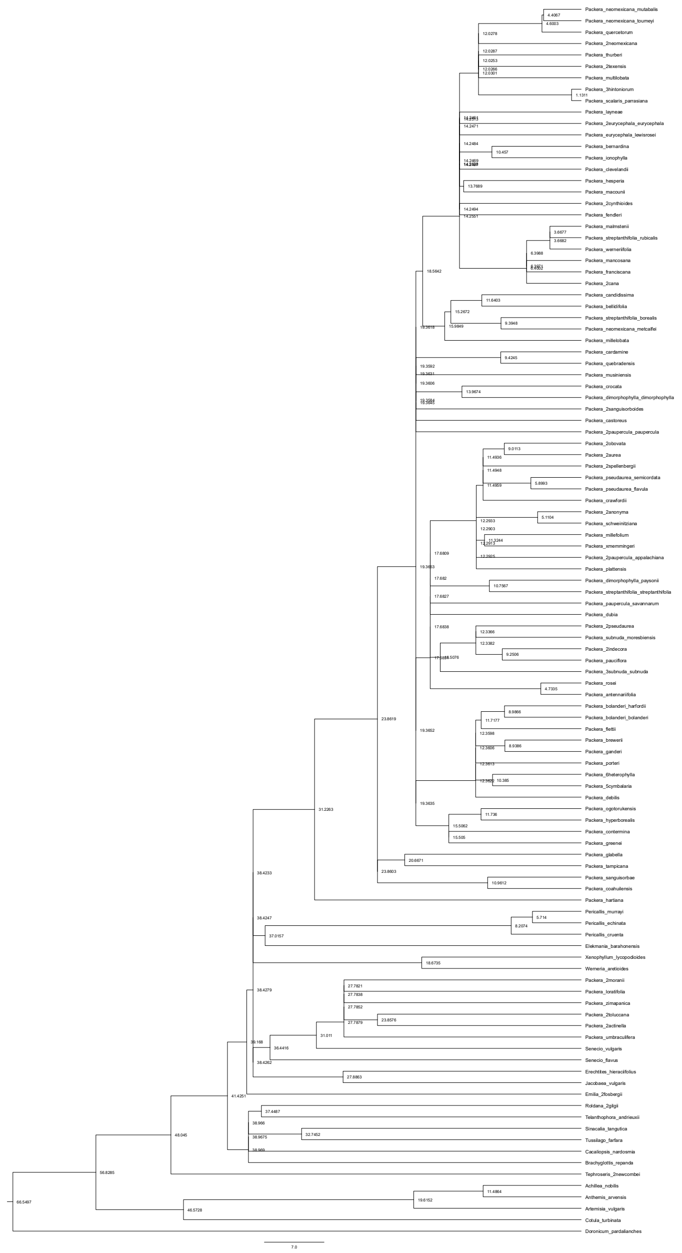

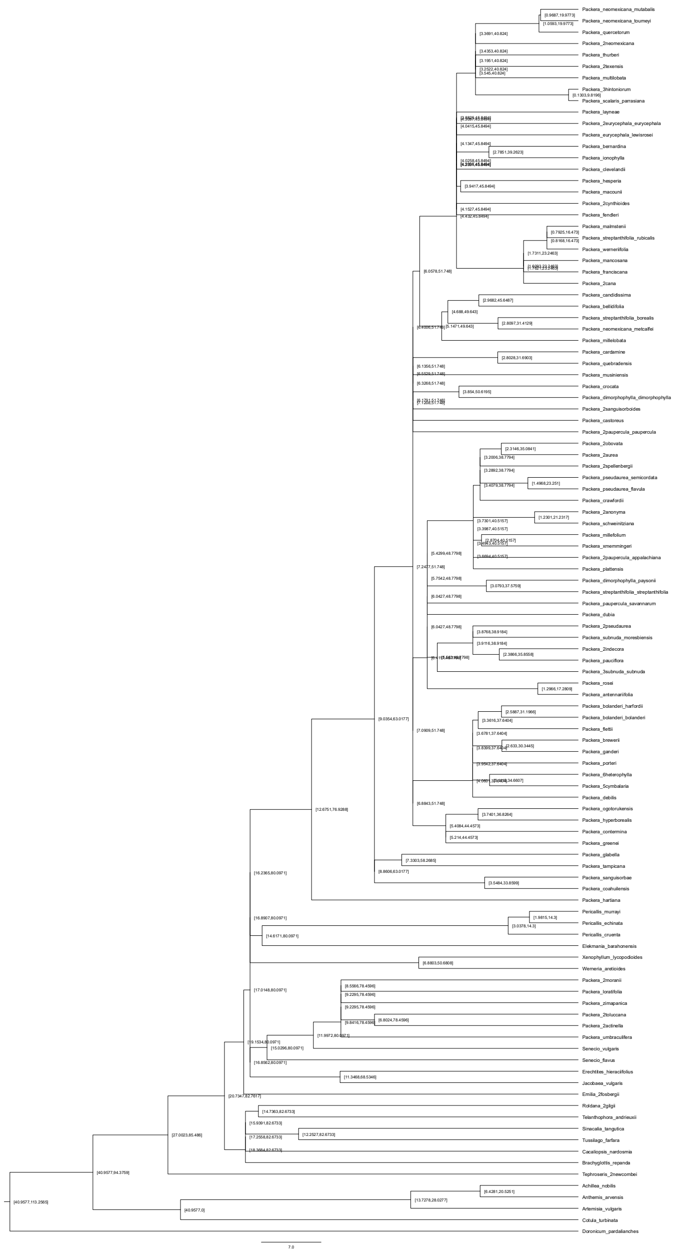
