## Supplementary Figure 9 for "Phylogenomic challenges in polyploid-rich lineages: Insights from paralog processing and reticulation methods using the complex genus *Packera* (Asteraceae: Senecioneae)"

**BioGeoBEARS DEC on Packera − HybPiper**
**ancstates: global optim, 4 areas max. d=0.0101; e=0.0038; j=0; LnL=−300.65**

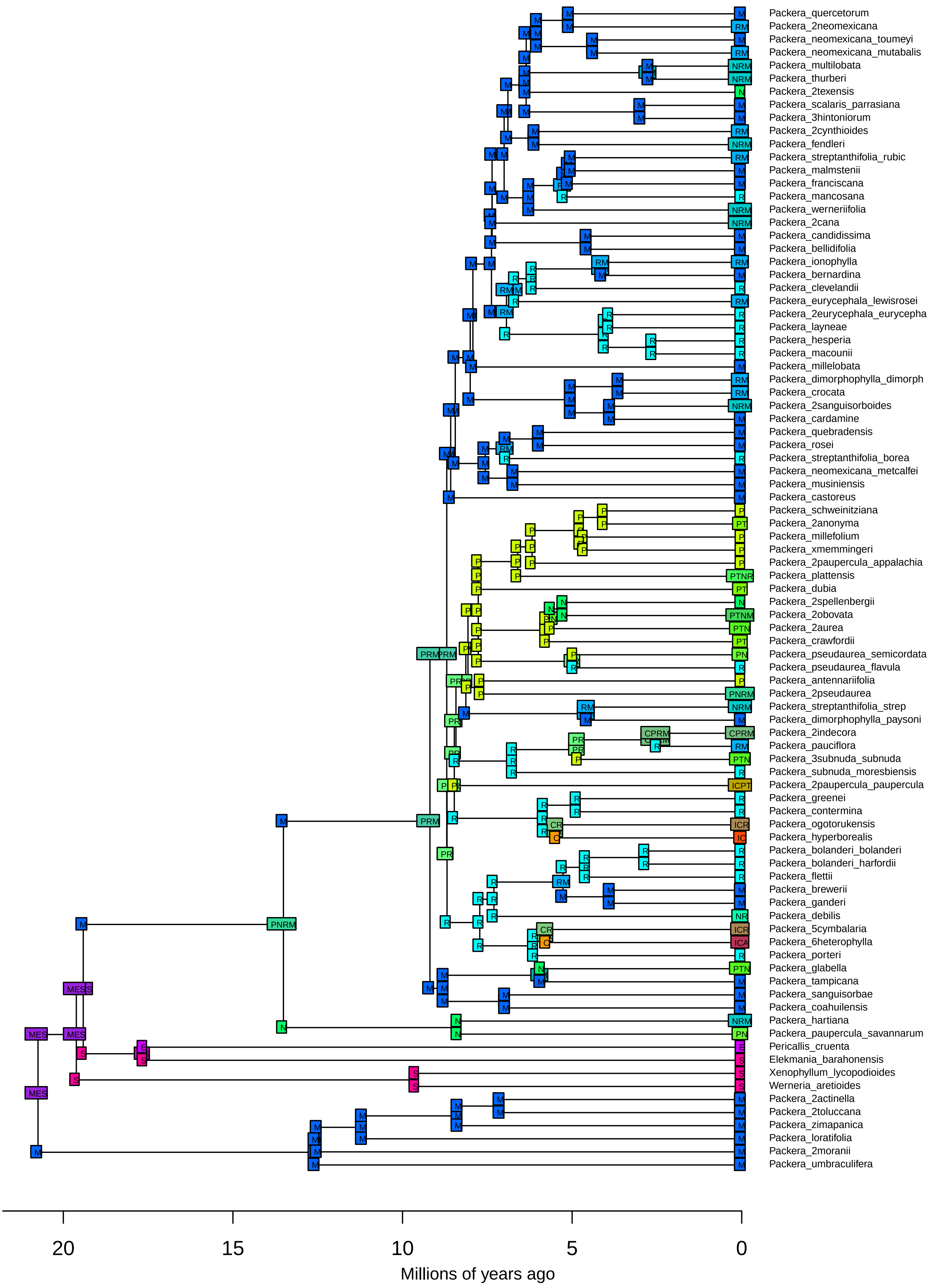

**BioGeoBEARS DEC on Packera − HybPiper**
**ancstates: global optim, 4 areas max. d=0.0101; e=0.0038; j=0; LnL=−300.65**

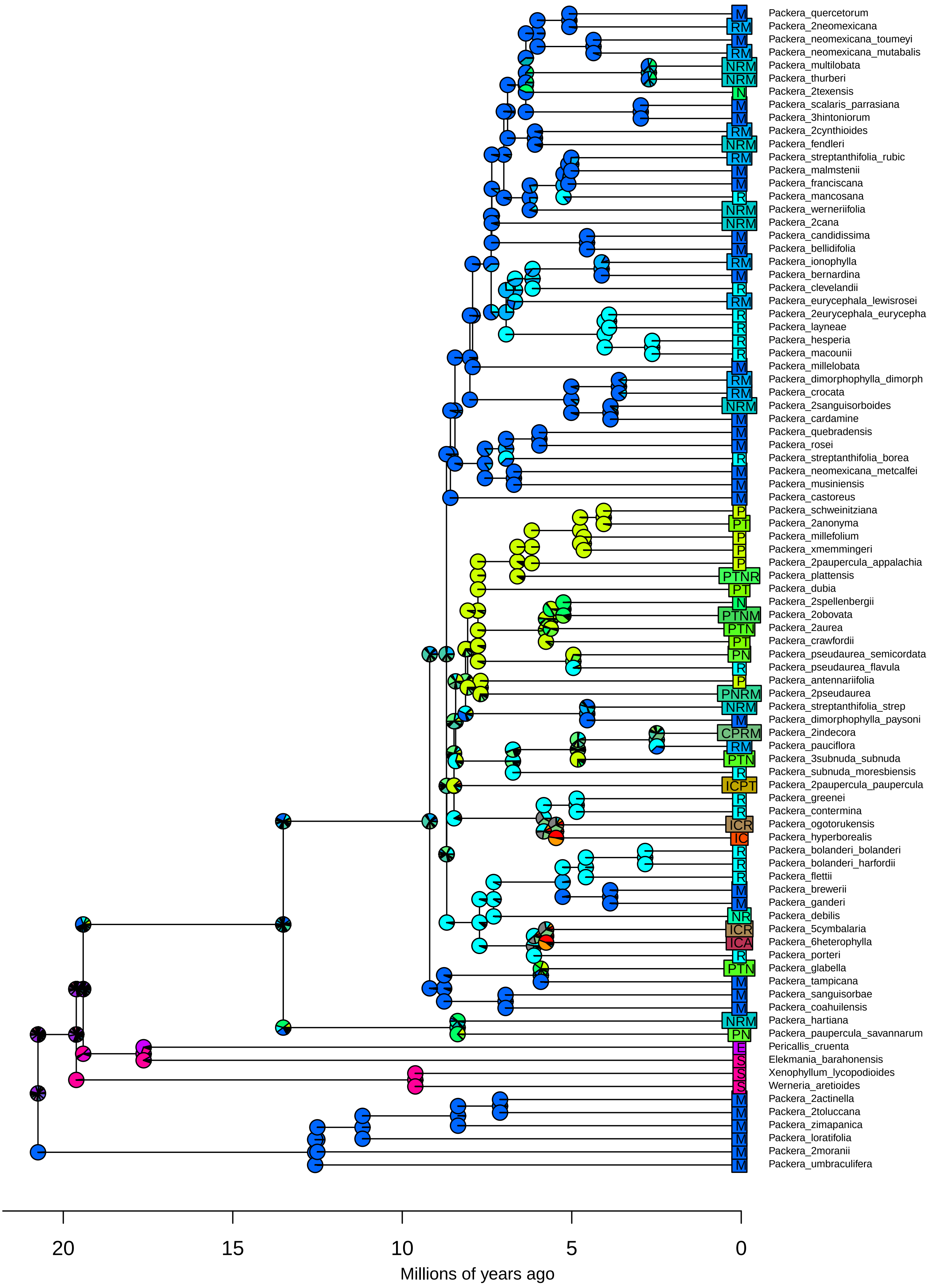

**BioGeoBEARS DEC+J on Packera − HybPiper**
**ancstates: global optim, 4 areas max. d=0.0091; e=0; j=0.0093; LnL=−293.05**

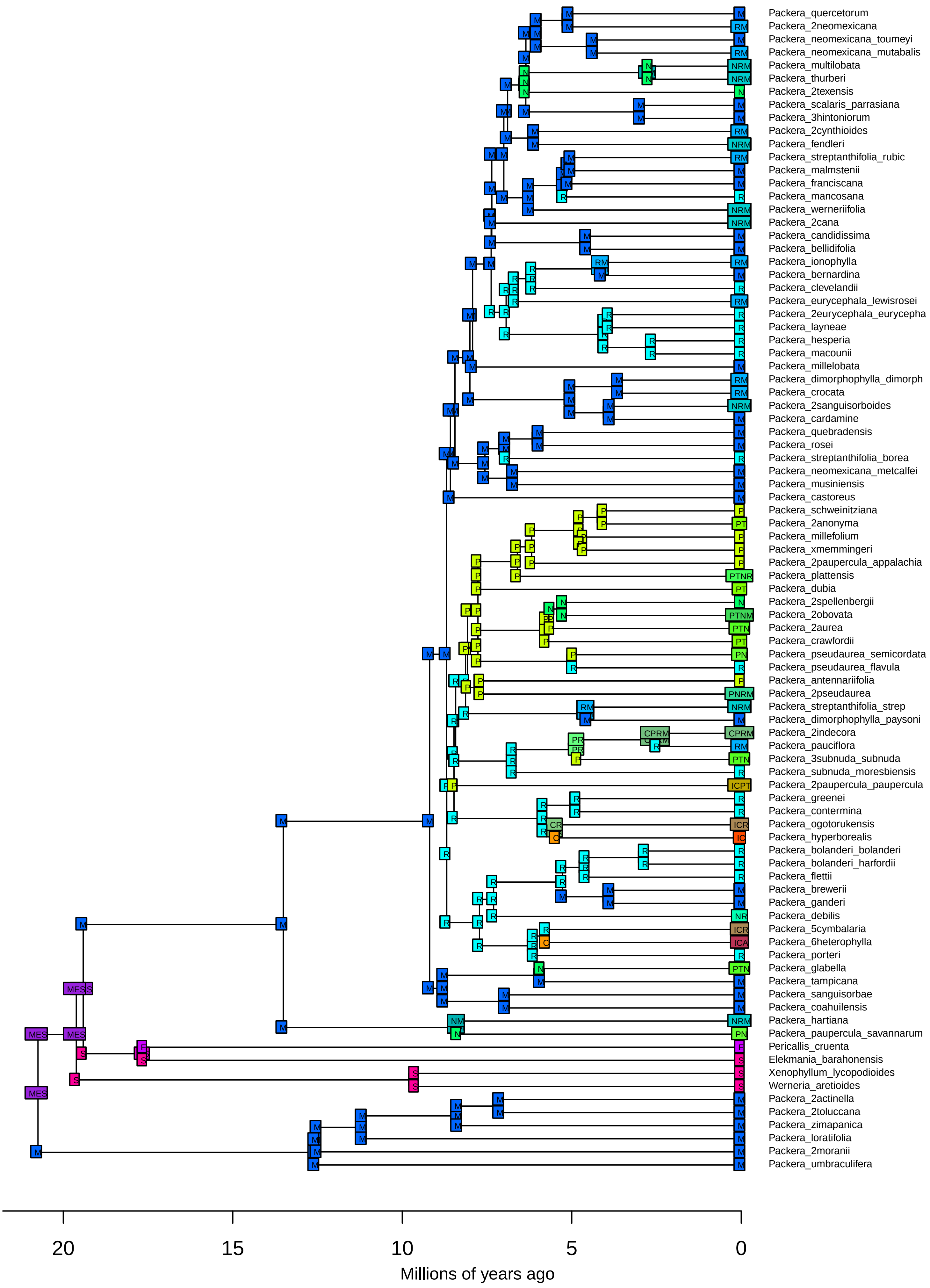

**BioGeoBEARS DEC+J on Packera − HybPiper**
**ancstates: global optim, 4 areas max. d=0.0091; e=0; j=0.0093; LnL=−293.05**

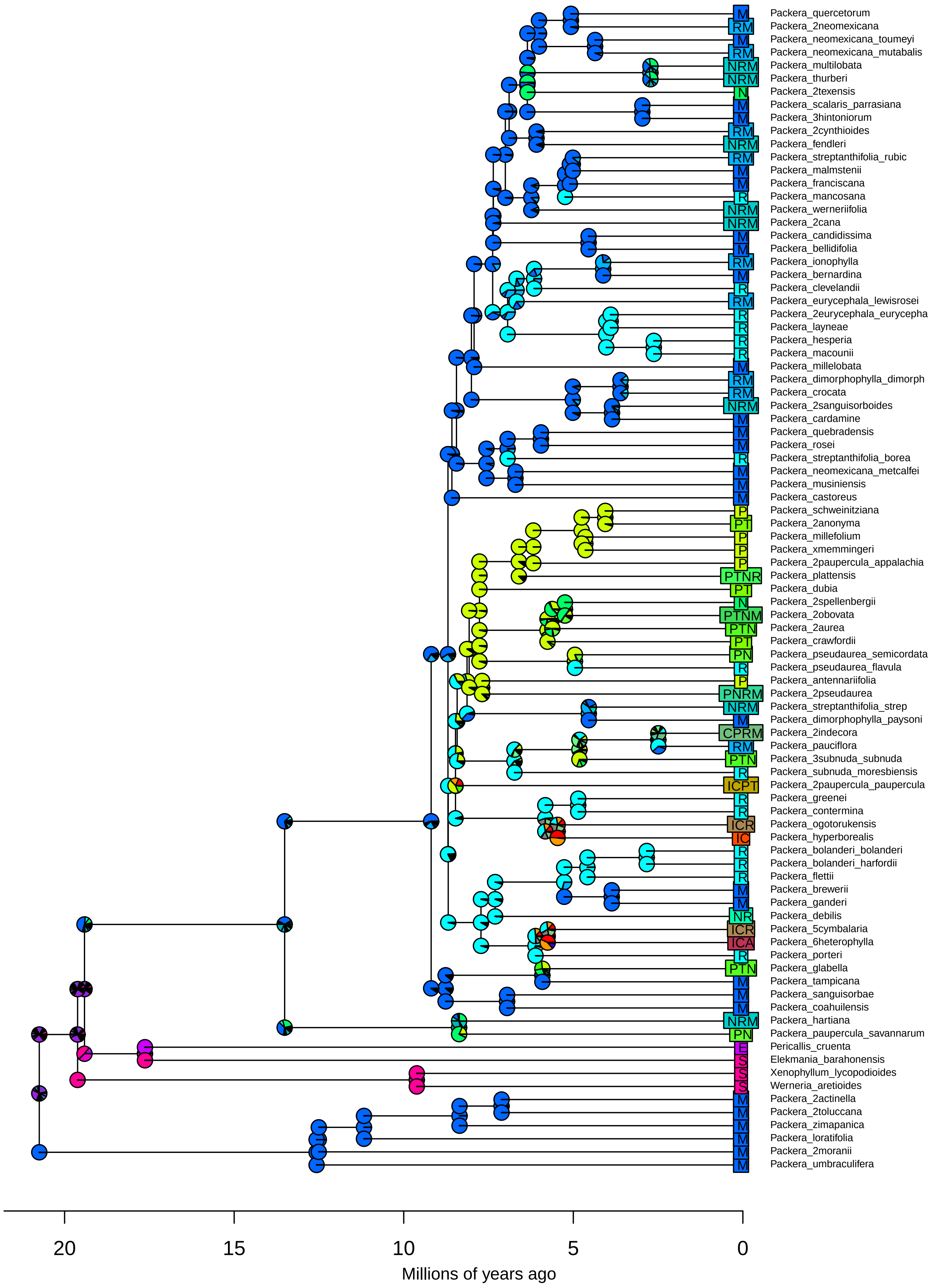

**BioGeoBEARS DIVALIKE on Packera − HybPiper**
**ancstates: global optim, 4 areas max. d=0.0119; e=0.0073; j=0; LnL=−309.94**

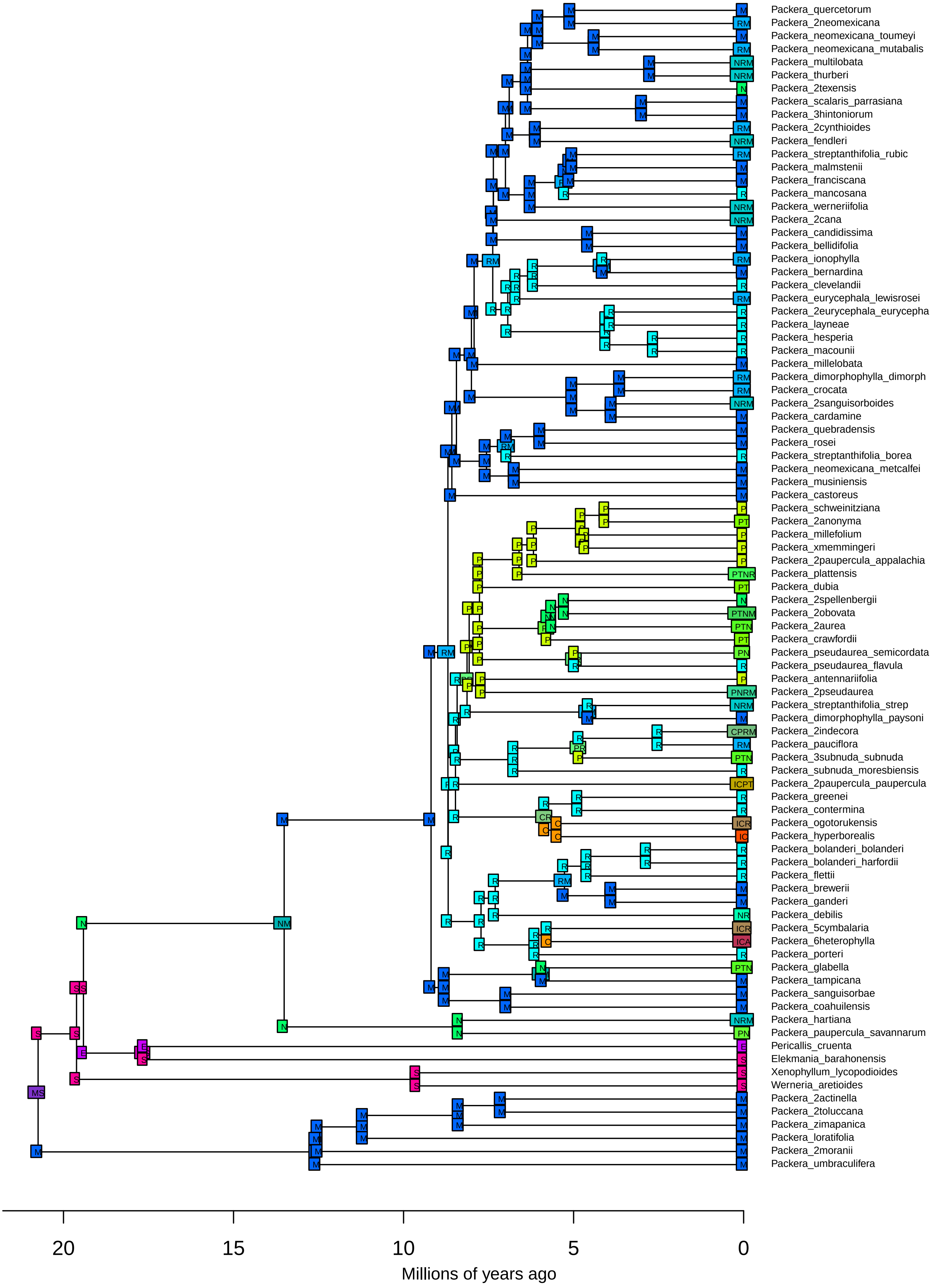

**BioGeoBEARS DIVALIKE on Packera − HybPiper**
**ancstates: global optim, 4 areas max. d=0.0119; e=0.0073; j=0; LnL=−309.94**

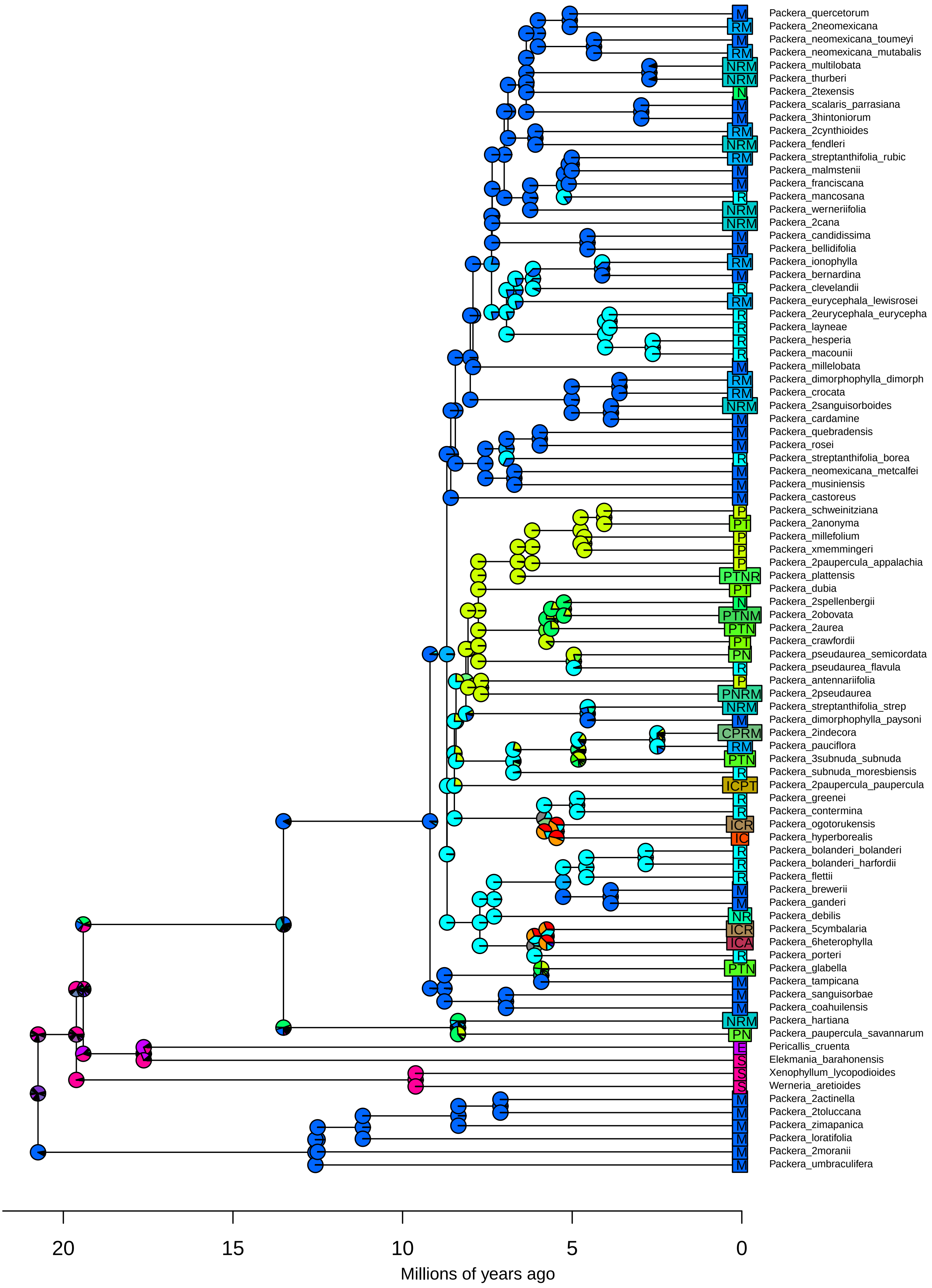

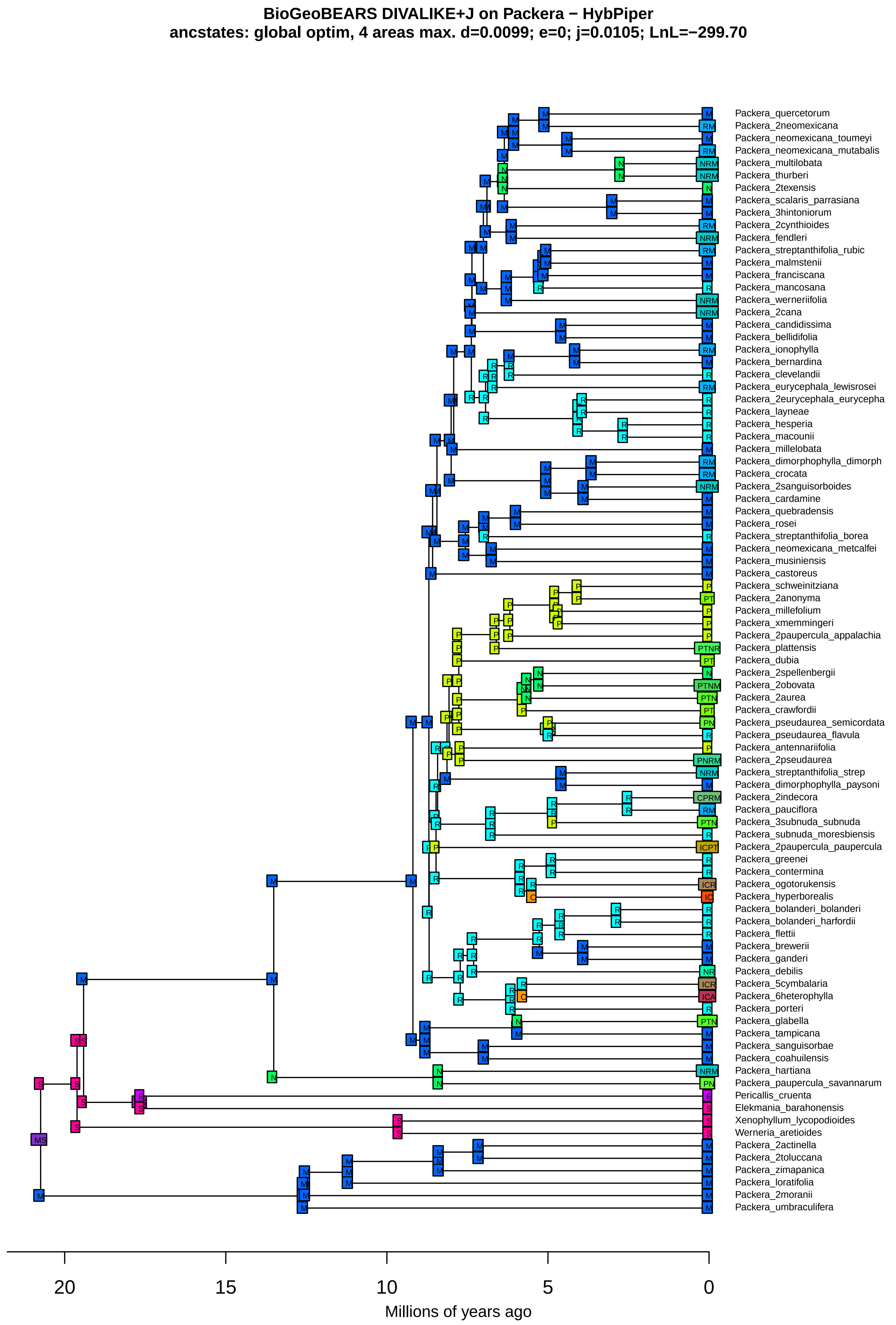

BioGeoBEARS DIVALIKE+J on Packera − HybPiper
ancstates: global optim, 4 areas max. d=0.0099; e=0; j=0.0105; LnL=−299.70
Packera_quercetorum
Packera_2neomexicana
Packera_neomexicana_toumeyi
Packera_neomexicana_mutabalis
Packera_multilobata
Packera_thurberi
Packera_2texensis
Packera_scalaris_parrasiana
Packera_3hintoniorum
Packera_2cynthioides
Packera_fendleri
Packera_streptanthifolia_rubic
Packera_malmstenii
Packera_franciscana
Packera_mancosana
Packera_werneriifolia
Packera_2cana
Packera_candidissima
Packera_bellidifolia
Packera_ionophylla
Packera_bernardina
Packera_clevelandii
Packera_eurycephala_lewisrosei
Packera_2eurycephala_eurycepha
Packera_layneae
Packera_hesperia
Packera_macounii
Packera_millelobata
Packera_dimorphophylla_dimorph
Packera_crocata
Packera_2sanguisorboides
Packera_cardamine
Packera_quebradensis
Packera_rosei
Packera_streptanthifolia_borea
Packera_neomexicana_metcalfei
Packera_musiniensis
Packera_castoreus
Packera_schweinitziana
Packera_2anonyma
Packera_millefolium
Packera_xmemmingeri
Packera_2paupercula_appalachia
Packera_plattensis
Packera_dubia
Packera_2spellenbergii
Packera_2obovata
Packera_2aurea
Packera_crawfordii
Packera_pseudaurea_semicordata
Packera_pseudaurea_flavula
Packera_antennariifolia
Packera_2pseudaurea
Packera_streptanthifolia_strep
Packera_dimorphophylla_paysoni
Packera_2indecora
Packera_pauciflora
Packera_3subnuda_subnuda
Packera_subnuda_moresbiensis
Packera_2paupercula_paupercula
Packera_greenei
Packera_contermina
Packera_ogotorukensis
Packera_hyperborealis
Packera_bolanderi_bolanderi
Packera_bolanderi_harfordii
Packera_flettii
Packera_breweri
Packera_ganderi
Packera_debilis
Packera_5cymbalaria
Packera_6heterophylla
Packera_porteri
Packera_glabella
Packera_tampicana
Packera_sanguisorbae
Packera_coahuilensis
Packera_hartiana
Packera_paupercula_savannarum
Pericallis_cruenta
Elekmania_barahonensis
Xenophyllum_lycopodioides
Werneria_aretioides
Packera_2actinella
Packera_2toluccana
Packera_zimapanica
Packera_loratifolia
Packera_2moranii
Packera_umbraculifera
20
15
10
5
0
Millions of years ago

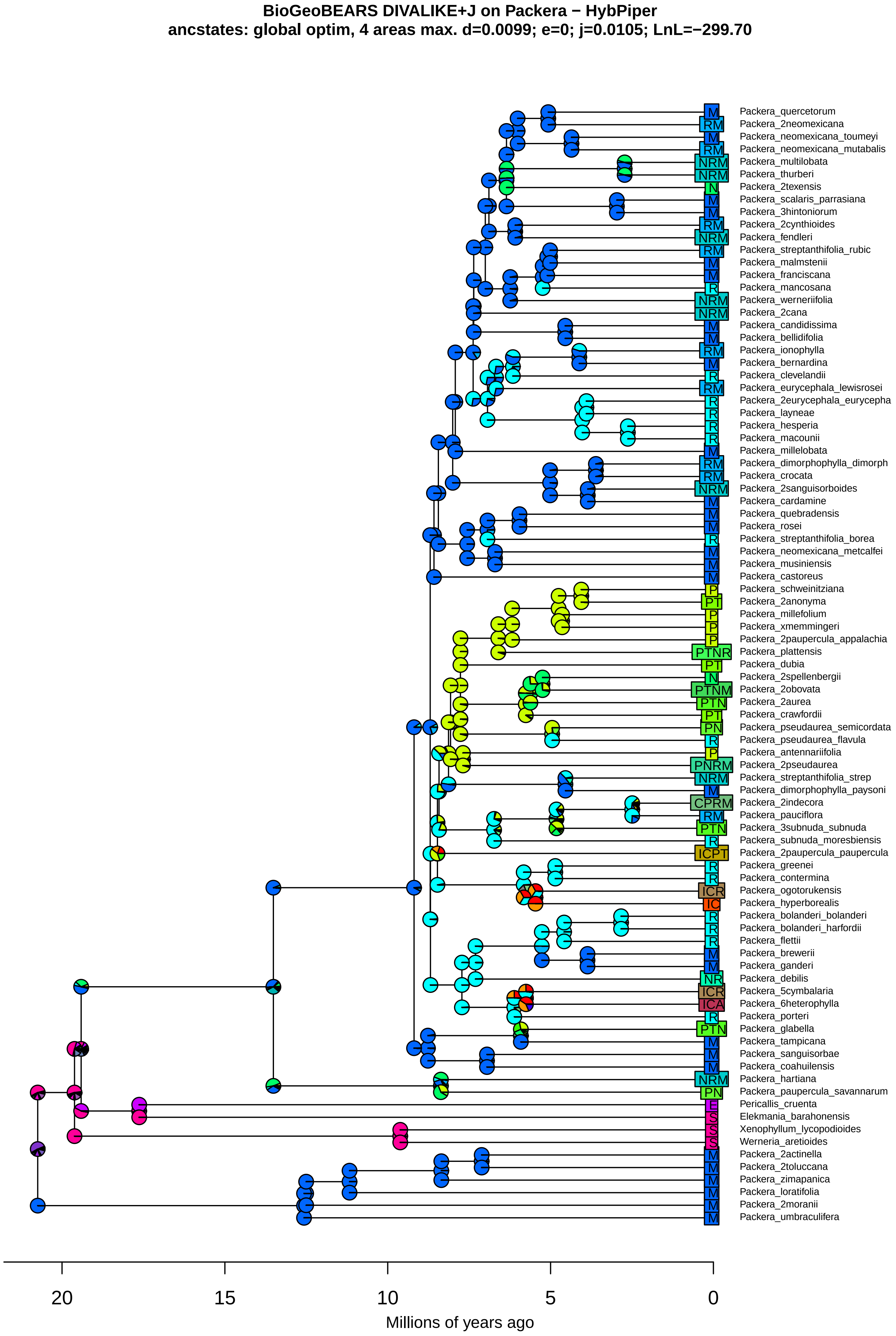
BioGeoBEARS DIVALIKE+J on Packera − HybPiper
ancstates: global optim, 4 areas max. d=0.0099; e=0; j=0.0105; LnL=−299.70
Packera_quercetorum
Packera_2neomexicana
Packera_neomexicana_toumeyi
Packera_neomexicana_mutabalis
Packera_multilobata
Packera_thurberi
Packera_2texensis
Packera_scalaris_parrasiana
Packera_3hintoniorum
Packera_2cynthioides
Packera_fendleri
Packera_streptanthifolia_rubic
Packera_malmstenii
Packera_franciscana
Packera_mancosana
Packera_werneriifolia
Packera_2cana
Packera_candidissima
Packera_bellidifolia
Packera_ionophylla
Packera_bernardina
Packera_clevelandii
Packera_eurycephala_lewisrosei
Packera_2eurycephala_eurycepha
Packera_layneae
Packera_hesperia
Packera_macounii
Packera_millelobata
Packera_dimorphophylla_dimorph
Packera_crocata
Packera_2sanguisorboides
Packera_cardamine
Packera_quebradensis
Packera_rosei
Packera_streptanthifolia_borea
Packera_neomexicana_metcalfei
Packera_musiniensis
Packera_castoreus
Packera_schweinitziana
Packera_2anonyma
Packera_millefolium
Packera_xmemmingeri
Packera_2paupercula_appalachia
Packera_plattensis
Packera_dubia
Packera_2spellenbergii
Packera_2obovata
Packera_2aurea
Packera_crawfordii
Packera_pseudaurea_semicordata
Packera_pseudaurea_flavula
Packera_antennariifolia
Packera_2pseudaurea
Packera_streptanthifolia_strep
Packera_dimorphophylla_paysoni
Packera_2indecora
Packera_pauciflora
Packera_3subnuda_subnuda
Packera_subnuda_moresbiensis
Packera_2paupercula_paupercula
Packera_greenei
Packera_contermina
Packera_ogotorukensis
Packera_hyperborealis
Packera_bolanderi_bolanderi
Packera_bolanderi_harfordii
Packera_flettii
Packera_brewerii
Packera_ganderi
Packera_debilis
Packera_5cymbalaria
Packera_6heterophylla
Packera_porteri
Packera_glabella
Packera_tampicana
Packera_sanguisorbae
Packera_coahuilensis
Packera_hartiana
Packera_paupercula_savannarum
Pericallis_cruenta
Elekmania_barahonensis
Xenophyllum_lycopodioides
Werneria_aretioides
Packera_2actinella
Packera_2toluccana
Packera_zimapanica
Packera_loratifolia
Packera_2moranii
Packera_umbraculifera
20
15
10
5
0
Millions of years ago

**BioGeoBEARS BAYAREALIKE on Packera − HybPiper**
**ancstates: global optim, 4 areas max. d=0.0035; e=0.086; j=0; LnL=−275.79**

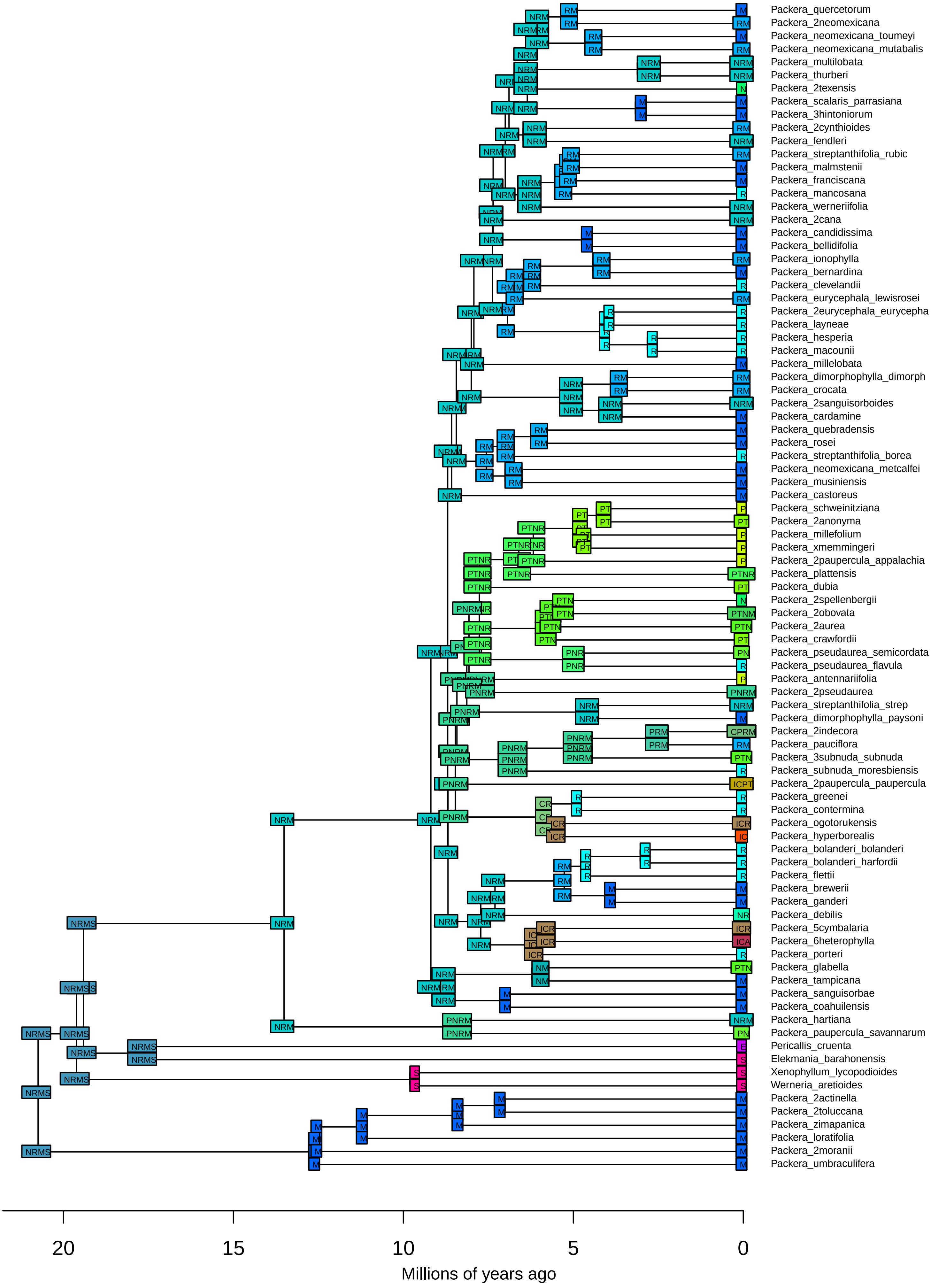

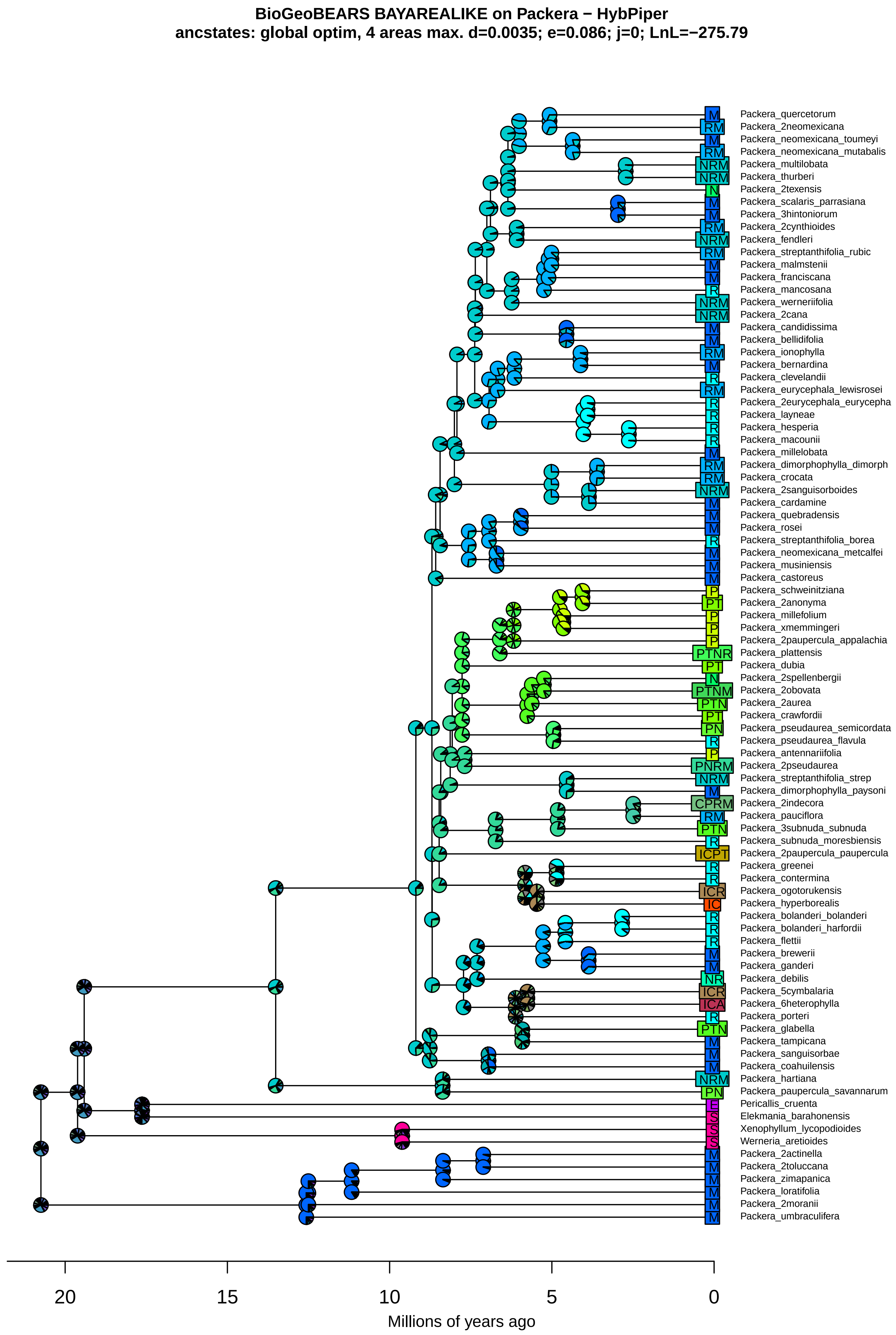

BioGeoBEARS BAYAREALIKE on Packera − HybPiper
ancstates: global optim, 4 areas max. d=0.0035; e=0.086; j=0; LnL=−275.79
M Packera_quercetorum
RM Packera_2neomexicana
M Packera_neomexicana_toumeyi
RM Packera_neomexicana_mutabalis
NRM Packera_multilobata
NRM Packera_thurberi
N Packera_2texensis
M Packera_scalaris_parrasiana
M Packera_3hintoniorum
RM Packera_2cynthioides
NRM Packera_fendleri
RM Packera_streptanthifolia_rubic
M Packera_malmstenii
M Packera_franciscana
R Packera_mancosana
NRM Packera_werneriifolia
NRM Packera_2cana
M Packera_candidissima
M Packera_bellidifolia
RM Packera_ionophylla
M Packera_bernardina
R Packera_clevelandii
RM Packera_eurycephala_lewisrosei
R Packera_2eurycephala_eurycepha
R Packera_layneae
R Packera_hesperia
R Packera_macounii
M Packera_millelobata
RM Packera_dimorphophylla_dimorph
RM Packera_crocata
NRM Packera_2sanguisorboides
M Packera_cardamine
M Packera_quebradensis
M Packera_rosei
R Packera_streptanthifolia_borea
M Packera_neomexicana_metcalfei
M Packera_musiniensis
M Packera_castoreus
P Packera_schweinitziana
PT Packera_2anonyma
P Packera_millefolium
P Packera_xmemmingeri
P Packera_2paupercula_appalachia
PTNR Packera_plattensis
PT Packera_dubia
N Packera_2spellenbergii
PTNM Packera_2obovata
PTN Packera_2aurea
PT Packera_crawfordii
PN Packera_pseudaurea_semicordata
R Packera_pseudaurea_flavula
P Packera_antennariifolia
PNRM Packera_2pseudaurea
NRM Packera_streptanthifolia_strep
M Packera_dimorphophylla_paysoni
CPRM Packera_2indecora
RM Packera_pauciflora
PTN Packera_3subnuda_subnuda
R Packera_subnuda_moresbiensis
ICPT Packera_2paupercula_paupercula
R Packera_greenei
R Packera_contermina
ICR Packera_ogotorukensis
IC Packera_hyperborealis
R Packera_bolanderi_bolanderi
R Packera_bolanderi_harfordii
R Packera_flettii
M Packera_brewerii
M Packera_ganderi
NR Packera_debilis
ICR Packera_5cymbalaria
ICA Packera_6heterophylla
R Packera_porteri
PTN Packera_glabella
M Packera_tampicana
M Packera_sanguisorbae
M Packera_coahuilensis
NRM Packera_hartiana
PN Packera_paupercula_savannarum
E Pericallis_cruenta
S Elekmania_barahonensis
S Xenophyllum_lycopodioides
S Werneria_aretioides
M Packera_2actinella
M Packera_2toluccana
M Packera_zimapanica
M Packera_loratifolia
M Packera_2moranii
M Packera_umbraculifera
20
15
10
5
0
Millions of years ago

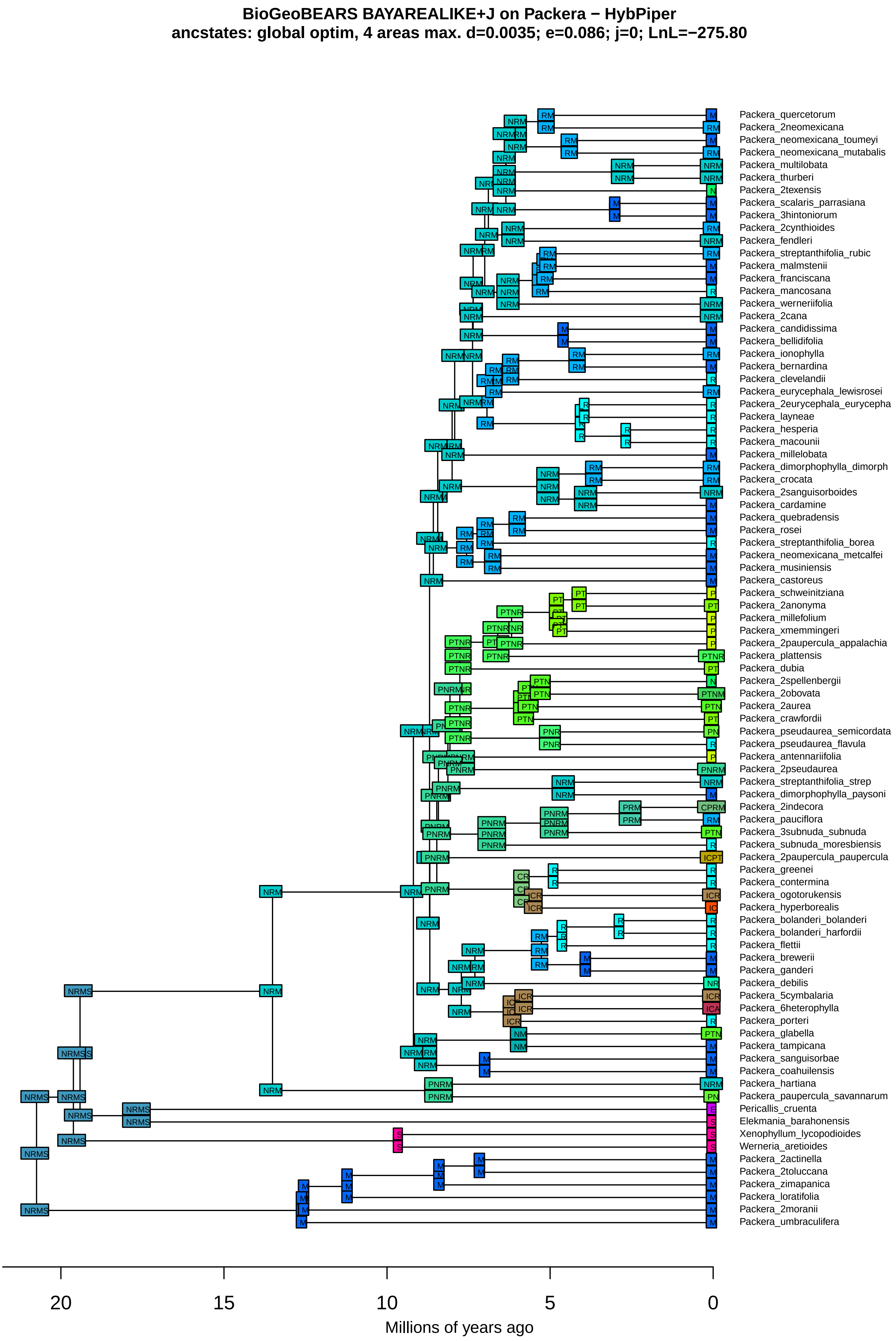

BioGeoBEARS BAYAREALIKE+J on Packera − HybPiper
ancstates: global optim, 4 areas max. d=0.0035; e=0.086; j=0; LnL=−275.80
Packera_quercetorum
Packera_2neomexicana
Packera_neomexicana_toumeyi
Packera_neomexicana_mutabalis
Packera_multilobata
Packera_thurberi
Packera_2texensis
Packera_scalaris_parrasiana
Packera_3hintoniorum
Packera_2cynthioides
Packera_fendleri
Packera_streptanthifolia_rubic
Packera_malmstenii
Packera_franciscana
Packera_mancosana
Packera_werneriifolia
Packera_2cana
Packera_candidissima
Packera_bellidifolia
Packera_ionophylla
Packera_bernardina
Packera_clevelandii
Packera_eurycephala_lewisrosei
Packera_2eurycephala_eurycepha
Packera_layneae
Packera_hesperia
Packera_macounii
Packera_millelobata
Packera_dimorphophylla_dimorph
Packera_crocata
Packera_2sanguisorboides
Packera_cardamine
Packera_quebradensis
Packera_rosei
Packera_streptanthifolia_borea
Packera_neomexicana_metcalfei
Packera_musiniensis
Packera_castoreus
Packera_schweinitziana
Packera_2anonyma
Packera_millefolium
Packera_xmemmingeri
Packera_2paupercula_appalachia
Packera_plattensis
Packera_dubia
Packera_2spellenbergii
Packera_2obovata
Packera_2aurea
Packera_crawfordii
Packera_pseudaurea_semicordata
Packera_pseudaurea_flavula
Packera_antennariifolia
Packera_2pseudaurea
Packera_streptanthifolia_strep
Packera_dimorphophylla_paysoni
Packera_2indecora
Packera_pauciflora
Packera_3subnuda_subnuda
Packera_subnuda_moresbiensis
Packera_2paupercula_paupercula
Packera_greenei
Packera_contermina
Packera_ogotorukensis
Packera_hyperborealis
Packera_bolanderi_bolanderi
Packera_bolanderi_harfordii
Packera_flettii
Packera_breweri
Packera_ganderi
Packera_debilis
Packera_5cymbalaria
Packera_6heterophylla
Packera_porteri
Packera_glabella
Packera_tampicana
Packera_sanguisorbae
Packera_coahuilensis
Packera_hartiana
Packera_paupercula_savannarum
Pericallis_cruenta
Elekmania_barahonensis
Xenophyllum_lycopodioides
Werneria_aretioides
Packera_2actinella
Packera_2toluccana
Packera_zimapanica
Packera_loratifolia
Packera_2moranii
Packera_umbraculifera
20
15
10
5
0
Millions of years ago

**BioGeoBEARS BAYAREALIKE+J on Packera − HybPiper**
**ancstates: global optim, 4 areas max. d=0.0035; e=0.086; j=0; LnL=−275.80**

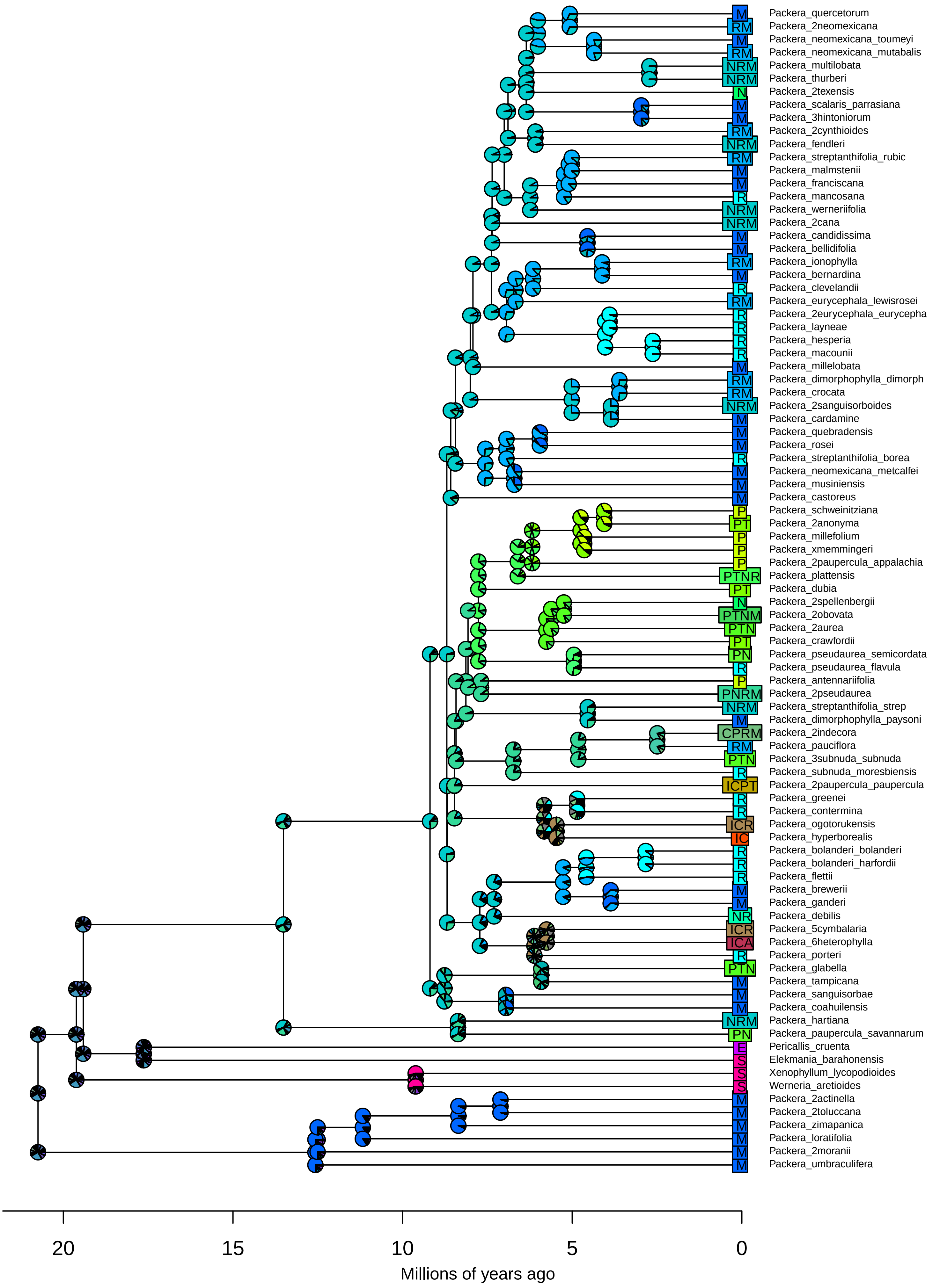

Legend

| | | | | | | | | | | | | | | | | |
|---|---|---|---|---|---|---|---|---|---|---|---|---|---|---|---|---|
| P | TS | ICM | INE | CTR | PTN | PES | NMA | ICPA | ICMS | IPME | INRM | CPTN | CPES | CNMA | PTNS | PNES | [illegible] |
| T | NR | ICA | INS | CTM | PTR | TNR | NME | ICPE | ICAE | IPMS | INRA | CPTR | CTNR | CNME | PTRM | PRMA | [illegible] |
| N | NM | ICE | IRM | CTA | PTM | TNM | NMS | ICPS | ICAS | IPAE | INRE | CPTM | CTNM | CNMS | PTRA | PRME | [illegible] |
| R | NA | ICS | IRA | CTE | PTA | TNA | NAE | ICTN | ICES | IPAS | INRS | CPTA | CTNA | CNAE | PTRE | PRMS | [illegible] |
| M | NE | IPT | IRE | CTS | PTE | TNE | NAS | ICTR | IPTN | IPES | INMA | CPTE | CTNE | CNAS | PTRS | PRAE | [illegible] |
| A | NS | IPN | IRS | CNR | PTS | TNS | NES | ICTM | IPTR | ITNR | INME | CPTS | CTNS | CNES | PTMA | PRAS | [illegible] |
| E | RM | IPR | IMA | CNM | PNR | TRM | RMA | ICTA | IPTM | ITNM | INMS | CPNR | CTRM | CRMA | PTME | PRES | [illegible] |
| S | RA | IPM | IME | CNA | PNM | TRA | RME | ICTE | IPTA | ITNA | INAE | CPNM | CTRA | CRME | PTMS | PMAE | [illegible] |
| T | RE | IPA | IMS | CNE | PNA | TRE | RMS | ICTS | IPTE | ITNE | INAS | CPNA | CTRE | CRMS | PTAE | PMAS | [illegible] |
| N | RS | IPE | IAE | CNS | PNE | TRS | RAE | ICNR | IPTS | ITNS | INES | CPNE | CTRS | CRAE | PTAS | PMES | [illegible] |
| R | MA | IPS | IAS | CRM | PNS | TMA | RAS | ICNM | IPNR | ITRM | IRMA | CPNS | CTMA | CRAS | PTES | PAES | [illegible] |
| M | ME | ITN | IES | CRA | PRM | TME | RES | ICNA | IPNM | ITRA | IRME | CPRM | CTME | CRES | PNRM | TNRM | [illegible] |
| A | MS | ITR | CPT | CRE | PRA | TMS | MAE | ICNE | IPNA | ITRE | IRMS | CPRA | CTMS | CMAE | PNRA | TNRA | [illegible] |
| E | AE | ITM | CPN | CRS | PRE | TAE | MAS | ICNS | IPNE | ITRS | IRAE | CPRE | CTAE | CMAS | PNRE | TNRE | [illegible] |
| S | AS | ITA | CPR | CMA | PRS | TAS | MES | ICRM | IPNS | ITMA | IRAS | CPRS | CTAS | CMES | PNRS | TNRS | [illegible] |
| N | ES | ITE | CPM | CME | PMA | TES | AES | ICRA | IPRM | ITME | IRES | CPMA | CTES | CAES | PNMA | TNMA | [illegible] |
| R | ICP | ITS | CPA | CMS | PME | NRM | ICPT | ICRE | IPRA | ITMS | IMAE | CPME | CNRM | PTNR | PNME | TNME | [illegible] |
| M | ICT | INR | CPE | CAE | PMS | NRA | ICPN | ICRS | IPRE | ITAE | IMAS | CPMS | CNRA | PTNM | PNMS | TNMS | [illegible] |
| A | ICN | INM | CPS | CAS | PAE | NRE | ICPR | ICMA | IPRS | ITAS | IMES | CPAE | CNRE | PTNA | PNAE | TNAE | [illegible] |
| E | ICR | INA | CTN | CES | PAS | NRS | ICPM | ICME | IPMA | ITES | IAES | CPAS | CNRS | PTNE | PNAS | TNAS | [illegible] |

**BioGeoBEARS DIVALIKE on Packera − paragone−nf:MO**
**ancstates: global optim, 4 areas max. d=0.0073; e=0.0085; j=0; LnL=−328.43**

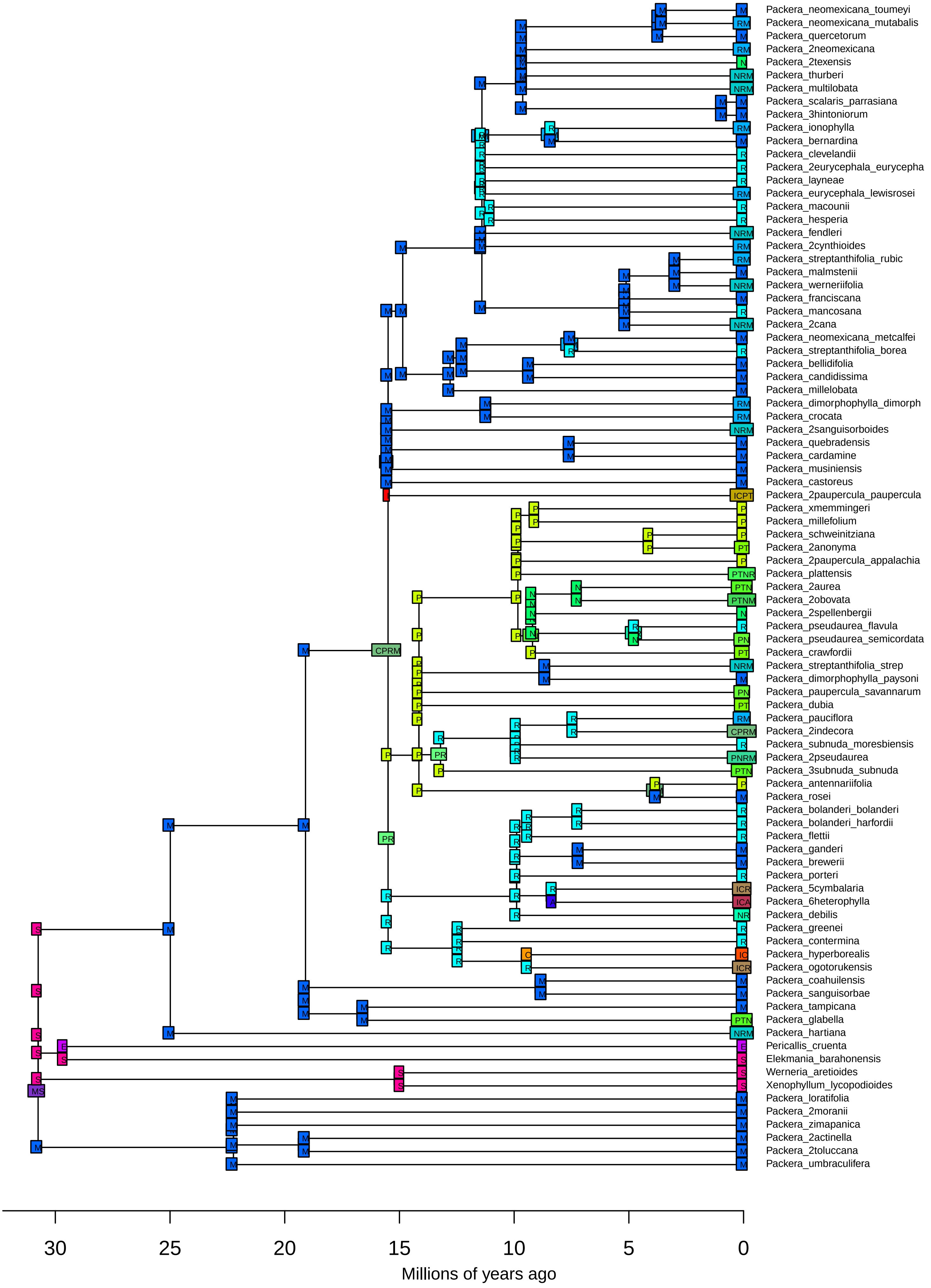

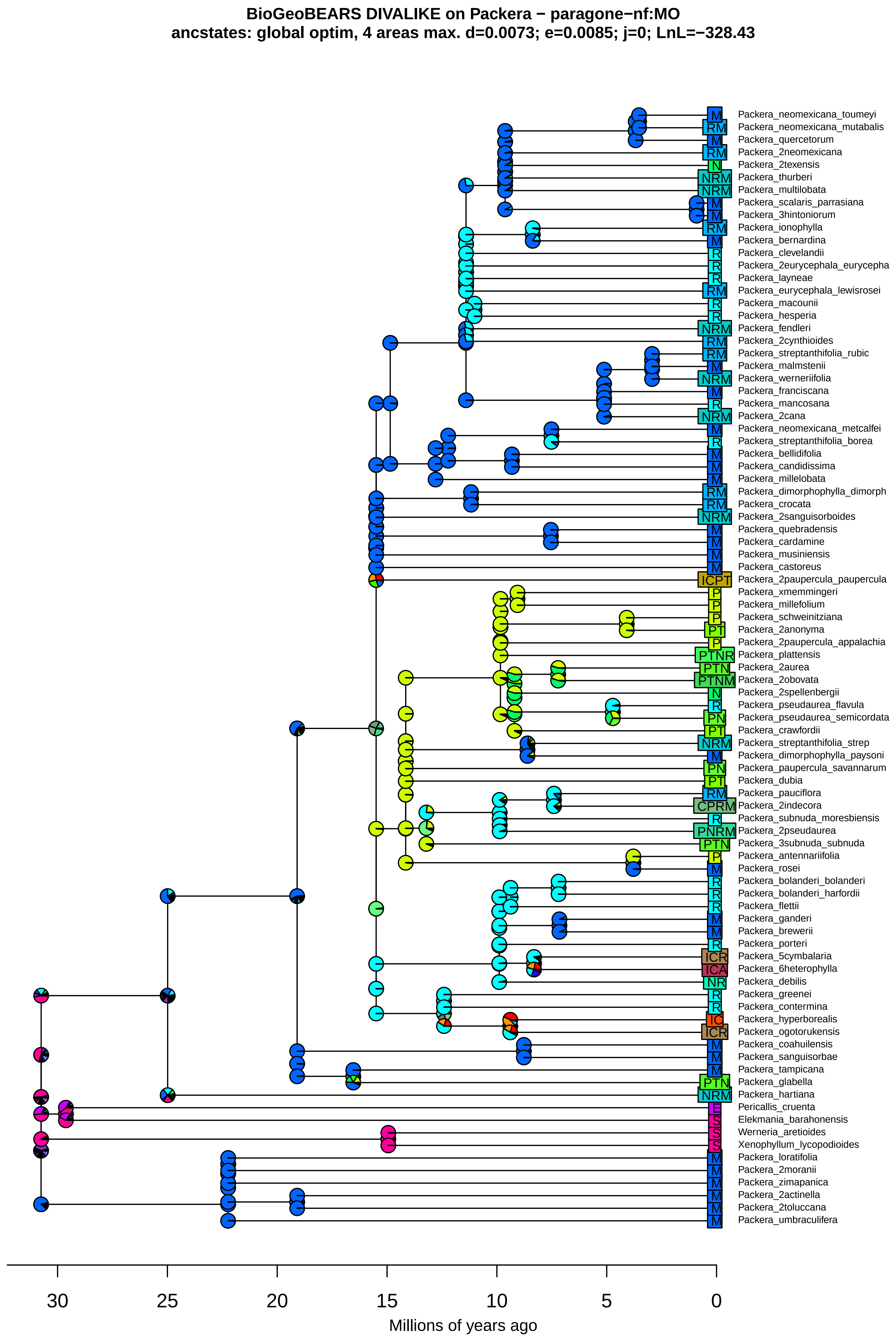
BioGeoBEARS DIVALIKE on Packera − paragone−nf:MO
anctstates: global optim, 4 areas max. d=0.0073; e=0.0085; j=0; LnL=−328.43
Packera_neomexicana_toumeyi
Packera_neomexicana_mutabalis
Packera_quercetorum
Packera_2neomexicana
Packera_2texensis
Packera_thurberi
Packera_multilobata
Packera_scalaris_parrasiana
Packera_3hintoniorum
Packera_ionophylla
Packera_bernardina
Packera_clevelandii
Packera_2eurycephala_eurycepha
Packera_layneae
Packera_eurycephala_lewisrosei
Packera_macounii
Packera_hesperia
Packera_fendleri
Packera_2cynthioides
Packera_streptanthifolia_rubic
Packera_malmstenii
Packera_werneriifolia
Packera_franciscana
Packera_mancosana
Packera_2cana
Packera_neomexicana_metcalfei
Packera_streptanthifolia_borea
Packera_bellidifolia
Packera_candidissima
Packera_millelobata
Packera_dimorphophylla_dimorph
Packera_crocata
Packera_2sanguisorboides
Packera_quebradensis
Packera_cardamine
Packera_musiniensis
Packera_castoreus
Packera_2paupercula_paupercula
Packera_xmemmingeri
Packera_millefolium
Packera_schweinitziana
Packera_2anonyma
Packera_2paupercula_appalachia
Packera_plattensis
Packera_2aurea
Packera_2obovata
Packera_2spellenbergii
Packera_pseudaurea_flavula
Packera_pseudaurea_semicordata
Packera_crawfordii
Packera_streptanthifolia_strep
Packera_dimorphophylla_paysoni
Packera_paupercula_savannarum
Packera_dubia
Packera_pauciflora
Packera_2indecora
Packera_subnuda_moresbiensis
Packera_2pseudaurea
Packera_3subnuda_subnuda
Packera_antennariifolia
Packera_rosei
Packera_bolanderi_bolanderi
Packera_bolanderi_harfordii
Packera_flettii
Packera_ganderi
Packera_brewerii
Packera_porteri
Packera_5cymbalaria
Packera_6heterophylla
Packera_debilis
Packera_greenei
Packera_contermina
Packera_hyperborealis
Packera_ogotorukensis
Packera_coahuilensis
Packera_sanguisorbae
Packera_tampicana
Packera_glabella
Packera_hartiana
Pericallis_cruenta
Elekmania_barahonensis
Werneria_aretioides
Xenophyllum_lycopodioides
Packera_loratifolia
Packera_2moranii
Packera_zimapanica
Packera_2actinella
Packera_2toluccana
Packera_umbraculifera
30 25 20 15 10 5 0
Millions of years ago

**BioGeoBEARS DIVALIKE+J on Packera − paragone−nf:MO**
**ancstates: global optim, 4 areas max. d=0.0056; e=0; j=0.0128; LnL=−304.16**

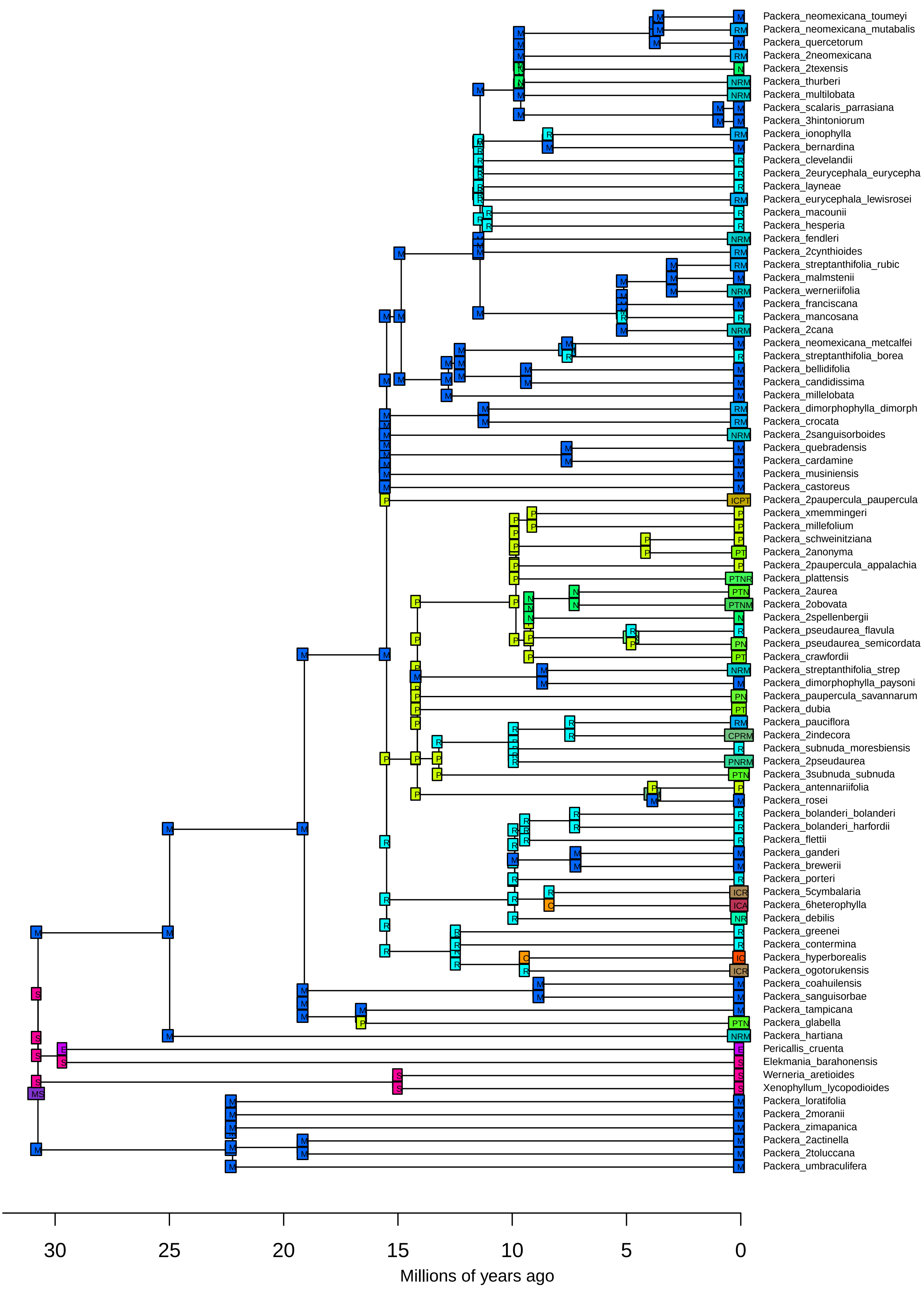

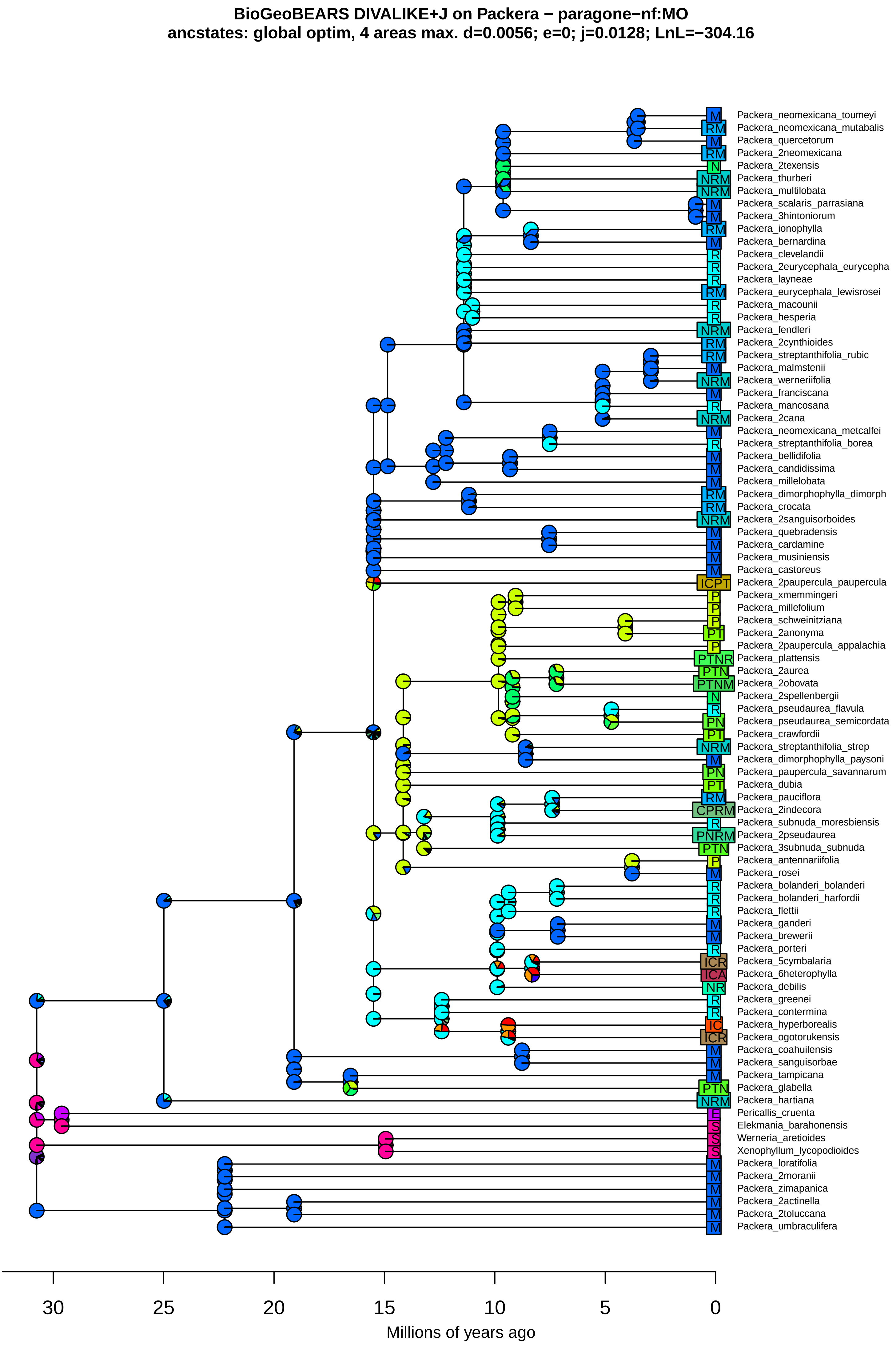
BioGeoBEARS DIVALIKE+J on Packera − paragone−nf:MO
ancstates: global optim, 4 areas max. d=0.0056; e=0; j=0.0128; LnL=−304.16
M Packera_neomexicana_toumeyi
RM Packera_neomexicana_mutabalis
M Packera_quercetorum
RM Packera_2neomexicana
N Packera_2texensis
NRM Packera_thurberi
NRM Packera_multilobata
M Packera_scalaris_parrasiana
M Packera_3hintoniorum
RM Packera_ionophylla
M Packera_bernardina
R Packera_clevelandii
R Packera_2eurycephala_eurycepha
R Packera_layneae
RM Packera_eurycephala_lewisrosei
R Packera_macounii
R Packera_hesperia
NRM Packera_fendleri
RM Packera_2cynthioides
RM Packera_streptanthifolia_rubic
M Packera_malmstenii
NRM Packera_werneriifolia
M Packera_franciscana
R Packera_mancosana
NRM Packera_2cana
M Packera_neomexicana_metcalfei
R Packera_streptanthifolia_borea
M Packera_bellidifolia
M Packera_candidissima
M Packera_millelobata
RM Packera_dimorphophylla_dimorph
RM Packera_crocata
NRM Packera_2sanguisorboides
M Packera_quebradensis
M Packera_cardamine
M Packera_musiniensis
M Packera_castoreus
ICPT Packera_2paupercula_paupercula
P Packera_xmemmingeri
P Packera_millefolium
P Packera_schweinitziana
PT Packera_2anonyma
P Packera_2paupercula_appalachia
PTNR Packera_plattensis
PTN Packera_2aurea
PTNM Packera_2obovata
N Packera_2spellenbergii
R Packera_pseudaurea_flavula
PN Packera_pseudaurea_semicordata
PT Packera_crawfordii
NRM Packera_streptanthifolia_strep
M Packera_dimorphophylla_paysoni
PN Packera_paupercula_savannarum
PT Packera_dubia
RM Packera_pauciflora
CPRM Packera_2indecora
R Packera_subnuda_moresbiensis
PNRM Packera_2pseudaurea
PTN Packera_3subnuda_subnuda
P Packera_antennariifolia
M Packera_rosei
R Packera_bolanderi_bolanderi
R Packera_bolanderi_harfordii
R Packera_flettii
M Packera_ganderi
M Packera_brewerii
R Packera_porteri
ICR Packera_5cymbalaria
ICA Packera_6heterophylla
NR Packera_debilis
R Packera_greenei
R Packera_contermina
IC Packera_hyperborealis
ICR Packera_ogotorukensis
M Packera_coahuilensis
M Packera_sanguisorbae
M Packera_tampicana
PTN Packera_glabella
NRM Packera_hartiana
E Pericallis_cruenta
S Elekmania_barahonensis
S Werneria_aretioides
S Xenophyllum_lycopodioides
M Packera_loratifolia
M Packera_2moranii
M Packera_zimapanica
M Packera_2actinella
M Packera_2toluccana
M Packera_umbraculifera
30
25
20
15
10
5
0
Millions of years ago

**BioGeoBEARS DIVALIKE on Packera − paragone−nf:MO**
**ancstates: global optim, 4 areas max. d=0.0073; e=0.0085; j=0; LnL=−328.43**

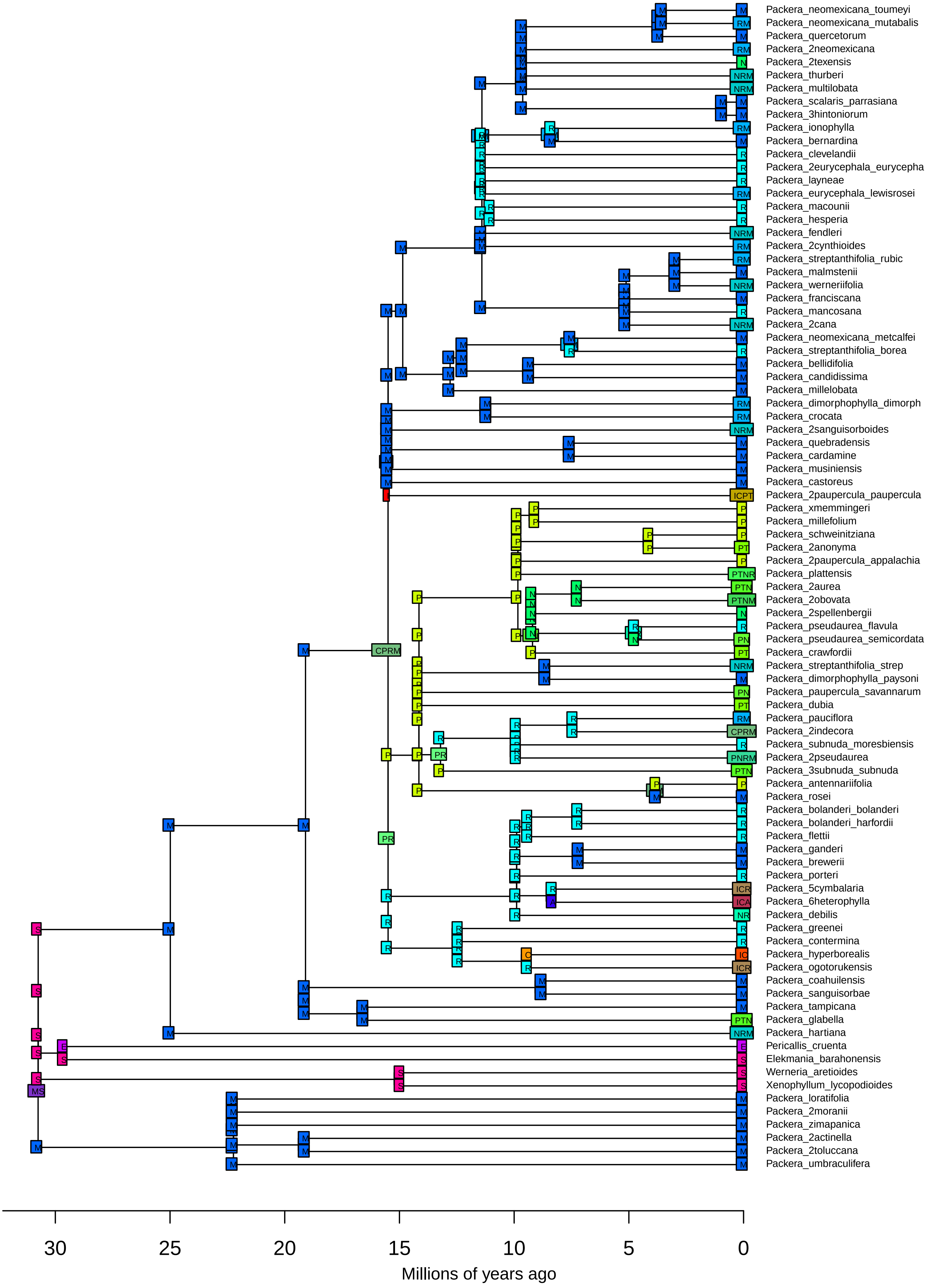

**BioGeoBEARS DIVALIKE on Packera − paragone−nf:MO**
**ancstates: global optim, 4 areas max. d=0.0073; e=0.0085; j=0; LnL=−328.43**

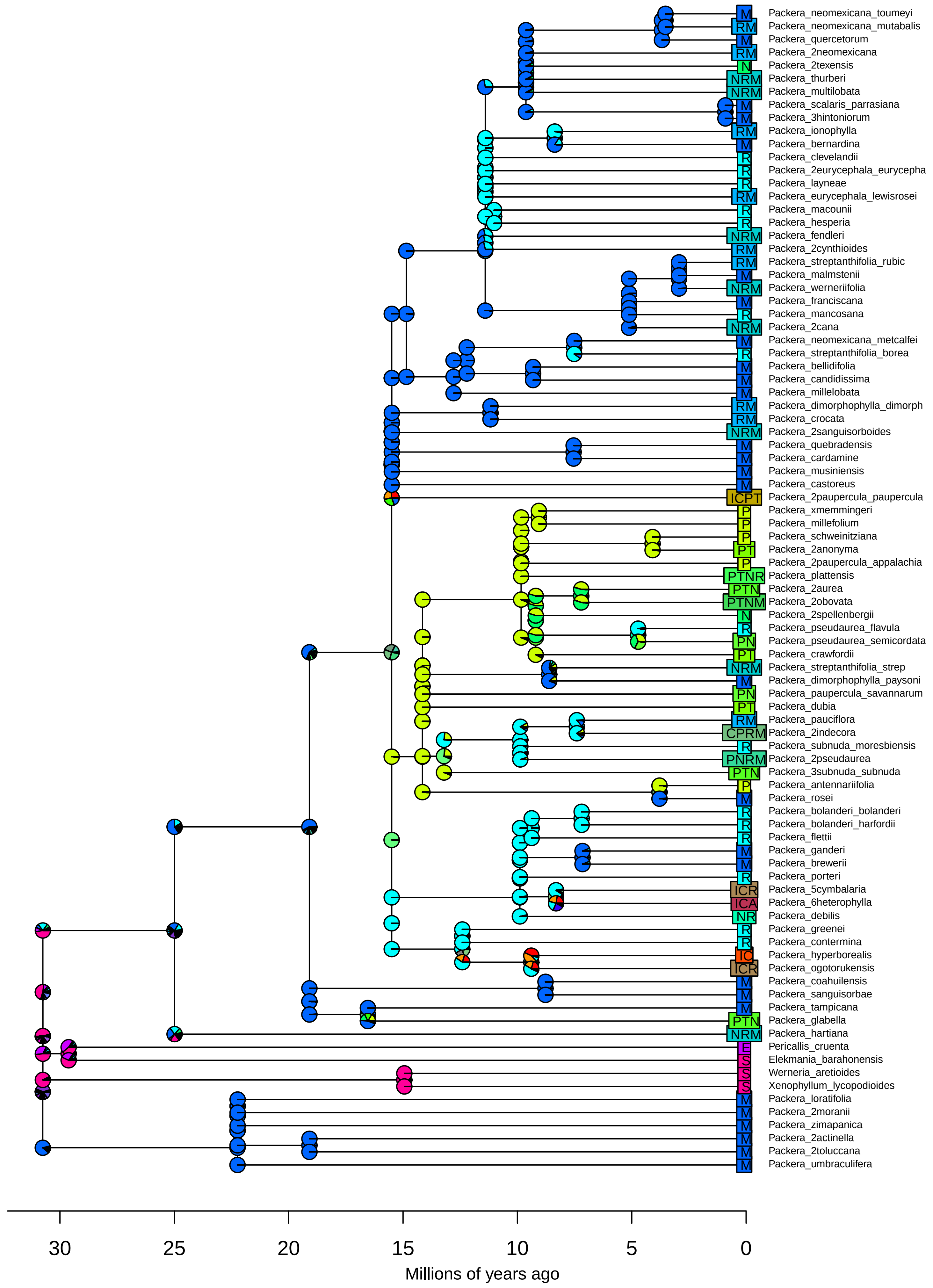

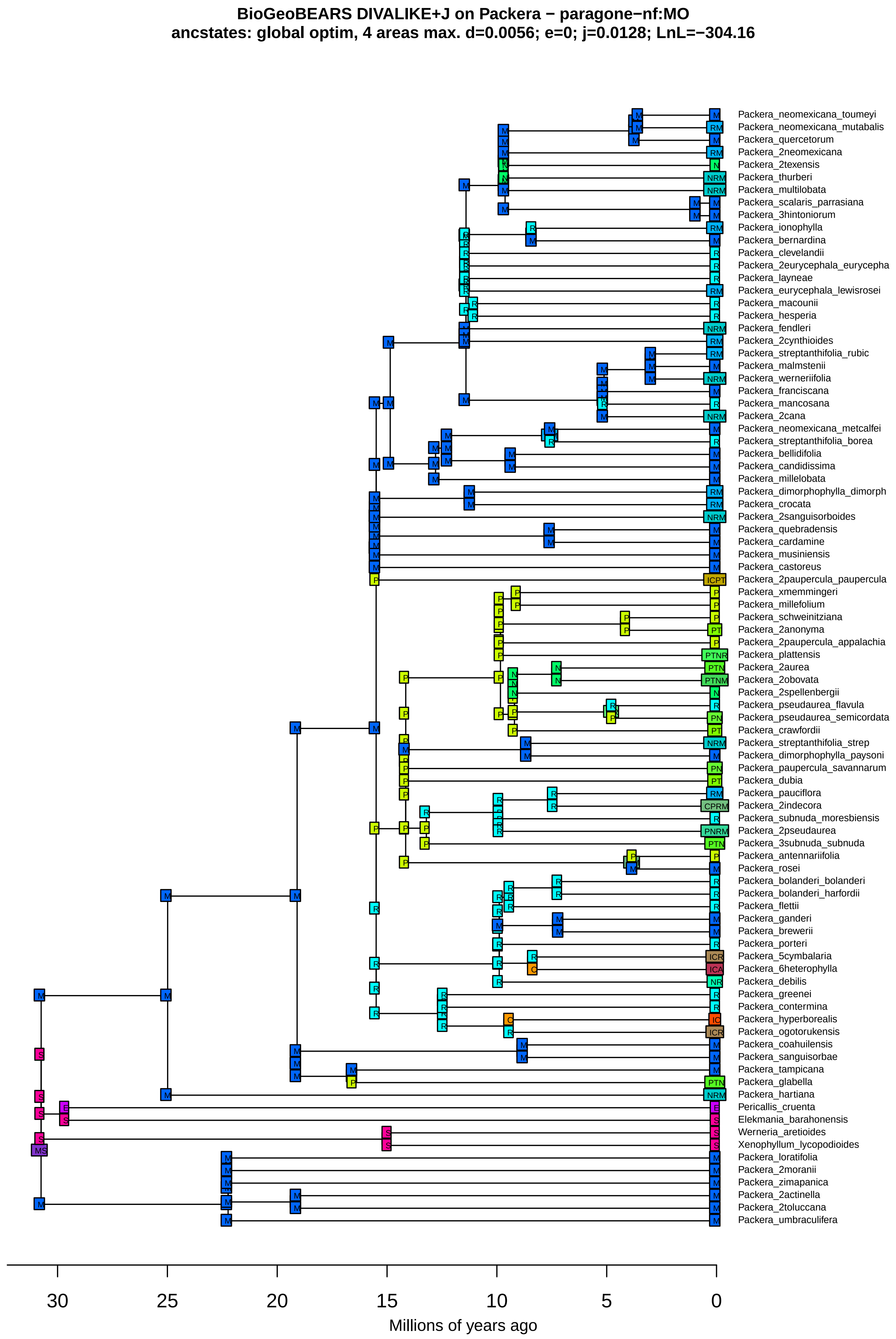

BioGeoBEARS DIVALIKE+J on Packera − paragone−nf:MO
ancstates: global optim, 4 areas max. d=0.0056; e=0; j=0.0128; LnL=−304.16
Packera_neomexicana_toumeyi
Packera_neomexicana_mutabalis
Packera_quercetorum
Packera_2neomexicana
Packera_2texensis
Packera_thurberi
Packera_multilobata
Packera_scalaris_parrasiana
Packera_3hintoniorum
Packera_ionophylla
Packera_bernardina
Packera_clevelandii
Packera_2eurycephala_eurycepha
Packera_layneae
Packera_eurycephala_lewisrosei
Packera_macounii
Packera_hesperia
Packera_fendleri
Packera_2cynthioides
Packera_streptanthifolia_rubic
Packera_malmstenii
Packera_werneriifolia
Packera_franciscana
Packera_mancosana
Packera_2cana
Packera_neomexicana_metcalfei
Packera_streptanthifolia_borea
Packera_bellidifolia
Packera_candidissima
Packera_millelobata
Packera_dimorphophylla_dimorph
Packera_crocata
Packera_2sanguisorboides
Packera_quebradensis
Packera_cardamine
Packera_musiniensis
Packera_castoreus
Packera_2paupercula_paupercula
Packera_xmemmingeri
Packera_millefolium
Packera_schweinitziana
Packera_2anonyma
Packera_2paupercula_appalachia
Packera_plattensis
Packera_2aurea
Packera_2obovata
Packera_2spellenbergii
Packera_pseudaurea_flavula
Packera_pseudaurea_semicordata
Packera_crawfordii
Packera_streptanthifolia_strep
Packera_dimorphophylla_paysoni
Packera_paupercula_savannarum
Packera_dubia
Packera_pauciflora
Packera_2indecora
Packera_subnuda_moresbiensis
Packera_2pseudaurea
Packera_3subnuda_subnuda
Packera_antennariifolia
Packera_rosei
Packera_bolanderi_bolanderi
Packera_bolanderi_harfordii
Packera_flettii
Packera_ganderi
Packera_brewerii
Packera_porteri
Packera_5cymbalaria
Packera_6heterophylla
Packera_debilis
Packera_greenei
Packera_contermina
Packera_hyperborealis
Packera_ogotorukensis
Packera_coahuilensis
Packera_sanguisorbae
Packera_tampicana
Packera_glabella
Packera_hartiana
Pericallis_cruenta
Elekmania_barahonensis
Werneria_aretioides
Xenophyllum_lycopodioides
Packera_loratifolia
Packera_2moranii
Packera_zimapanica
Packera_2actinella
Packera_2toluccana
Packera_umbraculifera
30
25
20
15
10
5
0
Millions of years ago

**BioGeoBEARS DIVALIKE+J on Packera − paragone−nf:MO**
**ancstates: global optim, 4 areas max. d=0.0056; e=0; j=0.0128; LnL=−304.16**

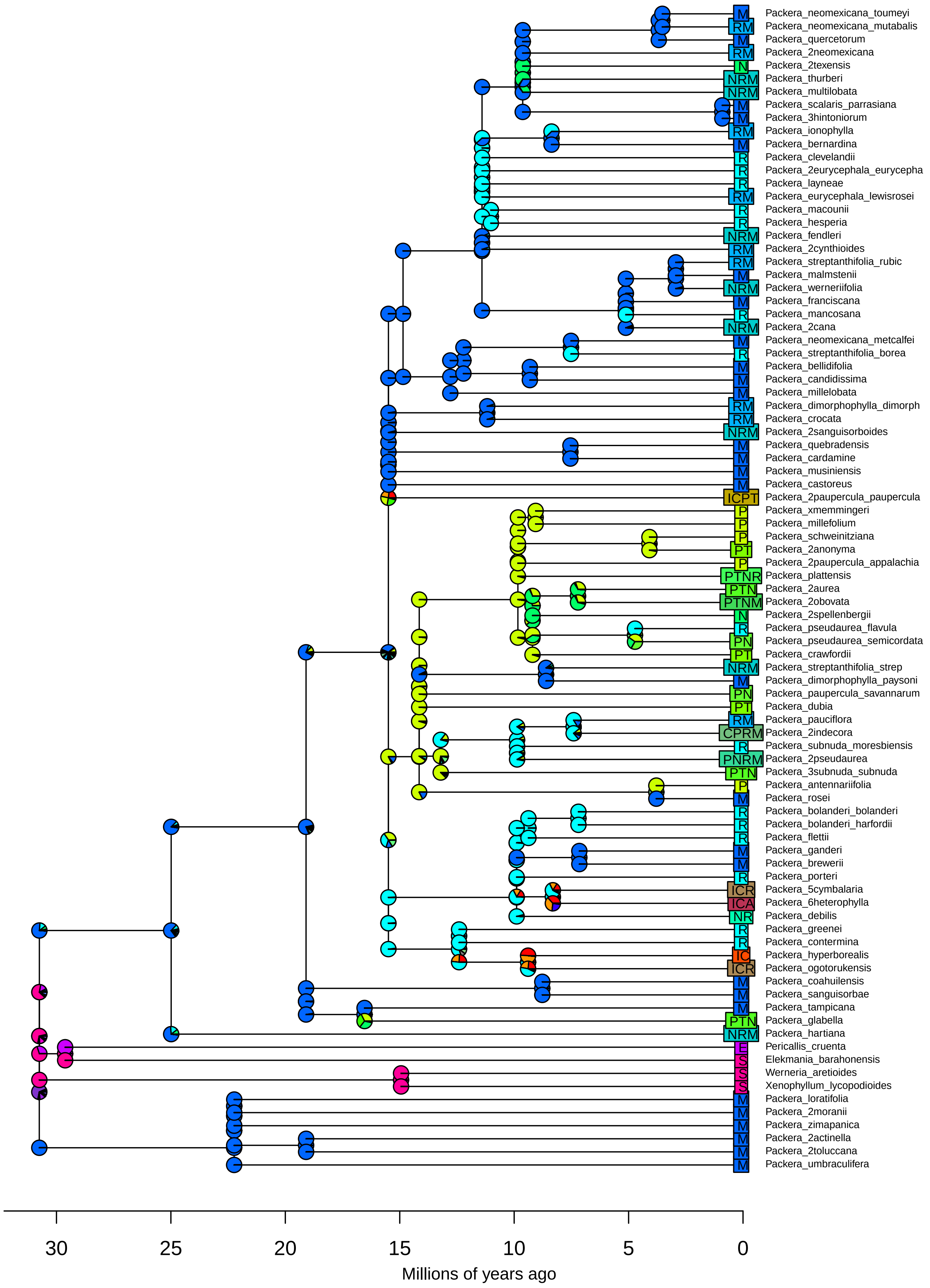

BioGeoBEARS BAYAREALIKE on Packera − paragone−nf:MO
ancstates: global optim, 4 areas max. d=0.0031; e=0.0475; j=0; LnL=−286.21

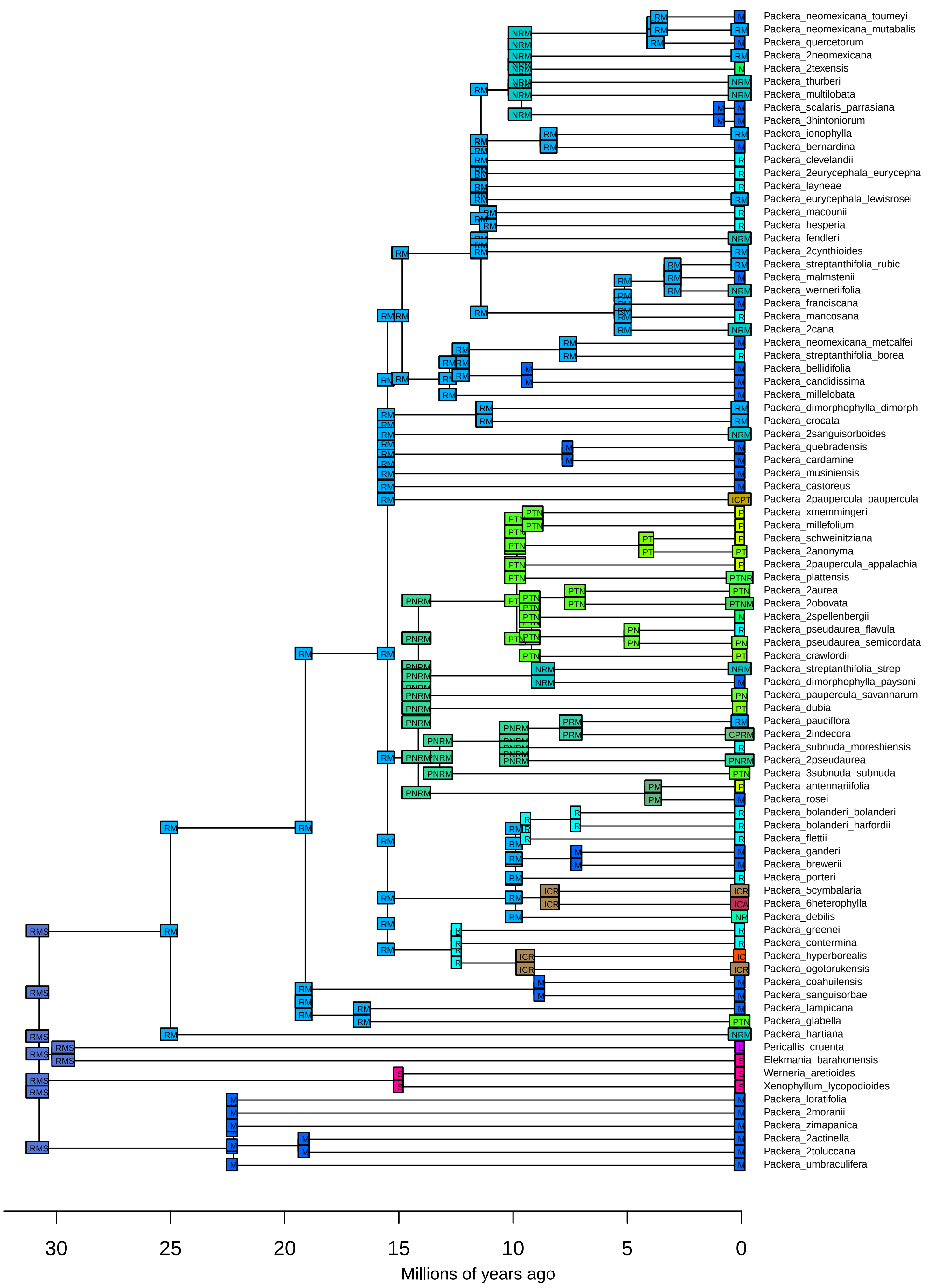

**BioGeoBEARS BAYAREALIKE on Packera − paragone−nf:MO**
**ancstates: global optim, 4 areas max. d=0.0031; e=0.0475; j=0; LnL=−286.21**

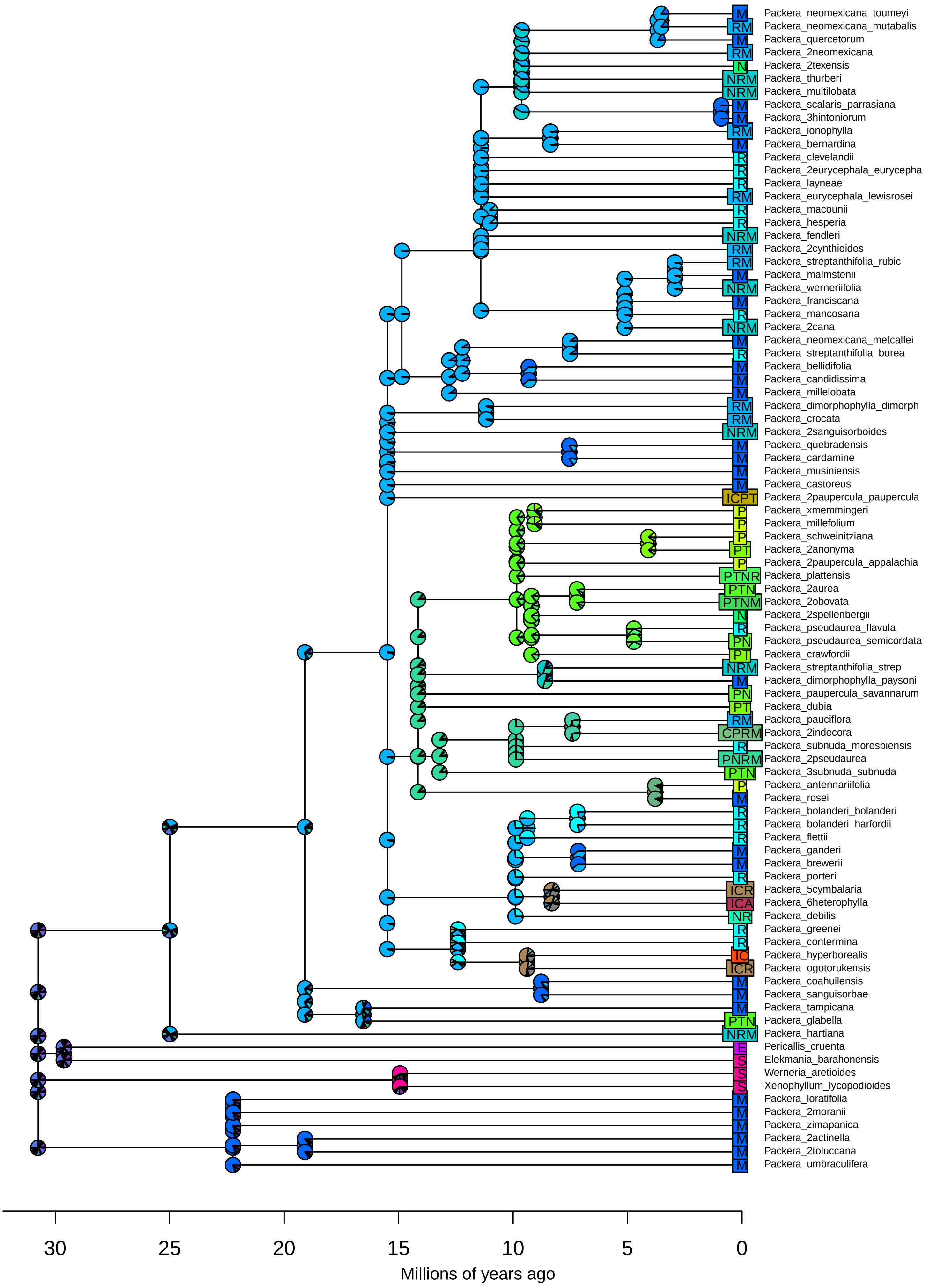

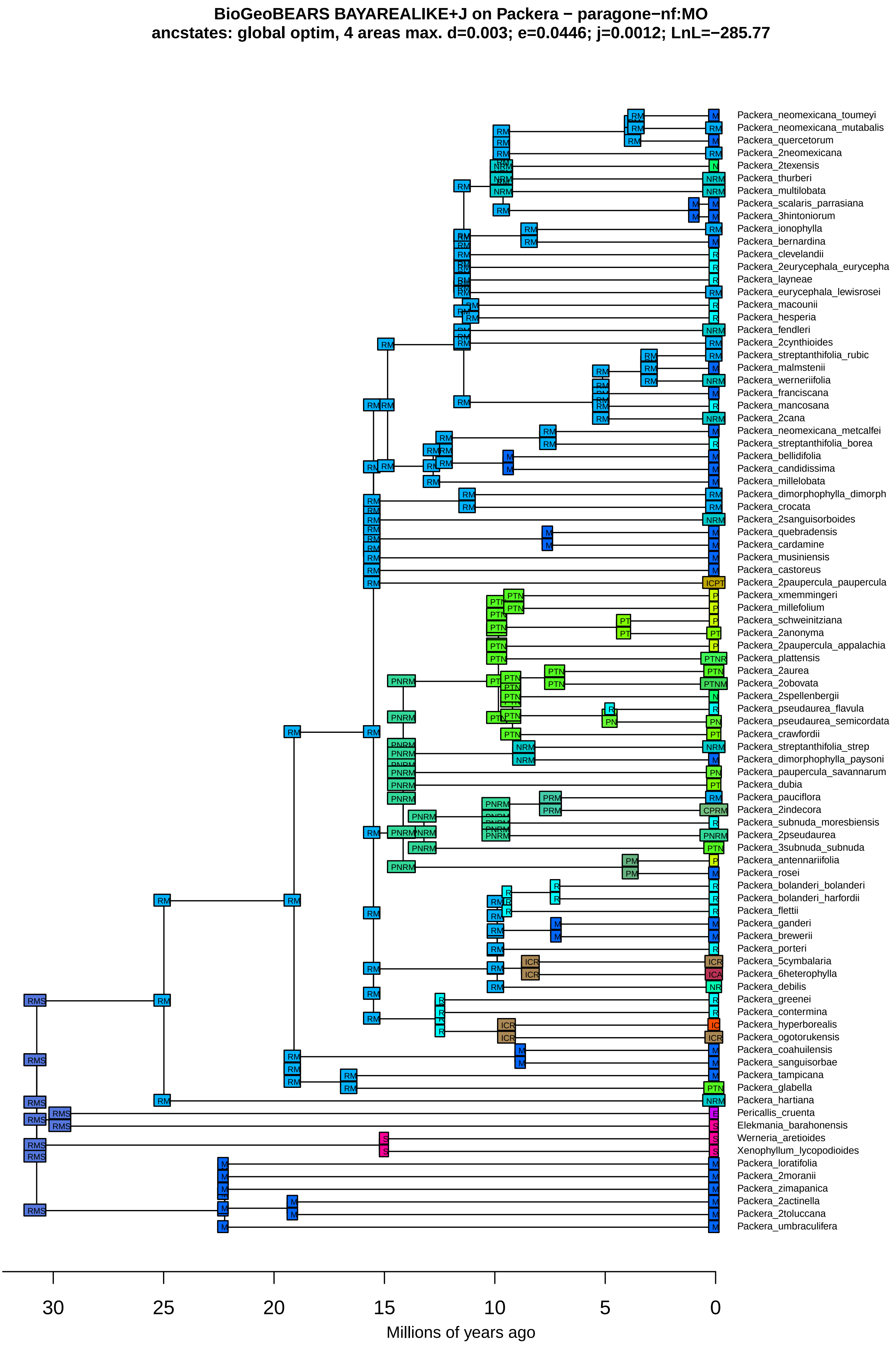
BioGeoBEARS BAYAREALIKE+J on Packera − paragone−nf:MO
ancstates: global optim, 4 areas max. d=0.003; e=0.0446; j=0.0012; LnL=−285.77
Packera_neomexicana_toumeyi
Packera_neomexicana_mutabalis
Packera_quercetorum
Packera_2neomexicana
Packera_2texensis
Packera_thurberi
Packera_multilobata
Packera_scalaris_parrasiana
Packera_3hintoniorum
Packera_ionophylla
Packera_bernardina
Packera_clevelandii
Packera_2eurycephala_eurycepha
Packera_layneae
Packera_eurycephala_lewisrosei
Packera_macounii
Packera_hesperia
Packera_fendleri
Packera_2cynthioides
Packera_streptanthifolia_rubic
Packera_malmstenii
Packera_werneriifolia
Packera_franciscana
Packera_mancosana
Packera_2cana
Packera_neomexicana_metcalfei
Packera_streptanthifolia_borea
Packera_bellidifolia
Packera_candidissima
Packera_millelobata
Packera_dimorphophylla_dimorph
Packera_crocata
Packera_2sanguisorboides
Packera_quebradensis
Packera_cardamine
Packera_musiniensis
Packera_castoreus
Packera_2paupercula_paupercula
Packera_xmemmingeri
Packera_millefolium
Packera_schweinitziana
Packera_2anonyma
Packera_2paupercula_appalachia
Packera_plattensis
Packera_2aurea
Packera_2obovata
Packera_2spellenbergii
Packera_pseudaurea_flavula
Packera_pseudaurea_semicordata
Packera_crawfordii
Packera_streptanthifolia_strep
Packera_dimorphophylla_paysoni
Packera_paupercula_savannarum
Packera_dubia
Packera_pauciflora
Packera_2indecora
Packera_subnuda_moresbiensis
Packera_2pseudaurea
Packera_3subnuda_subnuda
Packera_antennariifolia
Packera_rosei
Packera_bolanderi_bolanderi
Packera_bolanderi_harfordii
Packera_flettii
Packera_ganderi
Packera_brewerii
Packera_porteri
Packera_5cymbalaria
Packera_6heterophylla
Packera_debilis
Packera_greenei
Packera_contermina
Packera_hyperborealis
Packera_ogotorukensis
Packera_coahuilensis
Packera_sanguisorbae
Packera_tampicana
Packera_glabella
Packera_hartiana
Pericallis_cruenta
Elekmania_barahonensis
Werneria_aretioides
Xenophyllum_lycopodioides
Packera_loratifolia
Packera_2moranii
Packera_zimapanica
Packera_2actinella
Packera_2toluccana
Packera_umbraculifera
30
25
20
15
10
5
0
Millions of years ago

**BioGeoBEARS BAYAREALIKE+J on Packera − paragone−nf:MO**
**ancstates: global optim, 4 areas max. d=0.003; e=0.0446; j=0.0012; LnL=−285.77**

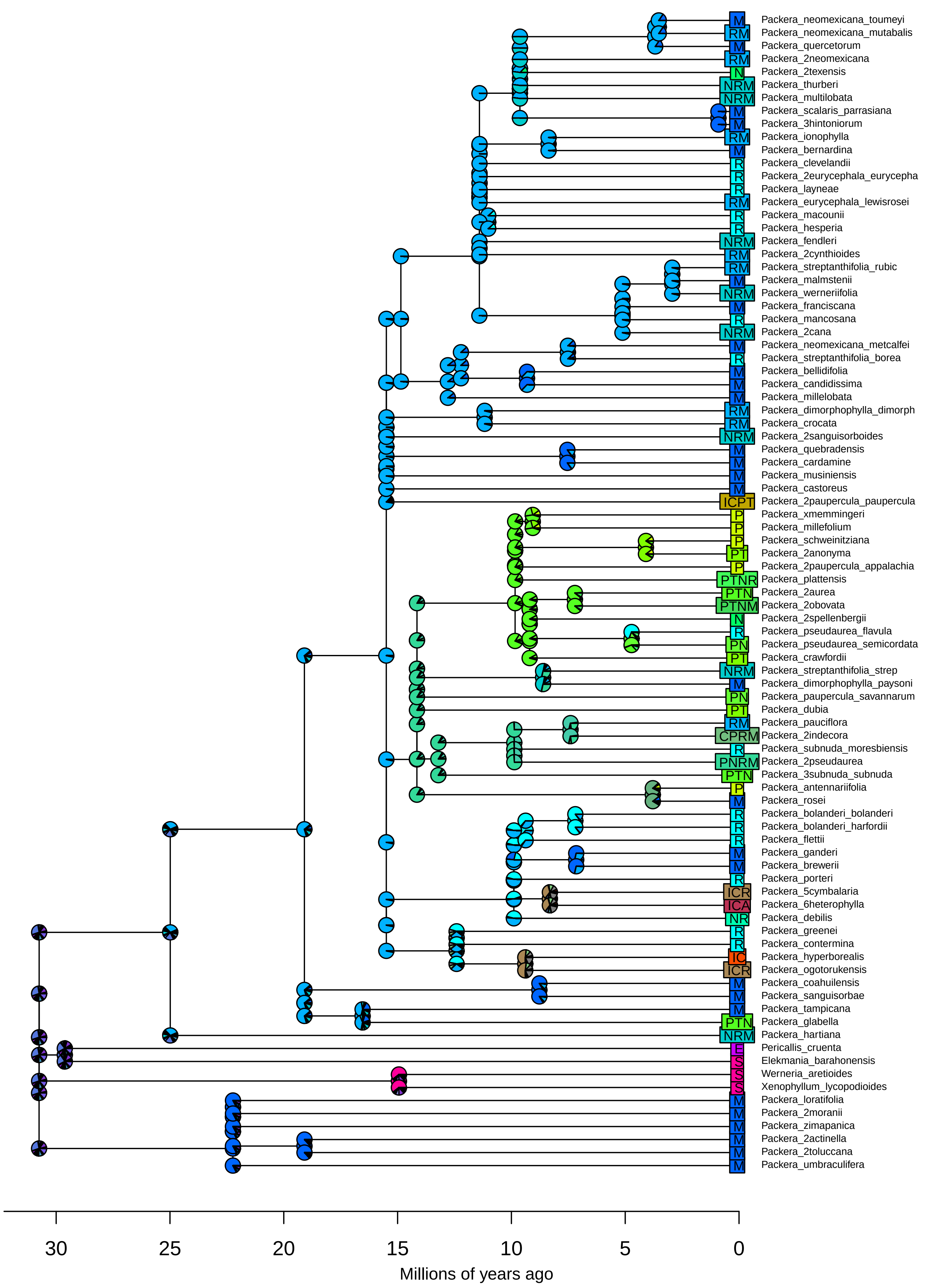

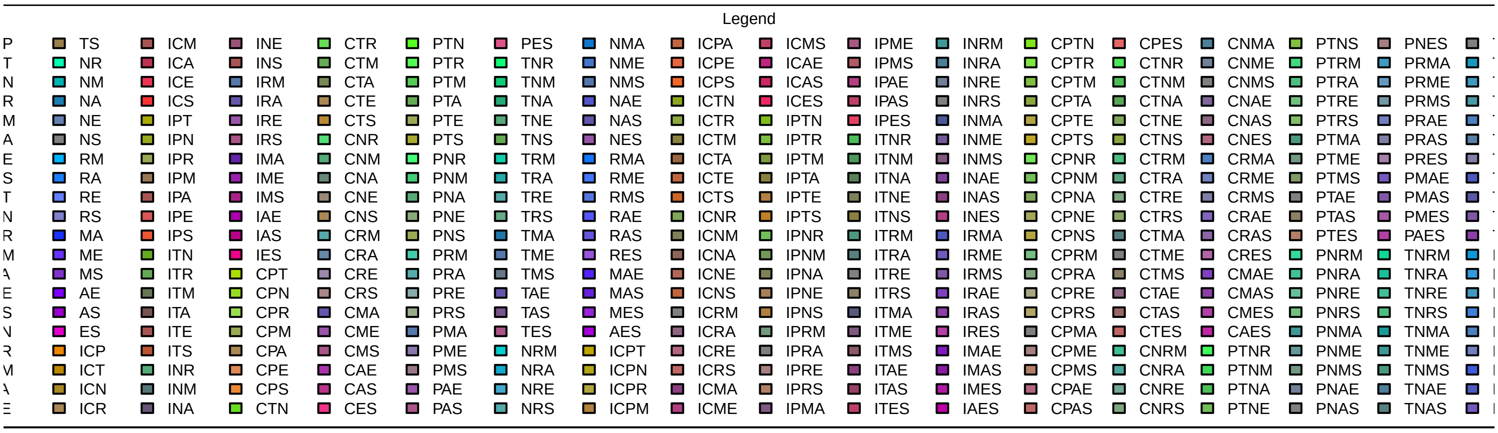
Legend
TS NR NM NA NE NS RM RA RE RS MA ME MS AE AS ES ICP ICT ICN ICR
ICM ICA ICE ICS IPT IPN IPR IPM IPA IPE IPS ITN ITR ITM ITA ITE ITS INR INM INA
INE INS IRM IRA IRE IRS IMA IME IMS IAE IAS IES CPT CPN CPR CPM CPA CPE CPS CTN
CTR CTM CTA CTE CTS CNR CNM CNA CNE CNS CRM CRA CRE CRS CMA CME CMS CAE CAS CES
PTN PTR PTM PTA PTE PTS PNR PNM PNA PNE PNS PRM PRA PRE PRS PMA PME PMS PAE PAS
PES TNR TNM TNA TNE TNS TRM TRA TRE TRS TMA TME TMS TAE TAS TES NRM NRA NRE NRS
NMA NME NMS NAE NAS NES RMA RME RMS RAE RAS RES MAE MAS MES AES ICPT ICPN ICPR ICPM
ICPA ICPE ICPS ICTN ICTR ICTM ICTA ICTE ICTS ICNR ICNM ICNA ICNE ICNS ICRM ICRA ICRE ICRS ICMA ICME
ICMS ICAE ICAS ICES IPTN IPTR IPTM IPTA IPTE IPTS IPNR IPNM IPNA IPNE IPNS IPRM IPRA IPRE IPRS IPMA
IPME IPMS IPAE IPAS IPES ITNR ITNM ITNA ITNE ITNS ITRM ITRA ITRE ITRS ITMA ITME ITMS ITAE ITAS ITES
INRM INRA INRE INRS INMA INME INMS INAE INAS INES IRMA IRME IRMS IRAE IRAS IRES IMAE IMAS IMES IAES
CPTN CPTR CPTM CPTA CPTE CPTS CPNR CPNM CPNA CPNE CPNS CPRM CPRA CPRE CPRS CPMA CPME CPMS CPAE CPAS
CPES CTNR CTNM CTNA CTNE CTNS CTRM CTRA CTRE CTRS CTMA CTME CTMS CTAE CTAS CTES CNRM CNRA CNRE CNRS
CNMA CNME CNMS CNAE CNAS CNES CRMA CRME CRMS CRAE CRAS CRES CMAE CMAS CMES CAES PTNR PTNM PTNA PTNE
PTNS PTRM PTRA PTRE PTRS PTMA PTME PTMS PTAE PTAS PTES PNRM PNRA PNRE PNRS PNMA PNME PNMS PNAE PNAS
PNES PRMA PRME PRMS PRAE PRAS PRES PMAE PMAS PMES PAES TNRM TNRA TNRE TNRS TNMA TNME TNMS TNAE TNAS

**BioGeoBEARS DEC on Packera − paragone−nf: 1−to−1**
**ancstates: global optim, 4 areas max. d=0.006; e=0.0053; j=0; LnL=−306.60**

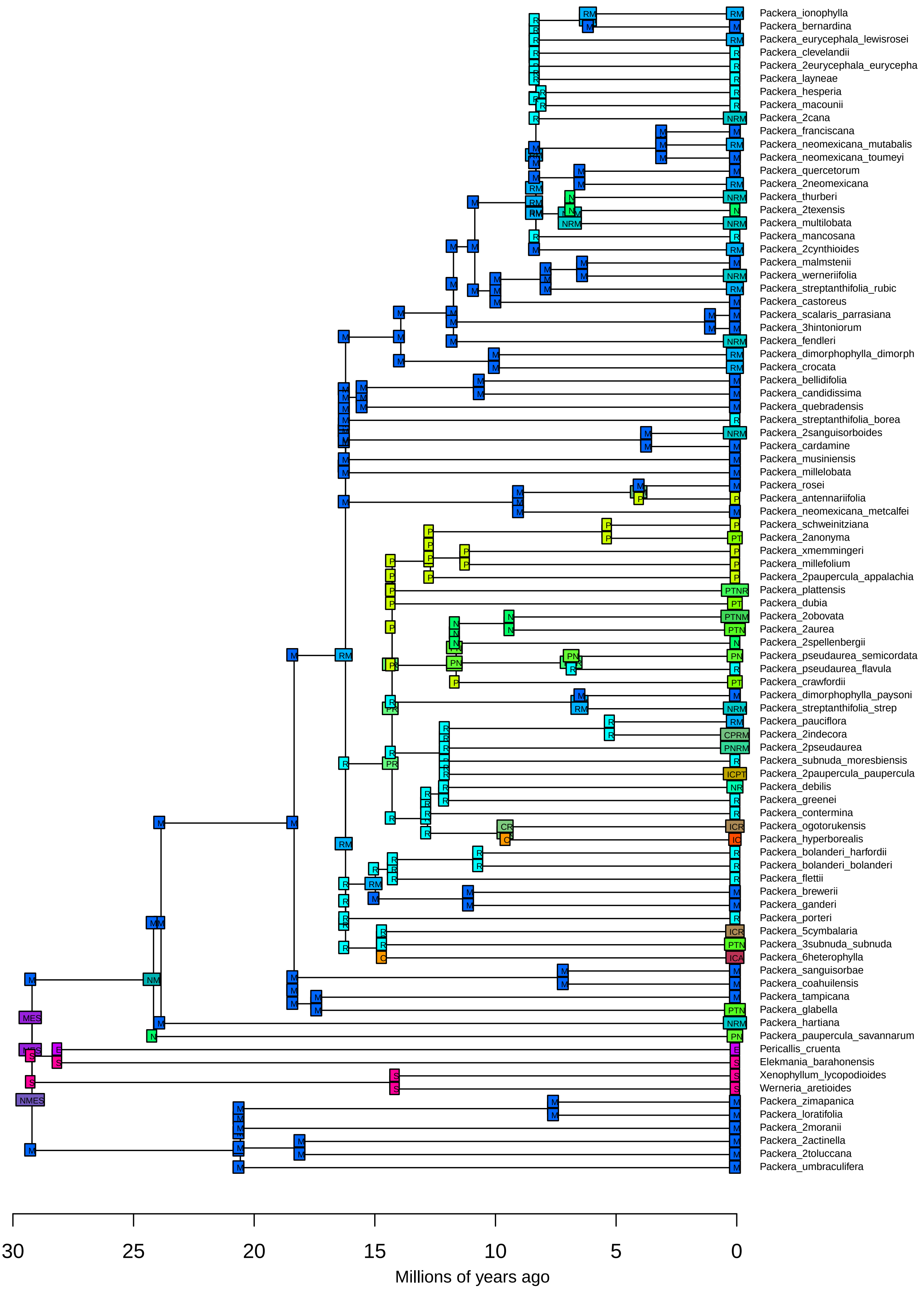

**BioGeoBEARS DEC on Packera − paragone−nf: 1−to−1**
**ancstates: global optim, 4 areas max. d=0.006; e=0.0053; j=0; LnL=−306.60**

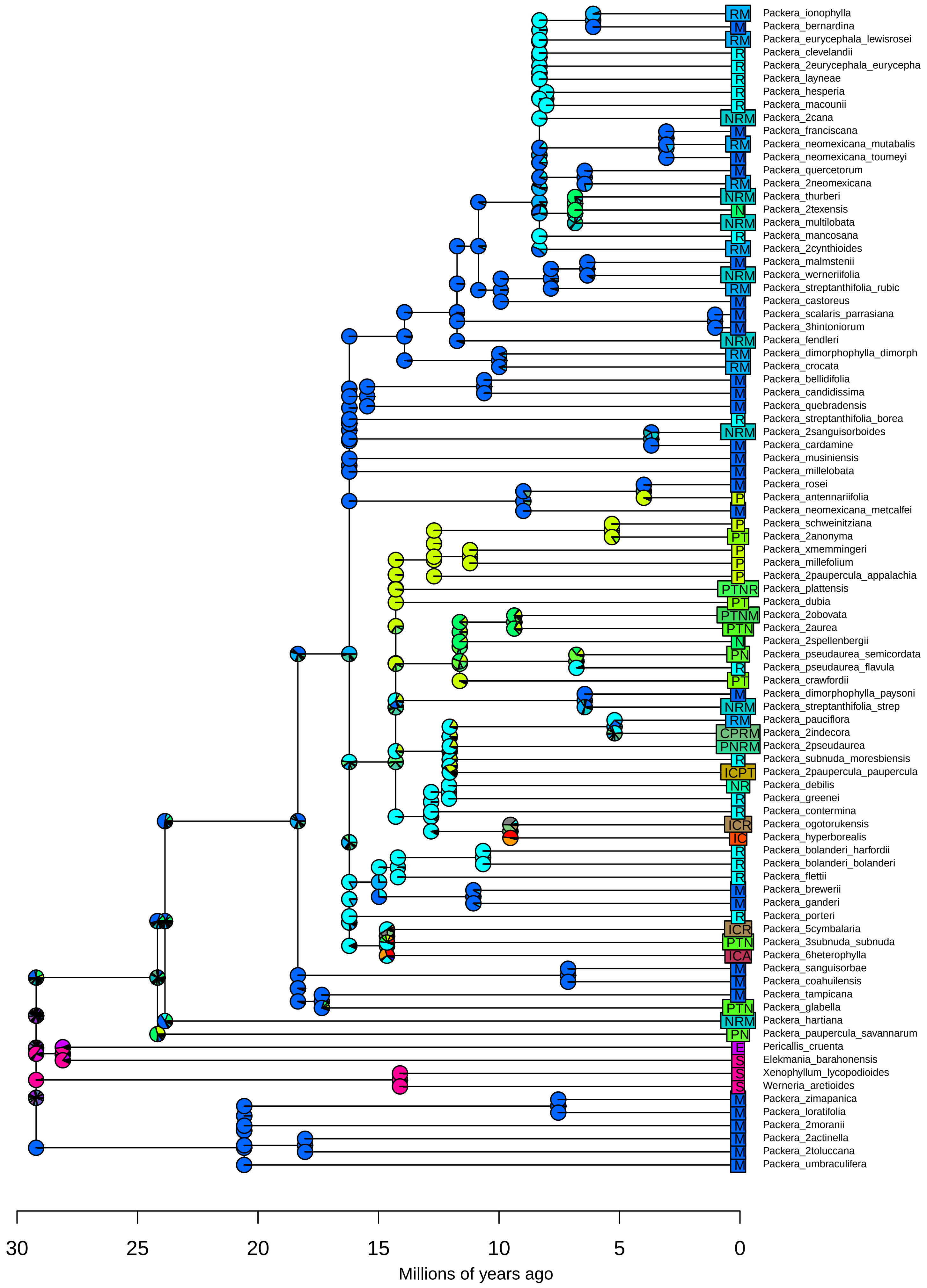

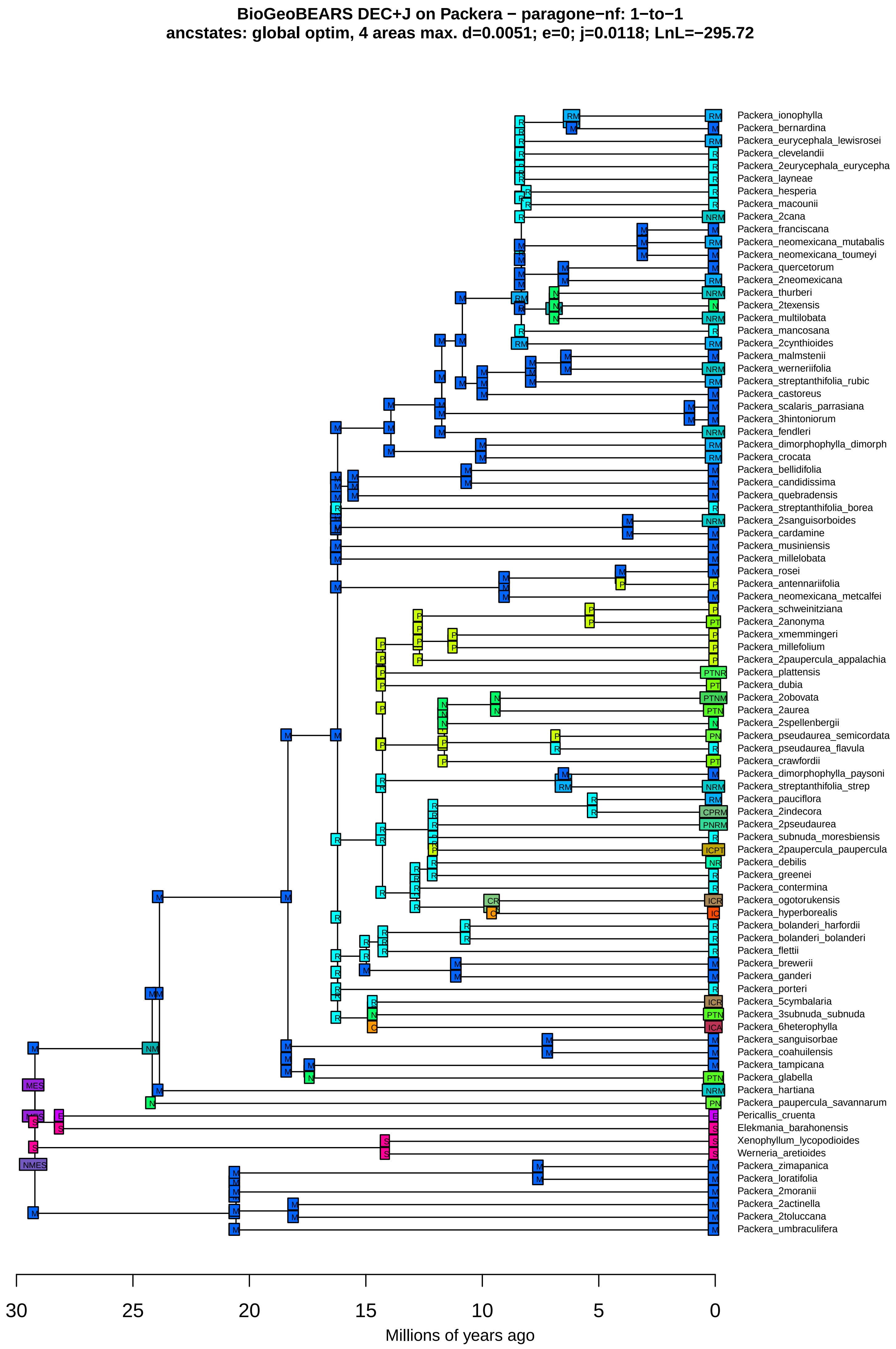
BioGeoBEARS DEC+J on Packera − paragone−nf: 1−to−1
ancstates: global optim, 4 areas max. d=0.0051; e=0; j=0.0118; LnL=−295.72
Packera_ionophylla
Packera_bernardina
Packera_eurycephala_lewisrosei
Packera_clevelandii
Packera_2eurycephala_eurycepha
Packera_layneae
Packera_hesperia
Packera_macounii
Packera_2cana
Packera_franciscana
Packera_neomexicana_mutabalis
Packera_neomexicana_toumeyi
Packera_quercetorum
Packera_2neomexicana
Packera_thurberi
Packera_2texensis
Packera_multilobata
Packera_mancosana
Packera_2cynthioides
Packera_malmstenii
Packera_werneriifolia
Packera_streptanthifolia_rubic
Packera_castoreus
Packera_scalaris_parrasiana
Packera_3hintoniorum
Packera_fendleri
Packera_dimorphophylla_dimorph
Packera_crocata
Packera_bellidifolia
Packera_candidissima
Packera_quebradensis
Packera_streptanthifolia_borea
Packera_2sanguisorboides
Packera_cardamine
Packera_musiniensis
Packera_millelobata
Packera_rosei
Packera_antennariifolia
Packera_neomexicana_metcalfei
Packera_schweinitziana
Packera_2anonyma
Packera_xmemmingeri
Packera_millefolium
Packera_2paupercula_appalachia
Packera_plattensis
Packera_dubia
Packera_2obovata
Packera_2aurea
Packera_2spellenbergii
Packera_pseudaurea_semicordata
Packera_pseudaurea_flavula
Packera_crawfordii
Packera_dimorphophylla_paysoni
Packera_streptanthifolia_strep
Packera_pauciflora
Packera_2indecora
Packera_2pseudaurea
Packera_subnuda_moresbiensis
Packera_2paupercula_paupercula
Packera_debilis
Packera_greenei
Packera_contermina
Packera_ogotorukensis
Packera_hyperborealis
Packera_bolanderi_harfordii
Packera_bolanderi_bolanderi
Packera_flettii
Packera_brewerii
Packera_ganderi
Packera_porteri
Packera_5cymbalaria
Packera_3subnuda_subnuda
Packera_6heterophylla
Packera_sanguisorbae
Packera_coahuilensis
Packera_tampicana
Packera_glabella
Packera_hartiana
Packera_paupercula_savannarum
Pericallis_cruenta
Elekmania_barahonensis
Xenophyllum_lycopodioides
Werneria_aretioides
Packera_zimapanica
Packera_loratifolia
Packera_2moranii
Packera_2actinella
Packera_2toluccana
Packera_umbraculifera
30
25
20
15
10
5
0
Millions of years ago

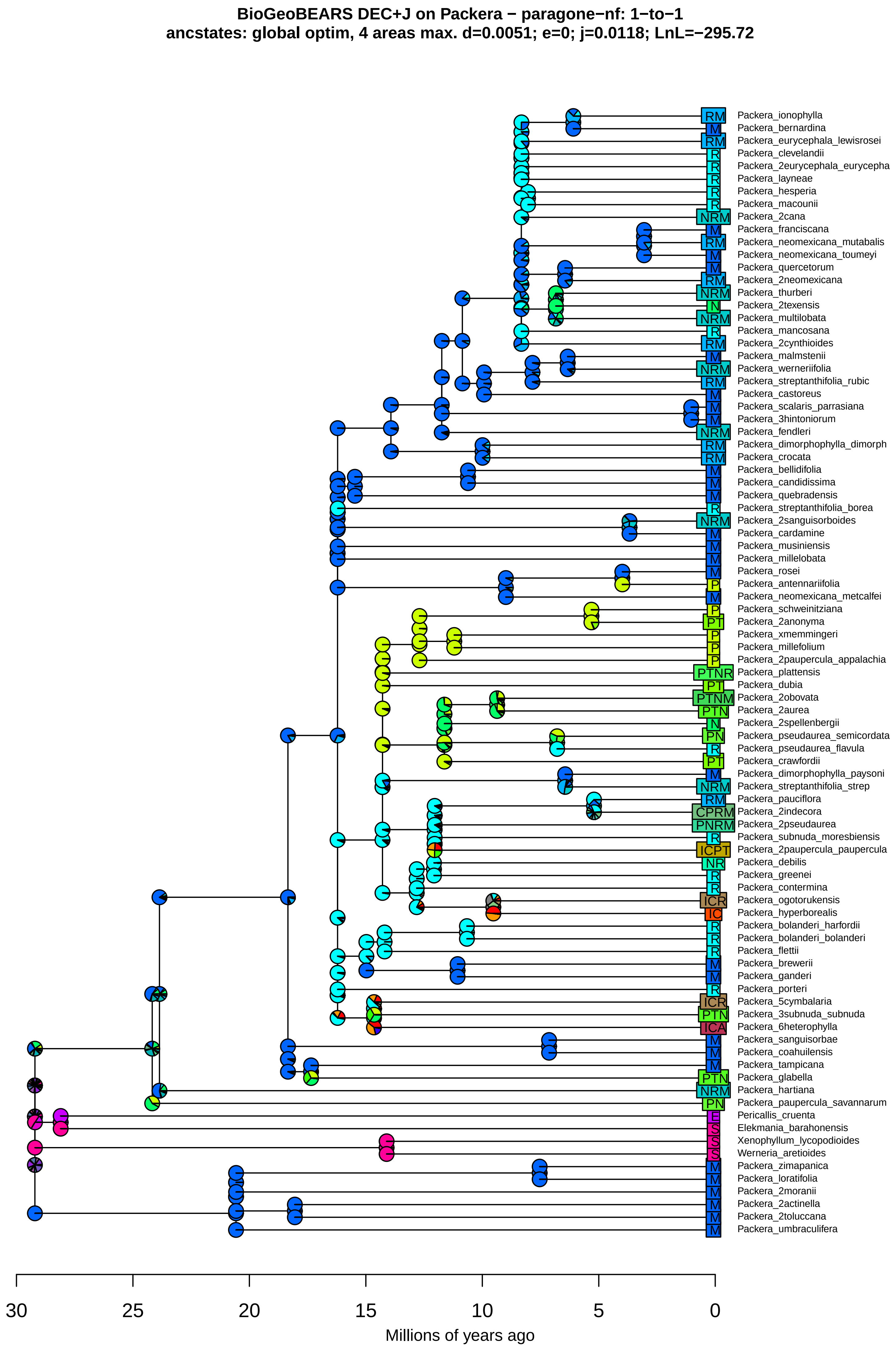
BioGeoBEARS DEC+J on Packera − paragone−nf: 1−to−1
ancstates: global optim, 4 areas max. d=0.0051; e=0; j=0.0118; LnL=−295.72
RM Packera_ionophylla
M Packera_bernardina
RM Packera_eurycephala_lewisrosei
R Packera_clevelandii
R Packera_2eurycephala_eurycepha
R Packera_layneae
R Packera_hesperia
R Packera_macounii
NRM Packera_2cana
M Packera_franciscana
RM Packera_neomexicana_mutabalis
M Packera_neomexicana_toumeyi
M Packera_quercetorum
RM Packera_2neomexicana
NRM Packera_thurberi
N Packera_2texensis
NRM Packera_multilobata
R Packera_mancosana
RM Packera_2cynthioides
M Packera_malmstenii
NRM Packera_werneriifolia
RM Packera_streptanthifolia_rubic
M Packera_castoreus
M Packera_scalaris_parrasiana
M Packera_3hintoniorum
NRM Packera_fendleri
RM Packera_dimorphophylla_dimorph
RM Packera_crocata
M Packera_bellidifolia
M Packera_candidissima
M Packera_quebradensis
R Packera_streptanthifolia_borea
NRM Packera_2sanguisorboides
M Packera_cardamine
M Packera_musiniensis
M Packera_millelobata
M Packera_rosei
P Packera_antennariifolia
M Packera_neomexicana_metcalfei
P Packera_schweinitziana
PT Packera_2anonyma
P Packera_xmemmingeri
P Packera_millefolium
P Packera_2paupercula_appalachia
PTNR Packera_plattensis
PT Packera_dubia
PTNM Packera_2obovata
PTN Packera_2aurea
N Packera_2spellenbergii
PN Packera_pseudaurea_semicordata
R Packera_pseudaurea_flavula
PT Packera_crawfordii
M Packera_dimorphophylla_paysoni
NRM Packera_streptanthifolia_strep
RM Packera_pauciflora
CPRM Packera_2indecora
PNRM Packera_2pseudaurea
R Packera_subnuda_moresbiensis
ICPT Packera_2paupercula_paupercula
NR Packera_debilis
R Packera_greenei
R Packera_contermina
ICR Packera_ogotorukensis
IC Packera_hyperborealis
R Packera_bolanderi_harfordii
R Packera_bolanderi_bolanderi
R Packera_flettii
M Packera_brewerii
M Packera_ganderi
R Packera_porteri
ICR Packera_5cymbalaria
PTN Packera_3subnuda_subnuda
ICA Packera_6heterophylla
M Packera_sanguisorbae
M Packera_coahuilensis
M Packera_tampicana
PTN Packera_glabella
NRM Packera_hartiana
PN Packera_paupercula_savannarum
E Pericallis_cruenta
S Elekmania_barahonensis
S Xenophyllum_lycopodioides
S Werneria_aretioides
M Packera_zimapanica
M Packera_loratifolia
M Packera_2moranii
M Packera_2actinella
M Packera_2toluccana
M Packera_umbraculifera
30
25
20
15
10
5
0
Millions of years ago

**BioGeoBEARS DIVALIKE on Packera − paragone−nf: 1−to−1**
**ancstates: global optim, 4 areas max. d=0.0071; e=0.008; j=0; LnL=−326.35**

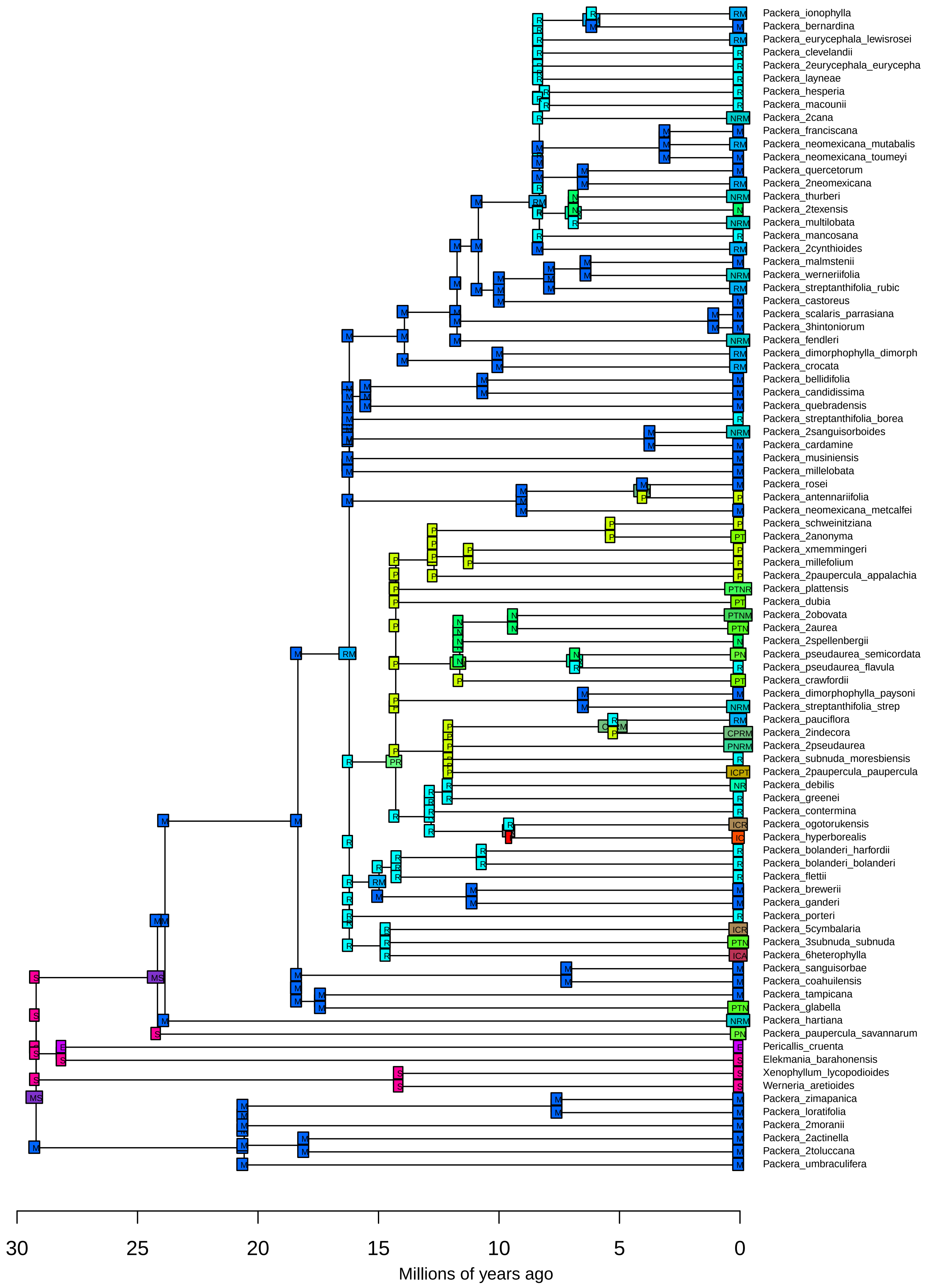

**BioGeoBEARS DIVALIKE on Packera − paragone−nf: 1−to−1**
**ancstates: global optim, 4 areas max. d=0.0071; e=0.008; j=0; LnL=−326.35**

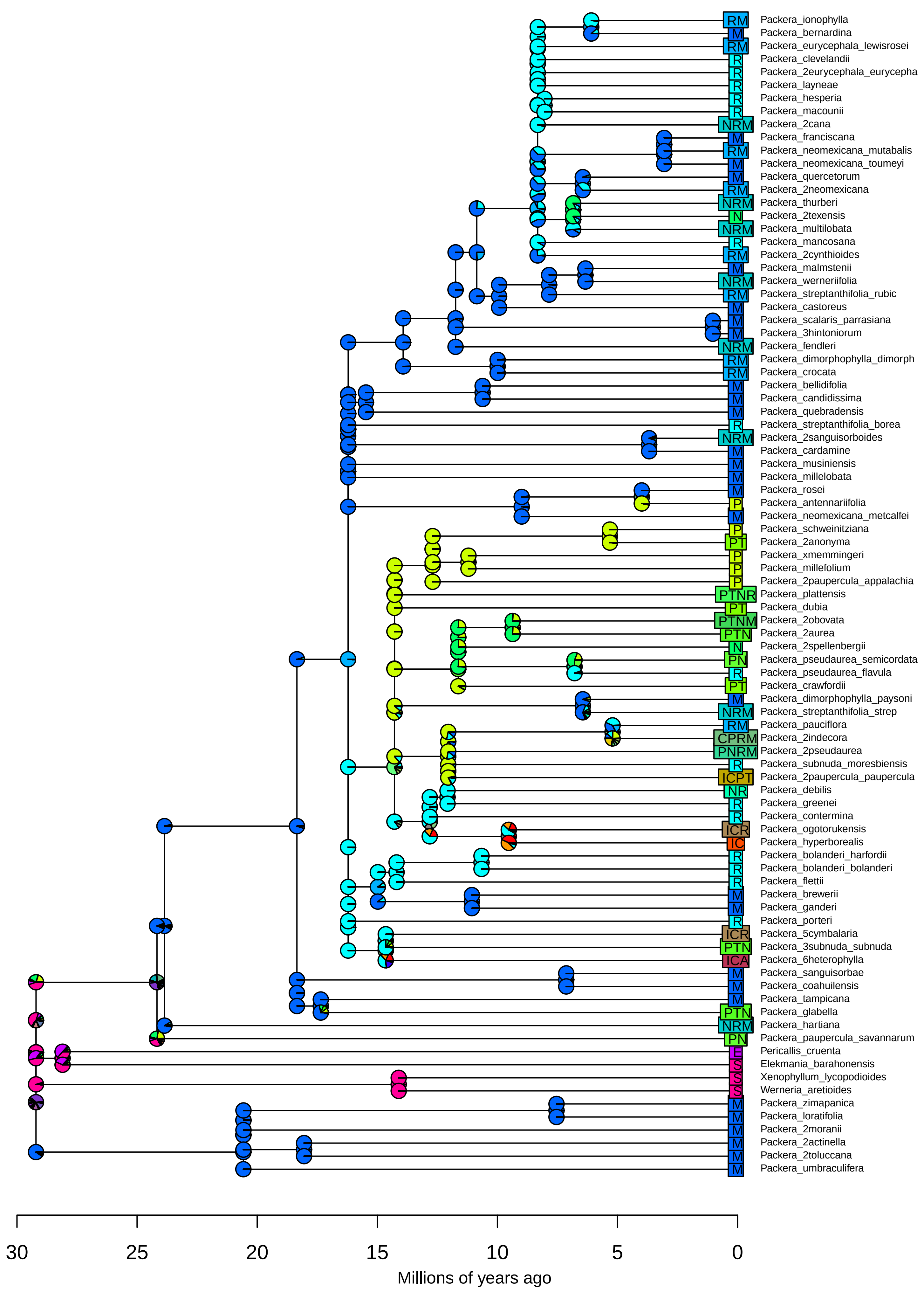

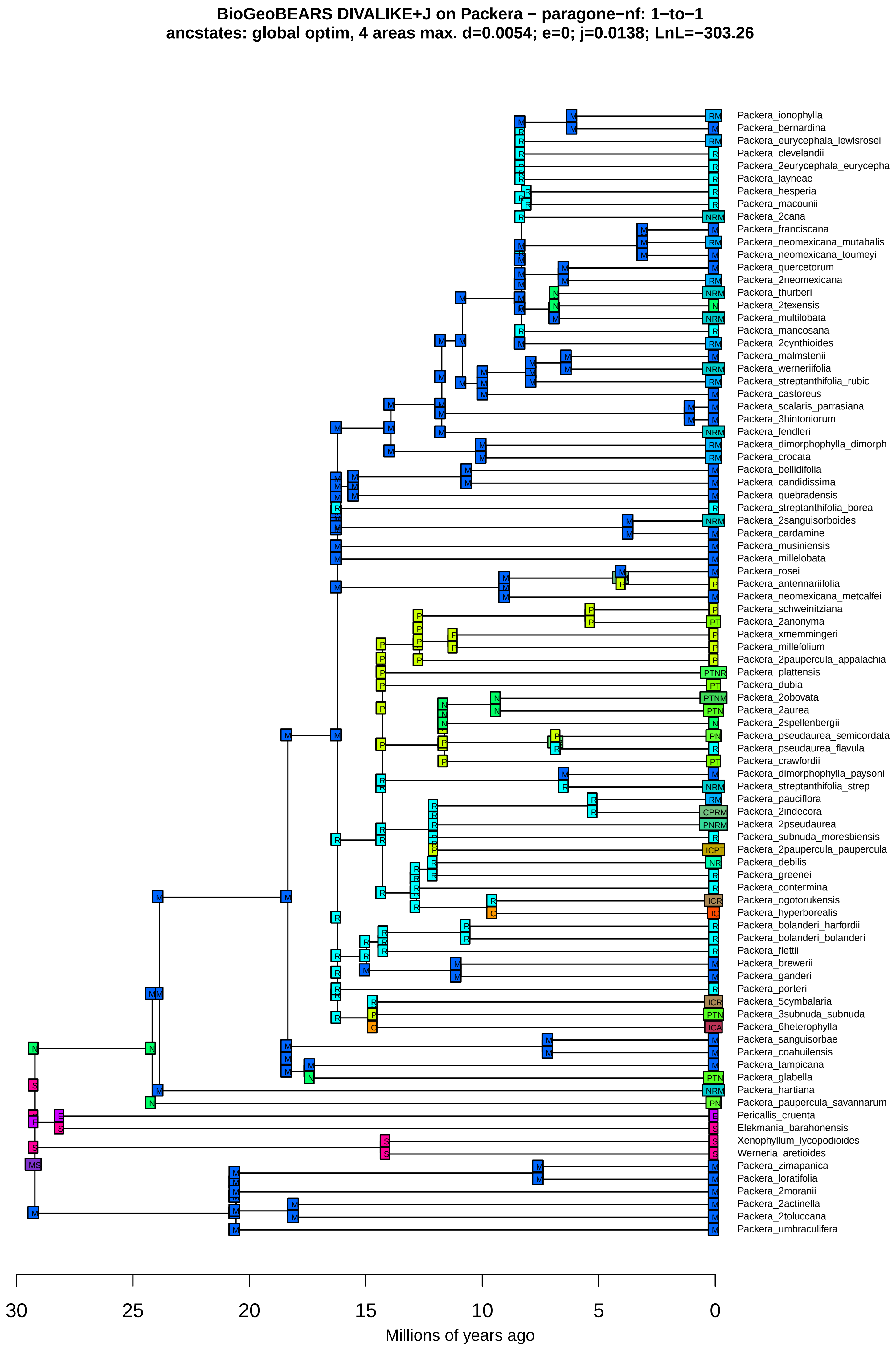
BioGeoBEARS DIVALIKE+J on Packera − paragone−nf: 1−to−1
ancstates: global optim, 4 areas max. d=0.0054; e=0; j=0.0138; LnL=−303.26
Packera_ionophylla
Packera_bernardina
Packera_eurycephala_lewisrosei
Packera_clevelandii
Packera_2eurycephala_eurycepha
Packera_layneae
Packera_hesperia
Packera_macounii
Packera_2cana
Packera_franciscana
Packera_neomexicana_mutabalis
Packera_neomexicana_toumeyi
Packera_quercetorum
Packera_2neomexicana
Packera_thurberi
Packera_2texensis
Packera_multilobata
Packera_mancosana
Packera_2cynthioides
Packera_malmstenii
Packera_werneriifolia
Packera_streptanthifolia_rubic
Packera_castoreus
Packera_scalaris_parrasiana
Packera_3hintoniorum
Packera_fendleri
Packera_dimorphophylla_dimorph
Packera_crocata
Packera_bellidifolia
Packera_candidissima
Packera_quebradensis
Packera_streptanthifolia_borea
Packera_2sanguisorboides
Packera_cardamine
Packera_musiniensis
Packera_millelobata
Packera_rosei
Packera_antennariifolia
Packera_neomexicana_metcalfei
Packera_schweinitziana
Packera_2anonyma
Packera_xmemmingeri
Packera_millefolium
Packera_2paupercula_appalachia
Packera_plattensis
Packera_dubia
Packera_2obovata
Packera_2aurea
Packera_2spellenbergii
Packera_pseudaurea_semicordata
Packera_pseudaurea_flavula
Packera_crawfordii
Packera_dimorphophylla_paysoni
Packera_streptanthifolia_strep
Packera_pauciflora
Packera_2indecora
Packera_2pseudaurea
Packera_subnuda_moresbiensis
Packera_2paupercula_paupercula
Packera_debilis
Packera_greenei
Packera_contermina
Packera_ogotorukensis
Packera_hyperborealis
Packera_bolanderi_harfordii
Packera_bolanderi_bolanderi
Packera_flettii
Packera_brewerii
Packera_ganderi
Packera_porteri
Packera_5cymbalaria
Packera_3subnuda_subnuda
Packera_6heterophylla
Packera_sanguisorbae
Packera_coahuilensis
Packera_tampicana
Packera_glabella
Packera_hartiana
Packera_paupercula_savannarum
Pericallis_cruenta
Elekmania_barahonensis
Xenophyllum_lycopodioides
Werneria_aretioides
Packera_zimapanica
Packera_loratifolia
Packera_2moranii
Packera_2actinella
Packera_2toluccana
Packera_umbraculifera
30
25
20
15
10
5
0
Millions of years ago

**BioGeoBEARS DIVALIKE+J on Packera − paragone−nf: 1−to−1**
**ancstates: global optim, 4 areas max. d=0.0054; e=0; j=0.0138; LnL=−303.26**

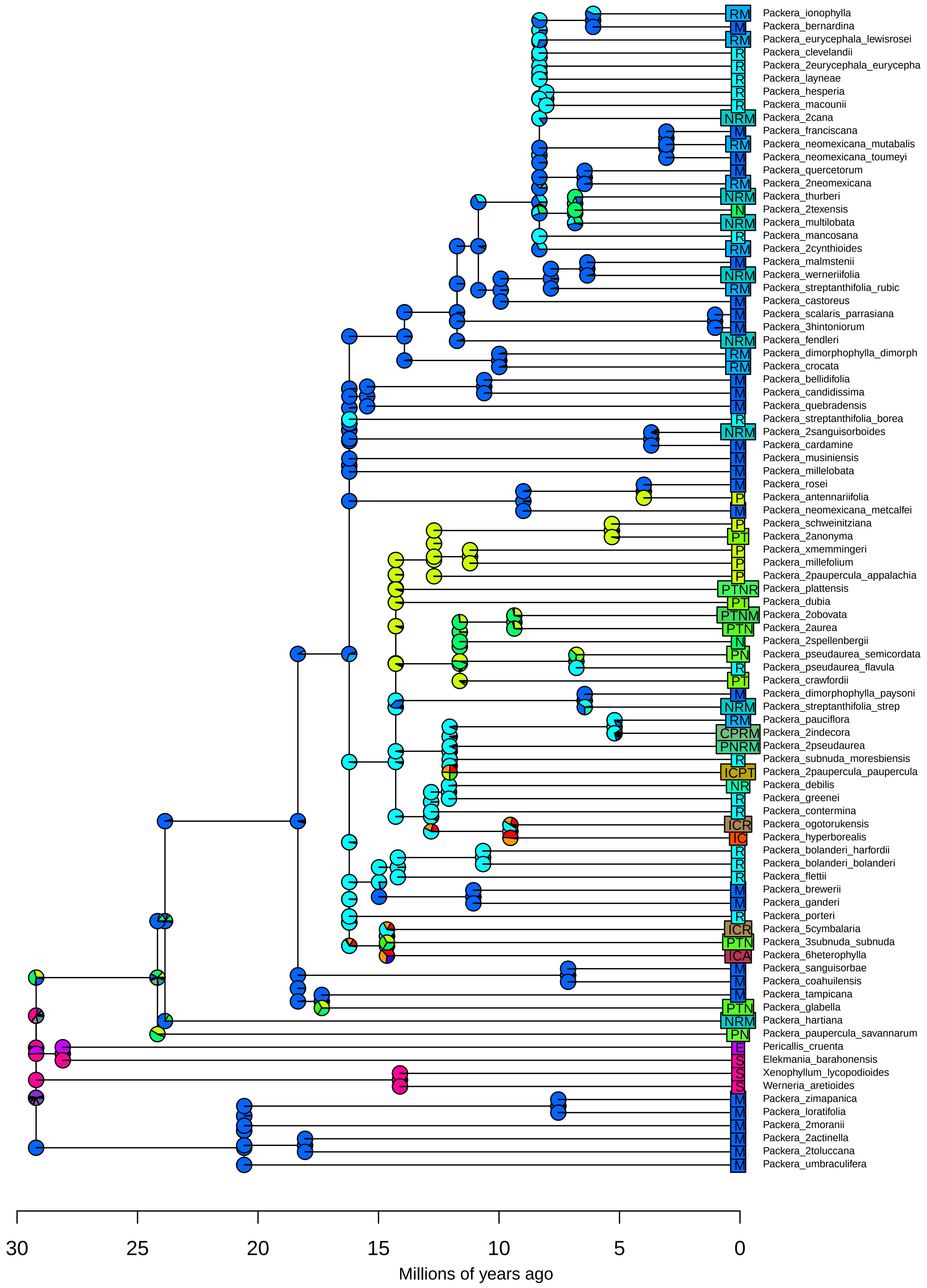

**BioGeoBEARS BAYAREALIKE on Packera − paragone−nf: 1−to−1**
**ancstates: global optim, 4 areas max. d=0.0035; e=0.0461; j=0; LnL=−289.35**

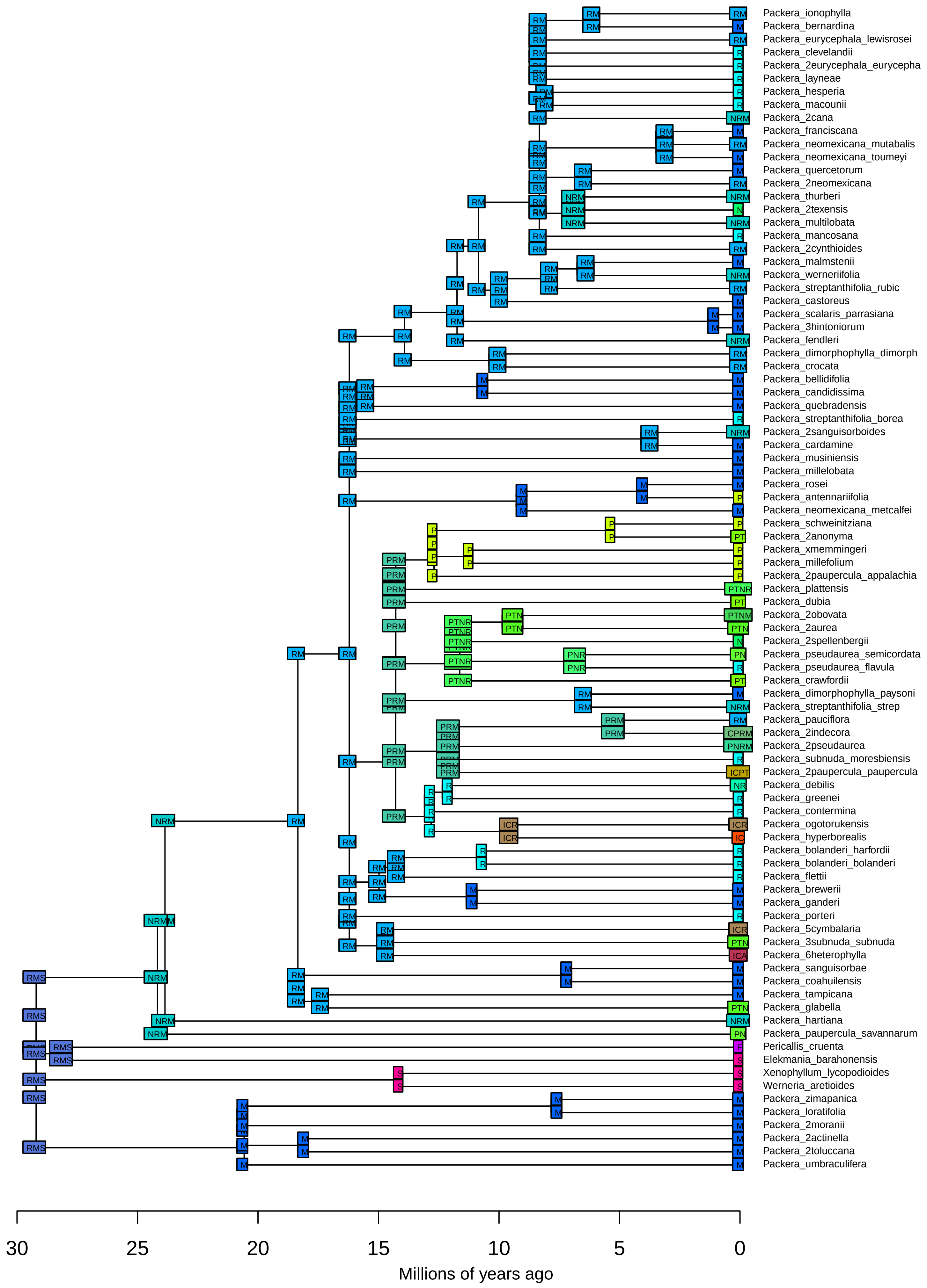

**BioGeoBEARS BAYAREALIKE on Packera − paragone−nf: 1−to−1**
**ancstates: global optim, 4 areas max. d=0.0035; e=0.0461; j=0; LnL=−289.35**

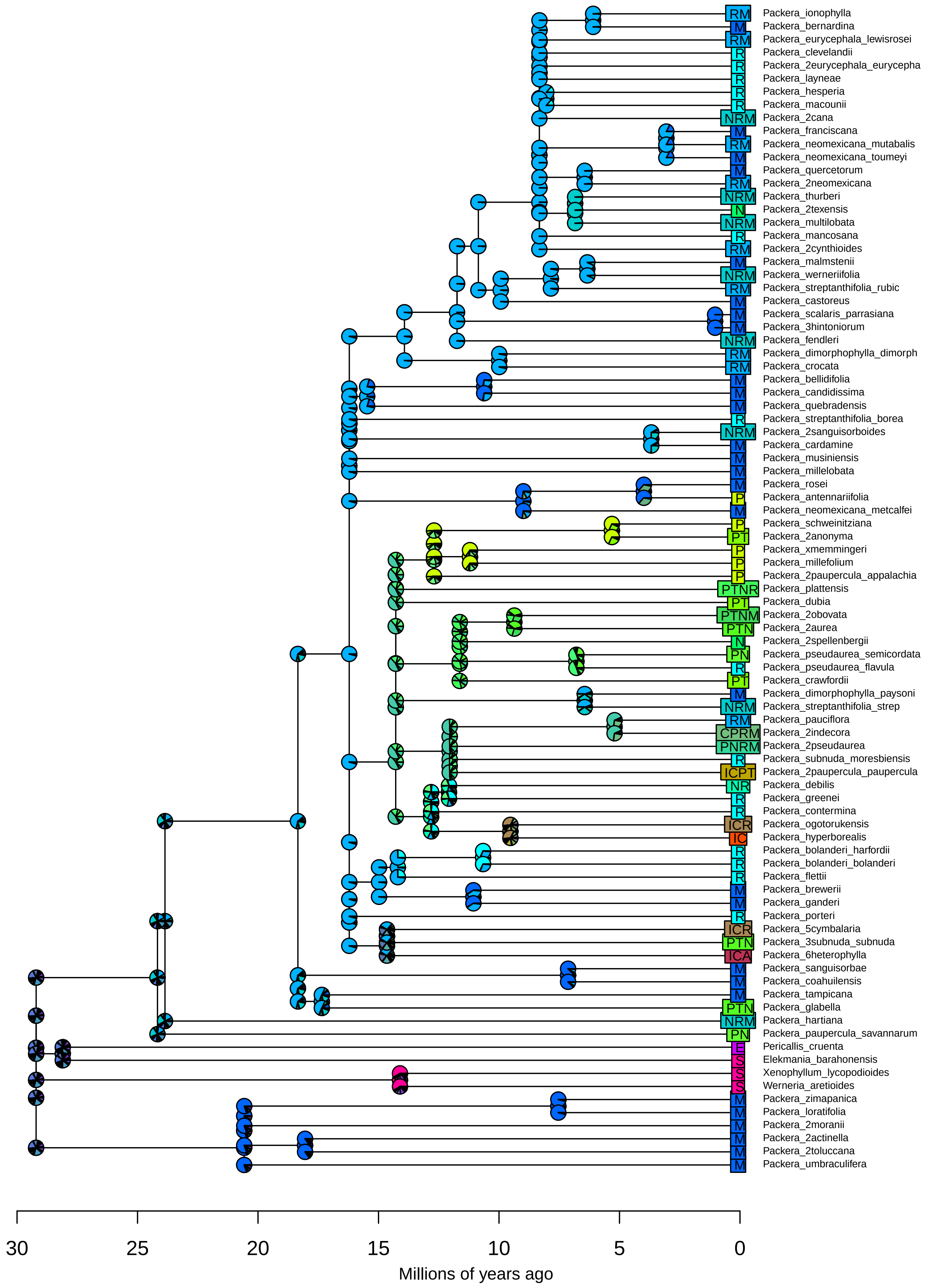

**BioGeoBEARS BAYAREALIKE+J on Packera − paragone−nf: 1−to−1**
**ancstates: global optim, 4 areas max. d=0.0035; e=0.0461; j=1e−04; LnL=−288.90**

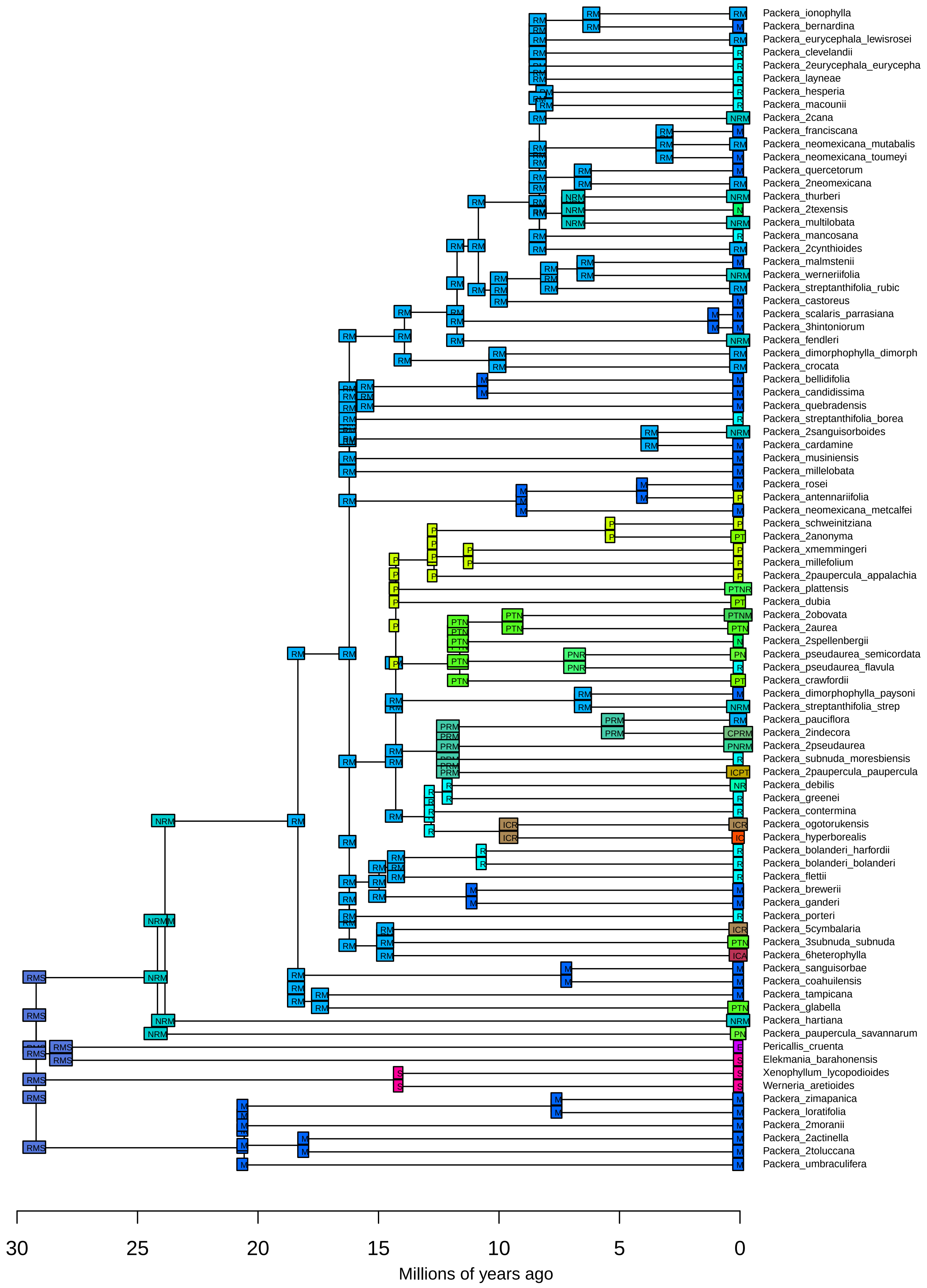

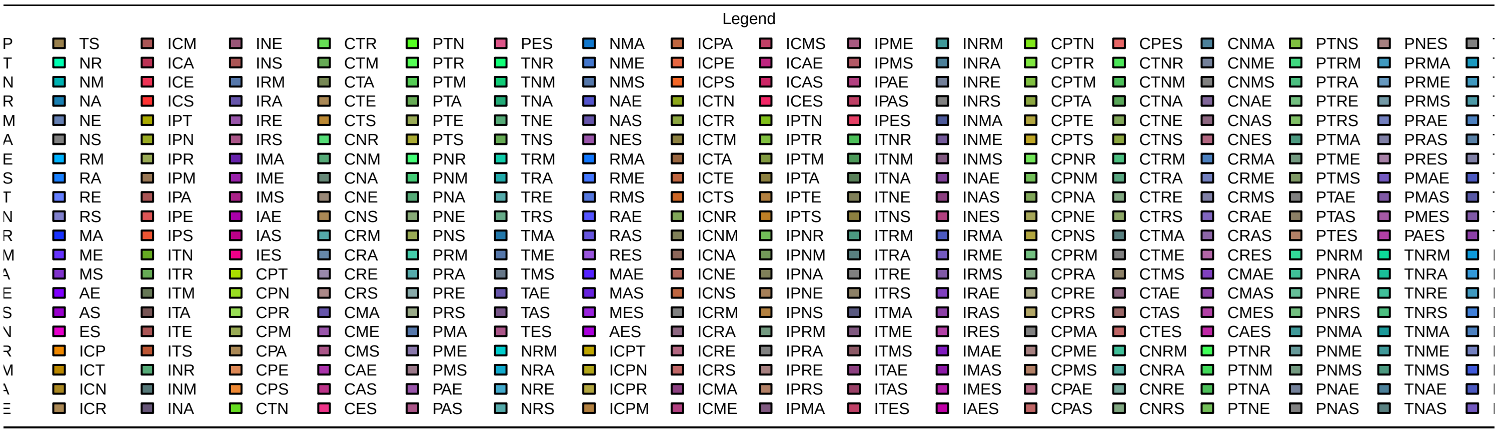
Legend
TS NR NM NA NE NS RM RA RE RS MA ME MS AE AS ES ICP ICT ICN ICR
ICM ICA ICE ICS IPT IPN IPR IPM IPA IPE IPS ITN ITR ITM ITA ITE ITS INR INM INA
INE INS IRM IRA IRE IRS IMA IME IMS IAE IAS IES CPT CPN CPR CPM CPA CPE CPS CTN
CTR CTM CTA CTE CTS CNR CNM CNA CNE CNS CRM CRA CRE CRS CMA CME CMS CAE CAS CES
PTN PTR PTM PTA PTE PTS PNR PNM PNA PNE PNS PRM PRA PRE PRS PMA PME PMS PAE PAS
PES TNR TNM TNA TNE TNS TRM TRA TRE TRS TMA TME TMS TAE TAS TES NRM NRA NRE NRS
NMA NME NMS NAE NAS NES RMA RME RMS RAE RAS RES MAE MAS MES AES ICPT ICPN ICPR ICPM
ICPA ICPE ICPS ICTN ICTR ICTM ICTA ICTE ICTS ICNR ICNM ICNA ICNE ICNS ICRM ICRA ICRE ICRS ICMA ICME
ICMS ICAE ICAS ICES IPTN IPTR IPTM IPTA IPTE IPTS IPNR IPNM IPNA IPNE IPNS IPRM IPRA IPRE IPRS IPMA
IPME IPMS IPAE IPAS IPES ITNR ITNM ITNA ITNE ITNS ITRM ITRA ITRE ITRS ITMA ITME ITMS ITAE ITAS ITES
INRM INRA INRE INRS INMA INME INMS INAE INAS INES IRMA IRME IRMS IRAE IRAS IRES IMAE IMAS IMES IAES
CPTN CPTR CPTM CPTA CPTE CPTS CPNR CPNM CPNA CPNE CPNS CPRM CPRA CPRE CPRS CPMA CPME CPMS CPAE CPAS
CPES CTNR CTNM CTNA CTNE CTNS CTRM CTRA CTRE CTRS CTMA CTME CTMS CTAE CTAS CTES CNRM CNRA CNRE CNRS
CNMA CNME CNMS CNAE CNAS CNES CRMA CRME CRMS CRAE CRAS CRES CMAE CMAS CMES CAES PTNR PTNM PTNA PTNE
PTNS PTRM PTRA PTRE PTRS PTMA PTME PTMS PTAE PTAS PTES PNRM PNRA PNRE PNRS PNMA PNME PNMS PNAE PNAS
PNES PRMA PRME PRMS PRAE PRAS PRES PMAE PMAS PMES PAES TNRM TNRA TNRE TNRS TNMA TNME TNMS TNAE TNAS

**BioGeoBEARS BAYAREALIKE+J on Packera − paragone−nf: 1−to−1**
**ancstates: global optim, 4 areas max. d=0.0035; e=0.0461; j=1e−04; LnL=−288.90**

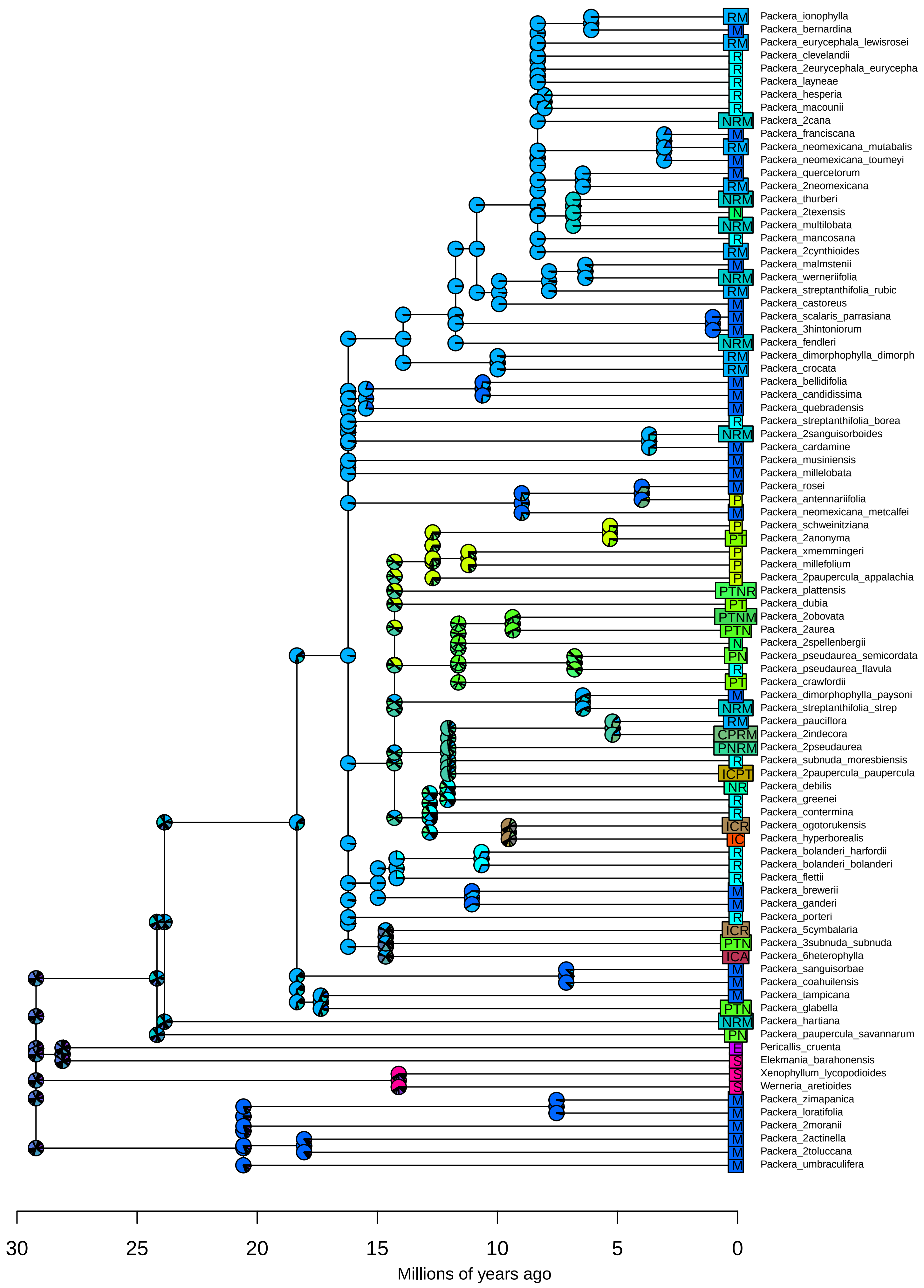
